# Conservation and divergence of regulatory architecture in nitrate-responsive plant gene circuits

**DOI:** 10.1101/2023.07.17.549299

**Authors:** C Bian, GS Demirer, MT Oz, Y Cai, SS Witham, GA Mason, Z Di, F Deligne, P Zhang, R Shen, A Gaudinier, SM Brady, NJ Patron

## Abstract

Nitrogen is an essential element for all life processes in plants. As such, plant roots dynamically respond to nitrogen availability below-ground by executing a signaling and transcriptional cascade resulting in altered plant growth, optimized for nutrient uptake. The NIN-LIKE PROTEIN 7 (NLP7) transcription factor senses nitrogen and along with its closely related paralog NLP6, partially coordinates these transcriptional responses. While post-translational regulation of NLP6/7 is well established, its upstream transcriptional regulation remains understudied in Arabidopsis and other plant species. Here, we dissect a previously identified sub-circuit upstream of NLP6/7 in Arabidopsis and which was predicted to contain multiple multi-node feedforward loops suggestive of an optimized design principle of nitrogen transcriptional regulation. This sub-circuit comprises AUXIN RESPONSE FACTOR 18 (ARF18), ARF9, DEHYDRATION RESPONSE ELEMENT BINDING-PROTEIN 26 (DREB26), A NAC-DOMAIN CONTAINING PROTEIN 32 (ANAC032), NLP6 and NLP7 and their regulation of NITRITE REDUCTASE 1 (NIR1). Conservation and divergence of this circuit and their influence on N-dependent root system architecture are similarly assessed in *Solanum lycopersicum*. The specific binding sites of these factors within their respective promoters and their putative *cis*-regulatory architecture are identified. The direct or indirect nature of these interactions are validated *in planta.* The resulting models were genetically validated in varying concentrations of available nitrate by measuring the transcriptional output of the network revealing rewiring of nitrogen regulation across distinct plant lineages.

**Significance Statement:** Nitrogen is a critical nutrient for plant growth and yield. While external N has facilitated modern agriculture, over-application of N-containing fertilizers has drastic ecological and environmental consequences. Here, we experimentally validate a six gene regulatory circuit with extensive feedforward loops identified to act upstream of the critical *NIN-LIKE PROTEIN 6/7* transcription factors which regulates a nitrogen metabolic enzyme. Our results indicate conservation and divergence in these circuits between Arabidopsis and tomato despite the similar role of NLP7 in N-dependent changes in root system architecture. The resulting network models complement existing knowledge of NLP7 regulation by providing a framework for targeted transcriptional engineering to increase plant nitrogen use efficiency.

## BACKGROUND

Nitrogen (N) is essential for plant growth and basic metabolic processes. The supply of N through the energy-intensive Haber-Bosch process and fertilizer application has been critical to increasing crop yields but also has negative ecological and environmental consequences (1, 2). Knowledge of the diverse regulatory events that crop species employ to respond and adapt to N deficiency can be used to breed plants with increased N use efficiency and ultimately, to lessen reliance on externally applied N. In response to limiting N, a complex series of molecular events are initiated (3–5) including rapid post-transcriptional, calcium- and phosphorylation-dependent signaling cascades that converge on transcriptional regulation of N transporters, assimilation enzymes, N signaling factors, carbon metabolism and hormone pathways (6–11). Below-ground, the outcome of these responses in the root includes altered lateral root elongation to forage for N and concomitant adjustment of N metabolism. Above-ground, changes are manifested in reduced plant growth and a reduction in yield.

Plant responses to nitrate have best been studied in the model dicot *Arabidopsis thaliana* (Arabidopsis). Evidence indicates that regulation of nitrogen- and nitrogen-associated metabolism is both transcriptional and post-translational in nature. Amongst the TFs identified to coordinate transcriptional responses to nitrate (5, 11–17), AtNLP7, and its paralog, AtNLP6 (14) play a key role by modulating the expression of downstream target genes following Ca^2+^-sensor protein kinase (CPK)-induced phosphorylation (8). AtNLP7 also directly binds nitrate (18). While our understanding of the post-translation modes by which NLP6/7 regulate gene expression, their own transcriptional regulation is less understood, but critical to our knowledge of this important factor.

TF networks consist of interconnected transcription factors and their target genes. These biological networks contain several types of regulatory interactions that are significantly over-represented including autoregulatory, feedforward and negative feedback loops <u>(Shen-Orr et al. 2002)</u>. Autoregulatory and feedforward loops are associated with assuring robustness of gene regulation, production of a toggle switch, a pulse of gene expression, and sustained activation of target genes (Figure 1A,B) <u>(Shen-Orr et al. 2002; Atkinson et al. 2003; An Introduction to Systems Biology: D…; Zong et al. 2023; Lee et al. 2002; Milo et al. 2011; Mangan and Alon 2003)</u>. We previously mapped a network of transcription factors that bind to promoters of nitrogen- and nitrogen-associated metabolic genes (Gaudinier et al. 2018). From these, a set of transcription factors, *AtARF9, AtARF18, AtANAC032* and *AtDREB26,* form a circuit consisting of multiple loops that are predicted to regulate expression of *AtNLP6, AtNLP7* and *AtNIR* (Figure 1C). These include auto-regulation (*AtDREB26*) and feedforward loops, where three genes interact in both a direct and indirect path (*AtARF9/AtDREB26/AtANAC032; AtARF18/AtDREB26/AtANAC032; AtARF9/AtNLP6/AtNIR; AtDREB26/AtANAC032/AtNLP7*). Given these network motifs, we hypothesized that these genes are critical to transcriptional regulation of nitrogen metabolic gene expression.

**Figure 1.**
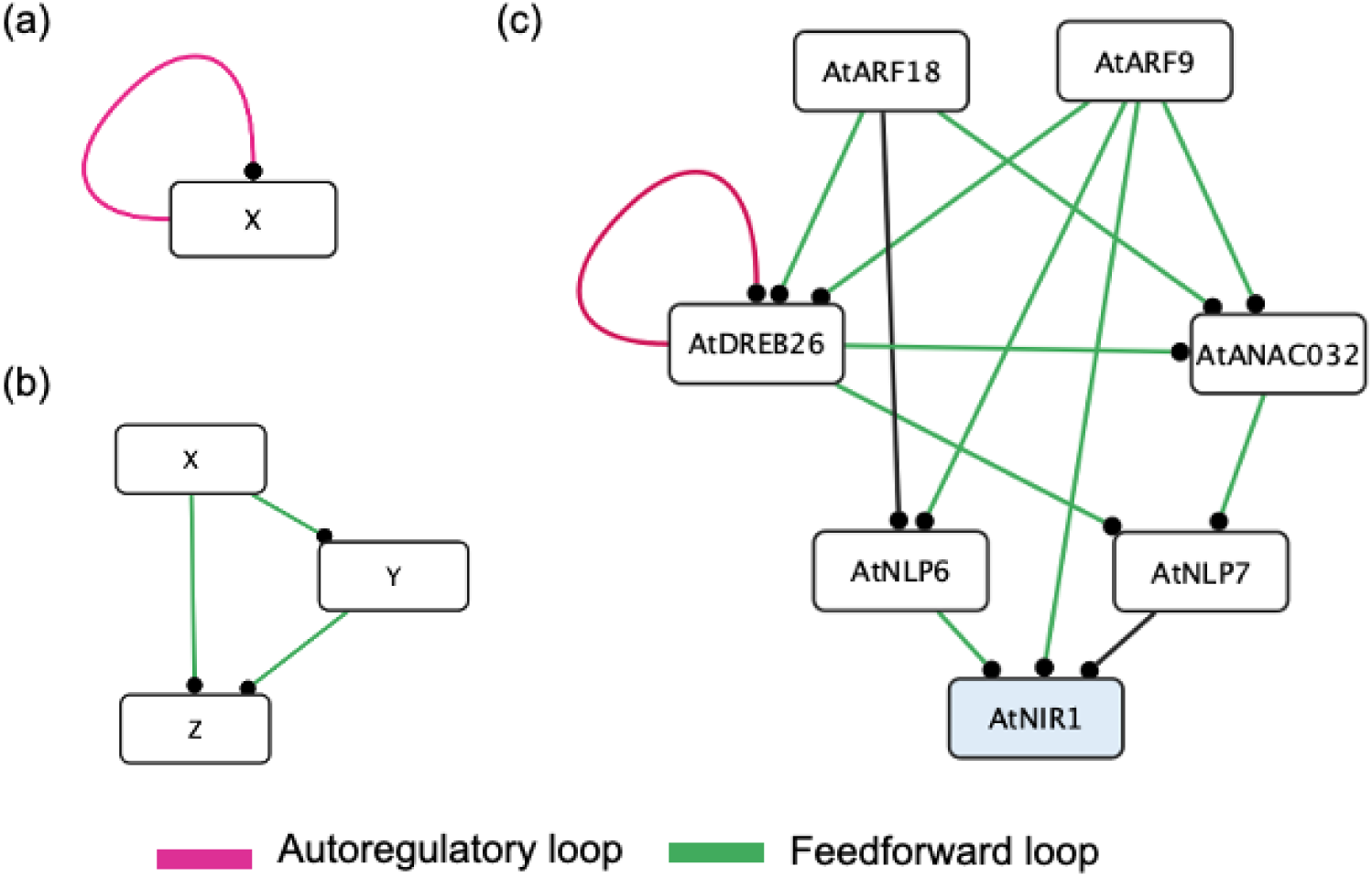
Nitrogen metabolism network sub-circuit in Arabidopsis that acts upstream of NLP6, NLP7 and NIR1 as determined by enhanced yeast one hybrid assays. (a) Autoregulatory loop. (b) Feedforward loop. (c) This gene sub-circuit contains multiple interconnected feedforward loops (green edges) and an autoregulatory interaction (pink edges). *AtNIR1* is indicated in blue as the representative metabolic target gene.

Interpreting the influence of these transcription factors and their regulation of each other as well as on the transcriptional regulation of N metabolism via Arabidopsis *NITRITE REDUCTASE 1* (*AtNIR1),* is non-trivial and depends on the sign of each regulatory interaction (activating or repressing) and the extent to which the gene is regulated by its upstream factors. Furthermore, to deploy the knowledge gained in Arabidopsis to the rational engineering of crop species, it is also necessary to obtain evidence of which aspects, if any, of the network are conserved. While NLP7 is conserved in some crop species (19, 20), we lack evidence for other TFs and the interconnected manner by which they genetically determine nitrate-responses. To date, few molecular players associated with N signaling and metabolism in tomato have been identified (21, 22). Although there is conservation of radial patterning between Arabidopsis and tomato, there are also key differences. Tomato has three cortex layers compared to the one in Arabidopsis, the outermost of which forms a barrier composed of lignin and suberin (23). A (24) predicted gene regulatory network for the exodermis demonstrated exodermal-enrichment of N metabolic genes with *SlNLP7* as a central regulatory factor. These differences between tomato and Arabidopsis further demonstrate the potential for changes in the underlying regulatory circuitry that controls responses to nitrate.

Transcriptional coordination of nitrogen metabolic gene expression occurs by multiple events, many of which have been mapped via genome-scale assays. These include chromatin accessibility assays <u>(Maher et al. 2018; Sullivan et al. 2014)</u>, *in vivo* binding of individual TFs by chromatin immunopurification coupled with RNA sequencing (Marchive et al. 2013), and transcriptional regulation by RNA sequencing of mutants or by Transient Transformation System for Genome-Wide Transcription Factor Target Discovery (TARGET) using individual TFs (Alvarez et al. 2020; Bargmann et al. 2013; Para et al. 2014; Brooks et al. 2019). Genetic elaboration of these interconnected regulatory interactions is laborious. Systems biology approaches that include gene editing and quantitative protoplast-based assays enable the iterative testing of regulatory interactions between multiple transcription factors and their target regulatory regions *in vivo*. To date, NLP7 has been identified to bind to nitrate responsive *cis* elements in the promoters of nitrate-responsive target genes including *NIR1* (Konishi and Yanagisawa 2010). In an effort to better elucidate the genetic architecture of these six genes predicted to act upstream of AtNLP6/7 with extensive feedforward loops (Figure 1C), we systematically identified and validated specific binding motifs for TFs and studied their *in vivo* DNA binding and regulation (17). Regions of open chromatin associated with each gene were mined for DNA binding motifs and the ability of TFs to bind to these targets were subsequently determined using an *in vitro* methodology (25). *In vivo* TF binding and regulation are confirmed using a modification of the TARGET assay (26) and protoplast co-expression. In parallel, conservation of this network in tomato was tested using phylogenomics coupled with the modified TARGET system. ARF18/ARF9 and NLP6/7 function at the top and bottom of this circuit, respectively, in both Arabidopsis and tomato. However, in between these factors, gene regulatory interactions differ between species. This network model was iteratively tested and validated using single, pairwise, and higher order genetic perturbations of all factors in a novel *in vivo* assay in multiple concentrations of available nitrogen. Collectively, this systems level approach illustrates the conservation and divergence of transcriptional regulation of N metabolism in two dicot species. Further, it identifies numerous sequences to target for future engineering of nitrate metabolism. Few plant regulatory networks have been genetically characterized to this extent. Our work provides a blueprint of how systems biology techniques can be used to inform synthetic biology approaches to crop engineering by predicting the outcomes of perturbations.

## RESULTS

### *In vitro* characterization of candidate TF binding sites supports interactions within a putative transcriptional network regulating Arabidopsis response to nitrate

In previous work, enhanced Yeast-1-Hybrid (eY1H) data was used to infer a putative transcriptional network that regulates the architecture of Arabidopsis root systems in response to N availability (17) (Figure 1). The yeast network contained many interconnected transcription factor-promoter interactions and was enriched for nitrate-responsive gene expression (Gaudinier et al. 2018). Genetic perturbation and mutant phenotyping of TFs predicted to be key in this network based on their connectivity properties showed that these genes regulate multiple aspects of root and shoot architecture. Understanding the cis-regulatory logic of these factors and how they work together in pathways is an important next step to understand how Arabidopsis coordinates the transcriptional response to available nitrogen. While Y1H is a powerful technique for identifying previously unknown interactions between TFs and target genes, false positives can result from transcription initiation by endogenous yeast TFs. Similarly, false negatives may result from improper folding, mislocalization, or because required post-translational modifications or protein interactions do not occur (27). In addition, though Y1H can identify candidate target genes, it is unable to identify the *cis*-regulatory motifs responsible for TF binding and cannot inform if interactions are likely to result in changes to the expression of target genes. This information is critical for understanding how regulatory networks control quantitative phenotypes, and for predicting the effects of perturbations. To address this, we first looked for evidence of TF-binding by identifying candidate binding sites in the regulatory regions of target genes.

To identify candidate TF binding sites, we performed a systematic analysis of the open chromatin sites upstream and downstream of each gene in the regulatory circuit. Regions for analysis were restricted to two kilobases upstream of the start of transcription, or until the next protein-coding gene. In addition, we analyzed one kilobase downstream of the start of transcription. Peaks were mapped from publicly available ATAC-seq data (28)(Figure 2a). Candidate binding sites for AtNLP7 and AtDREB26 were identified using the FIMO package within the MEME software suite (29) using publicly available data to infer position weight matrices (PWMs) (30) (Supplementary Data 1). Equivalent DNA binding data for AtNLP6, AtARF9, AtARF18 or AtANAC032 was unavailable and PWMs from a closely related TF, for which the DNA-binding domain was confirmed (Supplementary Data 2), were used to identify candidate sites (see Methods). This analysis identified candidate binding motifs that supported interactions within the putative Y1H regulatory circuit and identified the presence of further putative interactions (Figure 2a).

**Figure 2.**
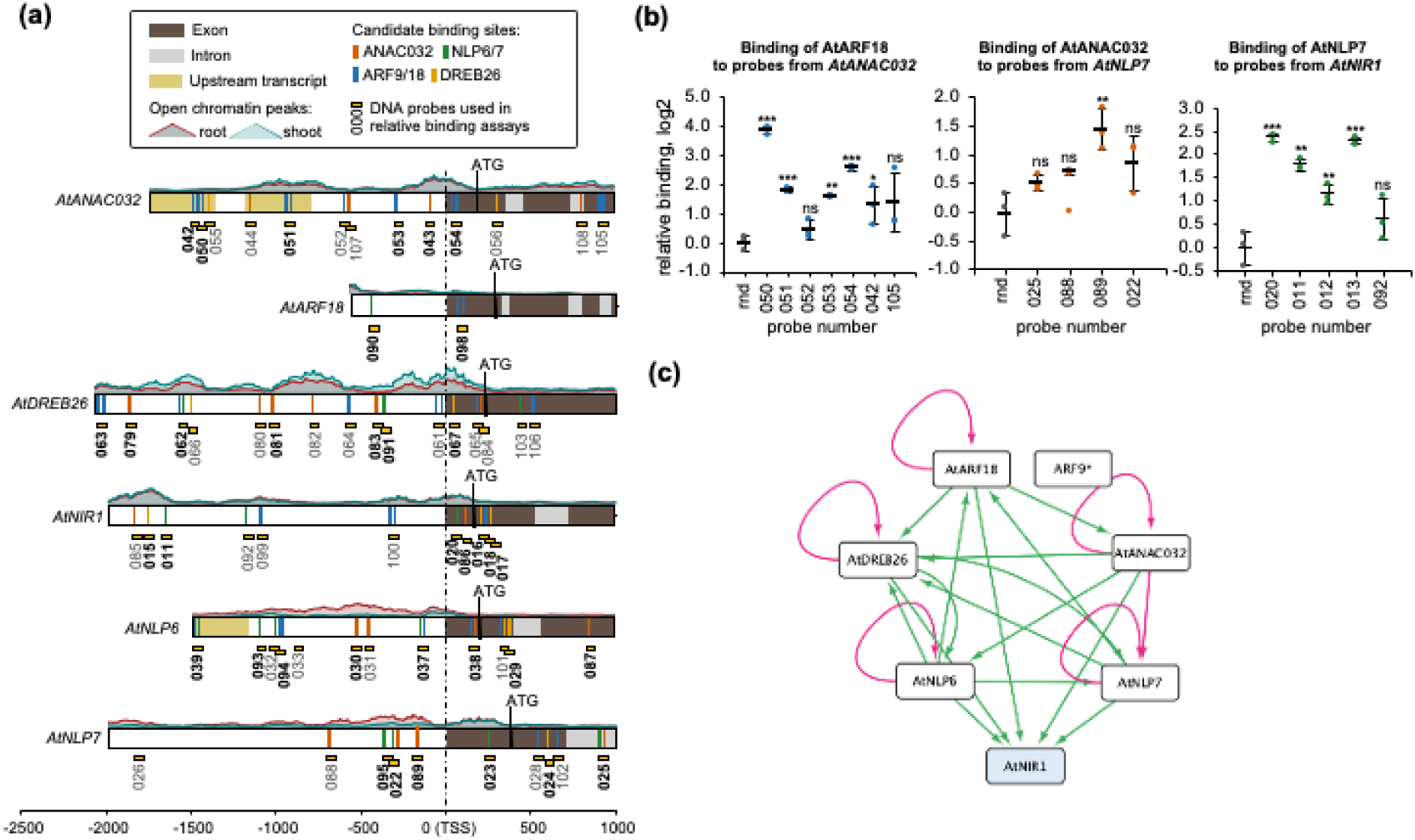
Transcription factor binding motifs and *in vitro* assays support and elaborate the sub-network. **(a)** Position weight matrices (PWMs) were used to identify candidate transcription factor binding sites in the 5’ regions of *AtANAC032*, *AtANR1, AtARF18, AtDREB26, AtNIR1, AtNLP6* and *AtNLP7* transcripts. Binding of TFs to short DNA probes containing candidate sites (numbered boxes) was evaluated using an *in vitro* assay; bold text indicates that significant binding of at least one transcription factor was detected (Supplementary Data 3) **(b)** Representative *in vitro* binding assays showing interactions between AtARF18, AtANAC032 and AtNLP7 and probes from *AtANAC032, AtNLP7 and AtNIR1,* respectively. Error bars = mean and standard deviation; n=3; *p*-values were calculated using an unpaired two-tailed Student’s t-test of each sample to the random (rnd) control probe; \**p*<0.05, ** *p*<0.01, *** *p*<0.001; ns = not significant **(c)** Protein-DNA interactions as determined by the *in vitro* assay. Edges that are part of a feedforward loop are indicated in green, and autoregulatory interactions in pink. AtNIR1 is indicated in blue as the representative metabolic target gene.

To validate the potential for TF-DNA binding, the ability of each TF to bind to each candidate binding site was experimentally determined using a plate-base method for specific DNA sequences (Cai et al. 2023). To do this, we expressed recombinant proteins of AtARF18, AtANAC032, AtNLP6, AtANLP7 and AtDREB26 with a genetically encoded N-terminal 9xHIS tag and a C-terminal HiBit tag. We were unable to purify sufficient soluble protein of AtARF9. We then tested the ability of purified protein to bind to 80 bp probes containing candidate binding sites, relative to random DNA probes. Example data supporting three edges within the network are shown in (Figure 2b). Data for all interactions is provided in Supplementary Data 3 and the sequences of all probes are provided in Supplementary Data 4.

From these data, we constructed an *in vitro* N network (Figure 2c) and identified several previously known interactions including between AtNLP6/ AtNLP7 and the Nitrogen Responsive Element (NRE) present in the promoter region of *AtNIR1* (31), as well as AtNLP7 with itself (32). These data also support four additional edges previously identified by Y1H: AtARF18 to *AtANAC032* and *AtDREB26*; AtANAC032 to *AtNLP7*; and AtDREB26 to *AtNIR1*. In addition, 12 further potential interactions were identified: AtNLP6 and AtNLP7 to *AtARF18* and *AtDREB26*; AtANAC032 to *AtDREB26;* AtDREB26, AtARF18 and AtANAC032 to *AtNIR1*; and AtDREB26 and AtANAC032 to *AtNLP6.* Some of these interactions could not have been identified by the earlier Y1H assay, as the motifs occur beyond the borders of the region used for eY1H assays. Finally, three instances of potential autoregulation were identified for AtANAC032, AtARF18, AtDREB26 and AtNLP6. Interestingly, AtNLP6, AtNLP7, AtDREB26, and AtANAC032 were all observed to bind significantly to both *AtNLP6* and *AtNLP7*. While these data indicate the potential for protein-DNA interactions, *in vitro* binding does not provide evidence that these 28 TF-DNA interactions occur *in planta* or are regulatory in nature.

### Refining the network by protoplast expression assays to determine the regulatory consequences of TF binding

To investigate if the TFs were able to physically bind to and modulate the expression of predicted targets *in vivo*, we used a variation of the Transient Transformation System for Genome-Wide Transcription Factor Target Discovery (TARGET) assay, in which glucocorticoid receptor (GR)-tagged TFs are expressed in protoplasts (26). This assay quantifies the expression of endogenous target genes following the treatment of cells with dexamethasone (DEX). Cycloheximide (CHX) in the presence of DEX determines if regulation is direct and regulatory. In TARGET, transcriptional changes were quantified using RNA-seq. However, in this variation, changes to the expression of specific target genes were quantified using qPCR. We additionally investigated predicted direct interactions using a ratiometric transactivation luciferase (LUC) assay in which each target promoter was fused to a nanoluciferase reporter (LucN) and coexpressed in Arabidopsis protoplasts with either a constitutively expressed TF or constitutively expressed control protein (YFP) (Supplementary Data 5).

Starting at the top of the predicted sub-network and working down, AtARF18 was found to directly repress *AtANAC032;* ANAC032 to directly repress *AtNLP7* and AtNLP7 to directly activate *AtNIR1* (Figure 3a-f; Supplementary Data 6). The expression of *AtDREB26* could not be monitored in a TARGET assay as CHX represses its expression (33). However, in the transactivation LUC assay, *AtDREB26* expression was activated by AtNLP7 and AtNLP6 and repressed by AtARF18 (Supplementary Data 7). Despite the identification of binding sites for AtDREB26 in the regulatory regions of *AtNIR1* and in the coding sequence of *AtNLP7* (Supplementary Data 3), we did not identify any instances of direct regulation by AtDREB26 in the TARGET assay. Finally, while *in vitro* binding to their own promoters was detected for AtNLP6 and AtNLP7 (Supplementary Data 3) direct autoregulation was not detected. The resulting regulatory network contains a coherent feedforward loop regulation of *AtNIR1* by AtANAC032 via AtNLP7.

**Figure 3.**
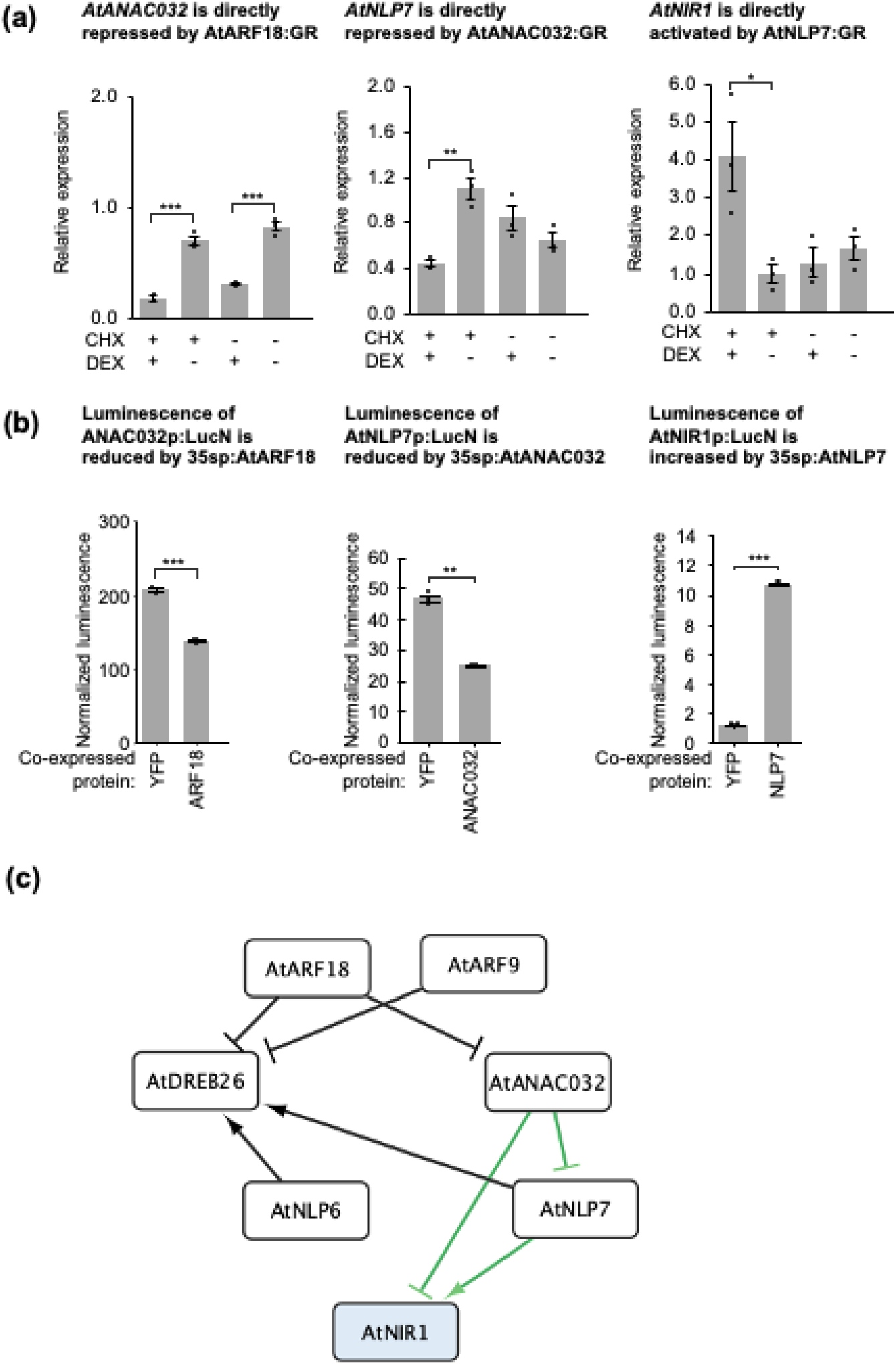
Examples of regulatory consequences of transcription factor expression in protoplasts. (**a**) Changes in endogenous expression of target genes in the modified TARGET assays. Values represent the mean and standard error of three biological replicates of which each is the mean of two technical replicates. CHX = cycloheximide; DEX = dexamethasone. (**b**) Changes in normalized luminescence of nanoluciferase (LucN) in response to transcription factor overexpression. Values represent the mean and standard error of three biological replicates. P-values were calculated using an unpaired two-tailed Student’s t-test *P<0.05, ** P<0.01, *** P<0.001 **(c)** Schematic of sub-network illustrating the regulatory consequences (modified TARGET and/or co-expression data) of transcription factors that directly bind their target genes. Edges that are part of a feedforward loop are indicated in green. *AtNIR1* is indicated in blue as the representative metabolic target gene.

### Testing conservation and divergence of nitrate transcriptional and developmental regulation in tomato

A conserved role for homologues of *AtNLP7* in nitrate responsiveness has been demonstrated in several species including rice (20), and maize (19). We therefore investigated if this regulatory network is conserved in another dicot lineage. We selected tomato as one of the most important horticultural crops, with a preference for nitrate as an inorganic N source (35). While tomato homologs of *AtNLP*7 have been identified (21), the molecular mechanisms of nitrate uptake and assimilation are poorly understood in this species. Further, recent data has demonstrated that some aspects of root development and gene regulation are not conserved between Arabidopsis and tomato (23).

We first confirmed that there is conservation of root system architecture and transcriptional regulatory responses to nitrate. To do this, we germinated and grew seeds for ten days on media supplemented with 0 to 10 mM KNO_3_. We observed that lateral root length and total root size increases from 0 mM to 1 mM with a decrease from 1 mM to 10 mM KNO_3_. However, this is non-linear, with growth decreasing at higher concentrations (Figure 4a-e). This is consistent with the scavenging behavior observed in Arabidopsis (36). We next tested the transcriptional influence of diverse concentrations of nitrogen. Samples were taken from roots grown in media supplemented with 0 mM, 1 mM or 10 mM KNO_3_ and RNA was extracted. RNAseq analysis revealed that canonical N regulatory genes increased in expression dependent on nitrate availability, including high-affinity nitrate transporters, low-affinity nitrate transporters, nitrate assimilation enzymes such as nitrate reductase and nitrite reductase, and LOB domain-containing proteins (Figure 4c). Analysis of GO category enrichment indicates a likely conservation of downstream biological processes including N and nitrate assimilation, glutamine and glutamate biosynthesis (Figure 4d and Supplementary Data 8). Many of these nitrate-responsive genes were also significantly differentially expressed in response to nitrate in other studies (e.g. *SlNRT1.1a, Solyc08g078950; SlNRT1.1b, Solyc08g007430; SlNIA1, Solyc11g013810; SlNRT2.2a, Solyc11g069750; SlNIR1, Solyc01g108630; SlNIR2, Solyc10g050890; SlLBD38, Solyc01g107190*) supporting their central nature in the nitrogen transcriptional response <u>(Wang et al. 2001; Julian 2022; Sunseri et al. 2023)</u>.

**Figure 4.**
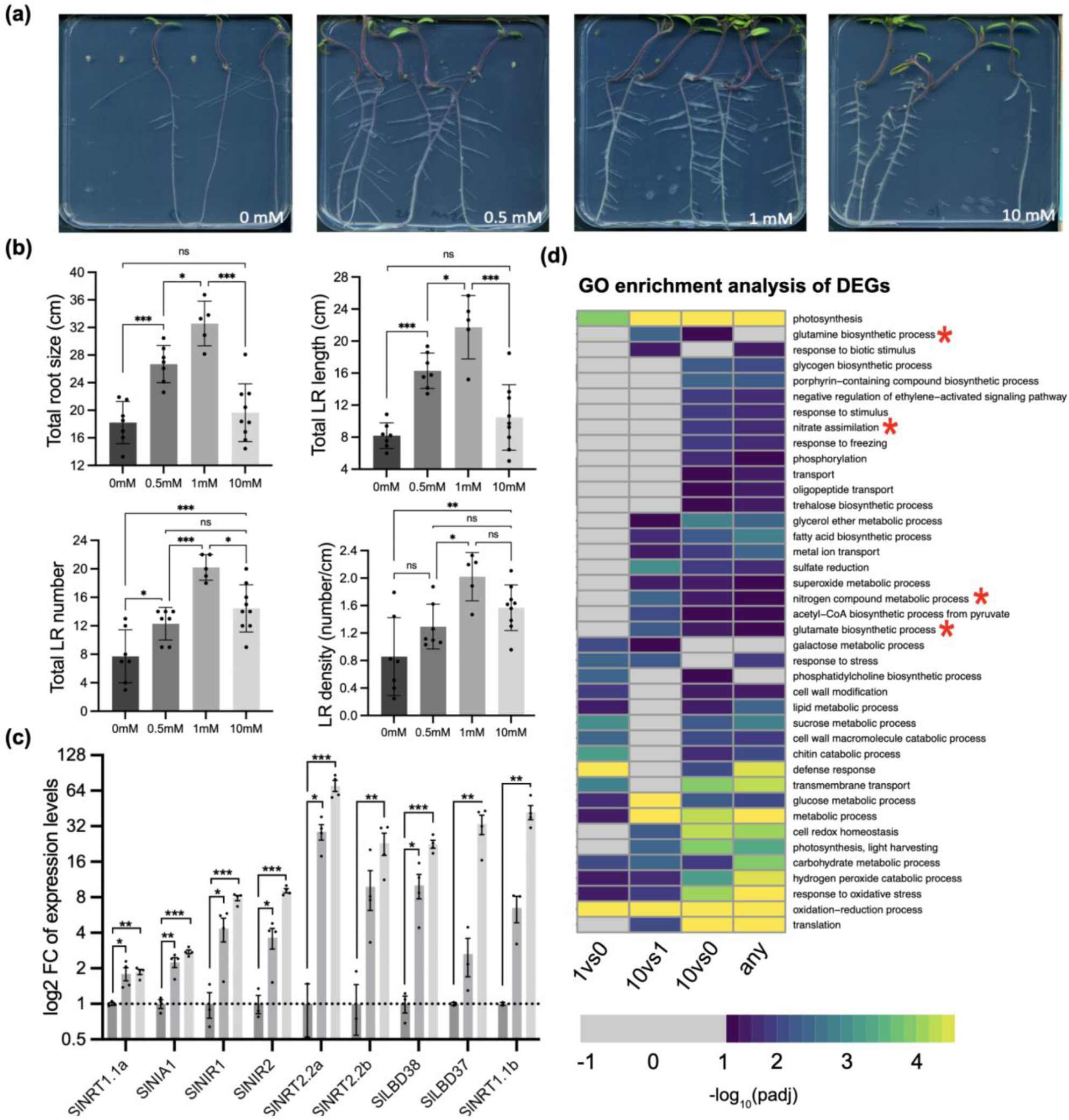
Root growth and gene expression response of tomato (M82) seedlings to potassium nitrate KNO3. **(a)** Exemplar images of root system architecture (RSA) in increasing nitrate. Plants were imaged 10 days after germination on MS media (no nitrogen) supplemented with 0, 0.5, 1, and 10 mM KNO3. **(b)** Quantification of root system architectures including total lateral root length, total root size (sum of main and lateral root lengths, total lateral root number and lateral root density (ratio of lateral root number to main root length from M82 seedlings 10 DAG grown on 0, 0.5, 1, and 10 mM KNO3 (n=7). Statistical analysis was performed using one-way ANOVA with Tukey post-hoc test (adjusted p-value <0.05). **(c)** Log2 fold-changes of nitrogen-responsive genes. Asterisks indicate differentially expressed between N conditions by limma (false discovery rate <0.05). *** p-value < 0.001, ** p-value < 0.01,* p-value < 0.05. **(d)** GO enrichment analysis of DEGs between nitrate conditions in M82 RNA-seq experiment. Canonical nitrogen-mediated Gene Ontology terms are marked with asterisks.

### What genes control the tomato nitrogen response?

Given the conservation in N-mediated changes in root system architecture and transcription, we hypothesized that the underlying regulatory network is likely conserved between Arabidopsis and tomato. Using the elaborated network from Arabidopsis (Figure 3c) as a base, we investigated if the tomato genome encoded likely orthologs of *AtARF18, AtARF9, AtDREB26, AtANAC032, AtNLP6, AtNLP7* and *AtNIR1.* As expected, given the history of tomato genome triplication (37), duplicates or triplicates of many of these Arabidopsis genes were identified. Putative orthologs were first identified using phylogenetic trees (23) (Supplementary Data 9), and this gene set was refined by identifying the homolog(s) with expression in the tomato root (23). A clear likely ortholog was identified for *AtARF18* (*Solyc01g096070*), two potential *AtARF9* paralogs (*Solyc08g008380* and *Solyc08g082630*), a single *AtDREB26* ortholog (*Solyc11g012980*), two genes corresponding to either *AtNLP6* or *AtNLP7* (*Solyc08g008410* and *Solyc08g082750*) and paralogs of *AtNIR1* whose expression was confirmed to be N concentration-dependent (*Solyc10g050890* and *Solyc01g108630,* Figure 4c, Supplementary Data 9, and Supplementary Data 16). *Solyc08g008380* was expressed in the tomato root, while Solyc08g082630 was not expressed in the root (Supplementary Data 9b). Therefore, *Solyc08g008380* was chosen for further analysis and was called *SlARF9B,* consistent with naming in (38). No clear *AtANAC032* ortholog was identifiable given the phylogeny. The first NLP gene, *Solyc08g008410* was found to be mis-annotated and to contain two NLP duplicates (Supplementary Data 10), the latter of which is extremely lowly expressed. The *NLP* genes, both of which contain RWP-RK nitrate sensing domains, were named *SlNLP7A* (*Solyc08g008410*) and *SlNLP7B* (*Solyc08g082750*), as it was impossible to differentiate which could be the correct ortholog for *AtNLP6* or *AtNLP7* (Supplementary Data 10).

Given the conservation of the wild type tomato root N response relative to Arabidopsis, and the presence of an additional gene within the circuit in Arabidopsis compared to tomato (AtANAC032), we next tested the degree to which they are genetically conserved. We therefore monitored phenotypes of mutated genes at the top (*AtARF18* and *SlARF18)* and bottom (*AtNLP7* and *SlNLP7A/B)* of the circuit, as well as of *ANAC032* which is present only in Arabidopsis (17). Root system architecture traits (primary root length, number of lateral roots, total lateral root length, average lateral root length, total root length, lateral root density and ratio of lateral root length to total root length) were measured in limiting (1 mM KNO_3_ for Arabidopsis, 0 and 1 mM KNO_3_ for tomato) and sufficient (10 mM KNO_3_) for mutant alleles of these genes in Arabidopsis and tomato. This included two previously described mutant alleles of Arabidopsis *ARF18*, *atarf18-2* and *atarf18-3* (17), new CRISPR-Cas9 generated alleles of *ARF18* (*atarf18-cc-1*) and *ANAC032* (*atnac032-cc-s-1*) (Supplementary Data 11 and 12). In tomato, guide RNAs were designed to target *SlARF18* and *SlARF9b* to reduce redundancy (Supplementary Data 12); and to *SlNLP7b*. In the Arabidopsis and tomato lines containing mutations of *ARF18* (*atarf18-2* and *arf18-3*) and *ANAC032* (*atnac032-1* and *atnac032-cc-s-1*), a smaller root system was observed relative to wild type in at least one concentration of KNO_3_ for any traits (Figure 5 and Supplementary Data 13, 14). In contrast with the Arabidopsis *ARF18* mutant alleles, *slarf18;9b-1* had a larger root system architecture. This larger root system architecture was also observed in Arabidopsis and mutant alleles of NLP7 (Figure 5, Supplementary Data 13, 14). Given these similarities and differences, we next looked at the degree of conservation of the network at molecular resolution.

**Figure 5.**
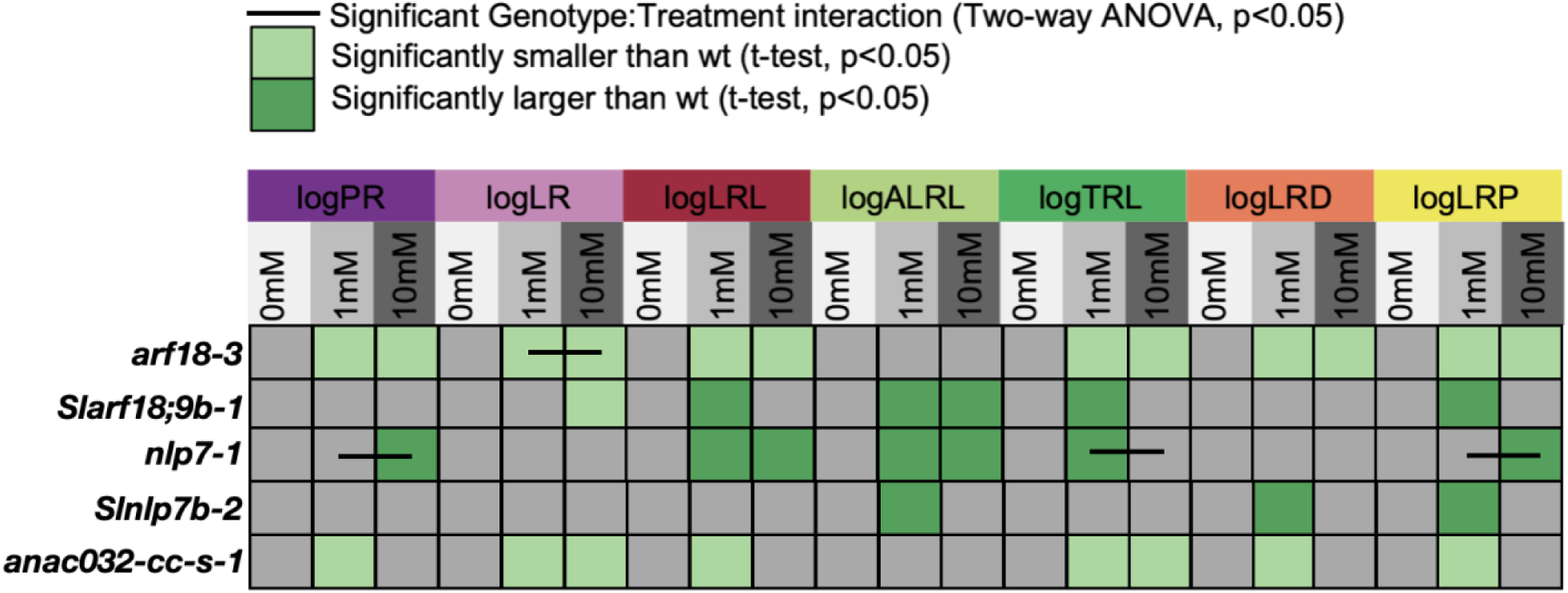
Tomato and Arabidopsis root system architecture traits in orthologous mutant backgrounds. Traits measured include the natural logarithm-transformed primary root length (logPR), number of lateral roots (logLR), total lateral root length (logLRL), average lateral root length (logALRL), total root length (logTRL), lateral root density (logLRD) and the ratio of lateral root length to total root length (logLRP). Traits were measured in 0, 1 and 10 mM KNO_3_ for tomato, and 1 and 10 mM KNO_3_ for Arabidopsis. Light green represents the trait as significantly smaller relative to wild type (wt), dark green represents the trait as significantly larger relative to wild type, white indicates no significant difference and grey indicates that this phenotype was not tested at this concentration (t-test, p<.05). A black bar indicates a significant genotype x treatment interaction as measured by a two-way ANOVA (p<.05).

### *In vitro* characterization of tomato *cis*-regulatory architecture

To investigate conservation of the network, we first investigated the upstream regulatory regions of *SlNIR1, SlNIR2, SLNLP7A, SLNLP7B, SLDREB26* to determine if there were candidate binding sites for ARF18/9; DREB26 and NLP6/7 TFs by using FIMO with publicly available PWM data (29, 30) (Figure 6a). There is extensive conservation of DNA binding domains between species (30, 39), and we tested if this was the case for the DNA-binding domains of the Arabidopsis and tomato TFs within the network. Indeed, extensive conservation between Arabidopsis and tomato was observed for the ARF, DREB and NLP factors (Supplementary Data 15). As a proof-of-principle, we tested if Arabidopsis NLP7 and ARF18 transcription factors were able to bind *in vitro* to candidate sites identified in the *SlNLP7A, SlNLP7B, SlNIR1* and *SlNIR2* promoters (Figure 6b). Interactions between AtARF18 and *SlDREB26* as well as *SlNIR1* were conserved, as were interactions between AtNLP7 and *SlNIR1/SlNIR2*. In contrast, AtARF18 was able to bind to the promoters of SlNLP7A and SlNLP7B – in Arabidopsis, interactions between AtARF18 and *AtNLP6*; as well as AtARF18 with *AtNLP7* were not observed and represent divergence between Arabidopsis and tomato (Supplementary Data 3).

**Figure 6.**
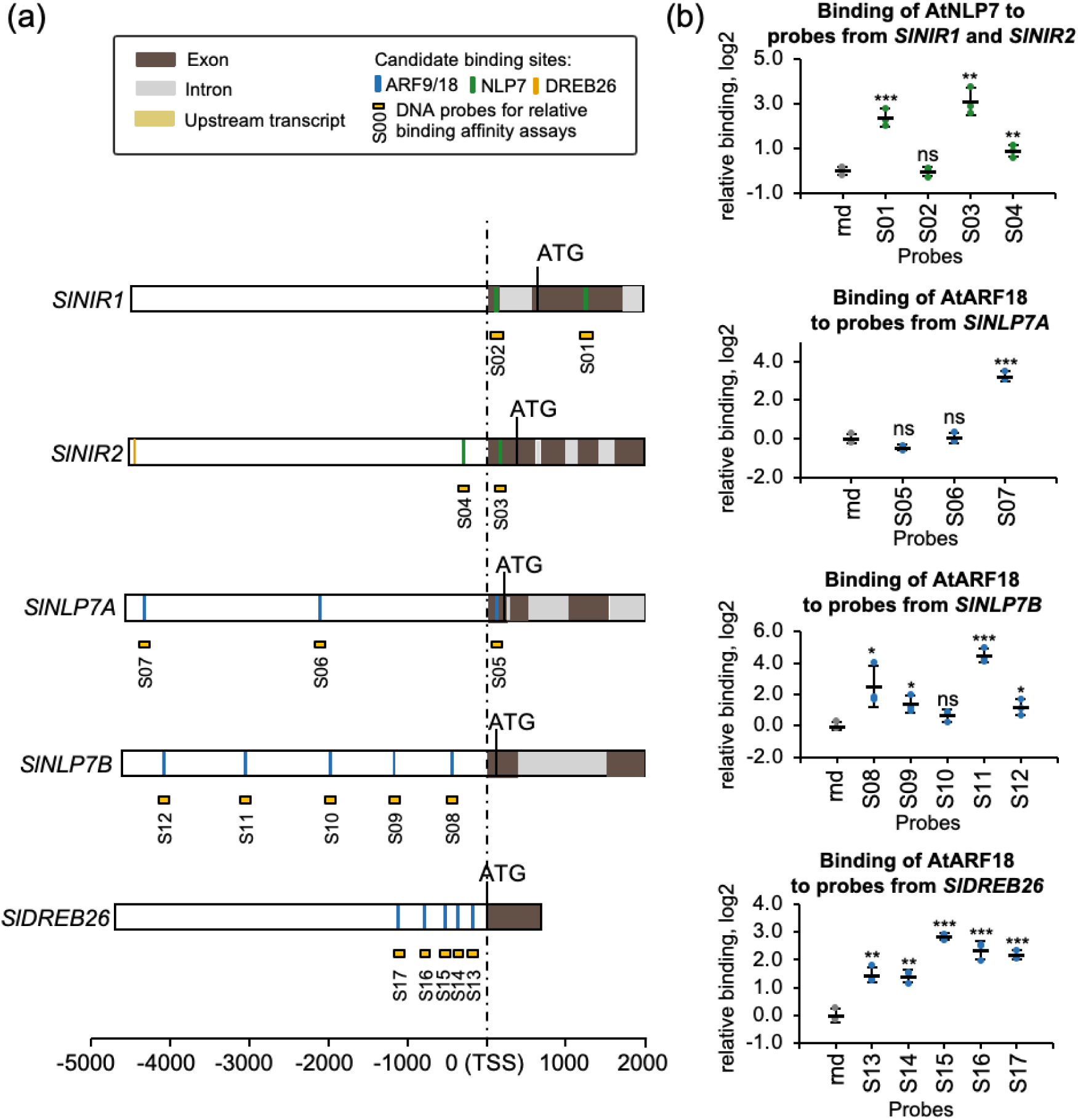
Transcription factor binding motifs and *in vitro* assays support conservation of the regulatory sub-network in tomato. **(a)** Position weight matrices (PWMs) were used to identify candidate transcription factor binding sites in the 5’ regions (up to 5 kilobases upstream and 2 kilobase downstream of the start of transcription) of *SlNIR1, SlNIR2, SlNLP7A, SlNLP7B and SlDREB26.* **(b)** *In vitro* relative binding assays showing binding of AtARF18 and AtNLP7 to candidate binding sites in *SlNIR1, SlNIR2, SlNLP7A, SlNLP7B and SlDREB26,* respectively. Error bars = mean and standard deviation; n=3; P-values were calculated using an unpaired two-tailed Student’s t-test of each sample to the random (rnd) control probe; *p<0.05, ** p<0.01, *** p<0.001; ns = not significant.

### Regulatory interactions are partially conserved between Arabidopsis and tomato

The *in vitro* TF-DNA binding assays, plus the lack of identification of a clear tomato ANAC032 ortholog, suggested distinct differences in the wiring of these N metabolic regulatory networks both in terms of binding and regulation. To determine which of these represent direct regulatory interactions, we generated glucocorticoid receptor (GR) fusions of the SlARF9B, SlARF18, SlDREB26, SlNLP7A and SlNLP7B transcription factors. In order to eliminate dosage-dependent effects, we introduced these constructs into their respective mutant backgrounds, as generated by CRISPR-Cas9 gene editing using *R. rhizogenes* mediated transformation. (23). These roots, hereby called “hairy roots’’ have similar growth and anatomy to primary roots (40) and the same cell type expression as observed with transcriptional reporters when compared with *A. tumefaciens*-transformed plant roots. Hairy roots also respond similarly to increasing concentrations of available nitrate, although with increased sensitivity at 1mM KNO_3_ compared to wild type roots. This was demonstrated by nitrate-dependent increases in the expression of *SlNIR1* and *SlNIR2* as determined by RT-qPCR, by comparing RNAseq of M82 roots with RNAseq from hairy roots, and as observed by nitrate-responsive nuclear-localized GFP expression driven by the *AtNIR1* and *AtNRP* promoters in 0, 1 and 10 mM KNO_3_ in hairy roots, compared to *A. tumefaciens*-transformed *AtNRP*:GFP (Supplementary Data 16). CRISPR-edited mutant alleles were confirmed by amplicon-based sequencing (Supplementary Data 12). As with the assays in Arabidopsis (Figure 3, Supplementary Data 6), transcription factor activity was induced by application with dexamethasone, and direct targets were confirmed by the addition of cycloheximide.

Although the Arabidopsis AtARF18 was able to bind to the *SlNLP7A* and *SlNLP7B* promoters, this was not reflected in the modified TARGET assay. However, an indirect repressive interaction was observed (Supplementary Data 17). SlARF18 and SlARF9B directly bound to the promoter of *SlDREB26* and repressed its expression (Figure 7a-b). In turn, SlDREB26 was found to directly repress the expression of *SlNLP7B* (Figure 7c). SlNLP7A and SlNLP7B both bind to and activate the expression of *SlNIR1* (Figure 7d-e). In several cases, these represent conserved regulatory interactions between Arabidopsis and tomato. At/SlARF18 and At/SlARF9 both repress the transcription of *At/SlDREB26*, and SlNLP7A/B/AtNLP6/7 activate the transcription of *At/S*l*NIR1*. Major differences in these networks include the tomato SlDREB26 repression of *SlNLP7B* which is not observed in Arabidopsis, the Arabidopsis AtNLP6 and AtNLP7 activation of *SlDREB26* which is not observed in tomato, and the Arabidopsis NAC032 interactions which were not able to be tested in tomato due to the lack of a clear ortholog. No feedforward loops were identified in tomato, although they were present in Arabidopsis (Figure 7f).

**Figure 7.**
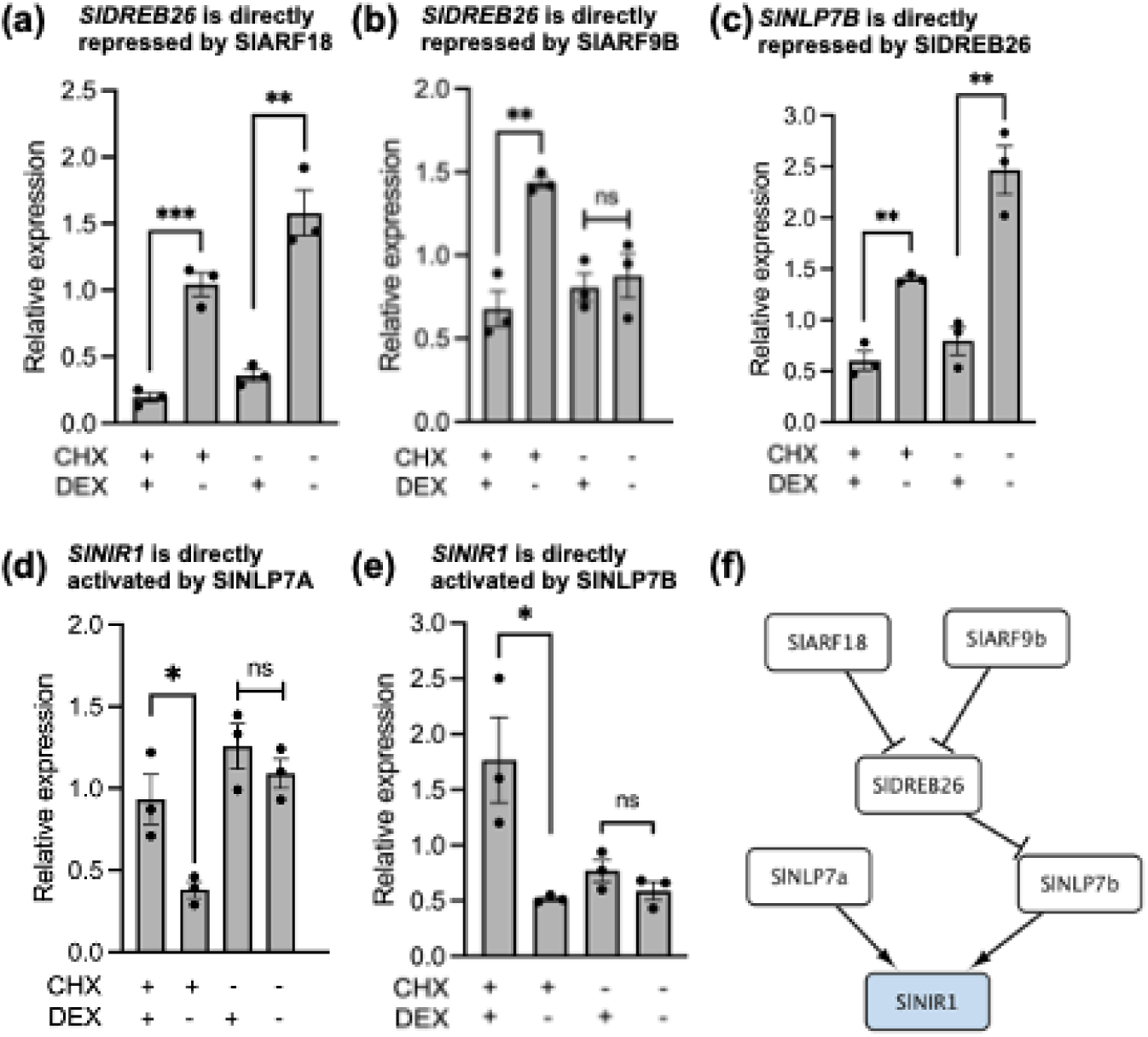
Examples of regulatory consequences of transcription factor expression in tomato root protoplasts via modified TARGET assays. Changes in endogenous gene expression of target genes in modified TARGET assays. *SlDREB26* expression levels are changed by (**a**) SlARF18 and (**b**) SlARF9B, (**c**) *SlNLP7B* expression levels are changed by SlDREB26, *SlNIR1* levels by (**d**) SlNLP7A and (**e**) SlNLP7B. DEX = dexamethasone; CHX = cycloheximide. Values represent the mean and standard error of three biological replicates of which each is the mean of two technical replicates. P-values were calculated using an unpaired two-tailed Student’s t-test, *P<0.05, ** P<0.01, *** P<0.001, and **** P<0.0001. N = 3. ns = not significant (**f**) Schematic of sub-network illustrating the regulatory consequences of transcription factors that directly bind their target genes in tomato. Light blue = metabolic gene (SlNIR1).

### Testing the network models using genetic perturbation

Collectively, these data in both Arabidopsis and tomato provide a predictive model of genetic regulation of plant N metabolism and demonstrate differences in their wiring and components. In summary, SlDREB26 is a key intermediary between SlARF18/9 and SlNLP7A/7B in tomato, as opposed to a target of both ARF18/9 and NLP6/7 in Arabidopsis. A coherent feedforward loop exists between ANAC032, AtNLP7 and *AtNIR1* in Arabidopsis, but ANAC032 is likely absent in tomato. NLP6 and NLP7 activate expression of *DREB26* and result in interconnected feedforward loops in Arabidopsis (Figure 3c), but these regulatory interactions are absent in tomato (Figure 7f).

In a classically defined systems biology approach, the predictive nature of models can be tested using genetic perturbation <u>(Ideker et al. 2001)</u>. Protoplast-based assays have been highly effective at elucidating multiple aspects of the Arabidopsis nitrogen transcriptional regulatory network (Alvarez et al. 2020; Para et al. 2014; Brooks et al. 2019). We hypothesized that a protoplast-based assay could be used to test multiple Arabidopsis and tomato mutant combinations in different concentrations of available nitrogen. We first tested the nitrate response of Arabidopsis and tomato protoplasts by monitoring the expression of *AtNIR1, AtNIA1, SlNIR1* and *SlNIA1* via qRT-PCR. Their expression increased as the concentration of nitrate increased (Supplementary Data 18). As a quantifiable output of the nitrogen network, we used the synthetic *NITRATE-REGULATED PROMOTER (NRP)* (41) fused to luciferase (Supplemental Data 18).This reporter is robustly activated as concentrations of available nitrate increase, and has been used in both Arabidopsis and crop plants (11, 19, 41) and in tomato (Supplemental Data 16). As the *NRP* is comprised of large promoter fragments that contain other binding sites, we additionally tested an optimized version of a previously reported 4xNRE minimal synthetic promoter (Konishi and Yanagisawa 2010), which exclusively contains four copies of an NLP7 binding motif. These reporters were transfected into protoplasts derived from wild type Arabidopsis or tomato roots. Their expression similarly increased as nitrate concentrations increased (0, 1 and 10 mM KNO_3_, Supplementary Data 18). Protoplast viability was maintained at a minimum of 60% at 24 hours after transfection (Supplementary Data 18). To test the influence of TF genetic perturbation, we monitored NRP and 4XNRE reporter activity in protoplasts from NLP6/7 mutant Arabidopsis and tomato roots (*tnlp6, atnlp7-1, slnlp7a/7b).* Luciferase was quantified relative to Col-0 primary roots or empty vector-transformed M82. In all mutants, responsiveness of the NRP and the 4XNRE was vastly reduced (Figure 8, Supplementary Data 18). We named this assay PAROT: <u>P</u>rotoplast <u>A</u>ssay <u>R</u>eporting <u>O</u>verall <u>E</u>ffects of <u>T</u>Fs and subsequently used the NRP reporter to monitor the influence of multiple mutant combinations on the transcriptional regulation of nitrogen metabolism in Arabidopsis and in tomato.

**Figure 8.**
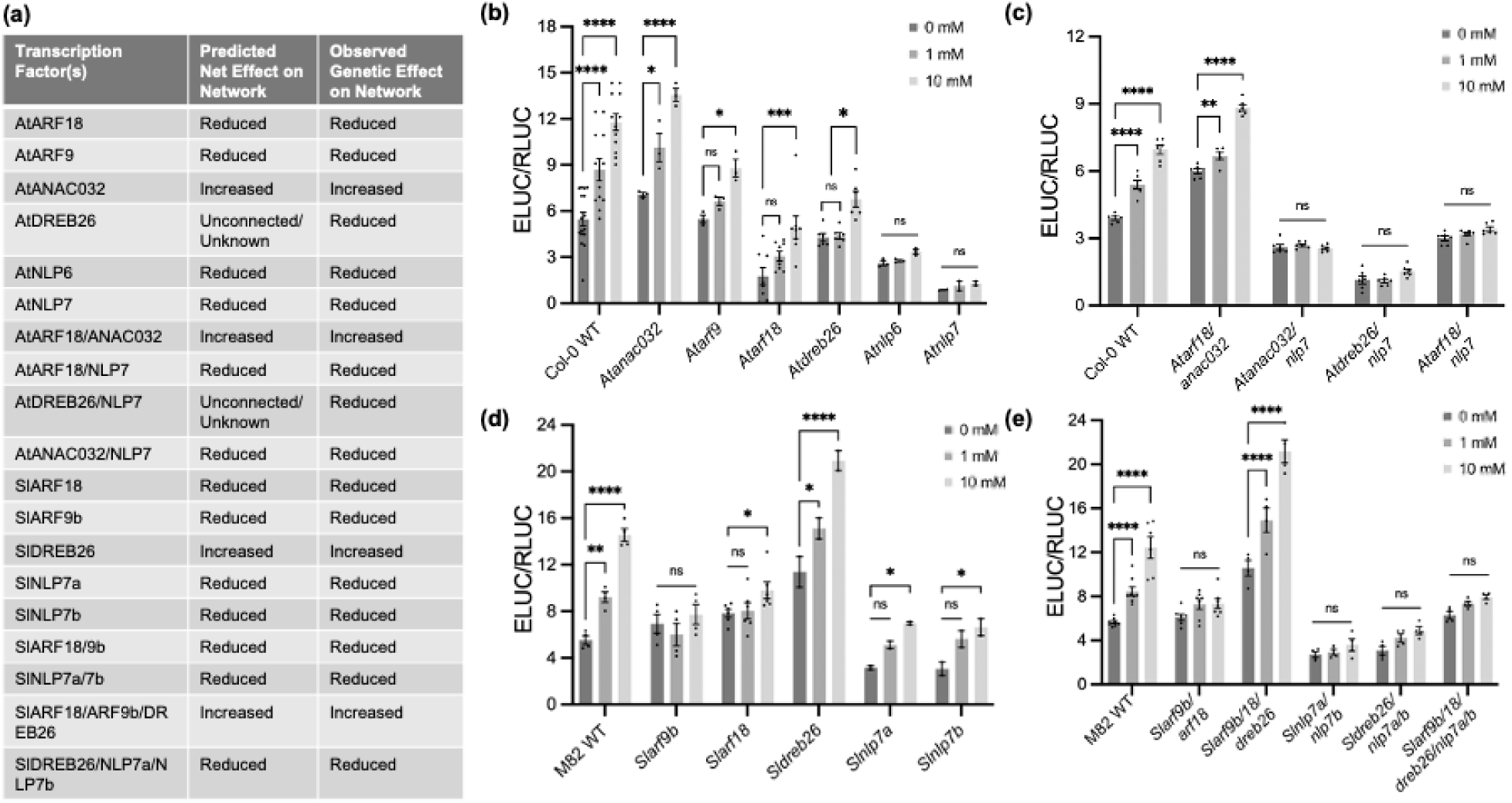
NRP:reporter (PAROT) assay in Arabidopsis and tomato mutants. (a) Regulatory models from Figure 3C and Figure 7F were used to predict the net effect on transcriptional regulation of nitrogen metabolism in the respective single and higher order mutant combinations as measured using the NRP reporter and the PAROT assay (predicted net effect on network). The observed output on the network is summarized in the third column. (b-e) Observed changes in network compared to wild type as quantified in a PAROT assay in which protoplasts are transformed with a dual vector setup where the nitrate-responsive *AtNRP* drives the expression of emerald luciferase (ELUC) and the constitutive NOS promoter drives expression of red luciferase (RLUC). Changes in normalized luminescence of ELUC in response to differing concentrations of nitrate in (b) Arabidopsis Col-0 WT, *Atarf9b*, *Atarf18-2, Atdreb26_c_, Atnlp6*, and *Atnlp7-1* (c) Arabidopsis Col-0 WT, *arf18-2/anac032, anac032/nlp7-1, dreb26_c_/nlp7_c_, and arf18-2/nlp7-1* (d) tomato M82 WT, *Slarf9b #8, Slarf18 #2, Sldreb26 #10, Slnlp7a #1, and Slnlp7b #13* (e) tomato M82 WT, *Slarf9b/18 #4, Slarf9b/18/dreb26 #9, Slnlp7a/7b #21, Sldreb26/nlp7a/7b #6, and Slarf9b/18/dreb26/nlp7a/7b #2* at 0, 1, and 10 mM KNO_3_. Error bars are standard error; N=4; P-values were calculated using two-way ANOVA with Tukey’s multiple comparisons test; *p<0.05, ** p<0.01, *** p<0.001; **** p < 0.0001, ns = not significant.

Ten mutant genotypes were generated for Arabidopsis, and nine for tomato, the latter using hairy roots (Supplementary Data 11 and 12). Given the regulatory network architecture in Figures 3c and 7f we first predicted the impact of these mutations and their combinations. In both Arabidopsis and tomato, ARF9 and ARF18 participated in a direct repressive and indirect repressive interaction upstream of *NIR1*. We thus predicted that NRP activity would be reduced. In both species, NLP6/7 directly activated expression of *NIR1*, and thus their predicted effect would also reduce NRP activity. DREB26 was unconnected to NIR1 in Arabidopsis and thus its effect could not be predicted, while in tomato it would act as a repressor and thus NRP activity would be increased. ANAC032 in Arabidopsis directly repressed both *NIR1* and *NLP7*, and its perturbation was thus predicted to increase NRP activity. In predicting higher order genetic interactions, we assumed that the downstream factor’s phenotype would be epistatic. For instance, in an Arabidopsis *arf18/anac032* double mutant, since ANAC032 acts as a repressor, we predicted that NRP activity would be increased. Similarly, in tomato, in an *arf18/9b/dreb26* mutant, the *dreb26* phenotype would be epistatic and NRP activity would be increased; while in a *dreb26/nlp7a/nlp7b* mutant, the *nlp7* phenotype would be epistatic and NRP activity would be reduced.

In all cases the experimentally observed interactions matched our predictions and supported the regulatory models. In both Arabidopsis and tomato, a single *Atnlp6* or *Atnlp7* or *Slnlp7a* or *Slnlp7b* mutant alone showed reduced sensitivity in its NRP response to increasing nitrogen, with the double in tomato showing near complete insensitivity (Figure 8c,f, Supplementary Data 19 a,d). The NLP7 mutant phenotypes are consistent with AtNLP7 as a major regulator of *AtNIR1* expression in Arabidopsis (32, 42), and tomato (Figure 5). The more extreme phenotype of the *Slnlp7a/7b* mutant and the *Slarf18/9b* mutants indicates that they both additively contribute to nitrogen responsiveness (Figure 8d,e). Our assumption regarding epistasis was also experimentally validated with the *Atanac032* phenotype being epistatic to *Atarf18*, the *Atnlp7* phenotype was epistatic to *Atanac032* and *Atarf18* (Figure 8c). In tomato the *dreb26* phenotype was epistatic to *Slarf18/arf9b*, and the *nlp7a/b* phenotype was epistatic to *Sldreb26*, and even to *slarf9b/18/dreb26* (Figure 8d,e). Collectively, these results support a hierarchy whereby ARF18/9 are at the top of the regulatory network, and NLP6/7 at the bottom of the network. These conserved interactions are demonstrated in Figure 9a and Supplementary Data 20. In Arabidopsis, ANAC032 acts as an intermediary in the network as does DREB26 in tomato (Figure 9a). Monitoring expression of *NIR1* as a readout in the modified TARGET assays was also an effective target to explore the output of the transcriptional nitrogen network.

**Figure 9:**
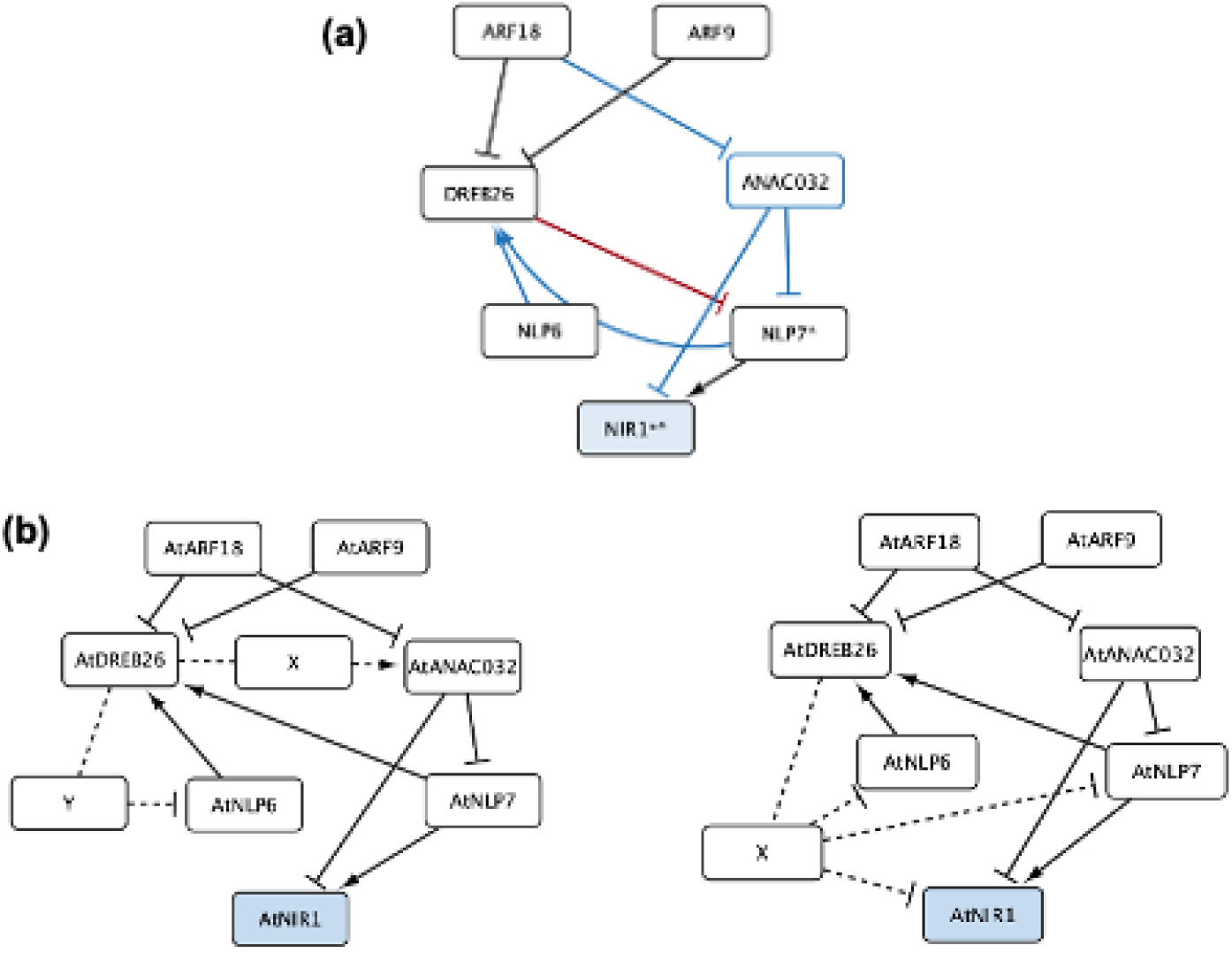
Conserved and species-specific direct regulatory interactions in Arabidopsis and tomato. (a) Three direct, regulatory interactions are conserved between Arabidopsis and tomato transcription factors and targets (black edges). Species-specific regulatory interactions are depicted in blue for Arabidopsis and red for tomato. * = SlNLP7a and SlNLP7b, at AtNLP7, ** = AtNIR1, and SlNIR1 and SlNIR2. Interactions were determined using the modified TARGET assays. (b) Two potential regulatory models that explain the observed influence on the nitrogen transcriptional network in the Arabidopsis DREB26 mutant allele (Figure 8) and which incorporates data regarding the indirect repressive interaction of AtDREB26 on *AtNLP6*, *AtNLP7* and *AtNIR1* (Figure 3 and Supplementary Data 6).

One remaining conundrum was the role of AtDREB26 in the network. In Arabidopsis, DREB26 indirectly repressed *NLP6, NLP7* and *NIR1*. In the PAROT assay, mutation of *dreb26* resulted in decreased sensitivity of the NRP reporter. Furthermore, the *adreb26/atnlp7* double mutant showed increased insensitivity relative to the single *atdreb26* mutant phenotype, but similar insensitivity to the *atnlp7* mutant phenotype. We thus predicted two potential regulatory scenarios. In the first, AtDREB26 indirectly represses *NLP7* and *NIR1* through an unknown gene X which activates *ANAC032*. AtDREB26 also represses *AtNLP6* via an unknown gene Y (Figure 9b). In a second scenario, AtDREB26 represses *NLP6, NLP7* and *NIR1* through an unknown gene X (Figure 9c).

## DISCUSSION

Changes in N availability are perceived and transduced via a series of signaling events culminating in rapid changes in transcription followed by changes in plant growth for adaptive purposes (7, 11). In order to engineer plants with increased N use efficiency via transcriptional regulation, we need specific knowledge of which transcription factors are involved, and which target sites they bind. We also need to understand the nature of these gene pathways, their direct and indirect interactions, and how transcriptional information flows through these pathways to transcriptionally regulate N metabolism.

NLP7 is a critical transcriptional regulator of N metabolism, and its activity is controlled by multiple factors - it directly senses nitrate, it’s nuclear localization is controlled by phosphorylation, and an understudied mode of its regulation is by control of its transcript abundance (Marchive et al. 2013; Liu et al. 2022). Increasing transcription of *NLP7* results in improved plant growth and C:N balance <u>(Yu et al. 2016)</u>, and is used as a measure for physiological significance of N-regulatory factors (Brooks et al. 2019), demonstrating the importance of its transcriptional regulation Transcriptional activity of NLP7 can be read out by changes in transcript abundance of its direct downstream targets, like that of *NIR1* (Alvarez et al. 2020; <u>Hummel et al. 2023</u>). NIR1 is a key enzyme in nitrogen assimilation, reducing nitrite to ammonium. Mutation of *NIR1*, similar to that of *NLP6/NLP7* and *NRT1.1*, results in chlorotic plant growth and perturbed nitrogen metabolism (<u>Costa-Broseta et al. 2020</u>; Cheng et al. 2023; <u>Tsay et al. 1993</u>). Therefore, precise and likely subtle changes in transcription of these genes will exert a meaningful influence on N metabolism and plant growth. Indeed, this is viewed in nature, where differences in transcription of *NIR1, NIA1* and *NRT1.1* are associated with changes in nitrogen sensitivity in genetically varying Arabidopsis populations <u>(North et al. 2009; Sakuraba et al. 2021)</u>.

N-mediated coordination of growth occurs via action in spatially restricted cell types (Zhang and Forde 1998; Zhang and Forde 2000; Gifford et al. 2008; <u>Vidal et al. 2010</u>). In addition, there is complex interconnection and feedback between N signals, transcription and hormone-mediated control of growth (<u>Vega et al. 2019</u>; Krouk et al. 2010; <u>Ma et al. 2014; Song et al. 2013</u>). Given this multi-layered and spatiotemporally diverse regulation, network-focused studies utilizing TARGET, protoplasts or other heterologous systems prove effective at elucidating the underlying architecture of nitrogen-mediated regulation (Alvarez et al. 2020; Brooks et al. 2019; Para et al. 2014; Bargmann et al. 2013; Gaudinier et al. 2018). Our approach and findings complement these many elegant network-focused studies (9, 10, 32, 44–49) by focusing on a five gene regulatory circuit predicted to contain multiple feedforward loops and an autoregulatory interaction which acts upstream of the largely understudied transcriptional regulation of *NLP6* and *NLP7* and its closely related paralog *NLP6*.

Dissecting the necessity of *cis*-regulatory elements or *cis*-regulatory modules has defined rules for cell type-specific expression in the bundle sheath (50) and for tomato fruit size and maize yield-related traits (51, 52). We first sought to identify *cis*-regulatory elements that define regulatory relationships within this gene circuit. These transcription factors had not been tested in previous genome-wide TF-directed approaches (O’Malley et al. 2016; Weirauch et al. 2014). We reduced the set to those present within accessible chromatin regions (28), and then our *in vitro* binding assays (Cai et al. 2023) further restricted these to elements which were able to directly recruit the factors under study (Figure 2b). Collectively these indicated a complex sub-circuit comprising 29 interactions via 46 *cis*-elements, albeit without the contribution of AtARF9 due to technical limitations (Figure 2c). The modified TARGET assay dramatically reduced the 29 possible interactions to eight via 15 putative *cis*-regulatory elements (Supplementary Data 3). This reduced set is highly conservative given that they do not reflect N-inducibility nor cellular context. Transient interactions are also under-represented in this assay type but are prevalent within the N regulatory network at large (32). The resulting sub-network architecture contains several interaction types that will be exciting in the future to monitor in the context of generating dynamic, non-linear behavior. *AtANAC032* is a newly identified gene that controls N-mediated regulation of root system architecture (Figure 5), and it participates in a coherent feedforward loop along with AtNLP7 and AtNIR1 coherent feedforward loop. The final outcome of this feedforward loop in terms of its temporal and spatial consequences remains to be determined. The mode by which AtDREB26 is able to influence *AtNIR1* expression also remains to be elucidated, using the predictive models guided by the experimentally generated genetic data (Figure 8-9). While Arabidopsis serves as an excellent organism to study genome-wide transcriptional regulation of N metabolism, an important question is how research in this species can be translated to a crop. tomato. The molecular genetic nature of tomato N-metabolic gene regulation has been minimally studied (Asins et al. 2017; Chapin et al. 1988; <u>Wang et al. 2001; Julian 2022; Sunseri et al. 2023</u>), likely due to its status as a specialty crop species that is most often grown in smaller-scale agricultural systems with unlimited access to mineral nutrients. Our analysis of root system architecture and the global transcriptional response to differing concentrations of available N indicate a remarkable similarity to the Arabidopsis response. At the molecular level, there were some differences in gene content with at least two tomato *NIR*, *NRT* and *ARF9* genes relative to their Arabidopsis counterparts, and the lack of an identifiable *ANAC032* ortholog. While both the Arabidopsis and tomato NLP7 mutant alleles demonstrated similar N-dependent changes in root system architecture, those of ARF18/9 demonstrated some similarities and differences, suggesting changes in their regulatory network (Figure 5). Dissection of the tomato circuit with both the *in vitro* and *in vivo* assays provided evidence of likely differences in their genetic architecture. The in vitro assays with the Arabidopsis TFs and tomato predicted *cis*-regulatory elements suggested that interactions could exist between the seven tomato genes via twelve *cis*-elements (Figure 6). Modified TARGET assays further confirmed five of these interactions in tomato as direct and regulatory (Figure 7f). The remaining that could not be validated could represent a true lack of interaction, or transient interactions not able to be captured by this system (32). No feedforward loops were identified in tomato (Figure 7). Key changes in circuitry between the two species include (i) the repressive function of SlDREB26 on *SlNLP7B* while in Arabidopsis AtDREB26 does not regulate any target genes; (ii) in Arabidopsis AtNLP6 and AtNLP7 activate *AtDREB26* transcription with no such interaction in tomato; and (iii) the lack of ANAC032 to the tomato module. Molecularly, the observed variation in regulatory interactions (Figure 3c, 7f), and the similarity in N-dependent root system architecture of the tomato and Arabidopsis *nlp7* mutants suggests the convergent function of their respective circuits.

The regulatory models presented in Figure 3C were experimentally tested by systematic genetic perturbation. We drew on approaches used in Arabidopsis to monitor the transcriptional outcomes of auxin and cytokinin hormone signaling pathways via a synthetic reporter (62–64) - the nitrogen-responsive NRP, and used a refined NRP with 4 repeats of the NLP7 binding site (Supplementary Data 18) (41). The advantage of this protoplast-based approach is that we could test the N-dependent inducibility of these different genes and their interactions. Remarkably, using single to quintuple mutant combinations, the regulatory models in Figure 3c and 7f were able to be genetically validated in the PAROT assays. In the future, the spatial and temporal action of these factors remains to be elucidated.

Collectively, these data illustrate divergence and conservation of a N-mediated transcriptional regulatory circuit in Arabidopsis and tomato. The circuit is undoubtedly more spatially and temporally complex, and involves the interaction with many more genes. However, the fact that these assays used to generate the two predicted and then genetically validated models, in terms of NLP7-directed transcriptional activity, suggests that they provide an important avenue to elucidate significant regulators of NLP7. Further, the tomato root system architecture mutant phenotypes demonstrate that these genes are physiologically relevant in a crop species. The protoplast assays reflect a whole-organ response and root system architecture traits are determined by the activities of discrete populations of cells in either xylem pole pericycle cells, lateral root founder cells, and the root stem cell niche. In the future, cellular-resolution assays must be used to untangle these aspects of the network and how they coordinate changes in N concentration with plant growth. These data provide a foundation to engineer robust nitrogen regulatory circuits in both species. The identified *cis*-regulatory modules serve as potential control points. It remains to be determined in each species exactly which *cis*-elements function in a given cell type or N concentration, as well as how or if these transcription factors coordinately act together to regulate target gene expression. Is this by distinct affinities of the transcription factor for its target or differing concentrations of the transcription factor itself? Is there competition between transcription factors in the case of feedforward loops? Indeed, many questions remain to be answered, but this work demonstrates how diverse assays can be used to provide blueprints for elucidation of network architecture and for future synthetic approaches to engineer improved N use efficiency in plant roots.

## MATERIALS AND METHODS

### Identification of candidate transcription factor binding sites

Upstream sequence regions consisting of 2 kilobases (kbs) upstream of the earliest annotated transcription start site (TSS) and 1 kb downstream of *AtANAC032* (AT1G77450), *AtARF18* (AT3G61830), *AtARF9* (AT4G23980), *AtNLP6* (AT1G64530), *At NLP7* (AT4G24020), *AtDREB26* (AT1G21910) and *AtNIR1* (AT2G15620) were extracted from the Arabidopsis TAIR 10 (66) genome assembly. These regions were truncated if they overlapped with the transcript of a protein coding gene. Alternatively, they were extended if a longer region was included in previously reported yeast-1-hybrid data (17). ATAC-seq data from (28) were used to identify regions of open chromatin in Arabidopsis roots and shoot tissues. Individual .bed files for each replicate were concatenated and BEDTools (v2.29.2) merge (67) was used to combine overlapping peaks. Candidate binding sites for AtNLP7 (TGNCYYTT) and AtDREB26 (YCRCCGHC) were identified using position weight matrices (PWMs) from DAP-seq (30) in FIMO (v5.4.1) (68) with a p-value cut-off of 10^−4^ and a zero-order background model created using fasta-get-markov (29). Motifs with q-value <0.05 were considered high-confident binding sites. PWMs are provided in Supplementary Data 1. As no empirical binding motif data was available for AtNLP6, AtARF9, AtARF18 or AtANAC032, we identified closely related proteins from the same TF families for which data was available and compared sequence identity within the DNA binding domains (Supplementary Data 2). Finding these domains to be highly conserved, the PWM for AtNLP7 was also used to predict sites for AtNLP6 (TGNCYYTT), and the PWM for ATAF1 (KACGTR) was used to predict binding sites for ANAC032 (Supplementary Data 1). The PWM for AtARF2 was used to predict binding sites for AtARF9 and AtARF18. In addition, TGTCTC and TGTCGG motifs were also annotated as candidate sites as they have previously been shown to bind auxin response factors (ARFs) with higher affinity than other auxin response elements (69). Finally, pairs of TGTCNN motifs spaced less than 14 bp apart were also annotated as candidate ARF TFBSs, as paired motifs were previously shown to be enriched and preferentially bound in DAP-seq data (69, 70).

For tomato, upstream regions consisting of 5 Kbs upstream and 2 Kbs downstream of the annotated TSS of *SlDREB26* (*Solyc11g012980*), *SlNIR1* (*Solyc10g050890)*, *SlNIR2* (*Solyc01g108630), SlNLP7A* (*Solyc08g008410*) and *SlNLP7B* (*Solyc08g082750*) were extracted from *Solanum lycopersicum* ITAG5.0 (https://phytozome-next.jgi.doe.gov/). To confirm similarity, the DNA binding domains of tomato and Arabidopsis orthologs of ARF9, ARF18, DREB26, NLP6 and NLP7 were aligned (Supplementary data 15). Candidate binding sites in the upstream regions of the tomato genes defined above were then analyzed using the same PWMs and parameters as defined above for Arabidopsis. A custom Jupyter Notebook (71) was used to create schematics of upstream regions illustrating gene features, candidate binding sites and open chromatin.

### *In vitro* transcription factor binding assays

Coding sequences of AtANAC032, AtARF18, AtARF9, AtNLP6, At NLP7, and AtDREB26 were amplified using primers to introduce the Gateway® recombinase sequences attB1 and attB2 and a C-terminal HiBIT tag. Amplicons were cloned into pDONR207 (attP1-ccdB-attP2) to create pENTR clones (attL1-TF:HiBiT-attL2) using Gateway®BP Clonase™II Enzyme Mix (Invitrogen). The resulting pENTR clones were combined with the Gateway®-ready pDEST vector, pH9GW(attR1-ccdB-attR2), a modified pET28a(+) vector (Novagen), using the Gateway®LR Clonase™II Enzyme Mix (Invitrogen). Details of all plasmids are available in Supplementary Data 21. Sequence-verified expression clones were transformed into *E. coli* BL21 and cultures were grown and protein purified as described in (Cai et al. 2023). Protein yield was quantified by measuring luminescence with the Nano-Glo® HiBiT Extracellular Detection System (N2420, Promega, Madison, WI) and HiBiT Control Protein (N3010, Promega). We were unable to obtain soluble protein of AtARF9. Details of all plasmids are available in Supplementary Data 21. Initially, binding of each TF to a double strand probe (DSPs) containing a known binding site was verified (Cai et al. 2023). Subsequently, the ability of each TF to bind to DSPs containing candidate binding sites from Arabidopsis and tomato promoters, relative to random DSPs, was quantified using the same method. Details of all probe sequences are available in Supplementary Data 4.

### Modified transient transformation system for genome-wide transcription factor target discovery (TARGET) assay

For Arabidopsis, constructs were generated using the Loop assembly toolkit (72) using one-step restriction-ligation protocols detailed in (73). Coding sequences of each TF were cloned (concurrently removing the stop codon as well as any internal instances of BsaI and SapI with point mutations to introduce synonymous codons with similar Arabidopsis codon usage) into pUAP4 (72) to create Level 0 standard parts comparable with the Phytobrick standard (74). These level 0 parts were assembled into the level 1 Loop acceptor (pCk2; Addgene #136696) in a one-step restriction-ligation reactions with a double CaMV35s-ΩTMV promoter/5′ UTR (pICH51288; Addgene #50269), a C-terminal GR tag (pEPOZ0CM0137; Addgene #197535) and the CaMV35s terminator (pICH41414). Details of plasmids are in Supplementary Data 21. Col-0 seeds were germinated and grown in potting medium (two-parts sieved compost to one-part sand) within controlled environment chambers with a 16 h photoperiod at 22 °C with 120–180 μmol/m^2^/s light intensity. The photoperiod was reduced to 8 h two days before leaves were harvested for the preparation of mesophyll protoplasts. Mesophyll protoplasts were prepared from *A. thaliana* Col 0 as previously described (75). For each transcription factor, protoplasts were quantified and divided into twelve aliquots of 200 µL in transfection buffer (0.4 M mannitol, 15 mM MgCl_2_, 4 mM MES, pH 5.6), each containing approximately 1 × 10^4^/mL intact protoplasts. In addition, for each batch of protoplasts, three aliquots were transfected with a control plasmid expressing nuclear-localized YFP. Plasmid DNA was prepared using the Plasmid Plus Midi kit (Qiagen, Hilden, Germany) except that three additional wash steps were performed before DNA elution from the column. 10 μg of purified DNA in transfection buffer (2 g of PEG Mn 4000 (Sigma, 81240) in 2 mL of 500 mM mannitol and 0.5 mL of 1M CaCl_2_) was mixed with each aliquot of protoplasts before protoplasts were washed and resuspended in 300 μL of washing buffer (154 mM NaCl, 125 mM, CaCl_2_, 5 mM KCl, 2 mM MES; pH5.6). Following incubation at 24°C with 70 µmol/m^2^/s constant light, the transformation efficiency of each batch was confirmed to be >70% in the control aliquot by quantification of YFP fluorescence in the nuclei using an inverted fluorescence microscope (Zeiss Axio Observer Z1 or ThermoFisher Evos). Aliquots transfected with TF:GR plasmids were treated with 20 mm KNO3 and 20 mm NH4NO3 for 2 hours. Following this, either 10 μM dexamethasone (in 100% EtOH) for 3 hrs; 35 μM cycloheximide (in DMSO) for 20 mins; 10 μM dexamethasone and 35 μM cycloheximide applied sequentially; or no treatment, were added to three aliquots of protoplasts. When a treatment was not given, equivalent volumes of EtOH or DMSO were added. After 3 hours, cells were harvested and RNA was extracted using the Sigma Spectrum total RNA kit and cDNA was synthesized using M-MLV reverse transcriptase (Thermo Fisher). Quantitative reverse-transcription-PCR (qRT-PCR) was performed on a QuantStudio™ 6 Pro Real-Time PCR System (Applied Biosystems A43182) in 384-well plates. Amplifications were done in 10 μL reactions containing 1× SYBR® Green JumpStart™ Taq ReadyMix™ (Sigma S4438), 0.2 μm each primer and 6 ng cDNA template. No template and no reverse-transcriptase controls were also performed. Each amplification was performed in duplicate (technical repeat). A QuantStudio™ 6 Pro 384-well standard, relative quantification with melt program (comparative Ct with melt) was used with the following parameters: 94 °C for 2 minutes then 40 cycles of 94 °C for 15 seconds, 58 °C for 1 minute. A melt curve was performed at the end of the run to confirm the specificity of the amplification. Primer sequences are provided in Supplementary Data 22.

For tomato, TF coding sequences were either amplified from cDNA using Phusion polymerase (Primer sequences are provided in Supplementary Data 22) and cloned into pENTR™ /D-TOPO™ (ThermoFisher), or synthesized (Genewiz, South Plainfield, NJ). Sequence verified clones were used in LR cloning reactions with the Gateway-enabled pBEACON DESTINATION vector (26), which contains a C-terminal GR-tag. The resulting EXPRESSION clones were sequence verified. Details of plasmids are provided in Supplementary Data 21. Protoplast isolation from hairy root cultures (see below for methods of hairy root production) was performed as previously described (75) with the following changes: Two-weeks after sub-cloning, protoplasts were isolated from hairy roots tissues harvested from three petri dishes using 1.5% cellulase R10, 1.5% cellulase RS, 0.4% macerozyme and 0.13% pectolyase enzyme solution. Hairy roots were lightly chopped with a sharp razor blade using ∼10 strokes and incubated in the enzyme solution for 4 hours at 28-30℃, in the dark, with shaking at 85 rpm. After incubation, the protoplast solution was filtered with 70 and 40 µm strainers, and centrifuged at 500 g 5 min at 4 ℃ once using 12 ml cold W5 solution in 14 ml round bottom culture tube, and second time using 2 ml W5 in a round bottom microcentrifuge tube. For downstream transfection, hairy root protoplasts are resuspended at a ∼4-8 × 10^5^ cells/ml concentration in MMG solution. Protoplast transfection, RNA purification and qRT-PCR was carried out as for Arabidopsis.

### Transactivation luciferase assays

Promoter regions of *AtARF18*, *AtANAC032*, *AtDREB26*, *AtNIR1*, *AtNLP6* and *AtNLP7* including ∼1000 bp upstream of the TSS and the 5′ UTR, were amplified and cloned into pUAP4 (72) to create Level 0 standard parts comparable with the Phytobrick standard (74). During cloning, internal BsaI sites in *ANAC032_P_* (301 bp upstream of TSS) and *ARF18_P_* (258 bp upstream of TSS) promoters were removed by the introduction of a mutation. These mutations did not disrupt any candidate TFBSs. These Level 0 promoter parts were used in in one-step digestion-ligation reactions with the Level 1 Loop pCk1 (Addgene #136695) backbones together with parts containing LucN (pEPYC0CM0133, Addgene #154595), a C-terminal 3×FLAG® epitope tag (pICSL50007; Addgene #50308), and 3′ UTR and terminator sequences from *Agrobacterium tumefaciens* nopaline synthase (*AtuNOS*_T_) (pICH41421, Addgene #50339). As a batch calibrator, an equivalent plasmid was constructed with *AtuNOS*_P_ (pEPSW1KN0035) and experiment calibrator plasmids (pEPSW1KN0034 and pEPSW1KN0072) for ratiometric quantification were assembled from *CaMV35s*_P_:tobacco mosaic virus (ΩTMV) (pICH51277, Addgene #50268) + firefly luciferase (LucF) (pEPAS0CM0008, Addgene #154594) + *CaMV35s*_T_ (pICH41414), or *AtuNOS*_P_ (pICH42211, Addgene #50255) + firefly luciferase (LucF) (pEPAS0CM0008, Addgene #154594) + *AtuOCS*_T_ (pICH41432, Addgene #50343). Finally, the coding sequence of each transcription factor and the YFP coding sequence were cloned into pUAP4 in one-step restriction-ligation reactions to create Level 0 parts (with stop codons) that were subsequently assembled into the level 1 Loop acceptor (pCk2; Addgene #136696) in one-step restriction-ligation reactions with a *CaMV35s_P_*-ΩTMV (pICH51277; Addgene #50268) and *CaMV35s_T_* (pICH41414). Details of plasmids are in Supplementary Data 21.

Arabidopsis mesophyll protoplasts were prepared as described above. For each promoter, six protoplast aliquots were prepared. Three aliquots received three plasmids: *TF*_P_-LucN-*AtuNOS*_T_ + *CaMV35s*_P_-ΩTMV-TF-*CaMV35s*_T_ + *CaMV35s*_P_-LucF-*CaMV35s*_T_ at equimolar ratios totalling 1000 fmol. In the other three aliquots *CaMV35s*_P_-ΩTMV-TF-CaMV35s_T_ was substituted for *CaMV35s*_P_-ΩTMV-YFP-*CaMV35s*_T_. The entire experiment was repeated three times and, in every experiment, three protoplast aliquots were transfected with a batch calibrator (*CaMV35s*_P_-LucN-*AtuNOS*_T_ + *CaMV35s*_P_-LucF-*CaMV35s*_T_). Full experimental details are provided in Supplementary Data 5. Protoplasts were transfected as described above and, after 18 hours, collected by centrifugation. To each aliquot, 30 μL lysis buffer protease inhibitor mix (1x passive lysis buffer (Promega E1941) and 1x protease inhibitor cocktail (Sigma P9599)) was added. After 15 minutes on ice, debris was collected by centrifugation and 30 μL of lysate was added to each well of a 4titude 96-well white polystyrene microplate flat bottom plate (Azenta Life Sciences). The NanoGlo® Dual-Luciferase® Reporter Assay System (Promega N1610) was used to quantify NanoLuc and LucF luminescence in a CLARIOstar Plus plate reader with each well measured for ten seconds at 3600 gain with one second settling time.

### Tomato transcriptome and gene expression analysis

Seeds were germinated on a nitrogen-free MS medium (Caisson labs, MSP07) supplemented with the appropriate concentration of KNO_3_ at 22 °C in constant light. The plates were placed vertically in growth chambers at 22 °C, 16 h light/8 h dark and whole roots were sampled 9 days post-germination. Tomato roots infected with competent *R. rhizogenes* not carrying any plasmid are referred to as WT hairy roots (see method for hairy root production for details). WT hairy roots were transferred to nitrogen-free MS medium (Caisson labs, MSP07) supplemented with the appropriate concentration of KNO_3_. Plates were placed horizontally in growth chambers at 22 °C, 16 h light/8 h dark. Newly-grown hairy roots were sampled 9 days post-transfer and used for both RNAseq and qPCR experiments. Total RNA was extracted from using the Quick-RNA Plant Miniprep Kit (Zymo Research). First-strand cDNA was synthesized using SuperScript IV Reverse Transcriptase (Thermo Fisher Scientific) with 500 ng of total RNA as a template.

RNA-seq libraries were prepared using the QuantSeq 3’ mRNA-Seq Library Prep Kit (Lexogen) according to the manufacturer’s protocol. Libraries were submitted to the UC Davis DNA Technologies Core and sequenced with an Illumina single-end HiSeq 4000 SR100. Four biological replicates and three technical replicates were used for each sample. After sequencing, reads were pooled, trimmed, and filtered using Trim Galore! (v0.4.5) <u>(Babraham Bioinformatics - Trim Galore!)</u> with the parameter - a GATCGGAAGAGCACA.<u>(Bray et al. 2016)</u> Trimmed reads were pseudo-aligned to the ITAG4.1 transcriptome (cDNA) using Kallisto (v0.43.1), with the parameters -b 100–single -l 200 -s 30. Raw RNA-seq read counts were filtered to remove genes with zero counts across all samples. Reads were converted to count per million (CPM) using the cpm() function in edgeR. Genes that were consistently lowly expressed across all samples were removed. Differential gene expression analysis was done using limma in R/Bioconductor, with empirical weights estimated using the voomWithQualityWeights function. Quantile normalization was used to account for different RNA inputs and library sizes. The linear model for each gene was specified as N treatment: log(counts per million) of an individual gene ∼0+N_concentration. Differentially expressed genes were selected based on a false discovery rate < 0.05. Raw sequence data generated during this study have been deposited in the National Center for Biotechnology Information (NCBI) BioProject database under accession number PRJNA975125.

For qPCR from hairy roots, the same cDNA was used in reactions with the SYBR Green Master Mix kit (Thermo Fisher Scientific). For qPCR from protoplasts, RNA was extracted from root protoplasts incubated in appropriate KNO_3_ concentration (see method for Protoplast Assay Reporting Overall Effects of TFs (PAROT) for details of protoplast isolation). RNA was purified as described above and cDNA was synthesised and amplified using the Luna Universal One-Step RT-qPCR Kit (NEB). Reactions were cycled in a CFX384 Touch Real-Time PCR Detection System (Bio-Rad). Three technical replicates were performed for each of three biological replicates. Expression values were normalized to the reference gene *SlEXP (Solyc07g025390*) using the ΔCT method. Data are presented as mean ± SD. Primer sequences are provided in Supplementary Data 22.

### Clustering and gene ontology (GO) enrichment analysis

Genes differentially expressed between N treatments were clustered in groups with similar expression patterns. The log2 CPM median across biological replicates were calculated for each gene and the expression of each gene was then scaled to the median expression across all samples. Hierarchical clustering was performed with the pheatmap v.1.0.12 R package with the Euclidean distance metric to quantify similarity. Genes assigned to each cluster are listed in Supplementary Data 8. Gene Ontology (GO) enrichment analysis was performed with the GOseq v.1.34.1 R package with correction for gene length bias <u>(Young et al. 2010)</u>. The odds ratio for each ontology was calculated with the formula: (number of genes in GO category / number of all genes in input) / (number of genes in GO category / number of genes in all clusters). Enriched ontology terms were selected based on a p-value < 0.05 and an odds ratio > 1. GO categories enriched in each cluster are listed in Supplementary Data 8.

### Phylogenetic tree construction

Phylogenetic trees were generated as previously described (23). Briefly, representative proteomes were downloaded from Phytozome, Ensembl, or consortia sites. Next, BLASTp (76) was used to identify homologous sequences within each proteome. To refine this set of sequences, a multiple sequence alignment was generated with MAFFT v7 (77) and a draft tree was generated with FastTree (78). A monophyletic subtree containing the relevant sequences of interest was then selected. For the final trees, MAFFT v7 using L-INS-i strategy was used to generate a multiple sequence alignment. Next, trimal was used with the -gappyout option. To generate a phylogenetic tree using maximum likelihood, RAxML was used with the option -m PROTGAMMAAUTO and 100 bootstraps. Finally, bipartitions with bootstrap values less than 25% were collapsed using TreeCollapserCL4 (http://emmahodcroft.com/TreeCollapseCL.html). The resulting trees were rooted on sequences from the earliest-diverging species represented in the tree. All trees are provided in Supplementary Data 9.

### Assembly of constructs for CRISPR/Cas9 mutagenesis

All constructs were generated using the Loop assembly toolkit (72) using the methods described in (79). New level 0 DNA parts (e.g. promoters, terminators, CDSs) were either synthesized (Twist Bioscience, San Francisco, CA and Genewiz, South Plainfield, NJ) or amplified by PCR with overhangs containing SapI recognition sites and assembled into pUAP4 (Addgene #136079) to create parts compatible with the Phytobrick standard (74). Targets for mutation were selected using CRISPR-P 2.0 (80). Spacer sequences were synthesized (Integrated DNA Technologies, Coralville, IA or Twist Biosciences, San Francisco, CA) and integrated into single guide RNA (sgRNA) Level 1 expression constructs as described in (79). For Arabidopsis, each final construct contained the FASTred (81) visual non-destructive selectable marker cassette (*AtOLE*_P_-OLE:RFP-*AtOLE*_T_) in position 1; a Cas9 expression cassette (*AtYAO*_P_-SpCas9-E9_T_) or (*AtYAO*_P_-SaCas9-*E9*_T_) in a reverse position 2; a tandem arrays of two to four sgRNA cassettes; and a plant kanamycin resistance cassette (*AtuNOS*_P_-nptII-*AtuOsc*_T_). For tomato, each final construct contained the FASTgreen selection cassette (*AtOLE*_P_-OLE:GFP-*AtOLE*_T_) in position 1; the SpCas9 expression cassette (*AtRPS5A*_P_-SpCas9-E9_T_) in reverse position 2; two sgRNA expression cassettes in tandem in positions 3 and 4; and a plant kanamycin resistance cassette (*AtuNOS*_P_-nptII-*AtuOsc*_T_) or basta resistance cassette (*CaMV35s*_P_-Bar-*CaMV35s*_T_) in position 5.

### Production of Arabidopsis lines with CRISPR/Cas9-induced mutations

Assembled CRISPR/Cas9 plasmids were transformed into *A. tumefaciens* GV3101 and liquid cultures were grown from single colonies in LB supplemented with 50 mg/mL rifampicin, 25 mg/mL gentamycin and 50 mg/mL kanamycin at 28 °C. Cells were collected by centrifugation and resuspended to OD_600_ 0.8 in 5% sucrose, 0.05% Silvet L-77 and sprayed onto the floral tissues of ten Col0 plants. Plants were sealed in black plastic bags for 24 hours. Seeds were collected from mature siliques and surface sterilized with 70% EtOH for 10 min followed by 3-5% sodium hypochlorite for 10 min. For selection of T_1_ transgenic plants, sterilized seeds were germinated and grown on MS supplemented with 75 mg/mL kanamycin with 16 h light 22 °C. T_1_ seedlings were transferred to soil and T_2_ seed was collected. Seed batches were examined with a fluorescence microscope to assess segregation of transgene by expression from the FAST-red marker cassette. Non-fluorescent T_2_ seeds from segregating seed batches were selected and germinated. DNA was extracted and PCR and sequencing was performed to confirm absence of the transgene and to identify mutations at the target loci. The genotypes of all lines analyzed in this study are provided in Supplementary Data 11 and 12. T_3_ progeny of each line were grown to bulk seed and to confirm the stability of the genotype and T_4_ seeds were used for subsequent analysis.

### Production of *Rhizobium rhizogenes*-induced tomato hairy roots with CRISPR/Cas9 mutations

Tomato hairy root transformation was performed as previously described (40). Briefly, competent *R. rhizogenes* was transformed by electroporation with each CRISPR/Cas9 construct, plated on nutrient agar (Difco, BD 247940) with the appropriate antibiotics, and incubated for 2-3 days at 28-30 ℃. Single colonies were selected and used to inoculate liquid cultures that were grown overnight at 30 ℃ with shaking. The resulting cultures were used to transform cotyledons. For each construct, 40 fully expanded cotyledons were cut and dipped into liquid *R. rhizogenes* culture and co-cultivated in the dark on MS agar (MS basal media pH 5.8 with 1X vitamins, 3% w/v sucrose, 1% w/v agar) without antibiotic selection for three days. Cotyledons were then transferred to selective MS agar (MS basal media pH 5.8 with 1X vitamins, 3% w/v sucrose, 1% w/v agar, 200 mg/L cefotaxime, 50 mg/L kanamycin). At least 15 independent antibiotic-resistant roots were subcloned for each transformation for genotyping and subsequent analyses. Generally, roots were subcloned on selective MS agar for two rounds before maintenance on MS agar with Cefotaxime (MS basal media pH 5.8 with 1X vitamins, 1% w/v sucrose, 1% w/v agar, 200 mg/L cefotaxime).

For genotyping, samples of root tissues were collected aseptically and genomic DNA was purified. Target regions were amplified, and adapters were ligated to produce barcoded pools of amplicon libraries. Libraries were normalized, pooled, and sequenced using Illumina chemistry (MGH CCIB DNA Core Facility, Massachusetts General Hospital, Boston). Sequencing reads were filtered, de-multiplexed, mapped to reference sequences using a customized script, and visualized in an Integrative Genomics Viewer genome browser (https://igv.org) to identify mutations. At least two independent lines, each with a different mutation, were selected from each transformation. The genotypes of all lines are provided in Supplementary Data 12.

### Production of stable tomato lines with CRISPR/Cas9-induced mutations

CRISPR/Cas9 constructs (as noted above) were introduced in *Agrobacterium tumefaciens* and transformed into *Solanum lycopersicum* cultivar M82 (LA3475) at the UC Davis Transformation Facility. For each construct, 12-20 first-generation (T0) seedlings were transferred to potting mix and cultivated in a growth chamber at 22 °C with 16-h light/8-h dark cycles for 14 days before transfer to the greenhouse. Genomic DNA was purified from leaf issues and target regions were amplified and sequenced as described above. T1 seeds were collected from lines in which mutations at the target were identified and the resulting plants were genotyped to identify lines with homozygous or biallelic mutations at target sites in which the T-DNA had segregated away. The genotypes of all lines are provided in Supplementary Data 12. Plants were self-pollinated and progeny seeds were collected for further analysis.

### Root system architecture (RSA) analysis

Arabidopsis seeds were sieved to obtain seeds between 250 and 300 µm, sterilized with 50% (v/v) bleach and stored at 4°C for 48 hours before germination on sterile growth media supplemented with the appropriate concentration of KNO_3_ as previously described (17). Arabidopsis mutants were grown in triplicate with Col-0 controls on the same plate. Nine seeds were placed on each plate. The plates were placed vertically in 22 °C growth chambers with 16-h light/8-h dark cycles for 9 days post germination before the plates were scanned and images processed using SmartRoot (82). Tomato seeds were germinated on a nitrogen-free MS medium (Caisson labs, MSP07) supplemented with Phytagel (Sigma-Aldrich,P8169) and appropriate amount of KNO_3_ to mimic different nitrogen treatments. Tomato mutant seeds and their M82 control were grown in triplicate separately with 5 seeds placed on each plate. The plates were placed vertically in 22 °C growth chambers with 16-h light/8-h dark cycles. Roots were scanned every day during 14 days after the first germination at 800 dpi with an Epson Perfection V700 scanner. Images were processed in Fiji software. For Tomato, RSA traits at 10 days after germination (DAG) were measured. For both species, RSA traits including primary root length, total lateral root number, total lateral root length, average lateral root length, total root length (primary root length + total lateral root length), lateral root density (total number of lateral roots divided by primary root length), and the ratio of total lateral root length to total root length were quantified.

All statistical analyses were performed in R version i386 4.3.0 For RSA assays, two statistical tests were performed: unpaired t-tests between wild-type and mutant seedlings for each treatment combination as well as a two-way ANOVA to test for genotype x treatment interactions. Statistical tests were performed on the natural log of a given trait for each individual. After log transformation, outliers that were more than two standard deviations away from the mean within a genotype-treatment group were removed. To test for wild-type response to nitrogen treatment, a one-way ANOVA was performed using the car package’s anova function (anova(lm(log.TraitValue ∼ Treatment + Plate))). To test for significant genotype by treatment interactions, a two-way ANOVA was performed (anova(lm(log.TraitValue ∼ Genotype*Treatment + Plate))). In the case of the tomato RSA analysis, “Plate” and “Genotype” were highly correlated, so the factor “Plate” was set to zero”. Additional t-tests for each pairwise treatment combination were also performed. To test for genotype-specific responses to nitrogen treatment, the natural log of a trait was modeled using a two-way ANOVA (anova(log.TraitValue ∼ Genotype*Treatment + Plate)). For the tomato data, the two-way ANOVA was run as anova(llog.TraitValue ∼ Genotype*Treatment). Please see Supplementary Data 14 for all calculated p-values.

### Assembly of nitrate-responsive reporter constructs

A promoter fragment containing 1 kb upstream of the start codon of *AtNIR1*_P_ was amplified from genomic DNA isolated from Arabidopsis Col-0 seedlings. In addition, 365 bps of a previously described synthetic arabidopsis nitrogen responsive promoter (*AtNRP*_P_) (41) was synthesized (Twist Bioscience, San Francisco, CA). A optimized minimal synthetic promoter containing four copies of the NRE motifs (4 x NRE), truncated to only include NLP7 TFBSs, with 21 bp of random spacer sequence with equal ATCG ratios downstream of each binding site followed by TATATAA fused to a minimal 35s promoter was synthesized. Promoter fragments were cloned into pENTR™/5’-TOPO™ (ThermoFisher) and the resulting ENTRY clones were used in a LR Clonase II reaction with the binary DESTINATION vector pMR099, which contains the coding sequence of nuclear-targeted GFP (GFP_nu_) and a cassette for constitutive expression of plasma membrane-targeted tagRFP (CaMV35s_P_-TagRFP:LTI6b) (40). The resulting expression clones encode *AtNIR*_P_-GFP_nu_ and *AtNRP*_P_-GFP_nu_. Similarly, *AtNRP*_P_ was assembled with the ELuc coding sequence to produce a AtNRP_P_-Eluc-Hsp18-2 expression vector.

### Confocal imaging

All confocal images were captured using a Zeiss Observer Z1 LSM700 (Zeiss) Confocal Laser Scanning Microscopy (water immersion, × 20 objective) with excitation at 488 nm and emission at 493– 550 nm for GFP and excitation at 555 nm and emission at 560–800 nm for mRFP. The tomato seeds or hairy roots were germinated/transferred to MS medium with different KNO_3_ concentrations for 10 days, and root tips of tomato seedlings or hairy roots were collected for imaging. Cell outlines were visualized using a constitutive mRFP expression cassette in the pMR099 vector. Confocal images were false-colored, and brightness/contrast was adjusted in Fiji software.

### Protoplast Assay Reporting Overall Effects of TFs (PAROT)

In this assay, vectors encoding a nitrate-responsive green luciferase (AtNRP_P_-Eluc-Hsp18-2) and constitutively-expressed red luciferase (pNOS-Rluc-tNOS; pGREAT27, Addgene #170915) are delivered to root protoplasts. To obtain Arabidopsis root tissues, seeds were sterilized with 50% (v/v) bleach and vernalized for 3 days before plating on MS media (10 g agar, 0.5 g MES, 10 g sucrose, 4.33 g MS salts, pH to 5.7 with KOH). Around 800 seeds were densely placed in two rows on 120 mm square plates with a Nitex mesh (Sefar) to facilitate recovery of roots from plates for protoplasting. The plates were placed vertically in growth chambers at 22°C with 16-h light/8-h dark cycles for 8-10 days before harvesting. Protoplasts were isolated as previously described (83) with modifications. Briefly, 20 mg pectolyase (Sigma-Aldrich P-5936) and 230 mg cellulase RS (Yakult) were dissolved in 15 mL of protoplast isolation buffer (600 mM mannitol, 20 mM MES, 20 mM KCl, 0.1% BSA, 10 mM CaCl_2_, pH 5.7) and passed through a 0.45 μm filter. Root tissues were harvested from six plates, chopped several times and placed within a 70 μm strainer set within a 6-well petri dish containing 5 ml of the enzyme solution. After 3 hours at room temperature (or 25°C) on an orbital shaker the solution was passed through a 40 μm cell strainer to remove any remaining tissues. Protoplasts were collected and washed in W5 buffer (2 mM MES, 154 mM NaCl, 125 mM CaCl_2_, 5 mM KCl, pH 5.7) by centrifugation at 4°C 500g for 10 minutes with acceleration and deceleration set to zero. Tomato root protoplasts were prepared as described in the modified TARGET assay section.

Protoplasts of both species were counted with a hemocytometer and diluted in MMG buffer (4 mM MES, 0.4 M mannitol, 15 mM MgCl_2_, pH 5.7) at a concentration of 2-8 × 10^5^ cells/ml for transfection with dual luciferase vectors. Fresh 40% PEG transfection solution (4 g of PEG4000 (Fluka, cat. no. 81240) added into 3 mL of H_2_O, 2.5 mL of 0.8 M mannitol, and 1 mL of 1 M CaCl_2_) was prepared at least one hour before transfection to allow enough time to dissolve. In a round bottom microcentrifuge tube, 10 μg of each plasmid was mixed with 100 μL protoplasts and 110 μL PEG solution and incubated for 15 minutes. Protoplasts were again washed with W5 after transfection and suspended in WI buffer (4 mM MES, 0.5 M mannitol, 20 mM KCl, pH 5.7) with appropriate KNO_3_ concentration (0, 1, or 10 mM) for 24 hours before quantification of luciferase in a plate reader, with each well measured for ten seconds at 3600 gain with one second settling time.

The viability of protoplasts was monitored by staining with an equal volume of 50 μg/mL fluorescein diacetate (FDA). Protoplasts were gently mixed by pipetting and immediately imaged using an EVOS fluorescence microscope (ThermoFisher) with an excitation at 470/22 nm and emission at 525/50 nm. Viability was assessed before transfection, immediately after transfection, and after 24 hours (Supplementary Data 18).

## Supporting information

Combined Supplementary Data

## ACKNOWLEDGEMENTS

We gratefully acknowledge the support of the United States Department of Agriculture (USDA), the National Science Foundation (NSF) of the United States of America, and the United Kingdom Research and Innovation’s Biotechnology and Biological Sciences Research Council (UKRI BBSRC). This work was funded by a joint USDA/NSF/BBSRC Breakthrough Technologies Award Grant numbers: USDA: 2019-67013-29012, BBSRC: BB/S020853/1). Additional funding was provided by Institute Strategic Programme Grants to the Earlham Institute: ‘Regulatory Interactions and Complex Phenotypes’ (BBS/E/T/000PR9819) and ‘Cellular Genomics’ (BBS/E/ER/230001C). SW was supported by the BBSRC Norwich Research Park Doctoral Training Partnership (BB/M011216/1 Project No. 2116916). Additional funding was provided by NSF 2118017 and 2119820 and a Howard Hughes Faculty Scholar award to SMB; and NSF 1907088 to GAM. We gratefully thank the UC Davis Genome Center for support of CB. We thank J. Ahn for contribution to protocols for analysis of root system architecture.

## CONTRIBUTIONS

NJP and SB conceived the study and were responsible for fundraising and supervision. SW, Y-MC, MTO, CB and GAM identified candidate binding sites; SW and Y-MC produced recombinant proteins and performed DNA binding assays; MTO, Y-MC, SW, GSD and CB cloned and produced constructs; MTO, SW and GSD performed transient protoplasts gene expression assays to detect regulatory interactions; CB produced and tested the N-responsive reporters; GSD, MTO and Y-MC conducted N-responsiveness assays in root protoplasts; CB generated and analyzed tomato transcriptome datasets; CB performed phylogenetic analysis of tomato genes; CB and GAM identified tomato homologs of Arabidopsis genes; MTO produced Arabidopsis CRISPR lines and conducted genotype analysis of all CRISPR and insertion lines; CB produced and genotyped tomato hairy root CRISPR lines and stable tomato CRISPR lines; MTO, Y-MC, GAM and RS conducted analyses of Arabidopsis plant root systems; CB conducted analyses of tomato root systems; all authors contributed to data analysis and the production of figures; NJP and SB drafted the manuscript text and all authors contributed to revisions.

## LIST OF SUPPLEMENTARY DATA

**Supplementary Data 1.** Logoplots of position weight matrixes used to identify candidate binding sites for transcription factors.

**Supplementary Data 2.** Alignments of the DNA binding domains of Arabidopsis transcription factors.

**Supplementary Data 3.** *In vitro* binding assays showing relative binding of transcription factors to probes from target genes.

**Supplementary Data 4.** Sequences of DNA probes used in transcription-factor DNA binding assays.

**Supplementary Data 5.** Methodology for protoplast transactivation assays.

**Supplementary Data 6.** Expression levels of Arabidopsis genes in modified TARGET assays.

**Supplementary Data 7.** Transactivation luciferase assays.

**Supplementary Data 8.** Nitrate responsive genes in tomato roots.

**Supplementary Data 9.** Phylogenetic analysis of transcription factors.

**Supplementary Data 10.** Domain analysis of tomato orthologues of AtNLP7.

**Supplementary Data 11.** Table of Arabidopsis and tomato mutant lines.

**Supplementary Data 12.** Genotypes of Arabidopsis and tomato CRISPR lines.

**Supplementary Data 13.** Analysis of root system architecture of Arabidopsis and tomato mutant alleles of NLP7 and ARF18.

**Supplementary Data 14.** Statistics for root system architecture analysis

**Supplementary Data 15.** Alignments of the DNA binding domains of tomato and arabidopsis transcription factors.

**Supplementary Data 16**. Characterisation of N responses in hairy roots

**Supplementary Data 17.** Expression levels of Arabidopsis genes in modified TARGET assays.

**Supplementary Data 18.** Exemplification of PAROT assay.

**Supplementary Data 19.** PAROT assay in Arabidopsis and tomato mutants.

**Supplementary Data 20.** Summary of all evidence for regulatory sub networks.

**Supplementary Data 21.** Plasmids used in this study.

**Supplementary Data 22.** Primers used in this study.

