## Supplementary material for "Conservation and divergence of regulatory architecture in nitrate-responsive plant gene circuits": Combined Supplementary Data

**Supplementary Data 1.** Logoplots of position weight matrixes used to identify candidate binding sites for transcription factors.

| TF | Gene ID | Family | PWM logo and data source |
|---|---|---|---|
| NLP7 | AT4G24020 | RW-PRK | 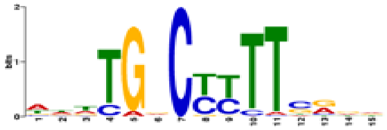 NLP7 (O'Malley et al 2016) |
| NLP6 | AT1G65430 | RW-PRK | |
| ARF9 | AT4G23980 | ARF | 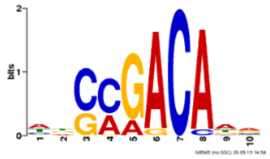 ARF2 (AT5G62000) (O'Malley et al 2016) |
| ARF18 | AT3G61830 | ARF | |
| ANAC032 | AT1G77450 | NAC | 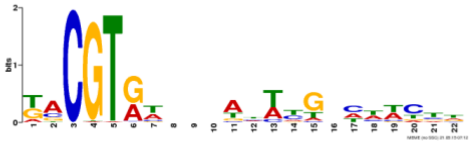 ATAF1 (AT1G01720) |
| DREB26 | AT1G21910 | ERF | 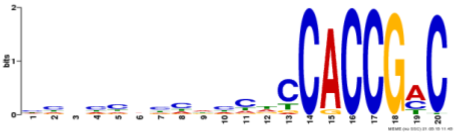 DREB 26 (O'Malley et al 2016) |

**Supplementary Data 2.**

Alignments of the DNA binding domains of Arabidopsis transcription factors.

Alignment of DNA binding domains of AtNLP6 and AtNLP7

```
AtNLP7        ----KKKTEKKRGKTEKTISLDVLQQYFTGSLKDAAKSLGVCPTTMKRICRQHGISRWPS
AtNLP6        EAKTVKKSERKRGKTEKTISLEVLQQYFAGSLKDAAKSLGVCPTTMKRICRQHGISRWPS

AtNLP7        RKIKKVNRSITKLKRVIESVQGTDGG
AtNLP6        RKINKVNRSLTRLKHVIDSVQGADGS
```

Alignment of DNA binding domains of AtARF9 and AtARF18 and AtARF2

```
AtARF9        FSKVLTASDTSTHGGFSVLRKHATECLPPLDMTQQTPTQELVAEDVHGYQWKFKHIFRGQ
AtARF18       FVKILTASDTSTHGGFSVLRKHATECLPSLDMTQATPTQELVTRDLHGFEWRFKHIFRGQ
AtARF2        FCKTLTASDTSTHGGFSVLRRHADECLPPLDMSRQPPTQELVAKDLHANEWRFRHIFRGQ

AtARF9        PRRHLLTTGWSTFVTSKRLVAGDTFVFLRGENGELRVGVRRAN
AtARF18       PRRHLLTTGWSTFVSSKRLVAGDAFVFLRGENGDLRVGVRRLA
AtARF2        PRRHLLQSGWSVFVSSKRLVAGDAFIFLRGENGELRVGVRRAM
```

Alignment of DNA binding domains of AtANAC032 and ATAF1

```
ANAC032       FPPGFRFHPTDEELVLMYLCRKCASQPIPAPIITELDLYRYDPWDLPDMALYGEKEWYFF
ATAF1         LPPGFRFHPTDEELVMHYLCRKCASQSIAVPIIAEIDLYKYDPWELPGLALYGEKEWYFF

ANAC032       SPRDRKYPNGSRPNRAAGTGYWKATGADKPIGRPKPVGIKKALVFYSGKPPNGEKTNWIM
ATAF1         SPRDRKYPNGSRPNRSAGSGYWKATGADKPIGLPKPVGIKKALVFYAGKAPKGEKTNWIM

ANAC032       HEYRLADVDRSVR-KKNSLRLDDWVLCRIYNK
ATAF1         HEYRLADVDRSVRKKKNSLRLDDWVLCRIYNK
```

**Supplementary Data 3.** *In vitro* binding assays showing relative binding of transcription factors to probes from target genes. Note: this dataset includes the three panels shown in Main Figure 2.

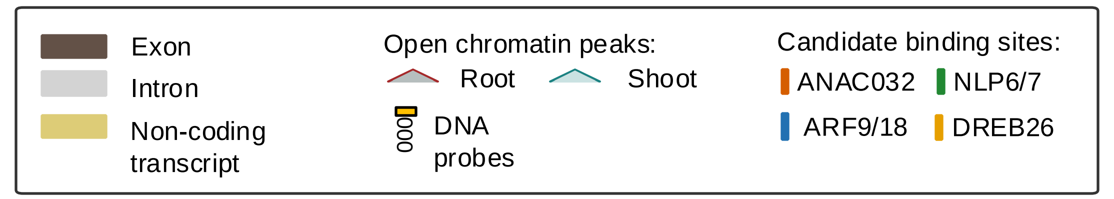

**Relative binding of AtDREB26, AtARF18 and AtANAC032 for candidate sites in *AtANAC032*.** Error bars = mean and standard deviation; n=3; P-values were calculated using an unpaired two-tailed Student's t-test of each sample to the random (rnd) control probe; *p<0.05, ** p<0.01, *** p<0.001; ns = not significant.

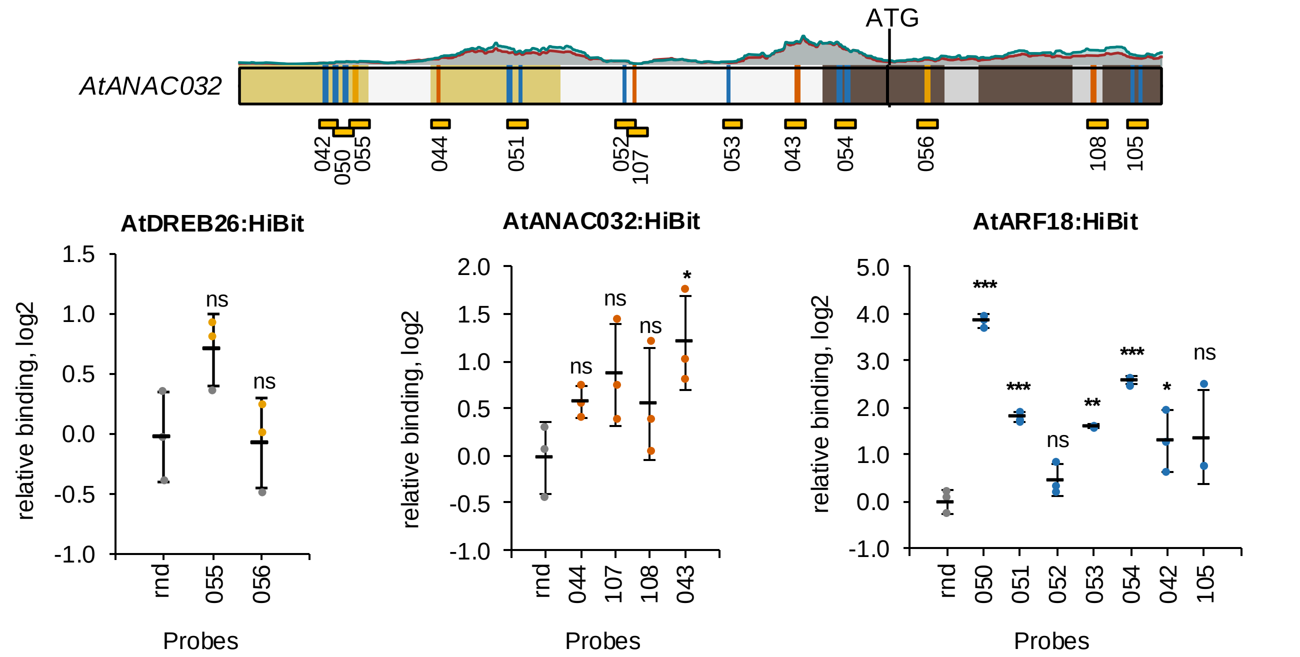

**Relative binding of AtNLP7, ATNLP6 and AtARF18 for candidate sites in *AtARF18*.** Error bars = mean and standard deviation; n=3; P-values were calculated using an unpaired two-tailed Student's t-test of each sample to the random (rnd) control probe; *p<0.05, ** p<0.01, *** p<0.001; ns = not significant.

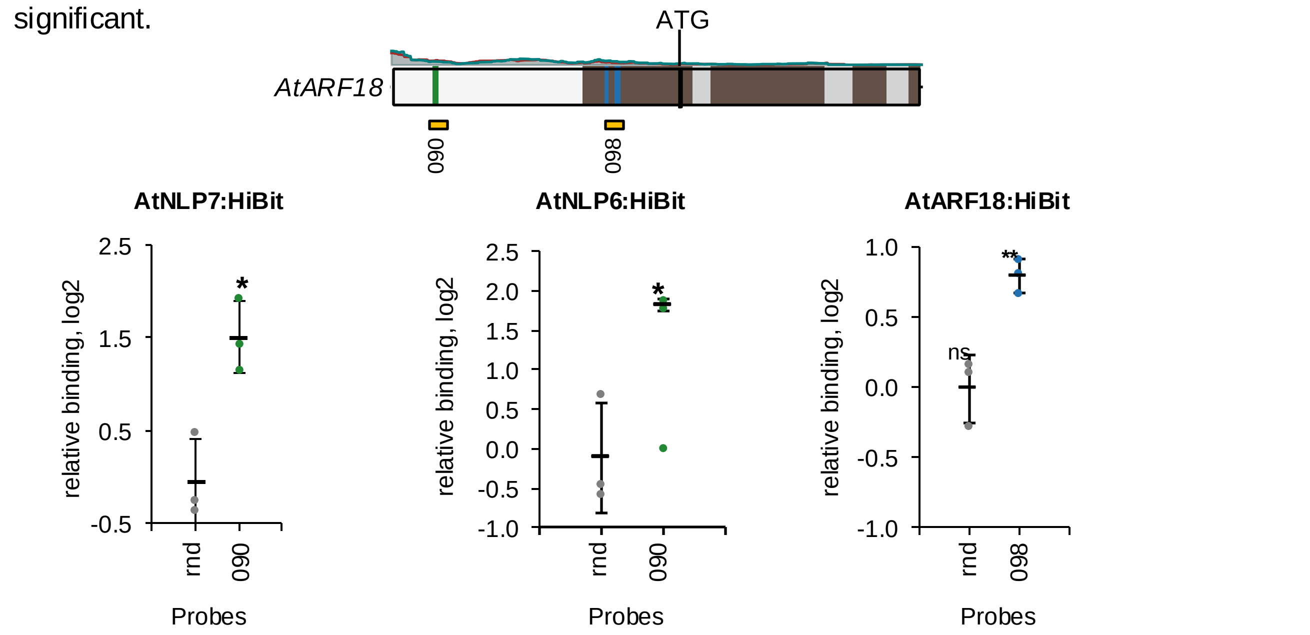

**Relative binding of AtNLP7, ATNLP6, AtDREB26, AtARF18 and AtANAC032 for candidate sites in *AtDREB26*.** Error bars = mean and standard deviation; n=3; P-values were calculated using an unpaired two-tailed Student's t-test of each sample to the random (rnd) control probe; *p<0.05, ** p<0.01, *** p<0.001; ns = not significant.

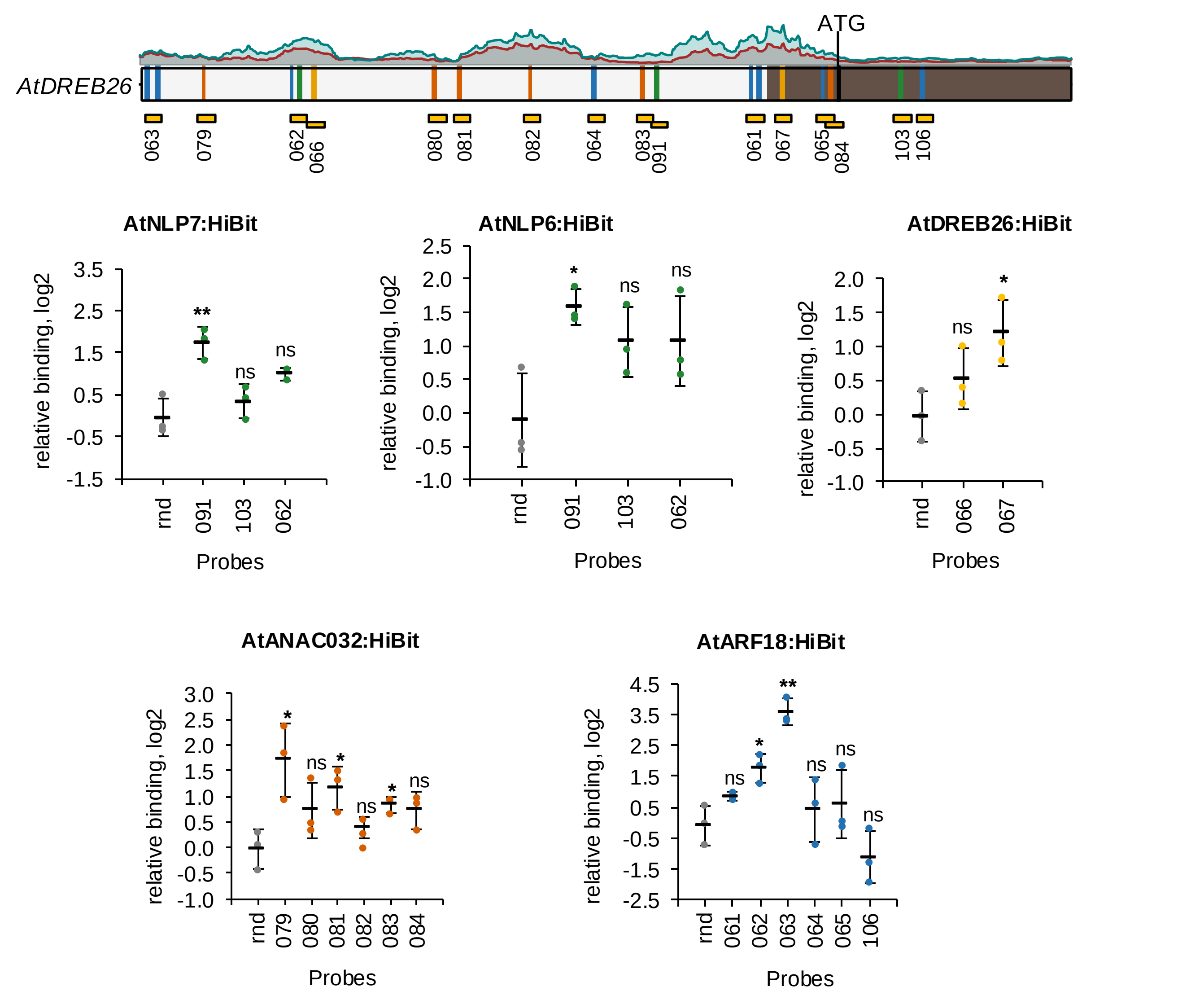

**Relative binding of AtNLP7, ATNLP6, AtDREB26, AtARF18 and AtANAC032 for candidate sites in *AtNIR1*.** Error bars = mean and standard deviation; n=3; P-values were calculated using an unpaired two-tailed Student's t-test of each sample to the random (rnd) control probe; *p<0.05, ** p<0.01, *** p<0.001; ns = not significant.

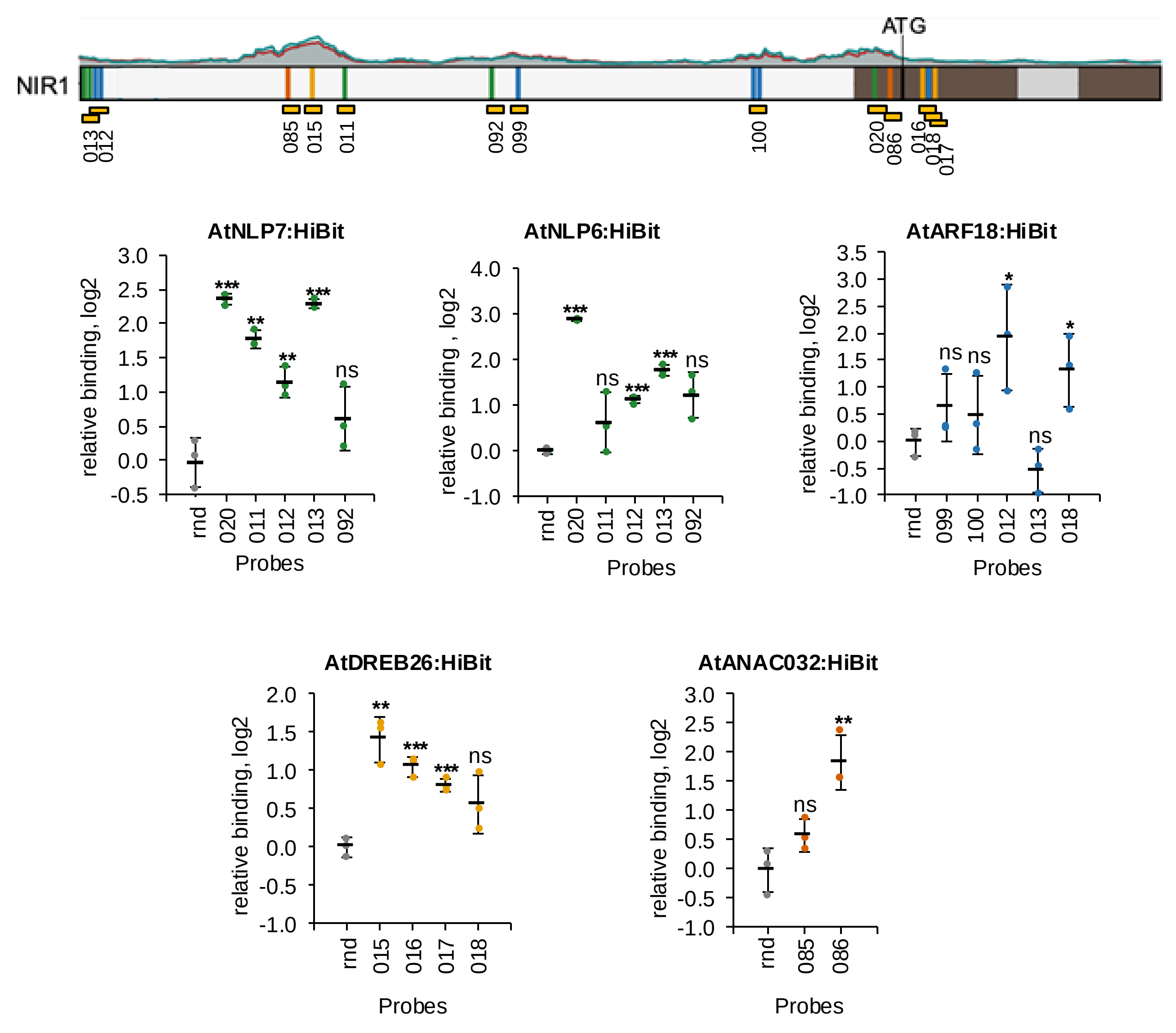

**Relative binding of AtNLP7, ATNLP6, AtDREB26, AtARF18 and AtANAC032 for candidate sites in *AtNLP6*.** Error bars = mean and standard deviation; n=3; P-values were calculated using an unpaired two-tailed Student's t-test of each sample to the random (rnd) control probe; *p<0.05, ** p<0.01, *** p<0.001; ns = not significant.

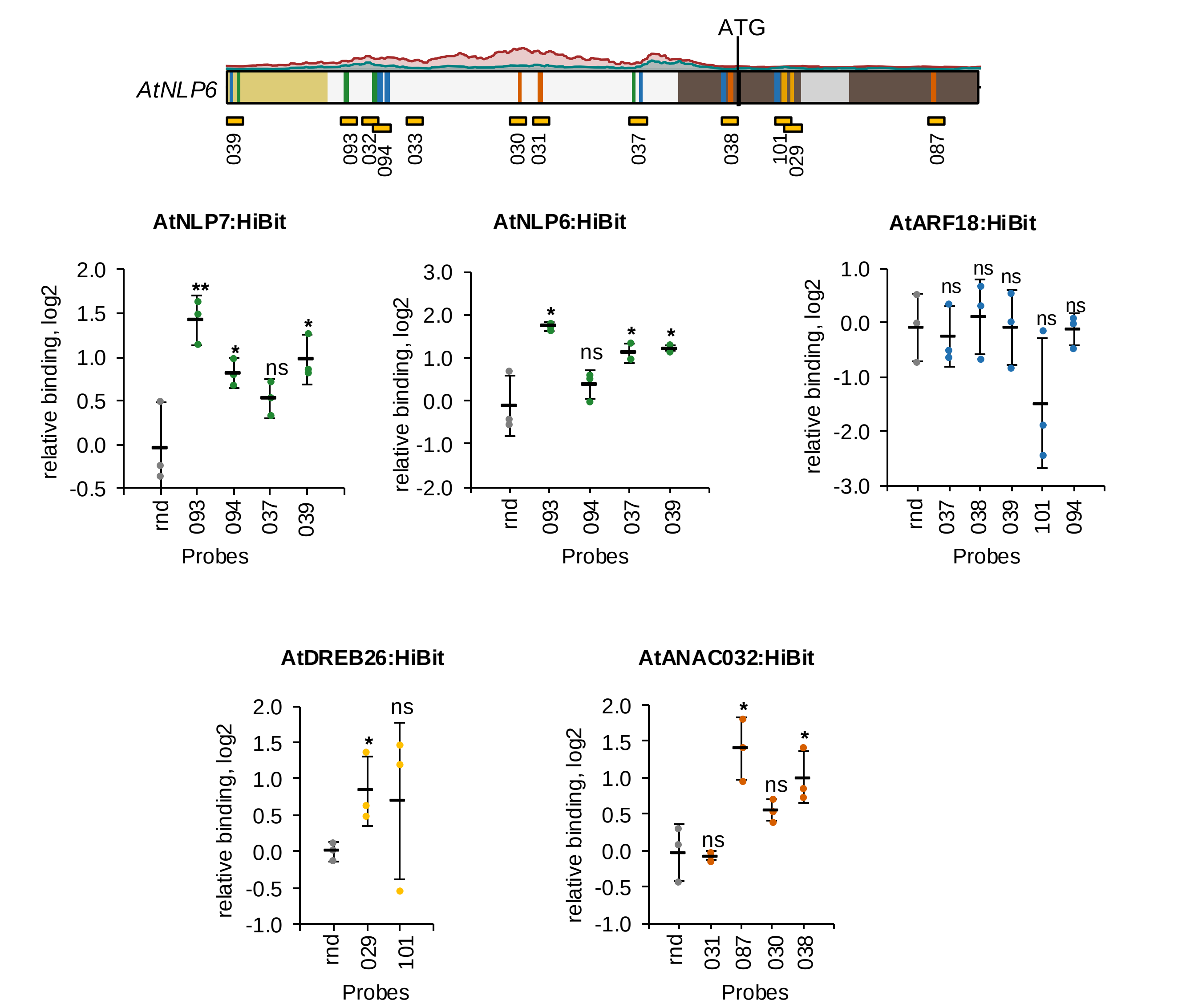

**Relative binding of AtNLP7, ATNLP6, AtDREB26, AtARF18 and AtANAC032 for candidate sites in *AtNLP7*.** Error bars = mean and standard deviation; n=3; P-values were calculated using an unpaired two-tailed Student's t-test of each sample to the random (rnd) control probe; *p<0.05, ** p<0.01, *** p<0.001; ns = not significant.

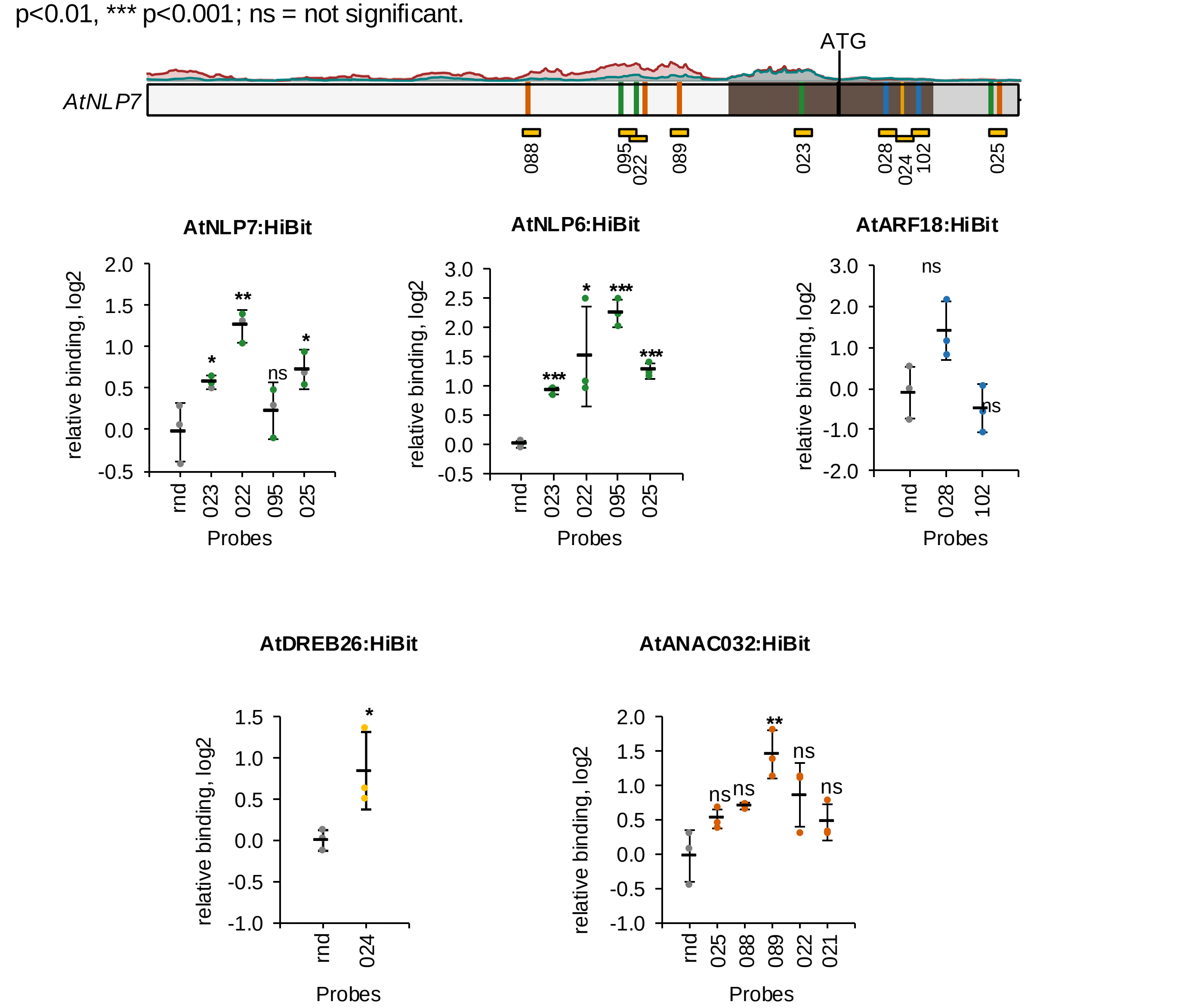

### Supplemetary Data 4

#### Sequences of double-stranded DNA probes used in in vitro binding assays

| Species | Promoter | robe II | Sequence (forward strand) | |
|---|---|---|---|---|
| Synthetic | n/a (random seqeunce) | Rnd | TAGCGAAGTACGATCCCATGAAGACGCTGGGTTTACATGGGAATGGTGCTTCTGTTCTAACAGGCTAGGATATAAGGCCATCACGCAGTA | |
| Synthetic | Synthetic (DAP-seq consensus) - positive control for DREB26 binding | 69 | TAGCGAAGTACGATCCCATGAAGACGCTGGGTccaccaccacctCCACCGACACAGGCTAGGATATAAGGCCATCACGCAGTA | (O'Malley et al.,2016) |
| Arabidopsis | NRE element from NIR1 - positive control for NLP6/7 binding | 20 | TAGCGAAGTACGATCCCATCAAAGAGAAACAACTTGACCCTTTACATTGCTCAAGAGCTCATCTCTTCCCTCTACGGCCATCACGCAGTA | (Konishi and Yanagisawa 2013) |
| Arabidopsis | ANAC032SO1 - positive control for ANAC032 binding | 71 | TAGCGAAGTACGATCCCATGAAGACGGAGGTAAGCAAATTGATCACGCAACTGGTGGATATAAGGCCATCACGCAGTA | (Allu et al., 2016) |
| Arabidopsis | DR5 (positive control for ARF9/19 binding) | 74 | CCTTTTGTCTCCCTTTTGTCTCCCTTTTGTCTCCCTTTTGTCTCCCTTTTGTCTCCCTTTTGTCTCCCTTTTGTCTC | (Liu et al., 2015) |
| Arabidopsis | ANAC032 | 42 | TAGCGAAGTACGATCCCCAAGTGTTCCAGATTTCTGGTTTGTCGCAGTCTTTAGTTTTCAAGGTTGGCCATCACGCAGTA | |
| Arabidopsis | ANAC032 | 43 | TAGCGAAGTACGATCCCGGACCGCTACATTCCAAATAGTCTGACGTAAGCAATGACAAAACTCACCTACATGGCCATCACGCAGTA | |
| Arabidopsis | ANAC032 | 44 | TAGCGAAGTACGATCCCGTTAGATCGAGCCAGAGAGGCAAAATCCATACGCATAAGCGTTTTGATGATTGGCCATCACGCAGTA | |
| Arabidopsis | ANAC032 | 50 | TAGCGAAGTACGATCCCGTTGGTACTGATCAGTGACACGTTTTGTTGTTTGTCTTCTTAGTGTTCATGTTCGGGCCATCACGCAGTA | |
| Arabidopsis | ANAC032 | 51 | TAGCGAAGTACGATCCCTTGATTATGAACCATCCACATCATGTCACGTGTCTAAACTCTTCGGTCACGGCCATCACGCAGTA | |
| Arabidopsis | ANAC032 | 52 | TAGCGAAGTACGATCCCTGGTTATAAAGAAAGAAACTTAATTTGTCTCGAATTTGAGTTTGGATGCGACGGCCATCACGCAGTA | |
| Arabidopsis | ANAC032 | 53 | TAGCGAAGTACGATCCCTGAACAAAGAAAAAAATAGACATTAAAAATAGTAAGATCGAAAATACGGCCATCACGCAGTA | |

| | | | |
|---|---|---|---|
| Arabidopsis | ANAC032 | 54 | TAGCGAAGTACGATCCCCCAACACAAAGCCCCTGTCTATCCGCCATGTGTCCACGACCTTTCTTATAGGCCATCACGCAGTA |
| Arabidopsis | ANAC032 | 55 | TAGCGAAGTACGATCCCTCATGTTCGATTCACTATGGGTGACTTTAGAAAGACGTTAACATGTCAACGGGCCATCACGCAGTA |
| Arabidopsis | ANAC032 | 56 | TAGCGAAGTACGATCCCTAAATGCGCGTCGCAGCCGATCCCTGCTCCGATTATCACCGAACTCGATTTGGCCATCACGCAGTA |
| Arabidopsis | ANAC032 | 105 | TAGCGAAGTACGATCCCCGGGTTAAAGCCTGTGACTGACACGTGTCCACCGGAATCTGTGGCGAGATGGCCATCACGCAGTA |
| Arabidopsis | ANAC032 | 107 | TAGCGAAGTACGATCCCGAGTTTGGATGCGACGTTTTACGTACTAGTTTTATAAAAGTAAATGGGCCATCACGCAGTA |
| Arabidopsis | ANAC032 | 108 | TAGCGAAGTACGATCCCAATCGGTTTAGTTAACCGGACGTGTTTTTGATATTCTTGAACAGTTGGCCATCACGCAGTA |
| Arabidopsis | ARF18 | 90 | TAGCGAAGTACGATCCCAATTCCCAAAACGCCGCGTGACCCTTCGTTTGCCTTTGAATACAATGGCCATCACGCAGTA |
| Arabidopsis | ARF18 | 98 | TAGCGAAGTACGATCCCCTGCAATCAATACGGACATAACCGTCCGTTGTGTCCTGTTTATAAAGTGGCCATCACGCAGTA |
| Arabidopsis | DREB26 | 61 | TAGCGAAGTACGATCCCGAAGAAAAACACCTGTCATGCACCACAATGTCACATTCATACACAAGAAAGGCCATCACGCAGTA |
| Arabidopsis | DREB26 | 62 | TAGCGAAGTACGATCCCGTTTGGATTTGGGTAGCAAAAGAGACAGAGTGAGAACAGCTGTAATTTGGGCCATCACGCAGTA |
| Arabidopsis | DREB26 | 63 | TAGCGAAGTACGATCCCGAGAAAGTCAAGGCTGTCCACAAAGAACATGTCATCTCCCATCTTTTCTTGGCCATCACGCAGTA |
| Arabidopsis | DREB26 | 64 | TAGCGAAGTACGATCCCGTTCACTCGGTTTATATTTTGGACAAATTCTAGAAAGAATCTAACACGGCCATCACGCAGTA |
| Arabidopsis | DREB26 | 65 | TAGCGAAGTACGATCCCCTCCACTTGATCACTACCACATTGTCCAATCACTCTACAAAGCCTGTACGGGCCATCACGCAGTA |
| Arabidopsis | DREB26 | 66 | TAGCGAAGTACGATCCCATCAATTTAGCATCATCACCTTCTCCCACACTCTTCTGACTTCACTCTTCGCGGCCATCACGCAGTA |

| | | | |
|---|---|---|---|
| Arabidopsis | DREB26 | 67 | TAGCGAAGTACGATCCCTAAATAATATTTCATCTCTTCCTCTCCCATACCCCTCCCAAATCAATCCATCGGCCATCACGCAGTA |
| Arabidopsis | DREB26 | 79 | TAGCGAAGTACGATCCCTATGTTGGTATATATTAAAGTTACGTAAGACAAAATTTATTAGAATTCGGGCCATCACGCAGTA |
| Arabidopsis | DREB26 | 80 | TAGCGAAGTACGATCCCCTTTAGACTTTGATGATACACGTAACAAATCTAATAAAAAGATTAGATGGGCCATCACGCAGTA |
| Arabidopsis | DREB26 | 81 | TAGCGAAGTACGATCCCGTAATAACATTTTCGTGCACGTATGTTTGCAGGTTTTACAGTCACGGCCATCACGCAGTA |
| Arabidopsis | DREB26 | 82 | TAGCGAAGTACGATCCCCTCCTGATAGTGATGATAGGTACGTATATAGTCAATATATAAAAGGGCCATCACGCAGTA |
| Arabidopsis | DREB26 | 83 | TAGCGAAGTACGATCCCACATGCAATTGATAAAAATTTACGTGATCGCATAATTAATTGTTGGGCCATCACGCAGTA |
| Arabidopsis | DREB26 | 84 | TAGCGAAGTACGATCCCCACTCTACAAAGCCTGTACGTACACAACAACATTACCATGGTGGGCCATCACGCAGTA |
| Arabidopsis | DREB26 | 91 | TAGCGAAGTACGATCCCTTGTGGTTCCTCAATAAAAAAAGCCAATCAAAATCATTATACGTTGGCCATCACGCAGTA |
| Arabidopsis | DREB26 | 103 | TAGCGAAGTACGATCCCTGAGATTAGGGCACCAAATCAAAAGACAAGGATTTGGTTAGGTTCTTACGGCCATCACGCAGTA |
| Arabidopsis | DREB26 | 106 | TAGCGAAGTACGATCCCAGCTTACGATGTTGCACTCTTATGTCTCAAAGGCCCTCAAGCCAATCTCAGGCCATCACGCAGTA |
| Arabidopsis | NIR1 | 11 | TAGCGAAGTACGATCCCTGTCCTTAAGATTTTAAAAGCTACAAGAGCCACTAGCTAtttttCAATTCCAAGGCCATCACGCAGTA |
| Arabidopsis | NIR1 | 12 | TAGCGAAGTACGATCCCCAAATCAGaaaaaCAAAACCCAAAAGACATGACAACCCTTTAATTATTGGCCATCACGCAGTA |
| Arabidopsis | NIR1 | 13 | TAGCGAAGTACGATCCCAAGACATGACAACaaaaaGTCCAAAAGACAAGTTATCCAAATCAGaaaGGCCATCACGCAGTA |

| | | | |
|---|---|---|---|
| Arabidopsis | NIR1 | 15 | TAGCGAAGTACGATCCCGAAGCCACACGTGGAATAGCGGTGGAGAATAATTTGGATGGATGGCGGTTTAGGCGGCCATCACGCAGTA |
| Arabidopsis | NIR1 | 16 | TAGCGAAGTACGATCCCCACCAAACCCAAAAGATCCGTCCTTGTCGCCGCCGCTCAGACCACAGGCCATCACGCAGTA |
| Arabidopsis | NIR1 | 17 | TAGCGAAGTACGATCCCCTCAGACCACAGCTCCGGCCGAATCCACCGCCTCTGTTGACGCAGGGCCATCACGCAGTA |
| Arabidopsis | NIR1 | 18 | TAGCGAAGTACGATCCCAAGATCCGTCCTTGTCGCCGCCGCTCAGACCACAGCTCCGGCCGAATCCACCGCCTGGCCATCACGCAGTA |
| Arabidopsis | NIR1 | 20 | TAGCGAAGTACGATCCCATCAAAGAGAAACAACTTGACCCTTTACATTGCTCAAGAGCTCATCTCTTCCCTCTACGGCCATCACGCAGTA |
| Arabidopsis | NIR1 | 85 | TAGCGAAGTACGATCCCTCACGTGGTCAATTATTTACGTCGAAGATGTGATAGATATTTAGTAGGCCATCACGCAGTA |
| Arabidopsis | NIR1 | 86 | TAGCGAAGTACGATCCCCCCTCTACAAAAATGGCCGCACGTCTCCAACCTTCTCCCAACTCCGGCCATCACGCAGTA |
| Arabidopsis | NIR1 | 92 | TAGCGAAGTACGATCCCTATAAATTAGAAAAAAAAAGACAATGCTAACGTTTTGGTGTGGAGGCCATCACGCAGTA |
| Arabidopsis | NIR1 | 99 | TAGCGAAGTACGATCCCGAACCTGCAGGTAGTTTGATTCATGTCTCTAGAATAACGCTAAAGAACACGGCCATCACGCAGTA |
| Arabidopsis | NIR1 | 100 | TAGCGAAGTACGATCCCGAAGTGTATGGTCGGTGTCAATAACTAATATGTCATATTTTTTACAGACTGGCCATCACGCAGTA |
| Arabidopsis | NLP6 | 29 | TAGCGAAGTACGATCCCTTTACTCCGCCGCAGTAGCTCCCACTCGCTTCACCGATTATTCCGTCTTACTCGCTGGCCATCACGCAGTA |
| Arabidopsis | NLP6 | 30 | TAGCGAAGTACGATCCCAACACGCACTTAGTGCCTAATTACGTCTTAATTTAATAttttttAAAGCGGCCATCACGCAGTA |
| Arabidopsis | NLP6 | 31 | TAGCGAAGTACGATCCCTTTGTTCATGGTGGTGGACGTGTAAGAAACGTCTAAAGATCTACTCTATATAACGGCCATCACGCAGTA |

| | | | |
|---|---|---|---|
| Arabidopsis | NLP6 | 32 | TAGCGAAGTACGATCCCTTTGATGAACAACCTAGTCCAAATATGGAAGAAGGAAGGAAGGATTGTCATTAGGCCATCACGCAGTA |
| Arabidopsis | NLP6 | 33 | TAGCGAAGTACGATCCCAATATTGATTAACAGCTAAATAACAAATTAGGGAAGTTATTATAACATTAAGGCCATCACGCAGTA |
| Arabidopsis | NLP6 | 37 | TAGCGAAGTACGATCCCGAAAAACAAAATTTCGATGCCTTTTGTCTCATAAAAATATTAAAGAAAATGGCCATCACGCAGTA |
| Arabidopsis | NLP6 | 38 | TAGCGAAGTACGATCCCAACACTTACTCTGACGAGTTTTTGTCTCGTCTTACGTCTTTGCTGCTGCGGCCATCACGCAGTA |
| Arabidopsis | NLP6 | 39 | TAGCGAAGTACGATCCCGATTCGACTTTTCTCACTGCTTGTCTCTTCTCTTCTGTTATGTTCCTGGGGCCATCACGCAGTA |
| Arabidopsis | NLP6 | 87 | TAGCGAAGTACGATCCCAATACAGGATGGTTTCATTGACGTATATGTTTTCCCTTGATGGTGGCCATCACGCAGTA |
| Arabidopsis | NLP6 | 93 | TAGCGAAGTACGATCCCTGGTTTAGTTGTGGATGAAAAGGACAGATTTGGAGAGCCTCTAACGTAGTGGCCATCACGCAGTA |
| Arabidopsis | NLP6 | 94 | TAGCGAAGTACGATCCCGAAGGATTGTCATTACAGAACAAGGGACACACAATTGTCAATTGCCAAACACGGCCATCACGCAGTA |
| Arabidopsis | NLP6 | 101 | TAGCGAAGTACGATCCCTCTCCGAAACTTCTGGAGATGTCGGCGGCGAGCTTTACTCCGCCGCAGGGCCATCACGCAGTA |
| Arabidopsis | NLP7 | 22 | TAGCGAAGTACGATCCCTTCTCAATTTCCCATTAATAATCCCGAAAAGGCATTACGTAATAATTAATATTTTACGGGCCATCACGCAGTA |
| Arabidopsis | NLP7 | 23 | TAGCGAAGTACGATCCCGAATCACACACTCTGCCAATTTGACCTTTCTCTCTCGCTTCTCTTTCTCTATCCAGGCCATCACGCAGTA |
| Arabidopsis | NLP7 | 24 | TAGCGAAGTACGATCCCCTGGGCTTTCTCCGACGGTGGAGGAAATGGTTTTCACCACGCAACCTCCGGTGGCGGGCCATCACGCAGTA |
| Arabidopsis | NLP7 | 25 | TAGCGAAGTACGATCCCTGTTTGTCCCTTCTTAGGAGGTTATGTCAATTACGTTATTTAGGTCTCTTTCCTTGGCCATCACGCAGTA |

| | | | |
|---|---|---|---|
| Arabidopsis | NLP7 | 26 | TAGCGAAGTACGATCCCACTTCATGCttttttATATTTCTCCTTTTGGACATCTAACATGTTTATAATTGGCCATCACGCAGTA |
| Arabidopsis | NLP7 | 27 | TAGCGAAGTACGATCCCTGGTAAAATCATAAAGTTGCCaaaaaTAGTTAAGTTCTATAAAAGGAGTGGGGGCCATCACGCAGTA |
| Arabidopsis | NLP7 | 28 | TAGCGAAGTACGATCCCGAATCGCATGATTTCTCCGATTTTTGTCTCCTCTTCCTCTGAGCAGCCTTGCGGCCATCACGCAGTA |
| Arabidopsis | NLP7 | 88 | TAGCGAAGTACGATCCCAAGCAACTAATTTTGGACGTACGAAATGCTTTGTAAATACATTGAGGCCATCACGCAGTA |
| Arabidopsis | NLP7 | 89 | TAGCGAAGTACGATCCCATTCCCACCACGAGTTTGCTTAACGTGTCGTTAAATGCAGCCGTCGGGGCCATCACGCAGTA |
| Arabidopsis | NLP7 | 95 | TAGCGAAGTACGATCCCGAATCAGGTAGGTCCAGGATTTGTCTTTTCTCCACTTCTCAATTTCCCAGGCCATCACGCAGTA |
| Arabidopsis | NLP7 | 102 | TAGCGAAGTACGATCCCCGATGATGAGAAGATCAGCTCTGTCTCCGGTGTTCCTTCTTTCCGTCTGGCCATCACGCAGTA |
| Tomato | SlNIR1 | S01 | TAGCGAAGTACGATCCCCTTATTTTAACATTAACGACCCTTTGCAACAATCAAGAGTCCACTAAACGTTTTGGGCCATCACGCAGTA |
| Tomato | SlNIR1 | S02 | TAGCGAAGTACGATCCCGACCAATACAAGGACTCTTGGCTCTTCTTGCTAACCTGCTTTTTCCTGTCTCTGAAATGGCCATCACGCAGTA |
| Tomato | SlNIR2 | S03 | TAGCGAAGTACGATCCCGTAATTTAGAAACAATAACGACCCTTTGGAACTTCCAAGAGTCCAAAAGCACCGTACGGCCATCACGCAGTA |
| Tomato | SlNIR2 | S04 | TAGCGAAGTACGATCCCGTATGCTGTAGCCTTAATCAAAAGGCCACACGTCTCTTTCCTTGACCCATGGGCCATCACGCAGTA |
| Tomato | SlNLP7-1 | S05 | TAGCGAAGTACGATCCCCTTCGTCATCTTTCTCTTCTCGCCACATACATAATAGAGAGAGAAAATCTCGGCCATCACGCAGTA |
| Tomato | SlNLP7-1 | S06 | TAGCGAAGTACGATCCCAGGACTTTAGTATGTTCTTATTTGTTTGATGTAAAATACACTTAAGCTTGGGCCATCACGCAGTA |

| | | | |
|---|---|---|---|
| Tomato | SlNLP7-1 | S07 | TAGCGAAGTACGATCCCTTTGCAATGAGTAATGACACATCTTTGGAGACATCTTTTGATCCCAACTCAGGCCATCACGCAGTA |
| Tomato | SlNLP7-3 | S08 | TAGCGAAGTACGATCCCCAAGTTTGGGGTTCATTCGTTTTCGGCGAAAAATTATGTTATTTTTACATAGGGCCATCACGCAGTA |
| Tomato | SlNLP7-3 | S09 | TAGCGAAGTACGATCCCATTTCCATATGTCATCGATTTATTATAATTGAGGATGTCTTTTTCGACCAACGGCCATCACGCAGTA |
| Tomato | SlNLP7-3 | S10 | TAGCGAAGTACGATCCCTAAATTTTTATTCTTTTTTTCTAAAAAATATAATTTCATATGAATAAATTGGCCATCACGCAGTA |
| Tomato | SlNLP7-3 | S11 | TAGCGAAGTACGATCCCCAACCTATGTTTGAAATCTCAGAGACACACTTGTACTATACTAAGGTCCGGCCATCACGCAGTA |
| Tomato | SlNLP7-3 | S12 | TAGCGAAGTACGATCCCATTTGATATGATTAGGACAGTAATTATAGAGCTAGTGACACTAAGATTCCGGCCATCACGCAGTA |
| Tomato | SlDREB26 | S13 | TAGCGAAGTACGATCCCCAACATCACTTCATCAATATCCATCTCTAAAACATAAACACACACAAAAGGCCATCACGCAGTA |
| Tomato | SlDREB26 | S14 | TAGCGAAGTACGATCCCCTGGAAAATGTCTGTCCTCCACTCTGATGTTTACATGTAGACATTATACTCGGCCATCACGCAGTA |
| Tomato | SlDREB26 | S15 | TAGCGAAGTACGATCCCAGCTAAGGACAAAATTGTCATTTGGATGGATAGTAAAAGGACATTTATCCGGCCATCACGCAGTA |
| Tomato | SlDREB26 | S16 | TAGCGAAGTACGATCCCGAAATCACATATTTTTGACACAAAATTATCATTTGTCTTTTATAAGTTTCTAGGCCATCACGCAGTA |
| Tomato | SlDREB26 | S17 | TAGCGAAGTACGATCCCAGGTATAAATGAGACAGGAGCACATGATAGTTTTGACATAGTCTTTATCGGCCATCACGCAGTA |

**Supplementary Data 5. Schematic showing the methodology for protoplast co-expression assays.** To assess changes in luminescence resulting from each of the promoters tested (pTest) in response to the expression of a given transcription factor (TF), luminescence values relative to an experiment calibrator are compared from samples with and without plasmids expressing each TF. To maintain equal transcriptional loads, in the absence of the TF, a control plasmid expressing YFP from the same promoter is included. The relative value for pTEST is then normalized to a batch calibrator to account for variation between protoplast batches providing a normalized value in arbitrary units (a.u.).

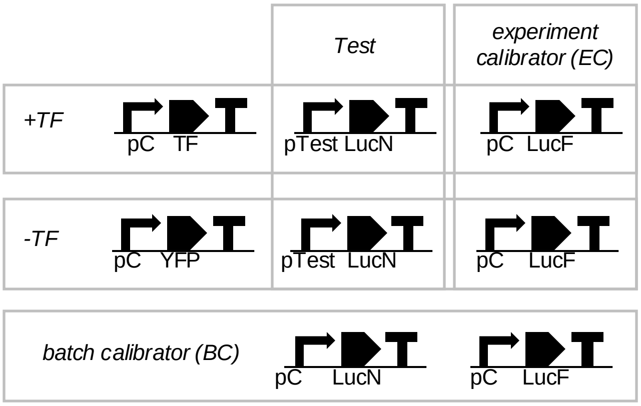

$$\textit{Normalized luminescence (a.u.)} = \frac{Test_{LucN}/EC_{LucF}}{BC_{LucN}/BC_{LucF}}$$

Appropriate pCs for each experiment were determined by evaluating responses to each TF, selecting the promoter that did not show a significant change in expression following co-expression. The identities of pC in each experiment are as follows:

| **TF** | **pC** |
|---|---|
| **AtANAC032** | **pCaMV35s** |
| **AtARF18** | **pAtuNos** |
| **AtDREB26** | **pAtuNos** |
| **AtNLP6** | **pCaMV35s** |
| **AtNLP7** | **pCaMV35s** |

**Supplementary Data 6. Expression levels of Arabidopsis genes in modified TARGET assays**. Values represent the mean and standard error of three biological replicates of which each is the mean of two technical replicates. P-values were calculated using an unpaired two-tailed Student's t-test, *P<0.05, ** P<0.01, *** P<0.001, and **** P<0.0001. Values represent the mean and standard error of three biological replicates (independent transfections) of which each is the mean of two technical replicates (qPCR assays).

**Expression of *AtANAC032* is not affected by 35sP-AtDREB26:GR**

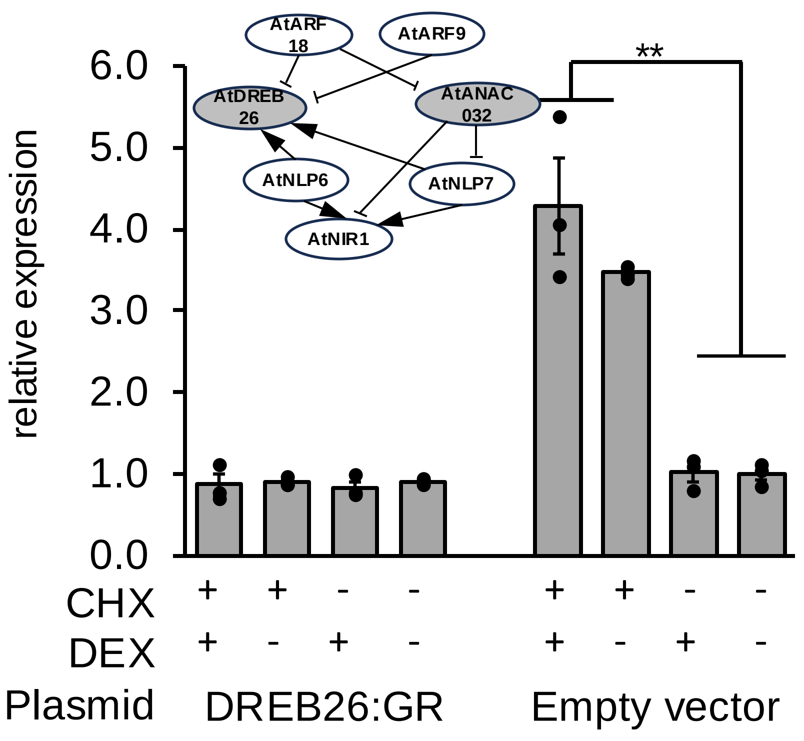

**Expression of *AtNIR1* is directly repressed by *35sP*-AtANAC032:GR**

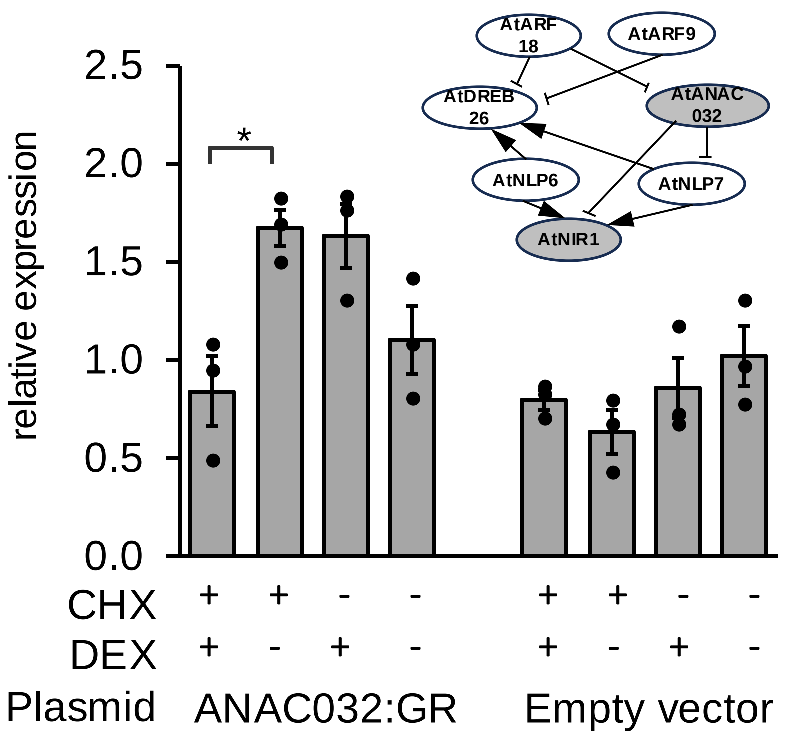

**Expression of *AtNIR1* is not affected by *35sP*-AtARF9:GR**

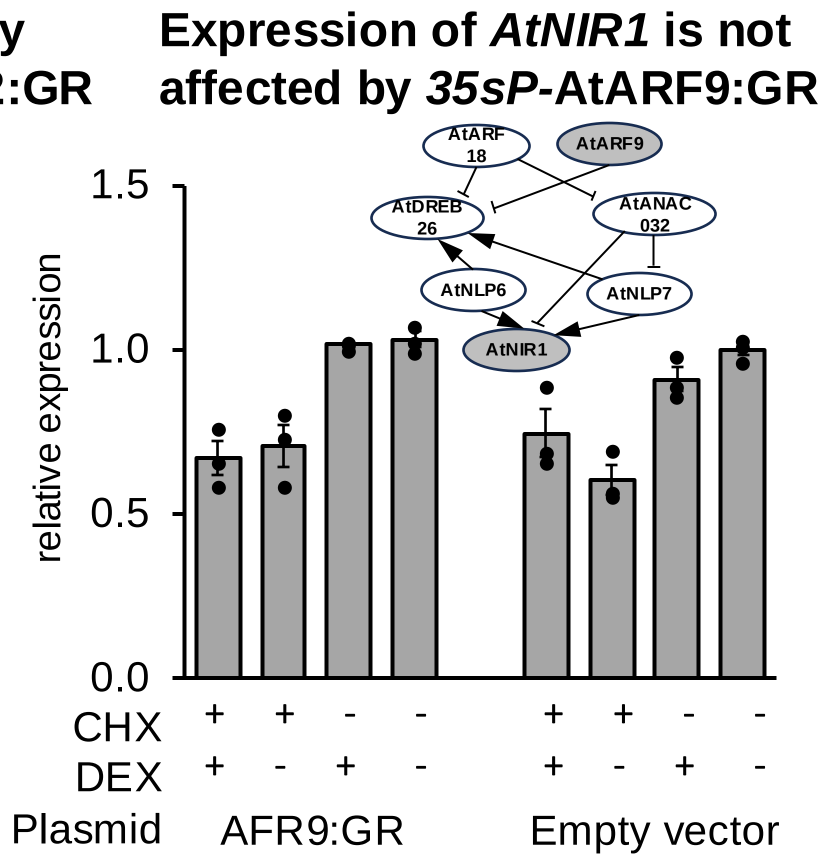

**Expression of *AtNIR1* is indirectly repressed by *35sP*-AtARF18:GR**

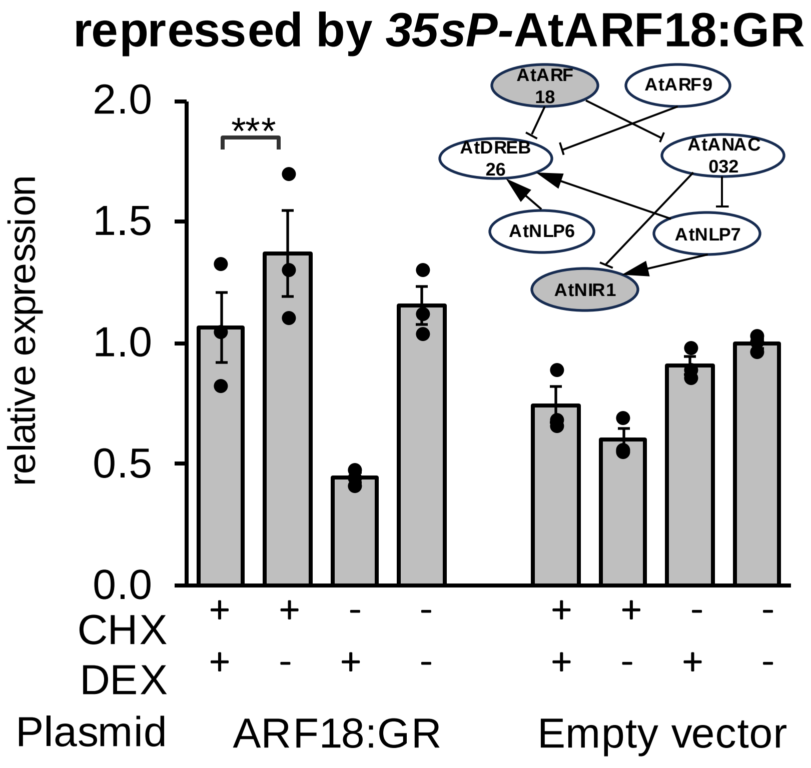

**Expression of *AtNIR1* is indirectly repressed by *35sP*-AtDREB26P:GR**

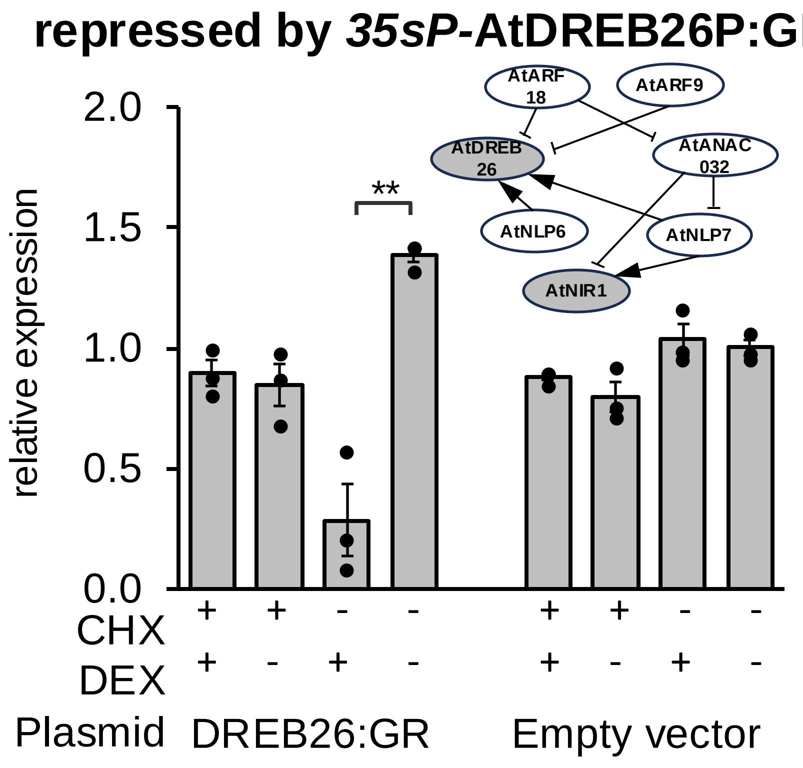

**Expression of *AtNIR1* is not affected by *35sP*-AtNLP6:GR**

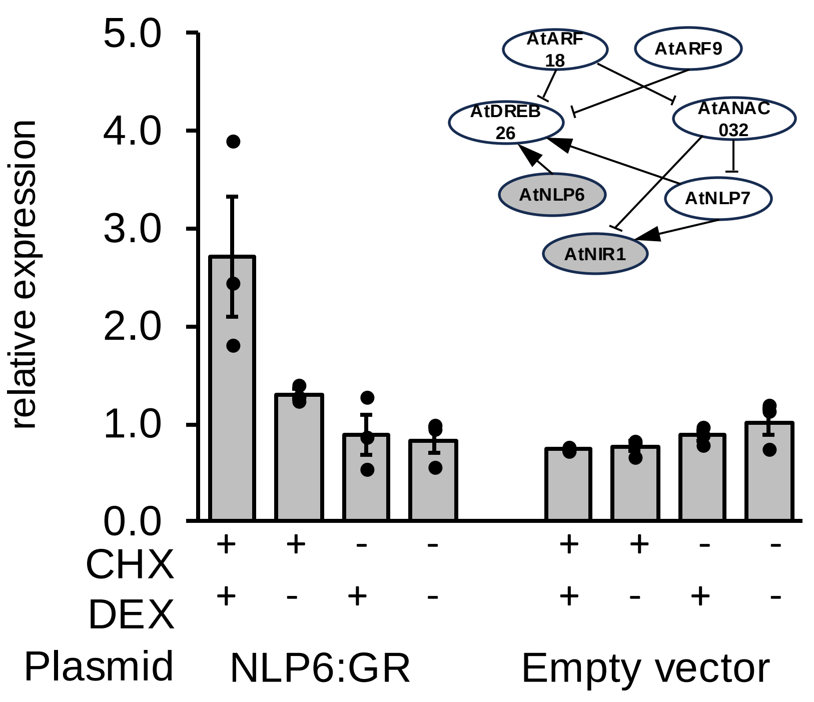

**Expression of *AtNLP6* not affected by *35sP*-AtDREB26:GR**

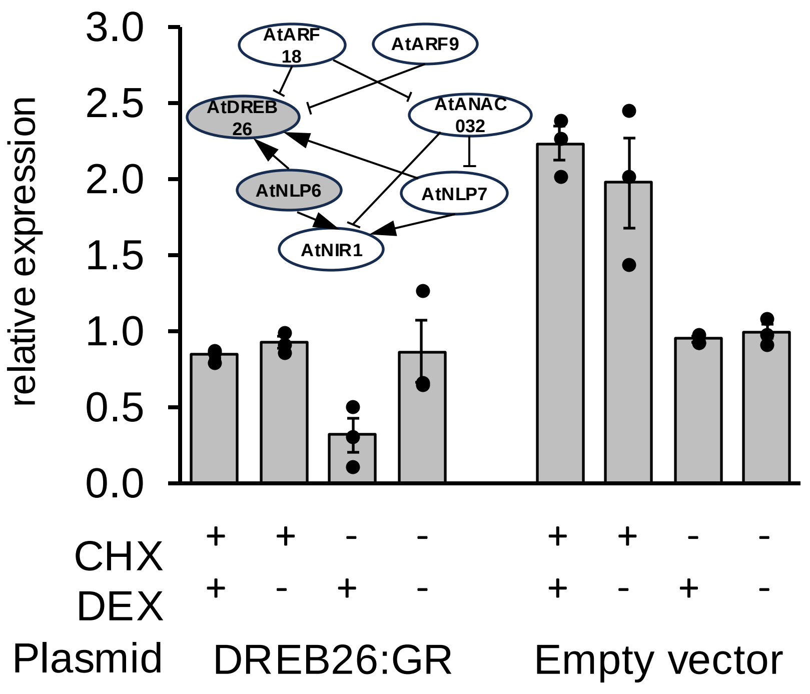

**Expression of *AtNLP6* is not affected by AtANAC032:GR**

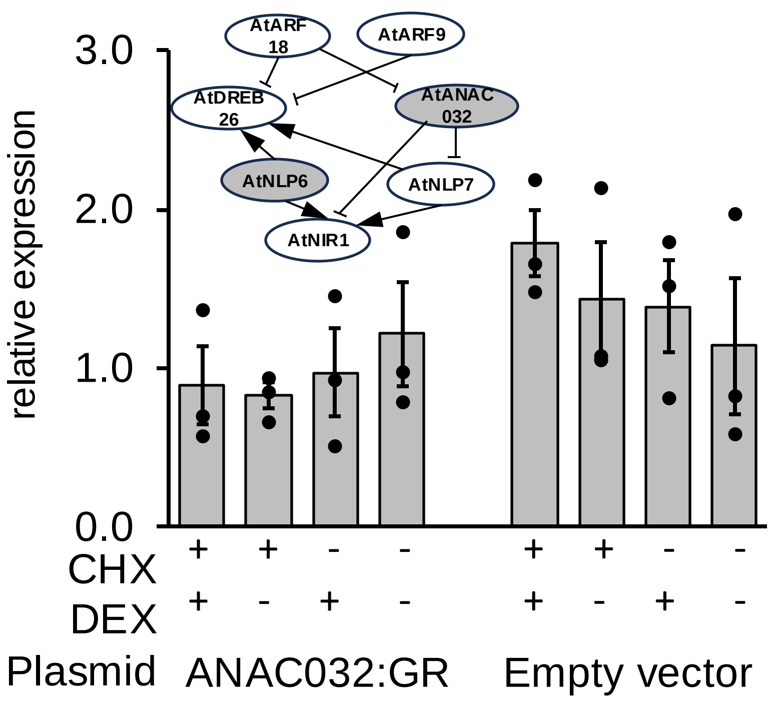

**Expression of *AtNLP6* is indirectly repressed by *35sP*-AtARF18:GR**

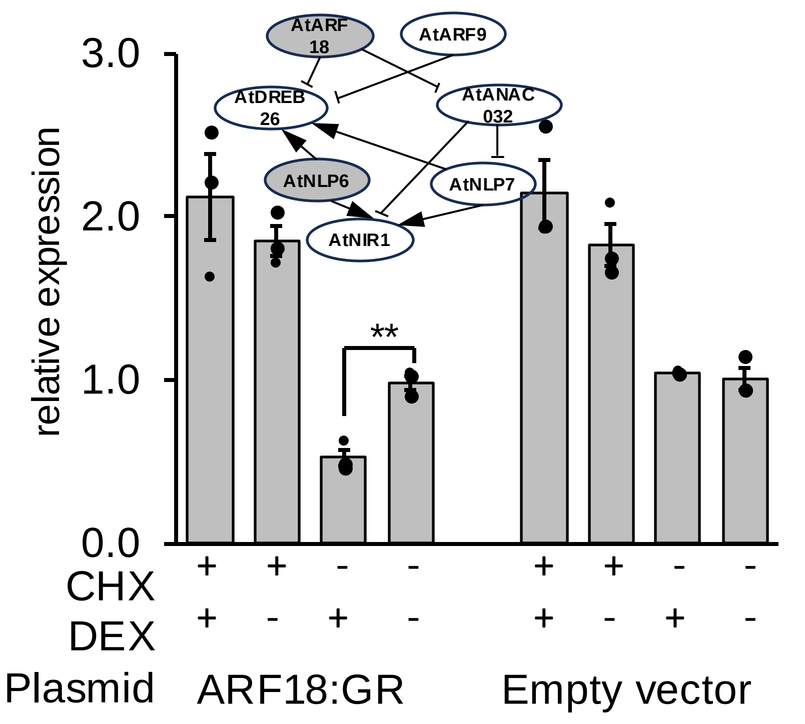

**Expression of *AtNLP6* is not affected by *35sP*-AtARF9:GR**

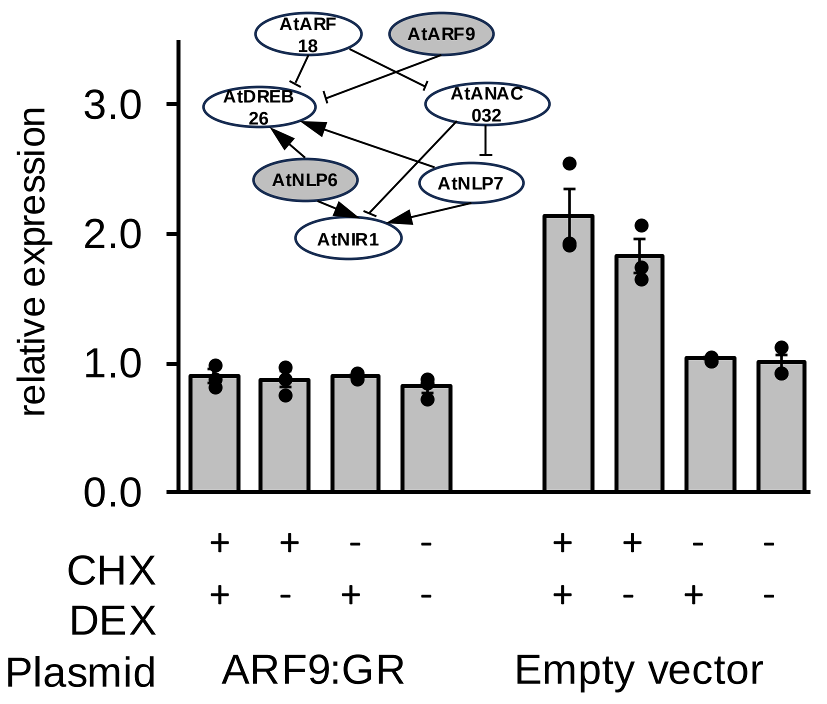

**Expression of *AtNLP7* is indirectly repressed by *35sP*-AtNLP6:GR**

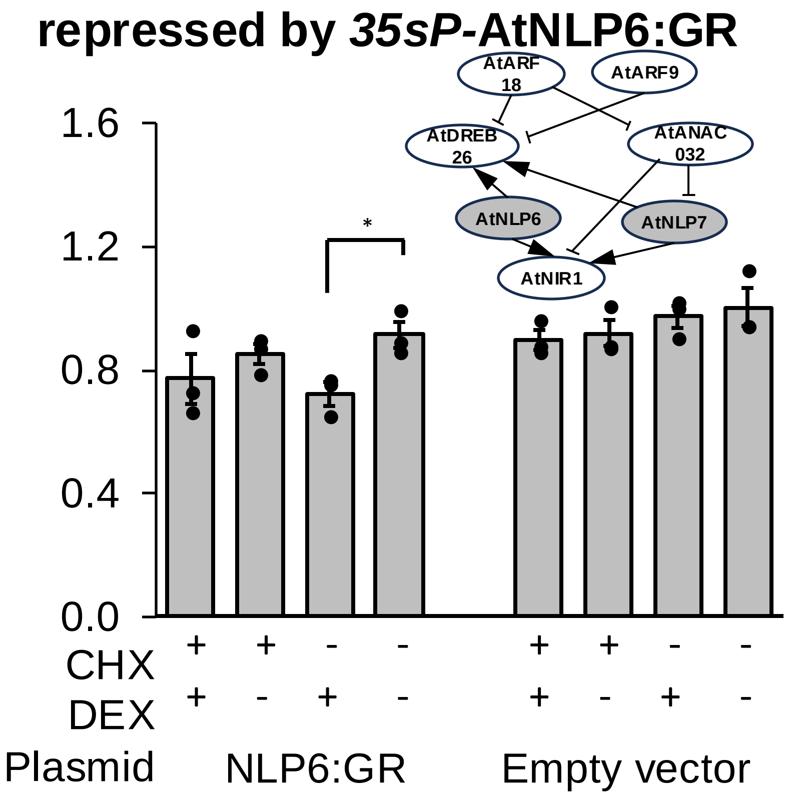

**Expression of *AtNLP7* is indirectly repressed by *35sP*-AtDREB26:GR**

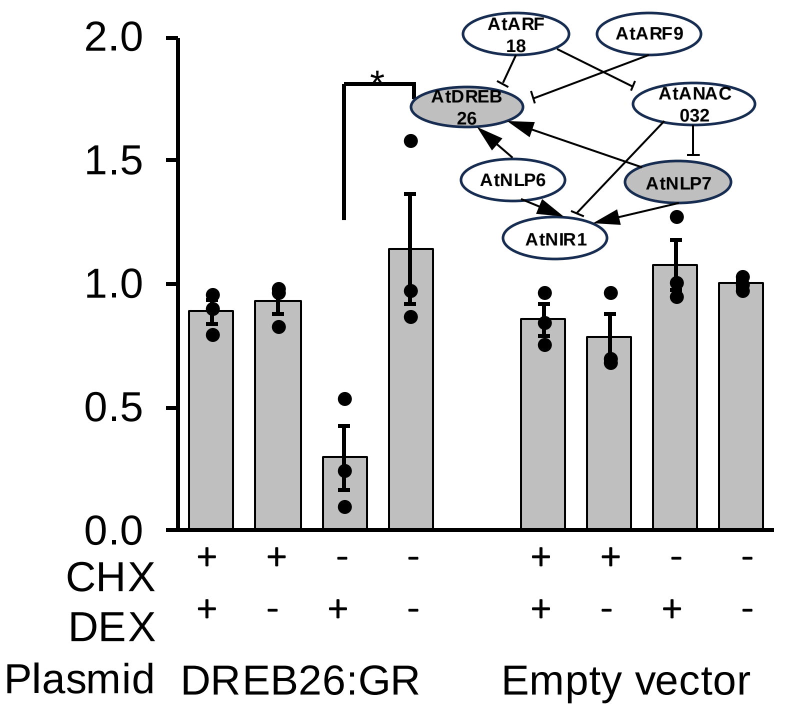

**Expression of *AtNLP7* is directly repressed by *35sP*-AtANAC032:GR**

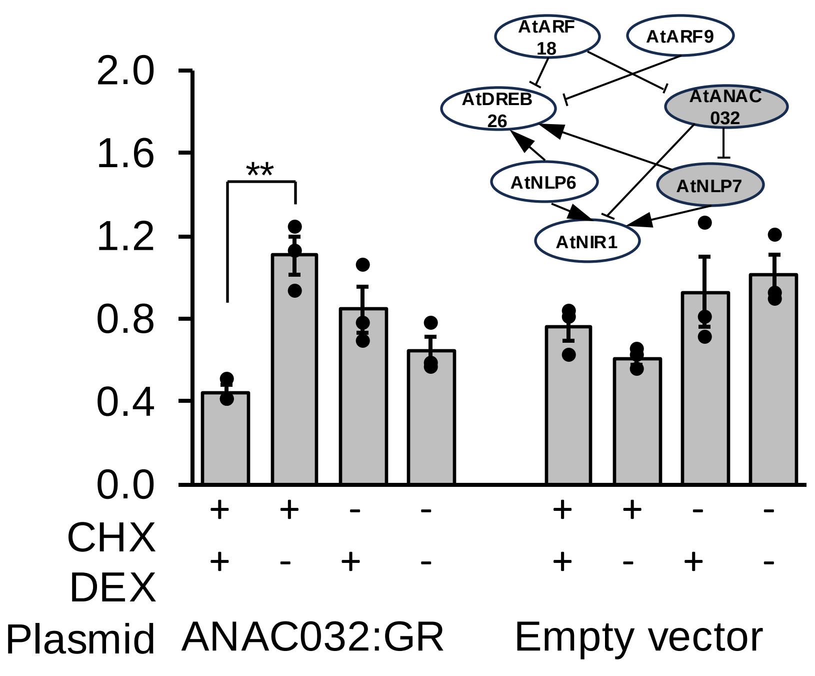

**Expression of *AtNLP7* is indirectly repressed by *35sP*-AtARF18:GR**

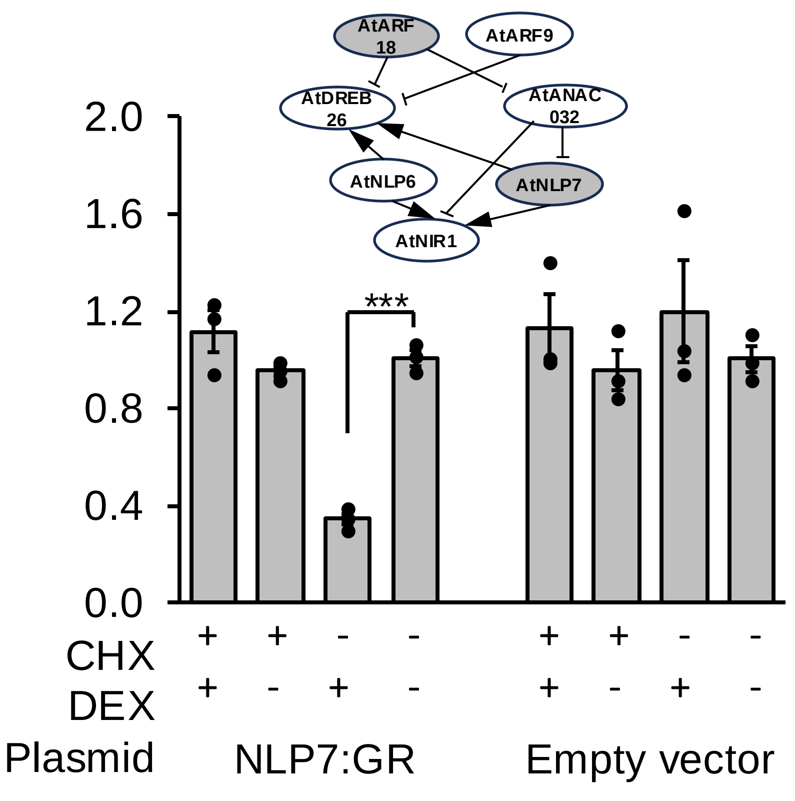

**Expression of *AtNLP7* is not affected by *35sP*-AtARF9:GR**

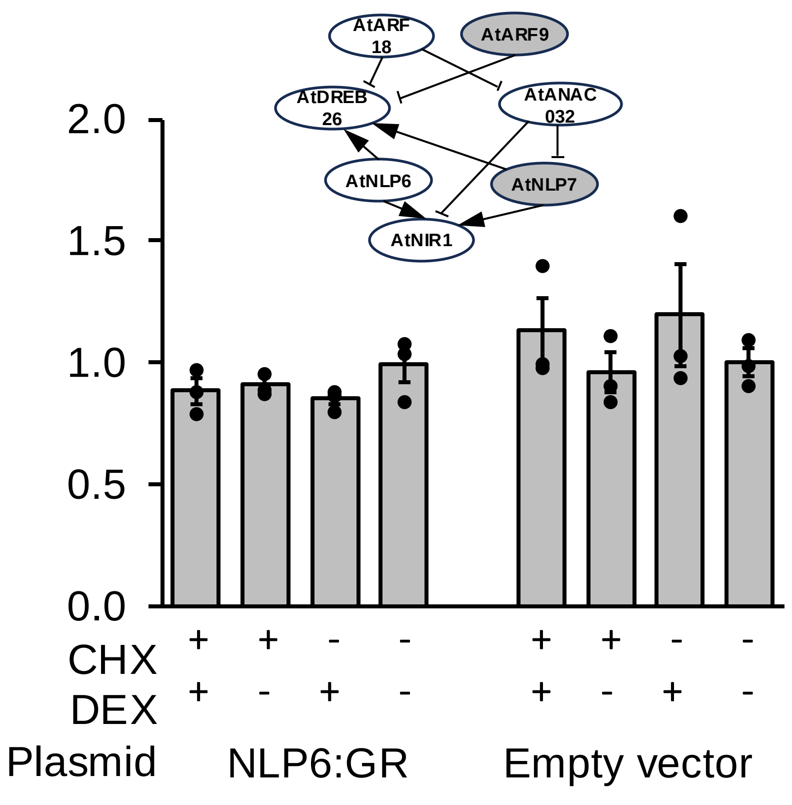

**Target Genes**

| Transcription Factor | *AtDREB26* | *AtNLP7* | *AtNLP6* | *AtNIR1* | *AtANAC032* |
|---|---|---|---|---|---|
| **AtDREB26:GR** | *nt | -0.06<br>**-2.21** | -0.12<br>**-1.60** | +0.09<br>**-2.69** | -0.08*<br>-0.14* |
| **AtNLP7:GR** | *nt | -0.17<br>-0.37 | nt | **+2.07**<br>-0.41 | nt |
| **AtNLP6:GR** | *nt | -0.61<br>-0.72 | nt | **+0.99**<br>+0.04 | nt |
| **AtARF18:GR** | *nt | -0.01<br>**-1.78** | -0.05<br>**-0.98** | -0.37<br>**-1.38** | **-3.91***<br>**-2.63*** |
| **AtARF9:GR** | *nt | -0.05<br>-0.21 | +0.04<br>+0.15 | -0.37<br>+0.12 | +0.07*<br>+0.32* |
| **AtANAC032:GR** | *nt | **-1.39**<br>-0.57 | +0.02<br>-0.35 | **-1.07**<br>+0.59 | nt |

**Summary of changes in gene expression of target genes in modified TARGET assays.** Figures indicate fold change in expression. Upper figures indicate the effects of direct interactions (cells treated with cycloheximide). Figures below indicate indirect effects (absence of cycloheximide). Red text indicates significant repression. Blue text indicates significant activation. nt, not tested; *, expression of the target gene responded to cycloheximide.

**Supplementary Data 7. Transactivation luciferase assays.** Graphs show luminescence from Arabidopsis promoter:nanoluciferase (LucN) constructs relative to CaMV35s:LucF and normalized CaMV35s:LucN/CaMV35s:LucF to with and without co-expression of transcription factor. Values represent the mean and standard error of three biological replicates (independent transfections). *P*-values were calculated using an unpaired two-tailed Student's t-test **P*<0.05, ** *P*<0.01, *** *P*<0.001).

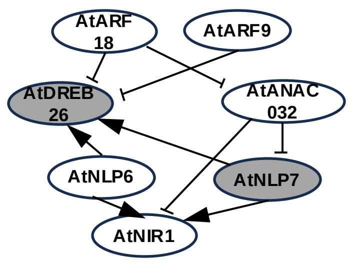

***DREB26$_P$*-LucN luminescence is increased by co-expression of *35sP*-AtNLP7**

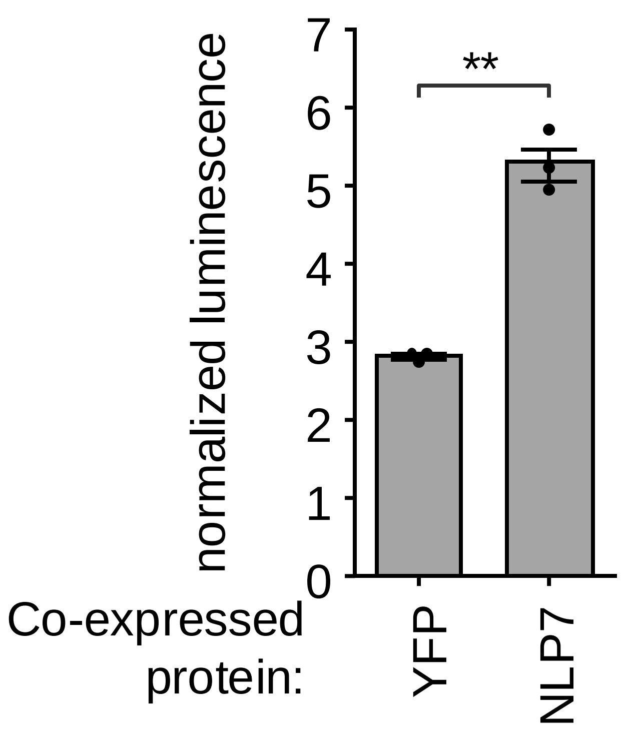

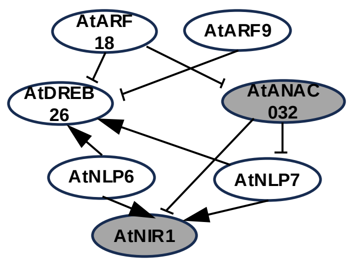

***AtNIR1$_P$*-LucN luminescence is increased by co-expression of *35sP*-AtANAC032**

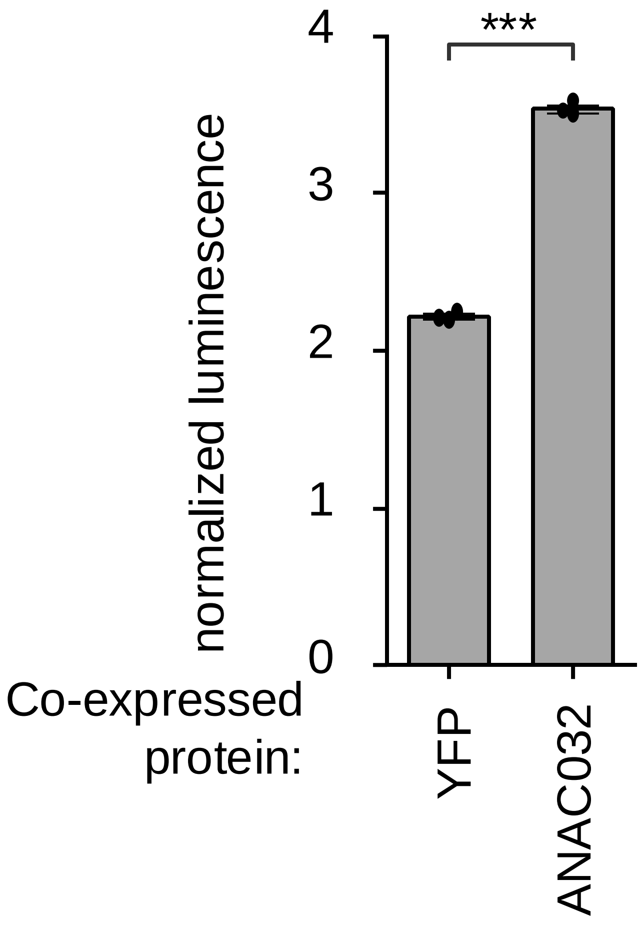

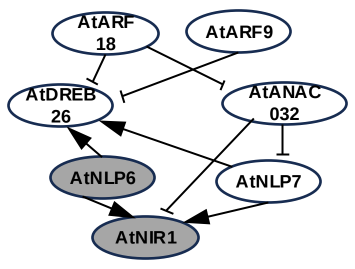

***AtNIR1$_P$*-LucN luminescence is increased by co-expression of *35sP*-AtNLP6**

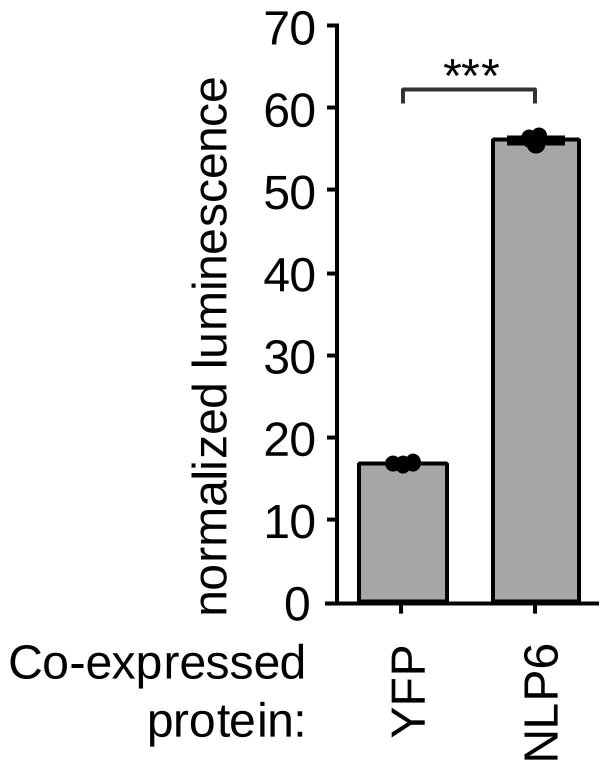

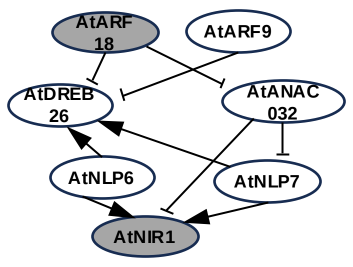

***AtNIR1$_P$*-LucN luminescence is reduced by co-expression of *35sP*-AtARF18**

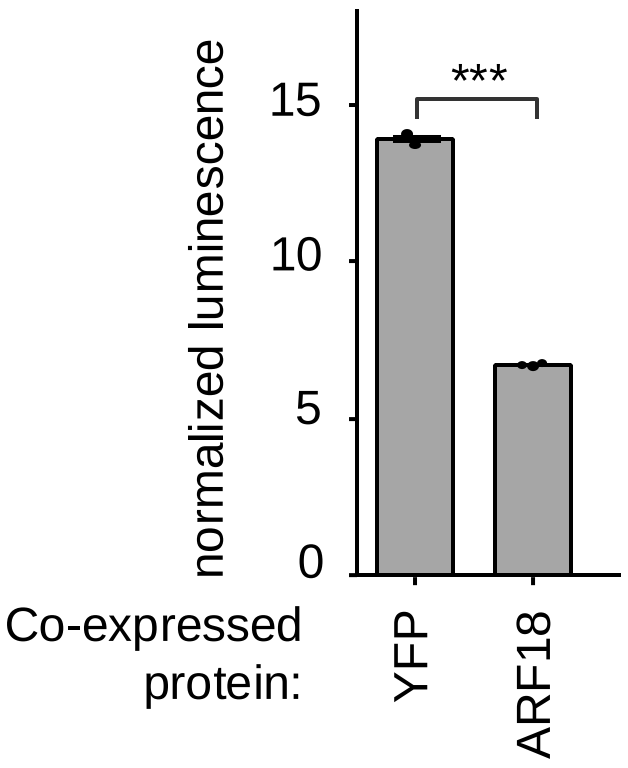

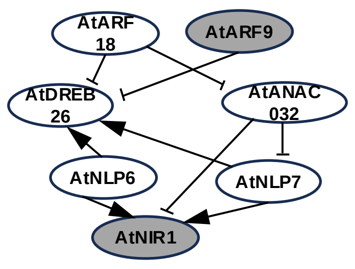

***AtNIR1$_P$*-LucN luminescence is reduced by co-expression of *35sP*-AtARF9**

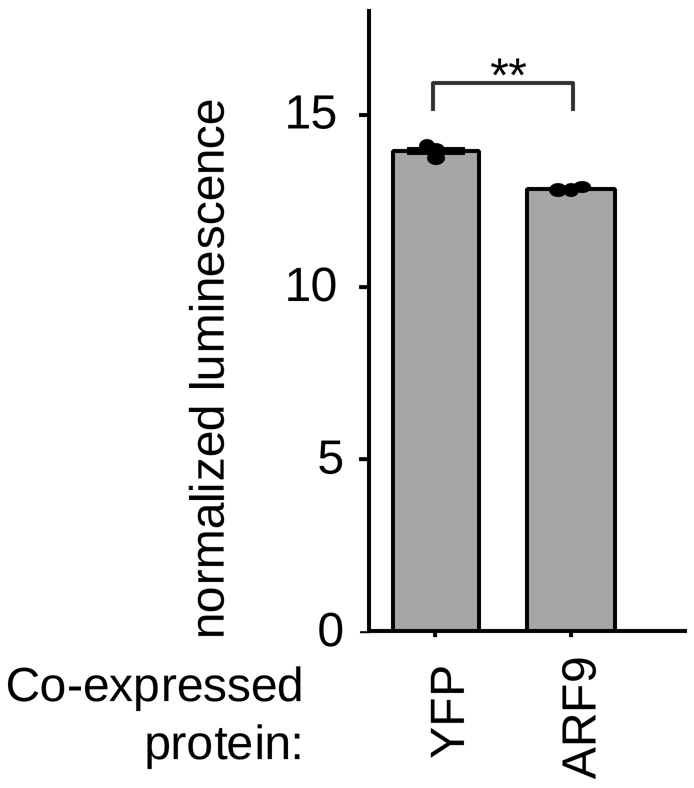

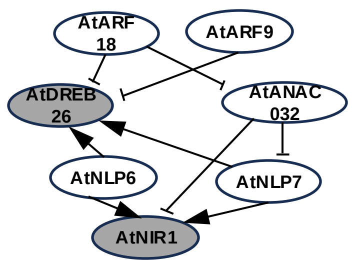

***AtNIR1$_P$*-LucN luminescence is reduced by co-expression of *35sP*-AtDREB26**

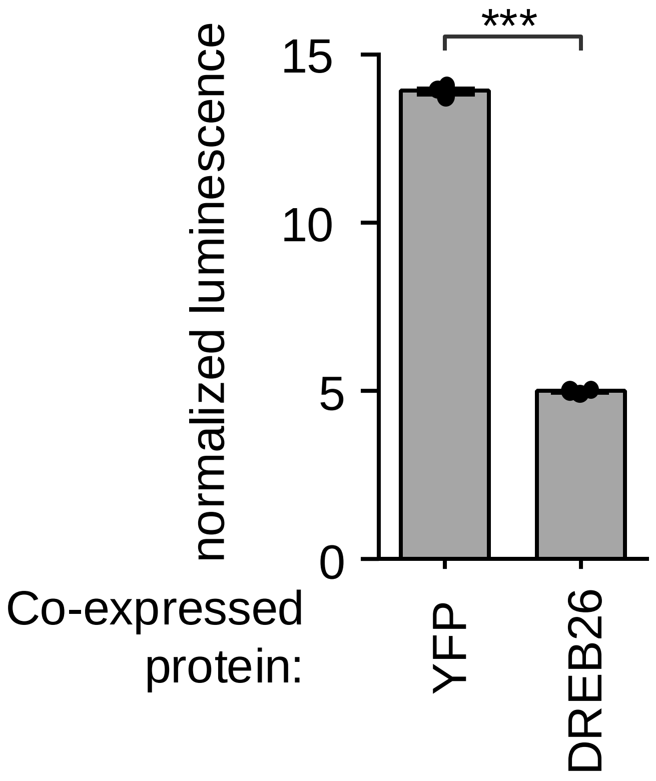

**Supplementary Data 8. Nitrate responsive genes in tomato roots. (a)** A heatmap shows scaled log2CPM of tomato nitrogen-responsive genes in the M82 root RNA-seq experiment. **(b-d)** GO enrichment analysis on 3 clusters of DEGs that are up-regulated at higher nitrate conditions in M82 RNA-seq experiment.

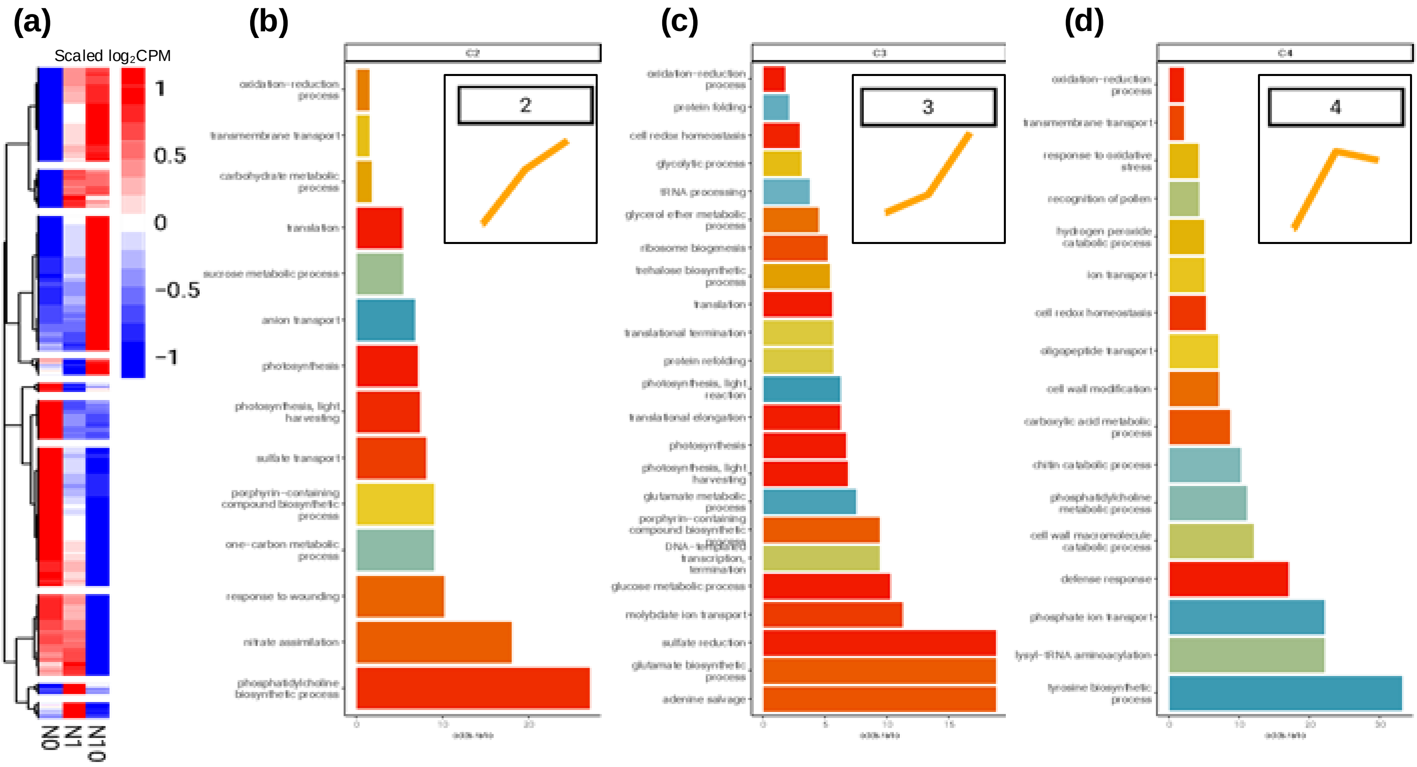

**Supplementary Data 9. Phylogenetic analysis of transcription factors. (a – g)** Maximum likelihood analysis of the amino acid sequences of transcription factors. Scale bars indicate the number of substitutions per site. Numbers at nodes indicate bootstraps. Sequences from *S. lycopersicum* are highlighted in red, while those from Arabidopsis are highlighted in blue.

### (a) ARF18

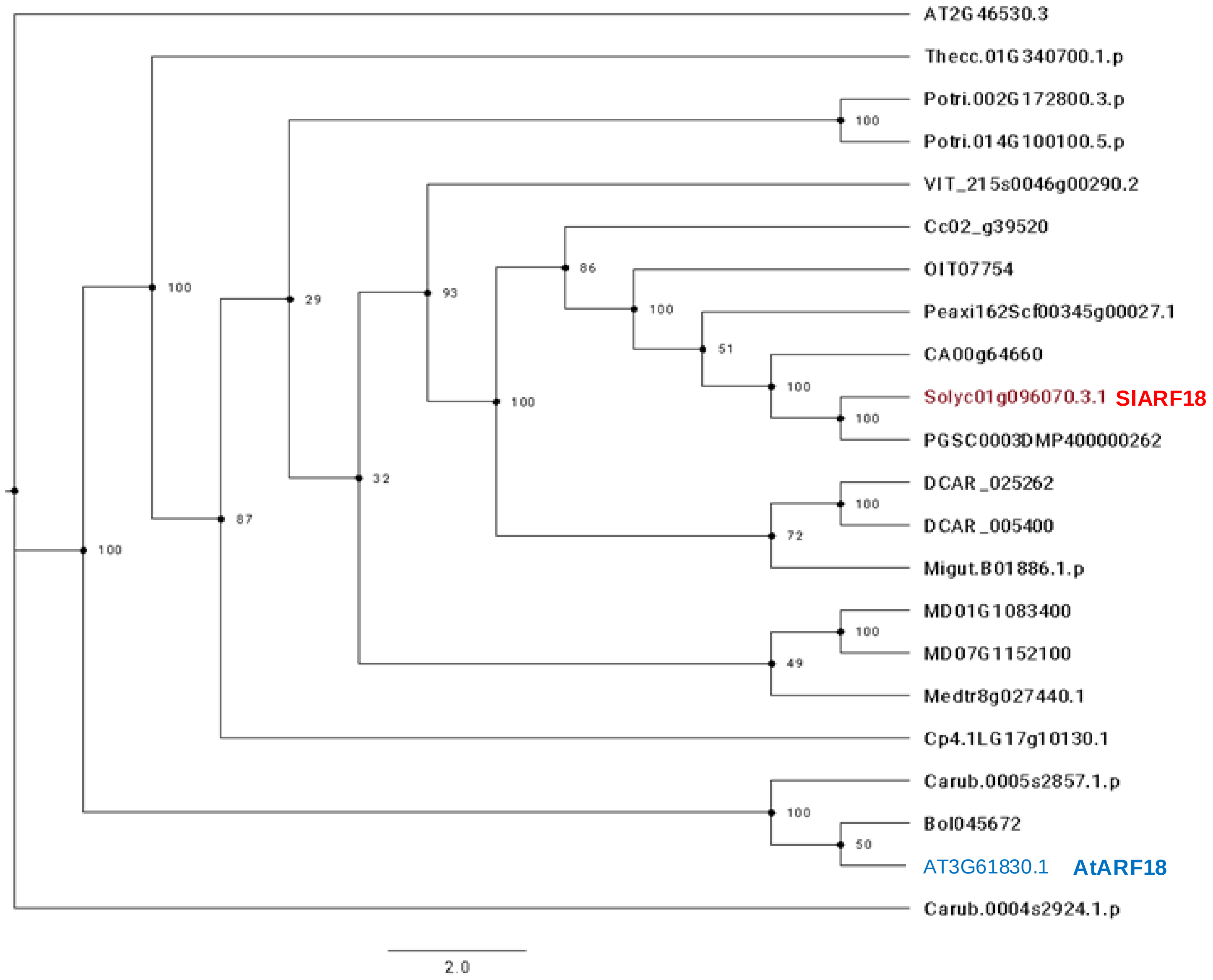

**Supplementary Data 9. Phylogenetic analysis of transcription factors in this study. (a – g)** Maximum likelihood analysis of the amino acid sequences of transcription factors. Scale bars indicate the number of substitutions per site. Numbers at nodes indicate bootstraps. Sequences from *S. lycopersicum* are highlighted in red, while those from Arabidopsis are highlighted in blue. (b) The tpm expression of two tomato *ARF9* orthologs showing *SlARF9A* gene is not expressed in tomato roots, while *SlARF9B* gene shows high expression in tomato roots. The source data is from Kajala et al (23) and X axis represents different cell types. COR, cortex; EN, endodermis; EP, epidermis; EXO, exodermis; MCO, meristematic cortex; MZ, meristematic zone; PH, phloem; V, vascular initials; WOX, *SlWOX5* expressed region; XY, xylem.

### (b) ARF9

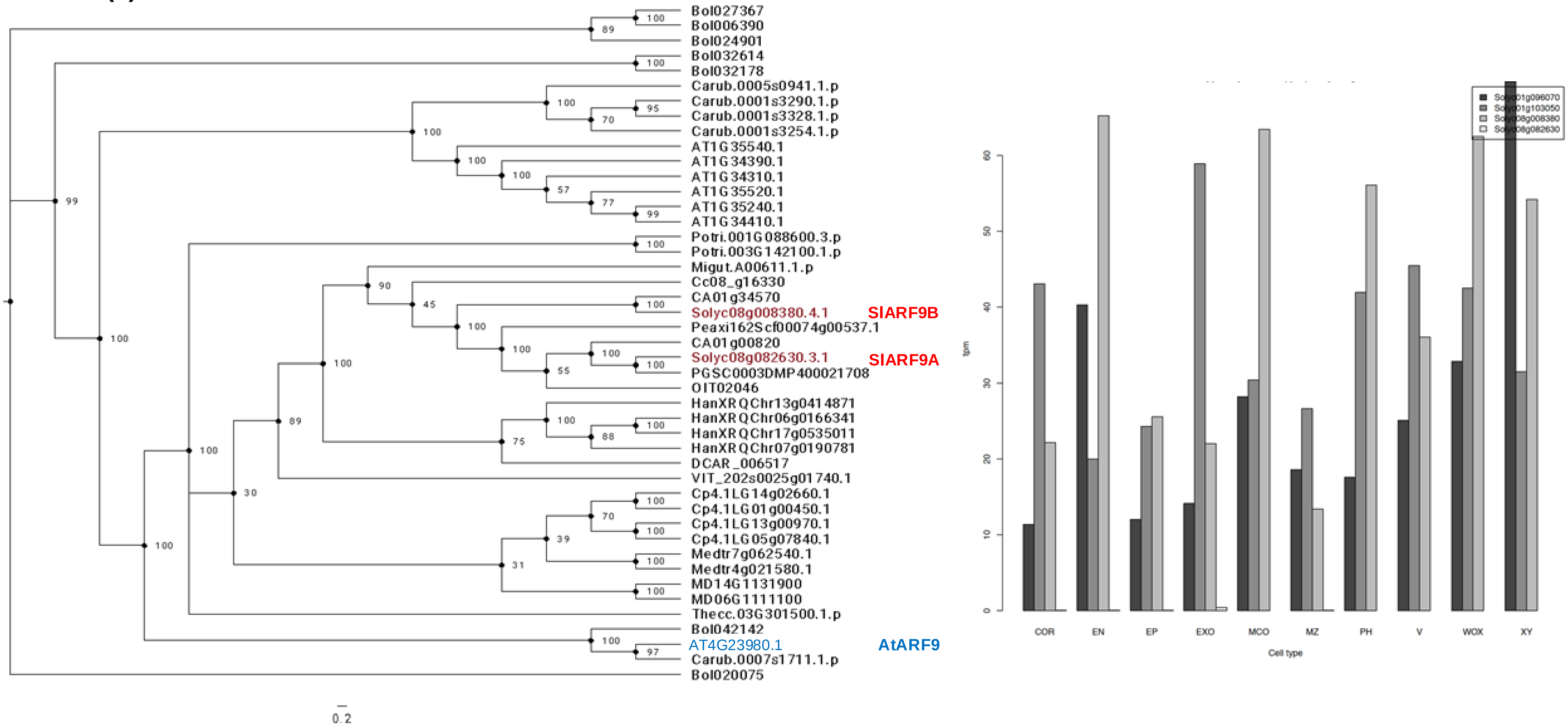

**Supplementary Data 9. Phylogenetic analysis of transcription factors in this study. (a – g)** Maximum likelihood analysis of the amino acid sequences of transcription factors. Scale bars indicate the number of substitutions per site. Numbers at nodes indicate bootstraps. Sequences from *S. lycopersicum* are highlighted in red, while those from Arabidopsis are highlighted in blue.

### (c) DREB26

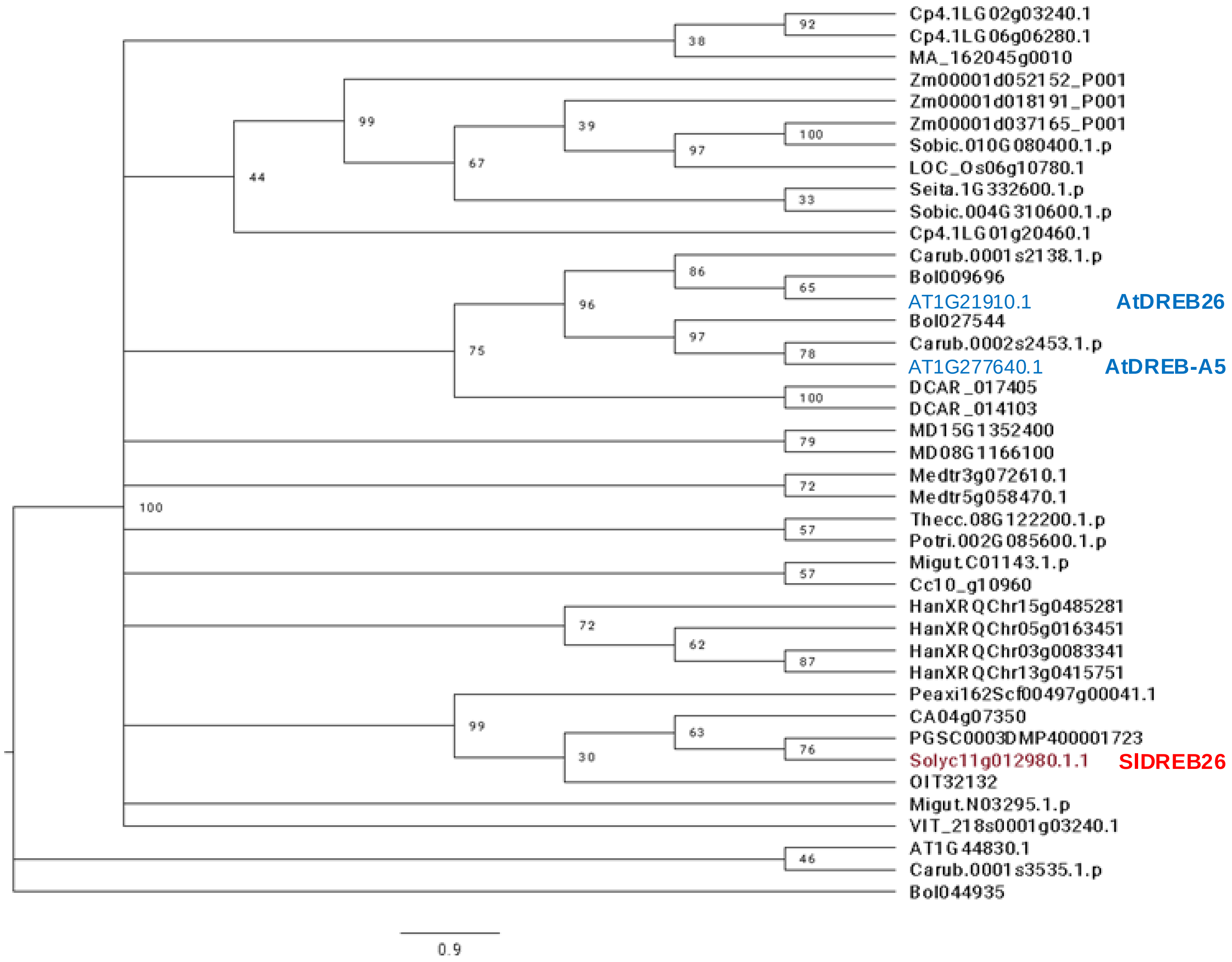

**Supplementary Data 9. Phylogenetic analysis of transcription factors in this study. (a – g)** Maximum likelihood analysis of the amino acid sequences of transcription factors. Scale bars indicate the number of substitutions per site. Numbers at nodes indicate bootstraps. Sequences from *S. lycopersicum* are highlighted in red, while those from Arabidopsis are highlighted in blue.

### (d) NLP6/7

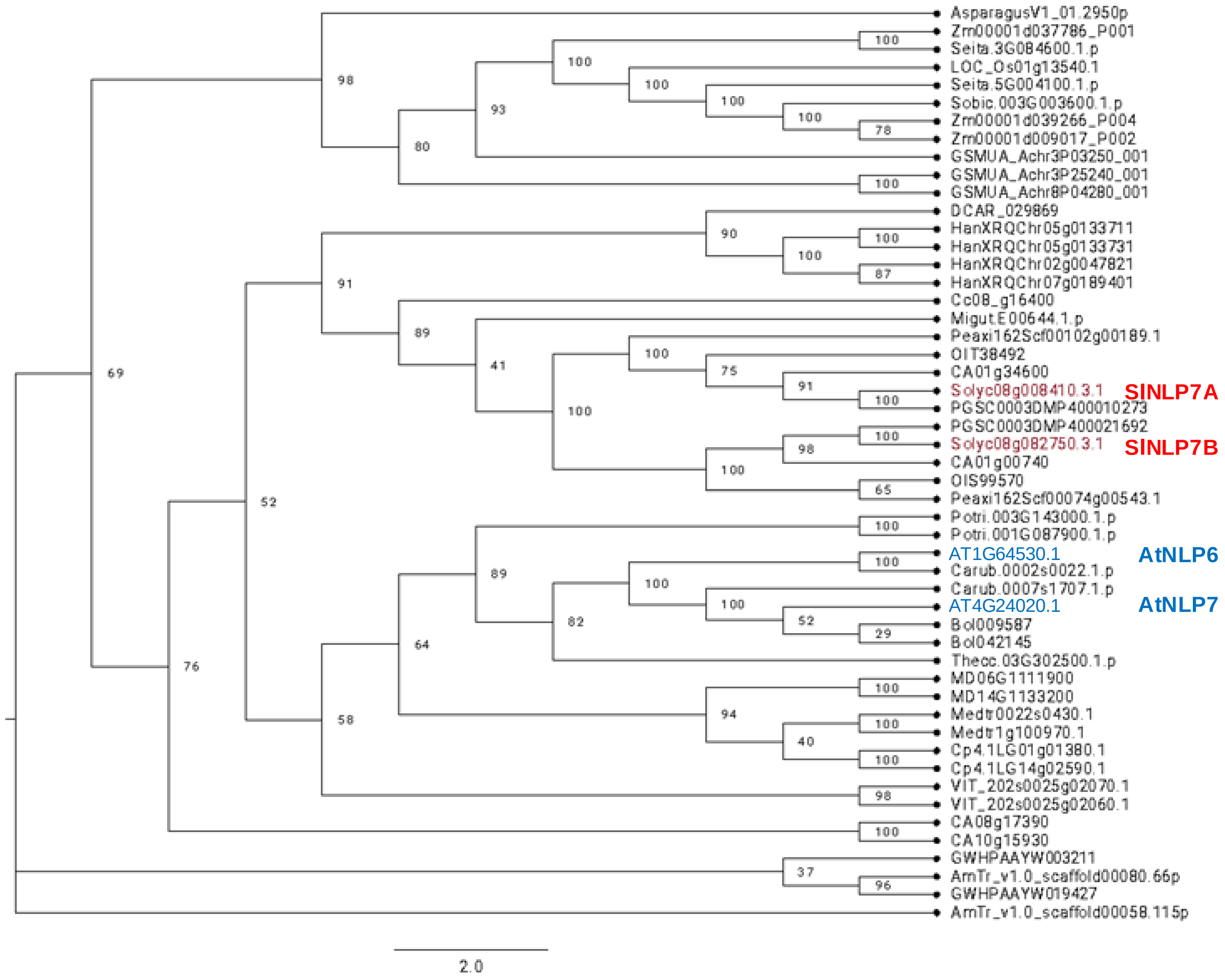

**Supplementary Data 9. Phylogenetic analysis of transcription factors in this study. (a – g)** Maximum likelihood analysis of the amino acid sequences of transcription factors. Scale bars indicate the number of substitutions per site. Numbers at nodes indicate bootstraps. Sequences from *S. lycopersicum* are highlighted in red, while those from Arabidopsis are highlighted in blue.

**(c) ANAC032**

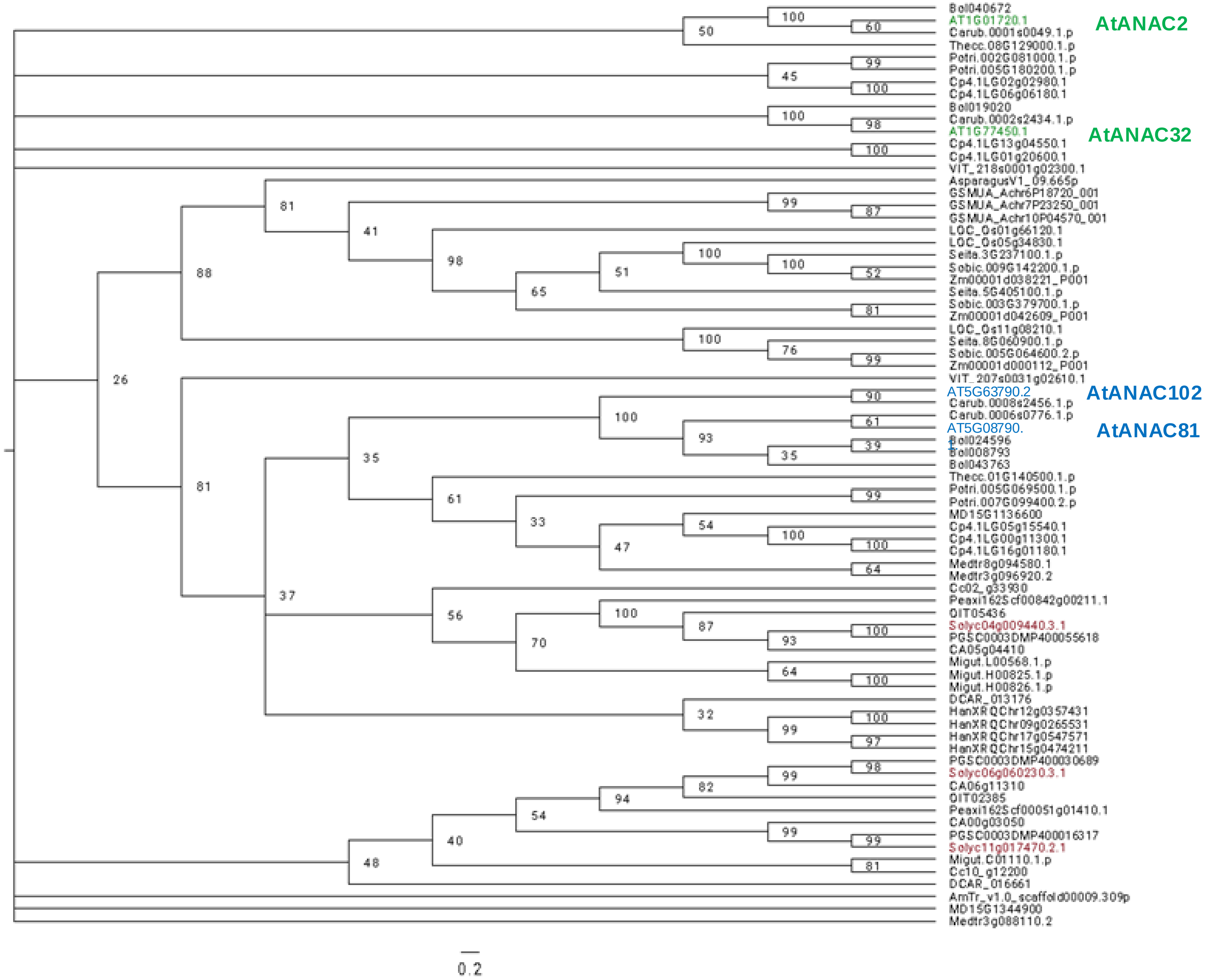

**Supplementary Data 9. Phylogenetic analysis of transcription factors in this study. (a – g)** Maximum likelihood analysis of the amino acid sequences of transcription factors. Scale bars indicate the number of substitutions per site. Numbers at nodes indicate bootstraps. Sequences from *S. lycopersicum* are highlighted in red, while those from Arabidopsis are highlighted in blue.

### (f) NIR1

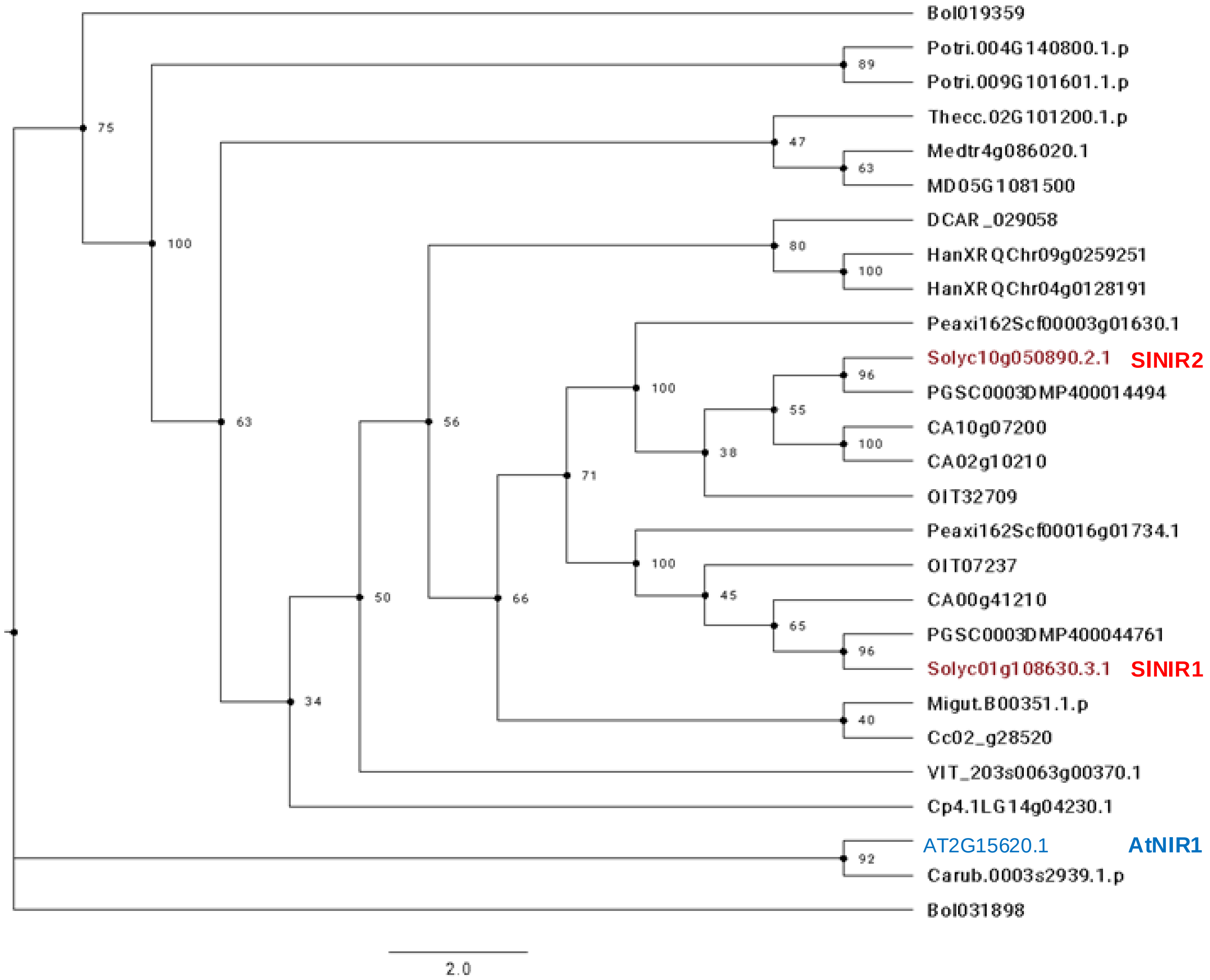

**Supplementary Data 10.** Domain analysis of tomato orthologues of *AtNLP7.* (A-B) RNAseq reads from Kajala, *et al.*, 2021 (*Cell* **184**, 3333-3348.e19) were mapped to predicted NLP7 orthologs in the tomato genome. *Solyc08g008410* appears mis-annotated and contains duplicate genes. We name these *SlNLP7A* and *SlNLP7C*, the latter of which has relatively low expression (B) RNAseq reads mapped to the annotated SlNLP7B gene. (C) Amino acid sequence alignment of SlNLP7A, SlNLP7B, and SlNLP7C showing similarity. (D-E) Domain analysis of SlNLP7A, SlNLP7B, and AtNLP7. All contain (D) RWP-RK, (D) PB1, and (E) nitrate binding domains as previously reported (Liu *et al.,* 2022. *Science* 377, 1419–1425).

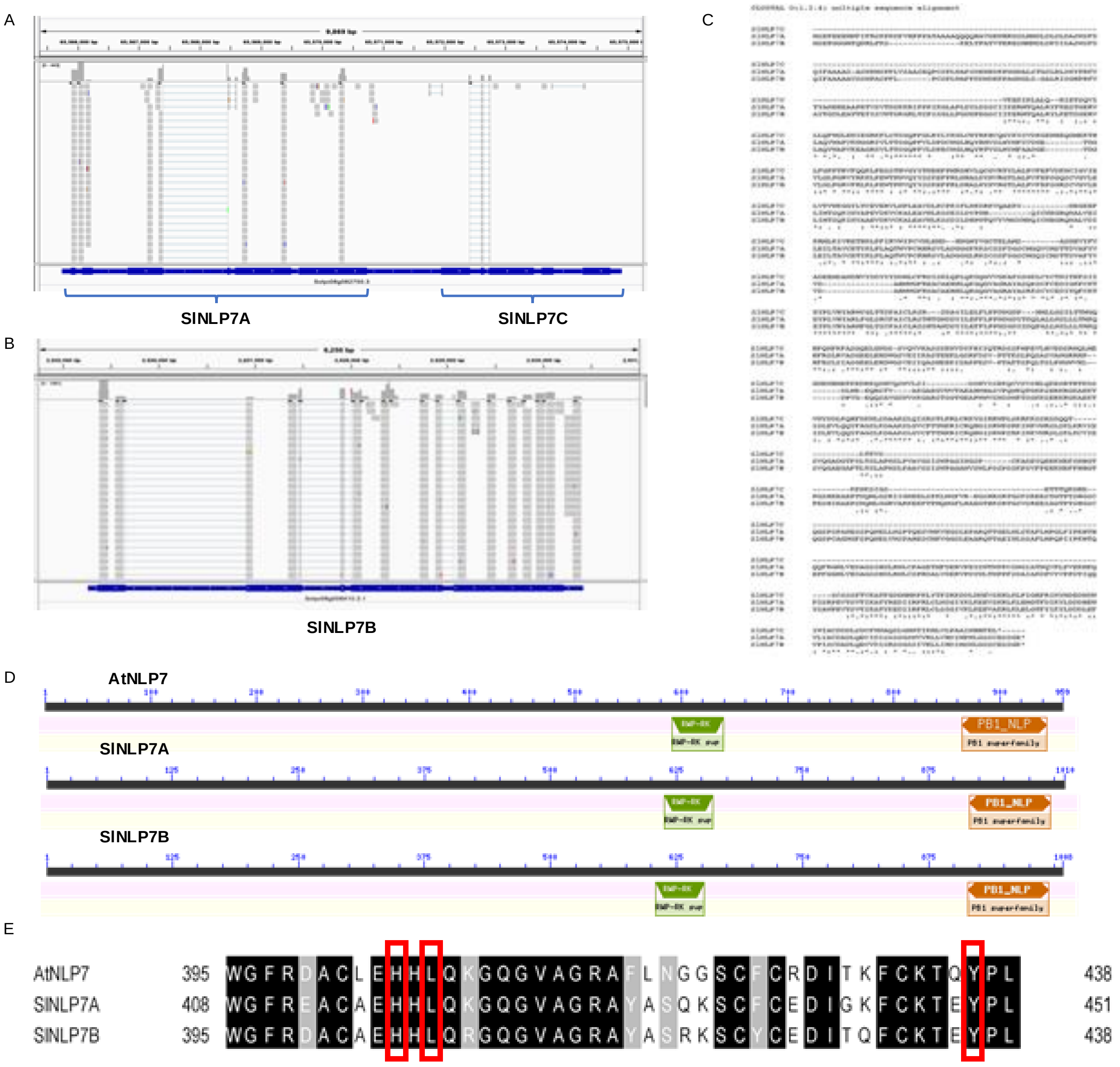

**Supplementary Data 11**
Table of Arabidopsis and Tomato mutant lines.
Bold text with subscript ‘c’ indicates mutation was created by CRISPR/Cas9 (details in Supplementary data 12). All other lines are T-DNA insertion lines.

| **Arabidopsis (At)** | **Tomato (Sl)** |
|---|---|
| *nlp6* | ***nlp7a*** |
| *nlp7-1* | ***nlp7b*** |
| *nlp6/nlp7-1* | ***nlp7a/nlp7b*** |
| *anac032* | |
| ***anac032***$_c$ | |
| *arf9b* | ***arf9b*** |
| *arf18-2* | |
| *arf18-3* | |
| ***arf18***$_c$ | ***arf18*** |
| *arf18-2/nlp7-1* | ***arf9b/arf18*** |
| ***dreb26***$_c$ | ***dreb26*** |
| ***dreb26***$_c$***/nlp7***$_c$ | ***arf9b/arf18/dreb26*** |
| *anac032/nlp7-1* | ***dreb26/nlp7a/nlp7b*** |
| *nlp7-1/arf18-2* | ***arf9b/arf18/dreb26/nlp7a/nlp7b*** |
| *arf18-2/anac032* | |

**Supplementary Data 12.** Genotypes of Arabidopsis and tomato CRISPR lines.

### Genotypes of Arabidopsis CRISPR/Cas9 lines

| Plant ID | Target TF(s) | sgRNA1 | sgRNA2 |
|---|---|---|---|
| dreb26-cc-1<br>(OZ0033T1-10T2-05T3-14) | AtDREB26 | GAAGATGGTGATGATGACAA**AGG**<br><br>**wild type** | TTAAGAAGTACAAAGGA-GTG**AGG**<br>TTAAGAAGTACAAAGGA**A**GTG**AGG**<br>**homozygous** |
| Anac032-cc-1<br>(OZ0033T1-19T2-29) | AtANAC032 | ATTTACGACAGAGATACATG**AGG**<br><br>**wild type** | AATATCCAGTACCAGCTGCA**CGG**<br>AAT--------------GCA**CGG**<br>**homozygous** |
| Arf18-cc-1<br>(OZ0033T1-19T2-02T3-21) | AtARF18 | GTTGAAGGTGATGATGATTT**CGG**<br><br>**wild type** | AGAGTTTTCTACTTCCC-TCA**GGG**<br>AGAGTTTTCTACTTCCC**T**TCA**GGG**<br>**homozygous** |
| dreb26-nlp7-cc-1<br>(OZ0103T1-47T2-35T3-06) | AtDREB26<br><br><br><br>AtNLP7 | CGTGTAAGAACAAGAACAAG**AAGAGT**<br>CGTGTAAGAACAAGAAC---------<br>**homozygous**<br><br>GTAGAGCTCGGACTCCCGGG**TCGAGT**<br>GTAGAGCTCGGACTCCC---------<br>**homozygous** | GATTAATTGAAACTCCAAAG**CGGAAT**<br>GATTAATTGAAACTCCA---------<br>**homozygous**<br><br>GTCTGAGCGAGAGGCAAGTT**ATGGGT**<br>GTCTGAGCGAGAGGCAA---------<br>**homozygous** |

**Supplementary Data 12.** Genotypes of Arabidopsis and tomato CRISPR lines.

*Sequencing chromatograms of homozygous CRISPR/Cas9 knockout lines*. Exons are schematically displayed as blue boxes and introns and untranslated regions as grey boxes. The sgRNA target sites are represented with pink arrowheads. The DNA sequence of sgRNAs are highlighted in grey and PAMs are shown in bold case. CRISPR/Cas9-induced mutations are shown below the chromatogram in red bold case letters or dashes.

DREB26-cc-1[HMZ]
OZ0033T1-10T2-05T3-14

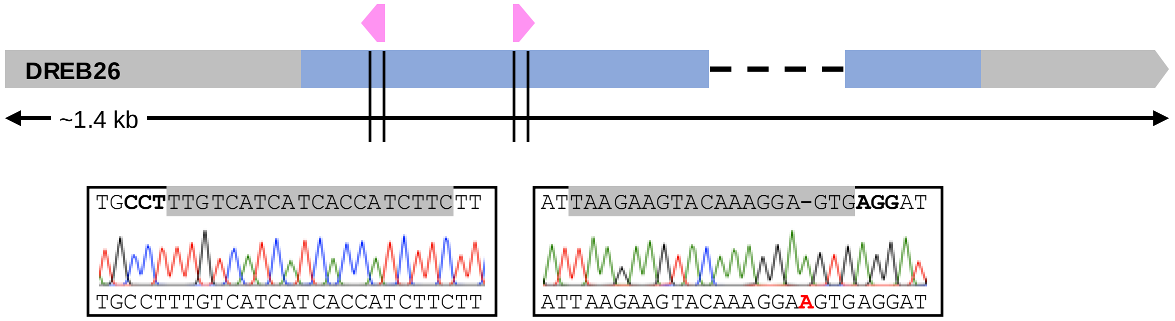

ANAC032-cc-1[HMZ]
OZ0033T1-19T2-29

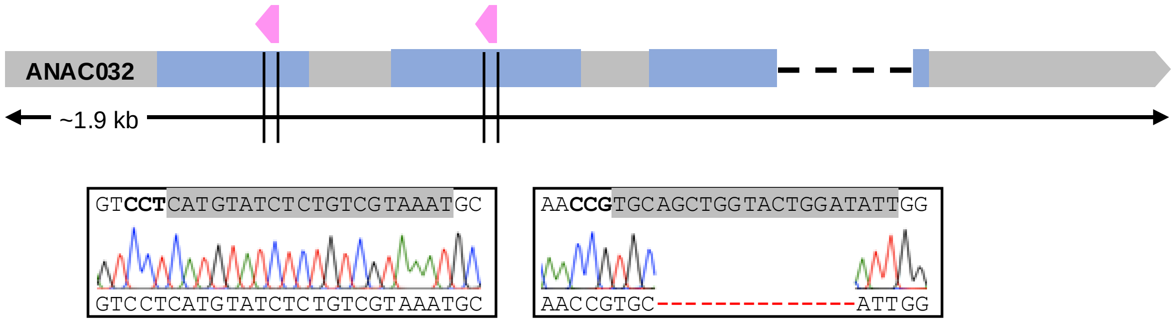

ARF18-cc-1
OZ0033T1-19T2-02T3-21

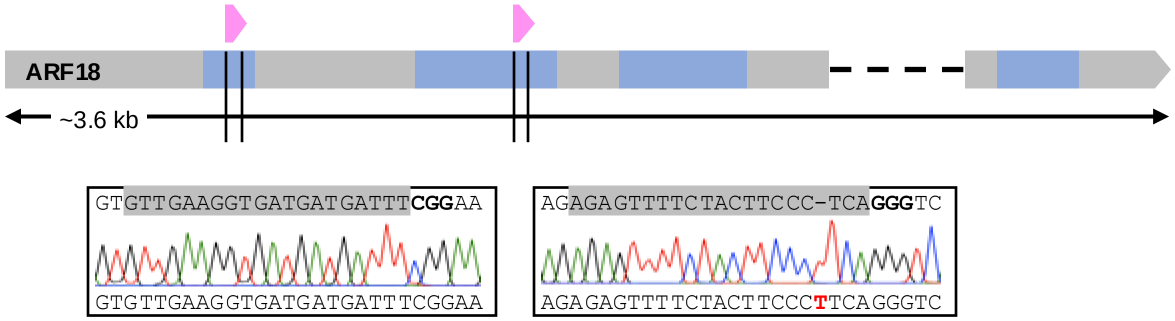

**Supplementary Data 12.** Genotypes of Arabidopsis and tomato CRISPR lines.

*Sequencing chromatograms of homozygous CRISPR/Cas9 knockout lines*. Exons are schematically displayed as blue boxes and introns and untranslated regions as grey boxes. The sgRNA target sites are represented with pink arrowheads. The DNA sequence of sgRNAs are highlighted in grey and PAMs are shown in bold case. CRISPR/Cas9-induced mutations are shown below the chromatogram in red bold case letters or dashes.

DREB26[HMZ] NLP7[HMZ]
OZ0103T1-47T2-35T3-06

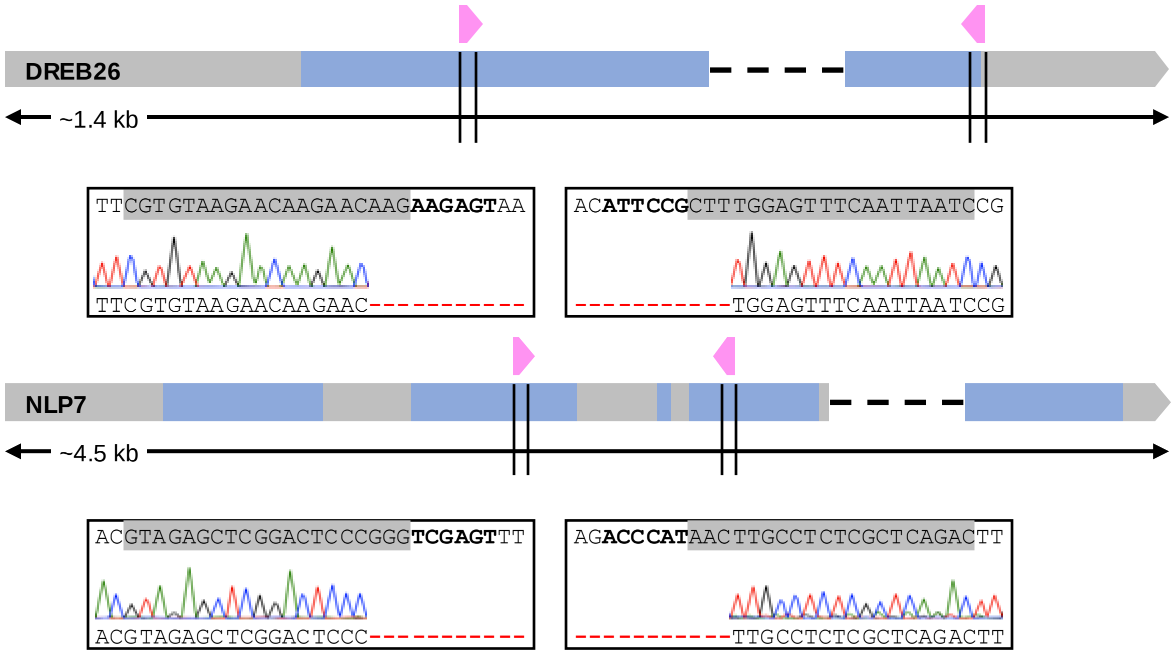

### Genotypes of Tomato CRISPR/Cas9 hairy root lines

| Plant ID | Target TF | sgRNA1 | sgRNA2 |
|---|---|---|---|
| ***slarf9b #8*** | SlARF9B | GAGTTATGGAGATTGTGTTGC**AGG** **+1bp**<br>**homozygous** | AGCGTAATTTAGGAGGGTTTG**TGG** **+1bp**<br>**homozygous** |
| ***slarf18 #2*** | SlARF18 | GATCTCTGGAAGGCA----TGC**AGG** **-2bp**<br>**homozygous** | GATCACATTGCACCCT--AGG**CGG** **-1bp**<br>**homozygous** |
| ***slarf9b/18 #4*** | SlARF9B<br><br><br><br>SlARF18 | GAGTTATGGAGATTGTG----------- **-13bp**<br>GAGTTATGGAGAT----TGTGC**AGG** **-2bp**<br>GAGTTATGGAGATTGTTGTGC**AGG** **+1bp**<br>**mosaic**<br><br>GATCTCTGGAA------------TGC**AGG** **-6bp**<br>GATCTCTGGAAGGC-----TGC**AGG** **-3bp**<br>**biallelic** | AGCGTAATTTAGGAGGGTTTG**TGG** **+1bp**<br>AGCGTAATTTAGGAGGGTTTTG**TGG** **+2bp**<br>**biallelic**<br><br>GATCACATTGCACCCTGTAGG**CGG** **+1bp**<br>GATCACATTGCACCCT----AGG**CGG** **-1bp**<br>**biallelic** |
| ***sldreb26 #10*** | SlDREB26 | GAAGAACAAG---------------------------- **-71bp**<br>**homozygous** | **CCT**----AAGGCCCCTCTGCTTCCT **-2bp**<br>**homozygous** |
| ***slnlp7a #1*** | SlNLP7A | GGACTTTGTACATCCA--CGC**CGG** **-1bp**<br>GGACTTTGTACATCC----------------- **-174bp**<br>**biallelic** | TATGTCGCCGTTTC----------T**CGG** **-5bp**<br>---------------------------------GTAT**CGG** **-174bp**<br>**biallelic** |
| ***slnlp7b #13*** | SlNLP7B | AAAGGAGTTGACTCCGGCGA**CGG** **WT**<br>**wild type** | **CCT**CT-----------------TTTCCCGATGA **-7bp**<br>**CCT**CTTTTGGGCATTTCCCGATGA **+1bp**<br>**biallelic** |
| ***slnlp7a/b #21*** | SlNLP7A<br><br><br>SlNLP7B | GGACTTTGTACATCCACCGC**CGG** **WT**<br>**wild type**<br><br>AAAGGAGTTGACTCCGGCGA**CGG** **WT**<br>AAAGGAGTTGACTCCGGTCGA**CGG** **+1bp**<br>**heterozygous** | TATGTCGCCGTTTCTCGATAT**CGG** **+1bp**<br>**homozygous**<br><br>**CCT**C----------------TTTCCCGATGA **-8bp**<br>**homozygous** |
| ***slarf9b/18/dreb26 #9*** | SlARF9B<br><br><br>SlARF18<br><br><br>SlDREB26 | GAGTTATGGAGAT----TGTGC**AGG** **-2bp**<br>**homozygous**<br><br>GATCTCTGGAAGGCATG--GC**AGG** **-1bp**<br>**homozygous**<br><br>CAAAGGAGT----------------------------- **-130bp**<br>GAATGAGAAGCTGGGGA--CA**TGG** **-1bp**<br>**biallelic** | AGCGTAATTTAGGA------------------ **-11bp**<br>**homozygous**<br><br>----------------------------------AGG**CGG** **-20bp**<br>**homozygous**<br><br>------------AGGCCCCTCTGCTTCCT **-130bp**<br>**CCT**A---------------CCTCTGCTTCCT **-7bp**<br>**biallelic** |
| ***sldreb26/nlp7a/b #6*** | SlDREB26<br><br><br><br>SlNLP7A<br><br><br><br>SlNLP7B | GAATGAGAAGC-------------TCA**TGG** **-6bp**<br>GAATGAGAAGCT-----------TCA**TGG** **-5bp**<br>**biallelic**<br><br>GGACTTTGTACATCCACCGC**CGG** **WT**<br>GGACTTTGTA---------------CGC**CGG** **-7bp**<br>**heterozygous**<br><br>AAAGGAGTTGACTCCGGCGA**CGG** **WT**<br>AAAGGAGTTGACT--CGGCGA**CGG** **-1bp**<br>**heterozygous** | **CCT**----AAGG--CCCTCTGCTTCCT **-3bp**<br>**CCT**AAAAGGGCCCCTCTGCTTCCT **+1bp**<br>**biallelic**<br><br>TATGTCGCCGTTTCTCGTTAT**CGG** **+1bp**<br>**homozygous**<br><br>**CCT**C------------------TTTCCCGATGA **-8bp**<br>**CCT**CT----------------TTTCCCGATGA **-7bp**<br>**biallelic** |
| ***slarf9b/18/dreb26/nlp7a/b #2*** | SlARF9B<br><br>SlARF18<br><br>SlDREB26<br><br>SlNLP7A<br><br>SlNLP7B | GAGTTATGGAGATTGTG------**AGG** **-3bp**<br>GAGTTATGGAGATTGTGTGCT**AGG** **+1bp**<br>**biallelic**<br>GATCTCTGGAAGGCAT--TGC**AGG** **-1bp**<br>**homozygous**<br>GAAGAACAAGTAC--------------------- **-101bp**<br>**homozygous**<br>GGACTTTGTACATCCA--CGC**CGG** **-1bp**<br>**homozygous**<br>AAAGGAGTTGACTCCGGCGA**CGG** **WT**<br>**wild type** | AGCGTAATTTAGGAGGG----G**TGG** **-2bp**<br>AGCGTAATTTAGGAGGGTTTG**TGG** **+1bp**<br>**biallelic**<br>GATCACATTGCACCCT--AGG**CGG** **-1bp**<br>**homozygous**<br>**CCT**AAAAGGCCCCTCTGCTTCCT **WT**<br>**wild type**<br>----------------------TGCTTCTGAACAG **-111bp**<br>**homozygous**<br>**CCT**CTTTTGGGCATTTCCCGATGA **+1bp**<br>**homozygous** |

### Genotypes of Tomato CRISPR/Cas9 stable lines

| Plant ID | Target TF | sgRNA1 | sgRNA2 |
|---|---|---|---|
| ***slarf9b/18 #4-17*** | SlARF9B | GAGTTATGGAGATTGTGT----------- **-140bp**<br>**homozygous** | ----------------------------------------TG**TGG** **-140bp**<br>**homozygous** |
| | SlARF18 | GATCTCTGGAAGGCATGCAGGGT<br>**399bp inversion**<br>**homozygous** | CAACTAAAGGCCCTGCAAGG**CGG**<br>**399bp inversion**<br>**homozygous** |
| ***slnlp7b #8-3*** | SlNLP7B | AAAGGAGTTGACTCCGGCGA**CGG WT**<br>**wild type** | **CCT**CTT--GGGCATTTCCCGATGA **-1bp**<br>**homozygous** |

### gRNAs used to create Tomato CRISPR/Cas9 lines

| Gene | gRNA1 | gRNA2 |
|---|---|---|
| **SlDREB26** | GAATGAGAAGCTGGGGATCA | GAGGAAGCAGAGGGGCCTTTT |
| **SlNLP7B** | AAAGGAGTTGACTCCGGCGA | TCATCGGGAAATGCCCAAAG |
| **SlNLP7A** | TATGTCGCCGTTTCTCGTAT | GGACTTTGTACATCCACCGC |
| **SlARF18** | GATCTCTGGAAGGCATGTGC | GATCACATTGCACCCTGAGG |
| **SlARF9B** | GAGTTATGGAGATTGTGTGC | GAGCGTAATTTAGGAGGGTTG |

**Supplementary Data 12.** Genotypes of Arabidopsis and tomato CRISPR lines used in this study.

*Data supporting tomato CRISPR/Cas9 knockout lines.*

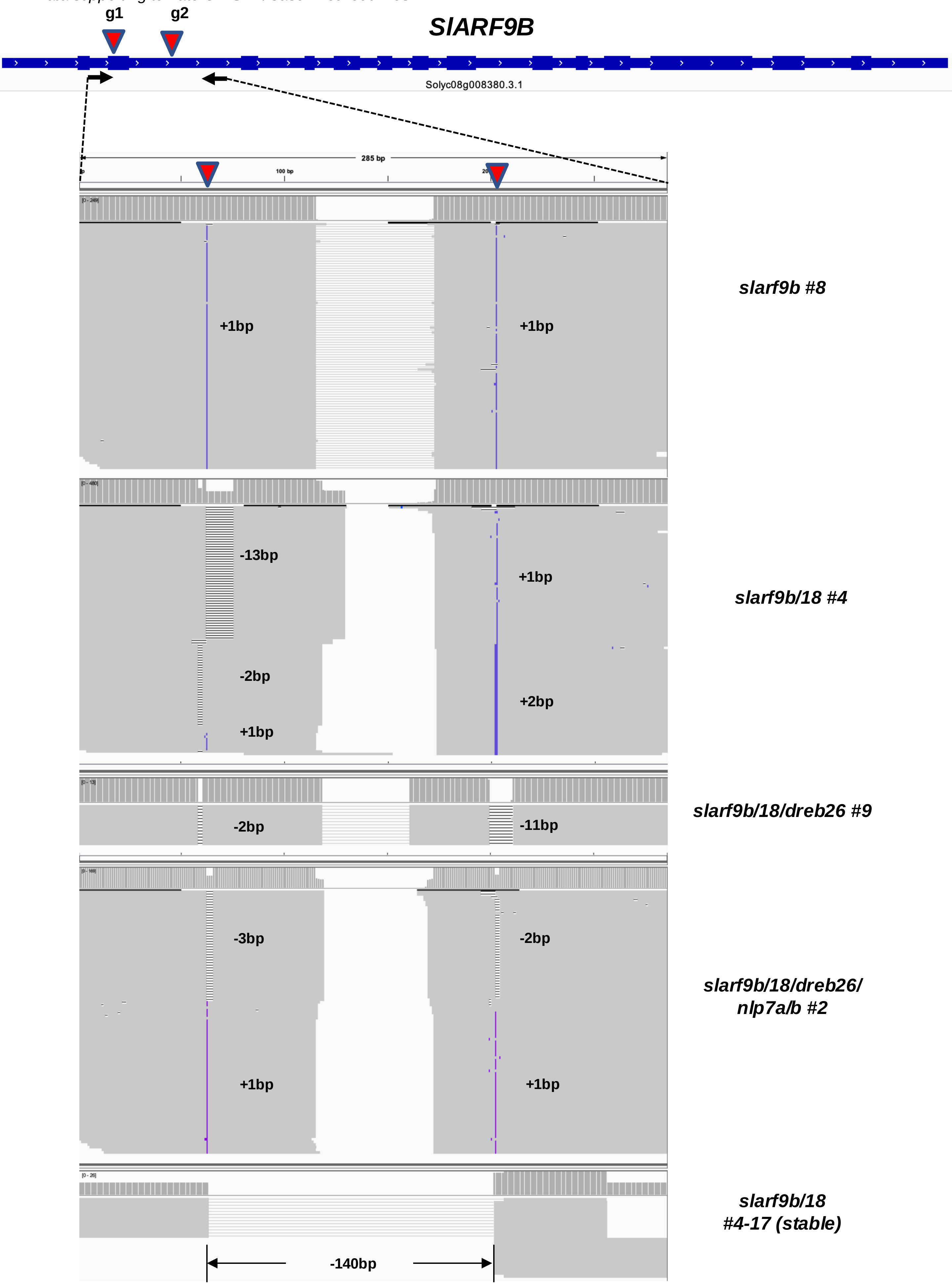

**Supplementary Data 12.** Genotypes of Arabidopsis and tomato CRISPR lines used in this study. *Data supporting tomato CRISPR/Cas9 knockout lines (Continued).*

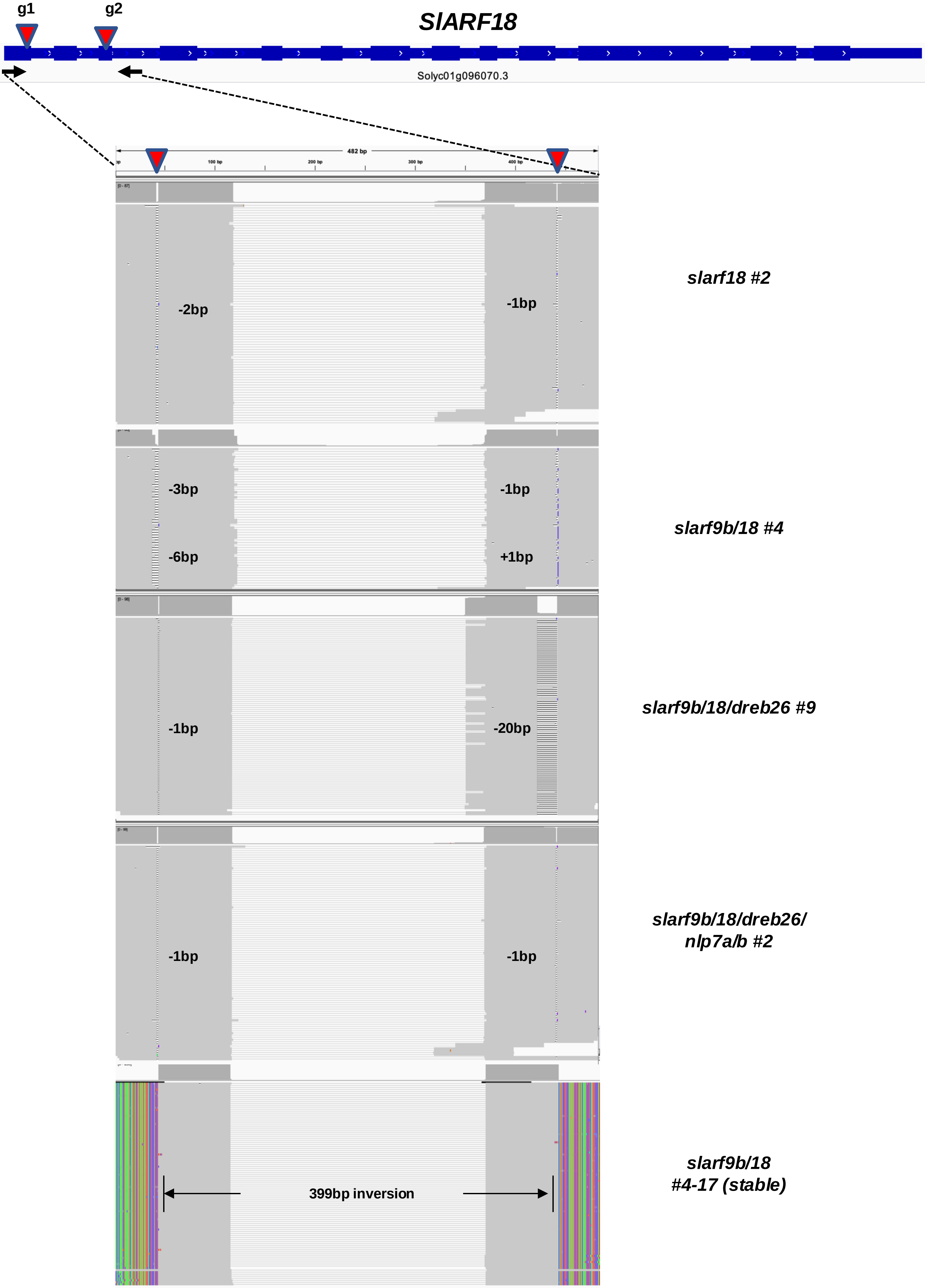

*Data supporting tomato CRISPR/Cas9 knockout lines (Continued).*

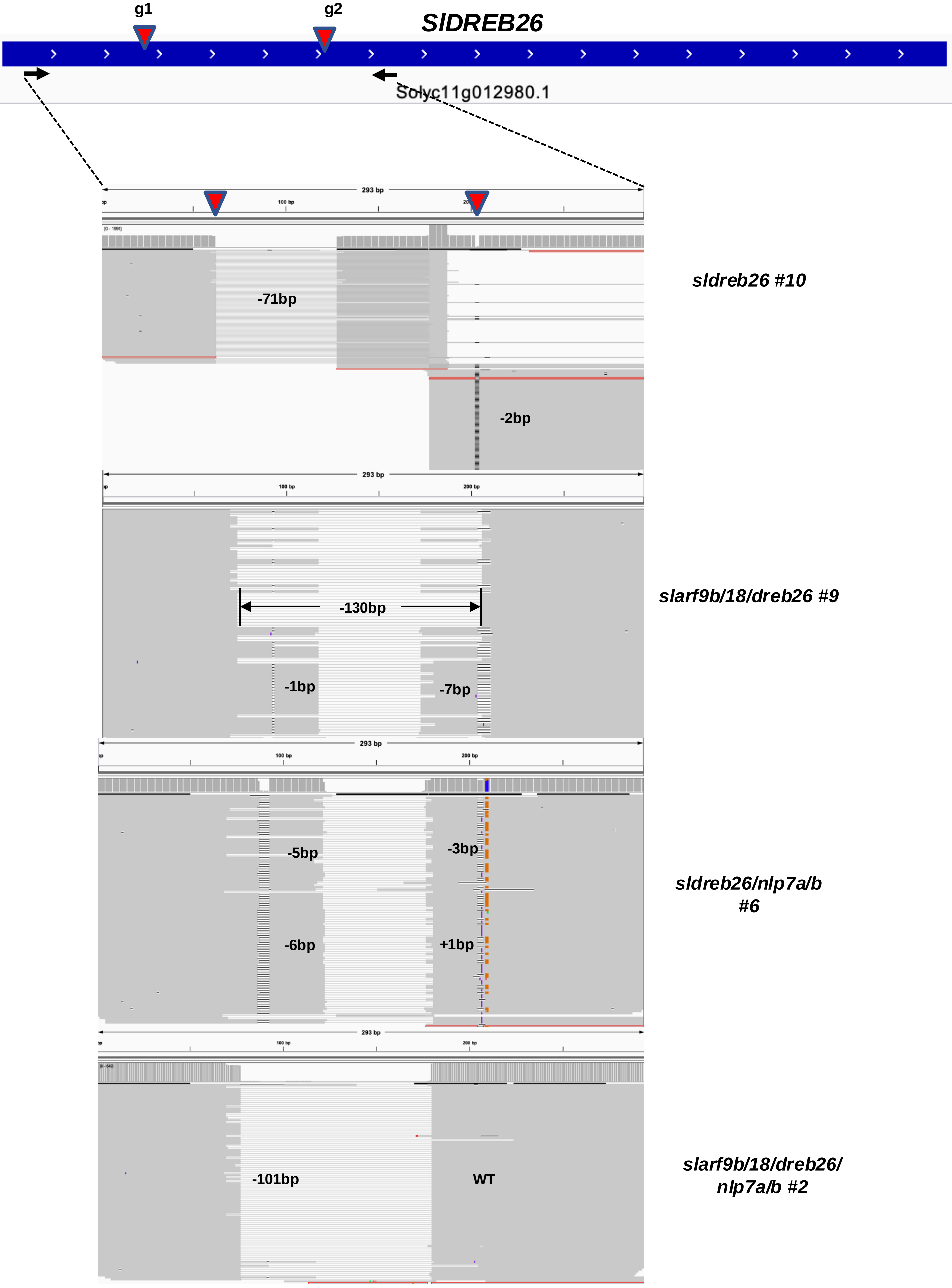

**Supplementary Data 12.** Genotypes of Arabidopsis and tomato CRISPR lines used in this study. *Data supporting tomato CRISPR/Cas9 knockout lines (Continued).*

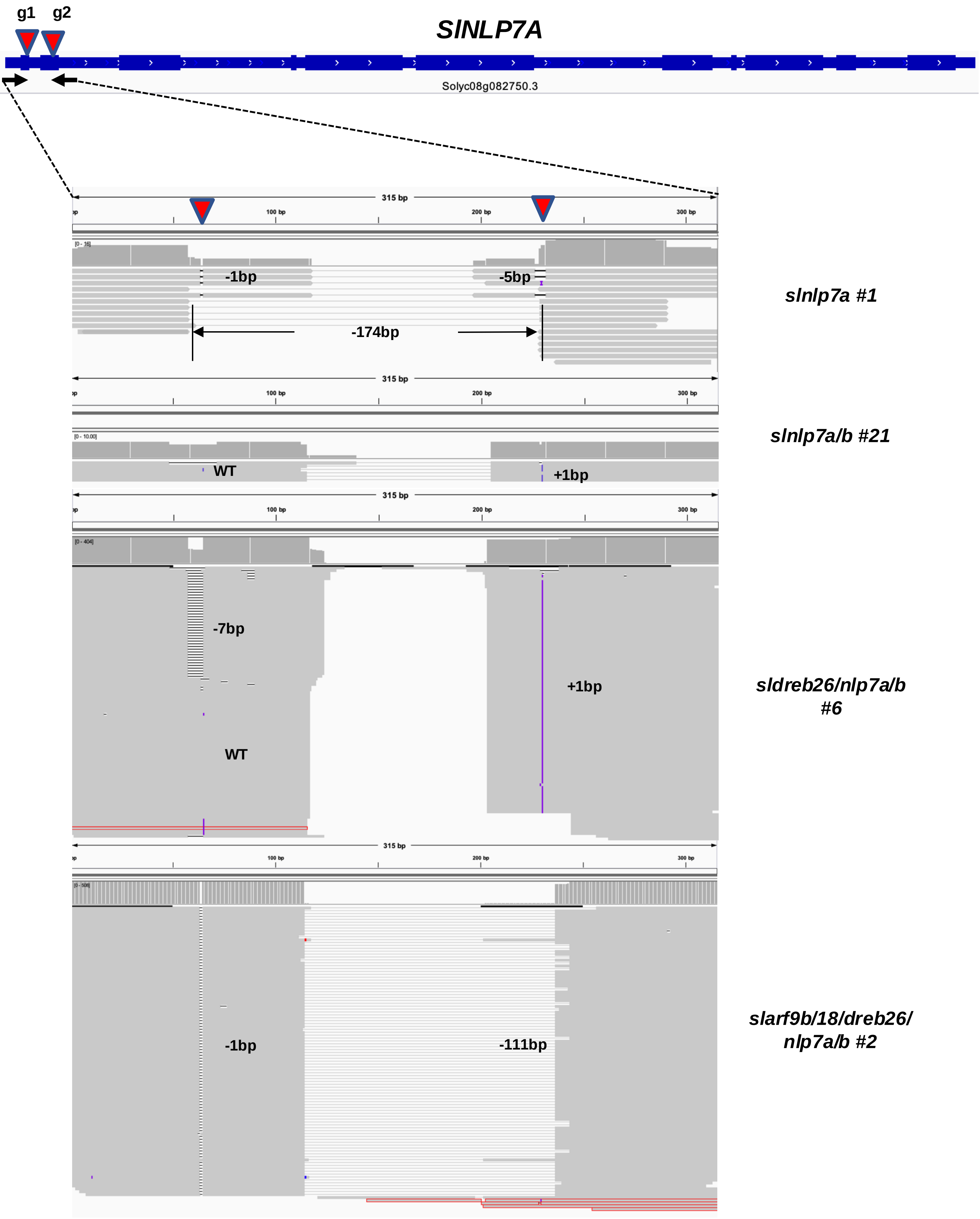

**Supplementary Data 12** Genotypes of Arabidopsis and tomato CRISPR lines used in this study. *Data supporting tomato CRISPR/Cas9 knockout lines (Continued).*

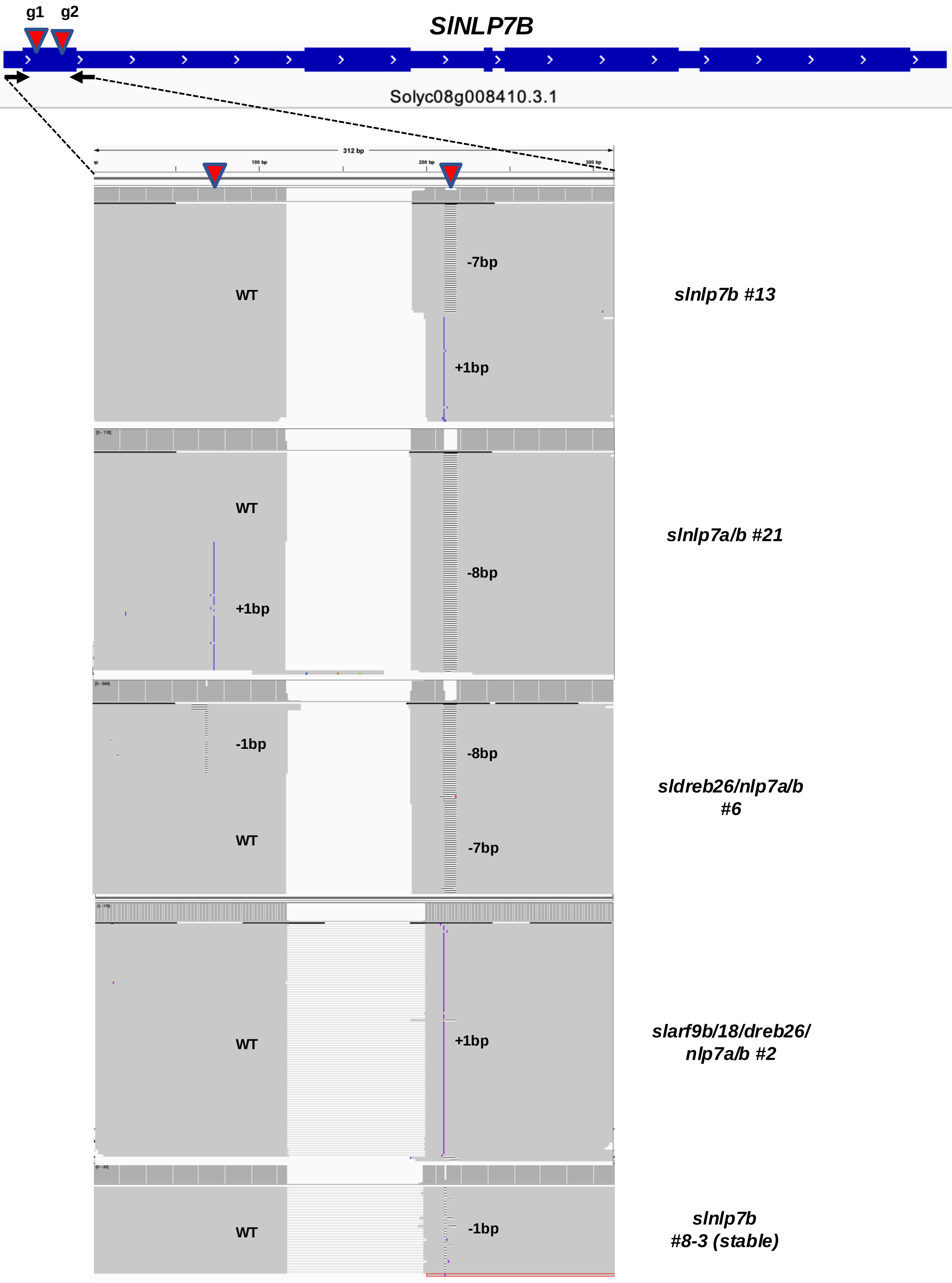

**Supplementary Data 13. Analysis of root system architecture of Arabidopsis and tomato mutant alleles of *NLP7, ARF18/9* and *ANAC032*.**
(a) RSA phenotypes of all Traits measured include the natural logarithm-transformed primary root length (logPR), number of lateral roots (logLR), total lateral root length (logLRL), average lateral root length (logALRL), total root length (logTRL), lateral root density (logLRD) and the ratio of lateral root length to total root length (logLRP). Traits were measured in 0, 1 and 10 mM $KNO_3$ for tomato, and 1 and 10 mM $KNO_3$ for Arabidopsis. Light green represents the trait as significantly smaller relative to wild type (wt), dark green represents the trait as significantly larger relative to wild type, white indicates no significant difference and grey indicates that this phenotype was not tested at this concentration (t-test, p<.05). A black bar indicates a significant genotype x treatment interaction as measured by a two-way ANOVA (p<.05).

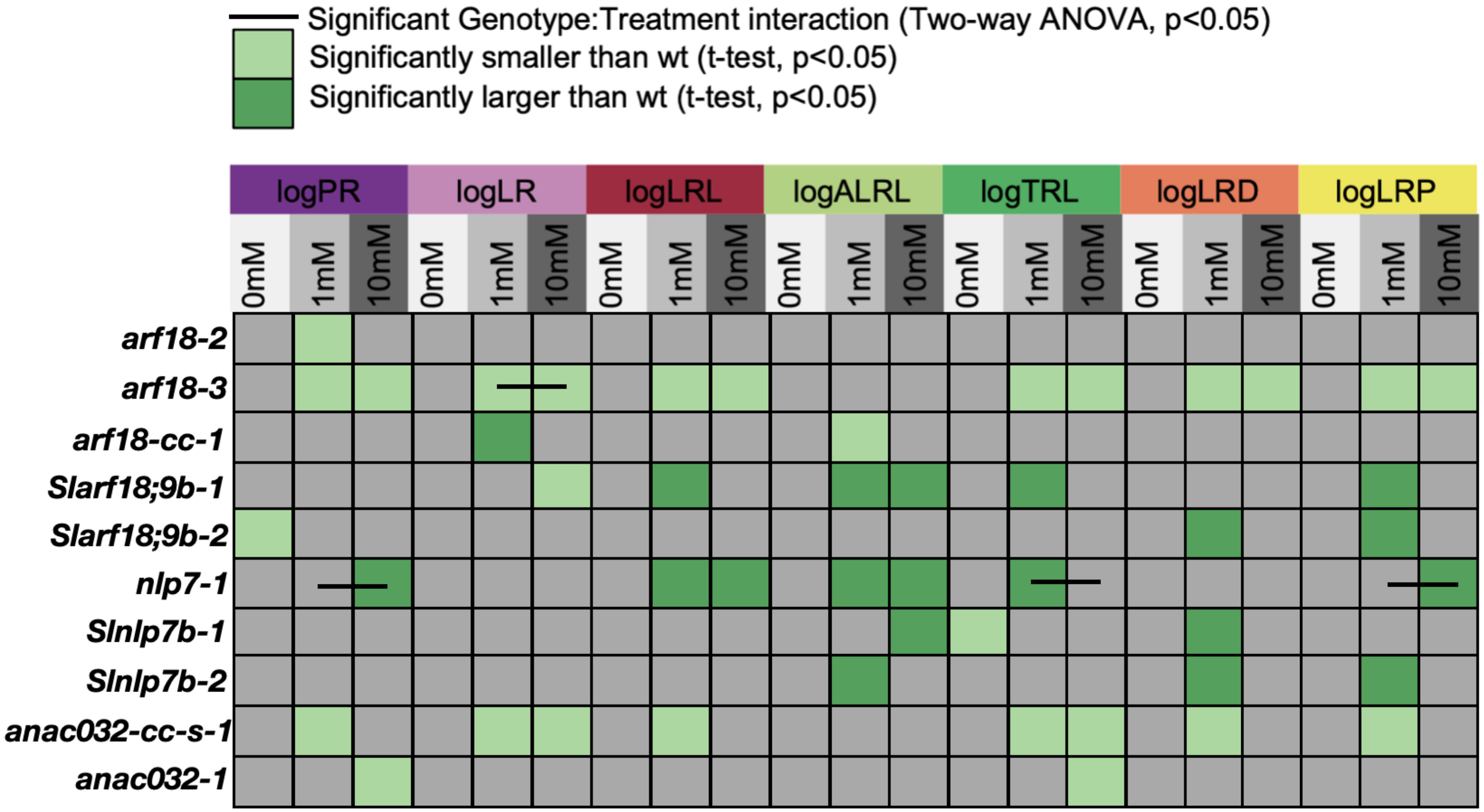

**(b) Root system architecture traits of Arabidopsis alleles of *NLP7* and *ARF18*.** Traits measured include the natural logarithm-transformed primary root length (logPR), number of lateral roots (logLR), total lateral root length (logLRL), average lateral root length (logALRL), total root length (logTRL), lateral root density (logLRD) and the ratio of lateral root length to total root length (logLRP). Traits were measured in 1 and 10 mM $KNO_3$. ***=p<.001, **=p<.01, *=p<.1.

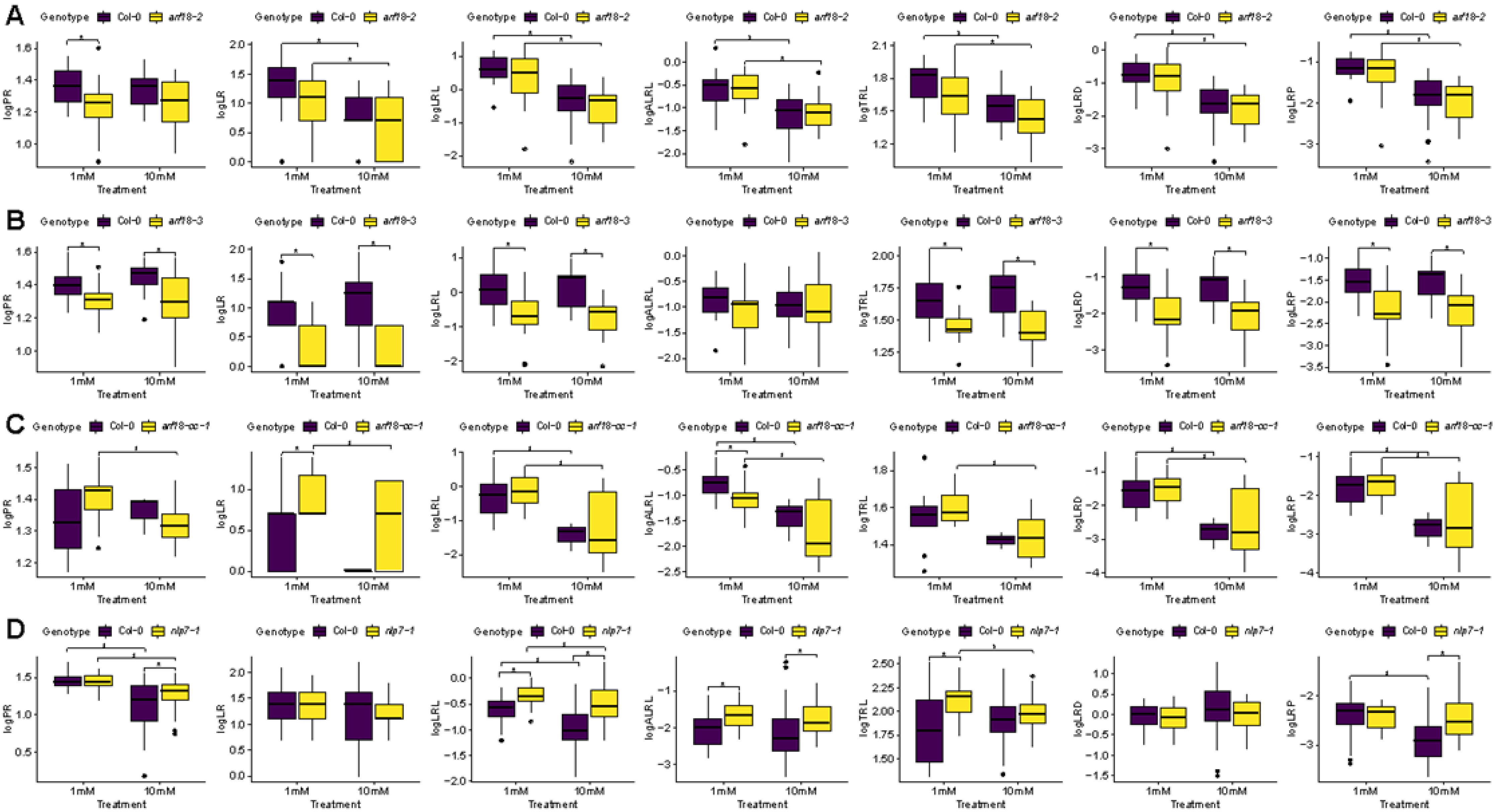

**(c) Arabidopsis *ANAC032* Mutant alleles display defects in root system architecture relative to wild type.** Traits measured include the natural logarithm-transformed primary root length (logPR), number of lateral roots (logLR), total lateral root length (logLRL), average lateral root length (logALRL), total root length (logTRL), lateral root density (logLRD) and the ratio of lateral root length to total root length (logLRP). Traits were measured in 1 and 10 mM $KNO_3$. ***=p<.001, **=p<.01, *=p<.1.

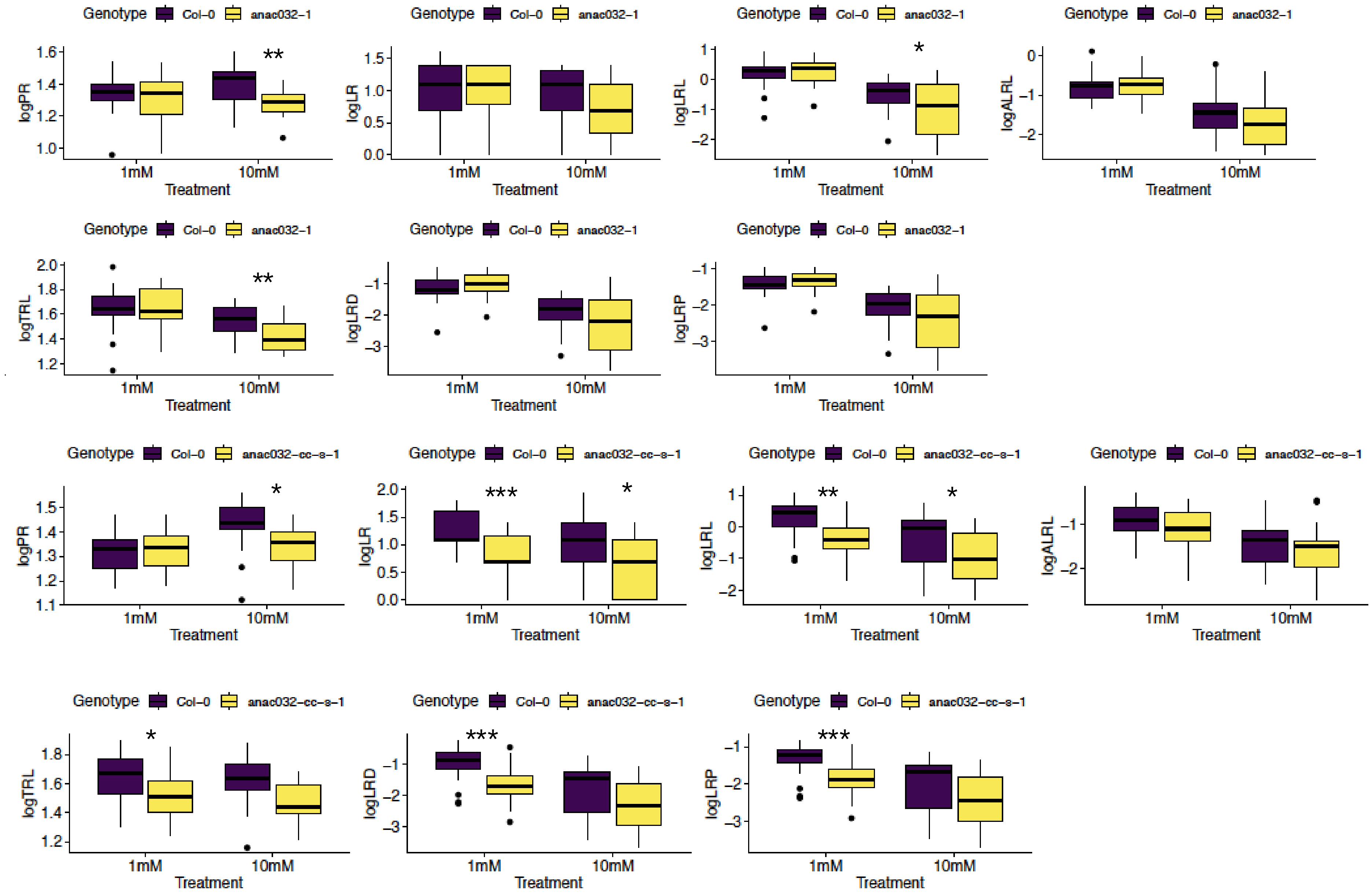

**(d) Analysis of root system architecture of tomato mutant alleles of *NLP7* and *ARF18*.** Traits measured include the natural logarithm-transformed primary root length (logPR), number of lateral roots (logLR), total lateral root length (logLRL), average lateral root length (logALRL), total root length (logTRL), lateral root density (logLRD) and the ratio of lateral root length to total root length (logLRP). ). Traits were measured in 0, 1 and 10 mM $KNO_3$.

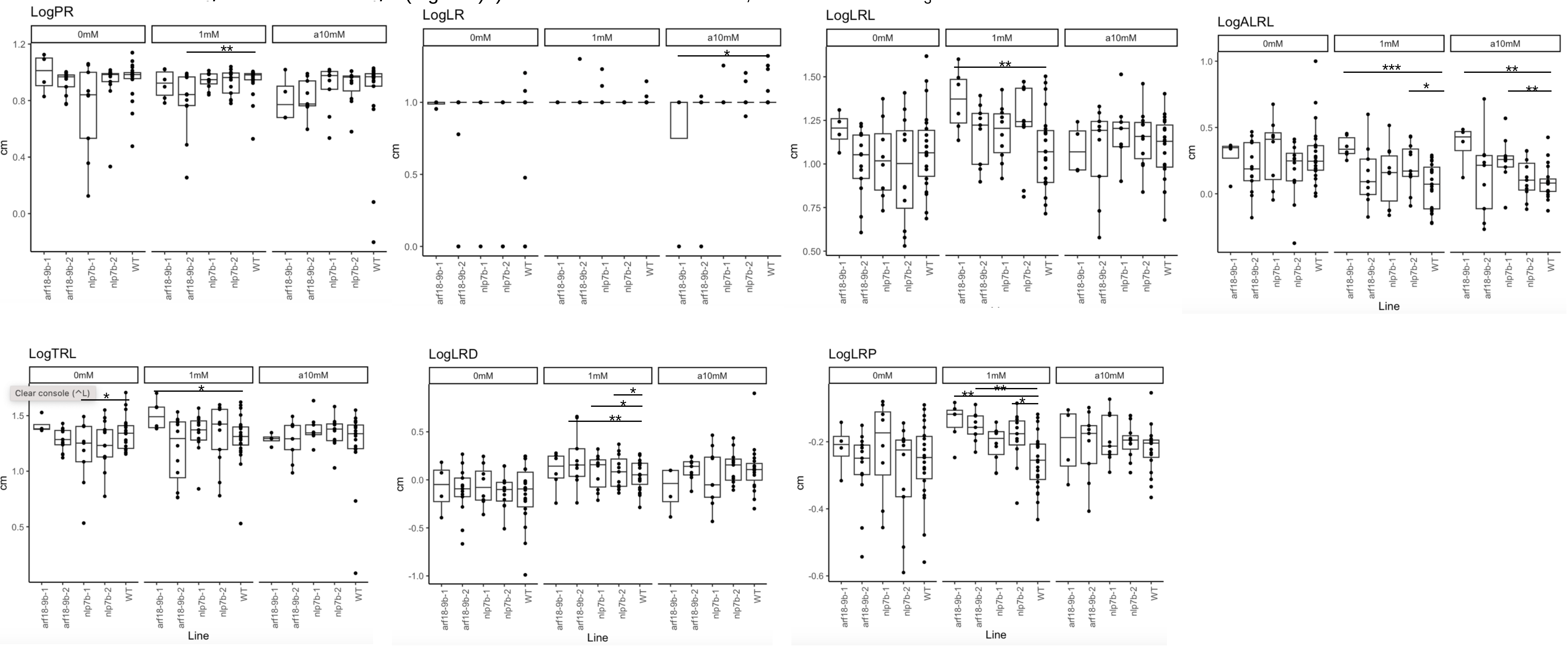

```
##################################################
#                                                #
#T-tests between each genotype within a treatment#
#                                                #
##################################################
[1] "anac032-1"
$logPR

	Two Sample t-test

data:  x by dataset_anac0321_1mM$Genotype
t = -0.44838, df = 36, p-value = 0.6566
alternative hypothesis: true difference in means between group anac032-1 
and group Col-0 is not equal to 0
95 percent confidence interval:
 -0.11119815  0.07093187
sample estimates:
mean in group anac032-1     mean in group Col-0 
               1.325065                1.345198 

$logLR

	Two Sample t-test

data:  x by dataset_anac0321_1mM$Genotype
t = 0.22151, df = 36, p-value = 0.8259
alternative hypothesis: true difference in means between group anac032-1 
and group Col-0 is not equal to 0
95 percent confidence interval:
 -0.2899639  0.3610712
sample estimates:
mean in group anac032-1     mean in group Col-0 
               1.036825                1.001272 

$logLRL

	Two Sample t-test

data:  x by dataset_anac0321_1mM$Genotype
t = 0.69392, df = 36, p-value = 0.4922
alternative hypothesis: true difference in means between group anac032-1 
and group Col-0 is not equal to 0
95 percent confidence interval:
 -0.2053797  0.4190221
sample estimates:
mean in group anac032-1     mean in group Col-0 
              0.2964885               0.1896673 

$logALRL

	Two Sample t-test

data:  x by dataset_anac0321_1mM$Genotype
t = 0.58434, df = 36, p-value = 0.5626
```

```
alternative hypothesis: true difference in means between group anac032-1
and group Col-0 is not equal to 0
95 percent confidence interval:
 -0.176082  0.318617
sample estimates:
mean in group anac032-1     mean in group Col-0
             -0.7403369              -0.8116044

$logTRL

	Two Sample t-test

data:  x by dataset_anac0321_1mM$Genotype
t = 0.17596, df = 36, p-value = 0.8613
alternative hypothesis: true difference in means between group anac032-1
and group Col-0 is not equal to 0
95 percent confidence interval:
 -0.1050800  0.1250457
sample estimates:
mean in group anac032-1     mean in group Col-0
               1.645463                1.635480

$logLRD

	Two Sample t-test

data:  x by dataset_anac0321_1mM$Genotype
t = 0.89401, df = 36, p-value = 0.3773
alternative hypothesis: true difference in means between group anac032-1
and group Col-0 is not equal to 0
95 percent confidence interval:
 -0.1610473  0.4149559
sample estimates:
mean in group anac032-1     mean in group Col-0
              -1.028577               -1.155531

$logLRP

	Two Sample t-test

data:  x by dataset_anac0321_1mM$Genotype
t = 0.87435, df = 36, p-value = 0.3877
alternative hypothesis: true difference in means between group anac032-1
and group Col-0 is not equal to 0
95 percent confidence interval:
 -0.1277836  0.3214602
sample estimates:
mean in group anac032-1     mean in group Col-0
              -1.348974               -1.445813

$logPR

	Two Sample t-test
```

```
data:  x by dataset_anac0321_10mM$Genotype
t = -2.8449, df = 31, p-value = 0.007802
alternative hypothesis: true difference in means between group anac032-1 
and group Col-0 is not equal to 0
95 percent confidence interval:
 -0.18490163 -0.03048934
sample estimates:
mean in group anac032-1     mean in group Col-0 
               1.285574                1.393269 

$logLR

      Two Sample t-test

data:  x by dataset_anac0321_10mM$Genotype
t = -1.1772, df = 31, p-value = 0.2481
alternative hypothesis: true difference in means between group anac032-1 
and group Col-0 is not equal to 0
95 percent confidence interval:
 -0.5491565  0.1472122
sample estimates:
mean in group anac032-1     mean in group Col-0 
              0.6818206               0.8827927 

$logLRL

      Two Sample t-test

data:  x by dataset_anac0321_10mM$Genotype
t = -1.8481, df = 31, p-value = 0.07415
alternative hypothesis: true difference in means between group anac032-1 
and group Col-0 is not equal to 0
95 percent confidence interval:
 -1.09057200  0.05370227
sample estimates:
mean in group anac032-1     mean in group Col-0 
             -1.0349740              -0.5165391 

$logALRL

      Two Sample t-test

data:  x by dataset_anac0321_10mM$Genotype
t = -1.4433, df = 31, p-value = 0.159
alternative hypothesis: true difference in means between group anac032-1 
and group Col-0 is not equal to 0
95 percent confidence interval:
 -0.7660784  0.1311530
sample estimates:
mean in group anac032-1     mean in group Col-0 
              -1.716795               -1.399332 

$logTRL
```

```
      Two Sample t-test

data:  x by dataset_anac0321_10mM$Genotype
t = -2.8313, df = 31, p-value = 0.00807
alternative hypothesis: true difference in means between group anac032-1
and group Col-0 is not equal to 0
95 percent confidence interval:
 -0.21653012 -0.03519891
sample estimates:
mean in group anac032-1     mean in group Col-0
               1.420799                1.546664

$logLRD

      Two Sample t-test

data:  x by dataset_anac0321_10mM$Genotype
t = -1.4607, df = 31, p-value = 0.1542
alternative hypothesis: true difference in means between group anac032-1
and group Col-0 is not equal to 0
95 percent confidence interval:
 -0.9842428  0.1627641
sample estimates:
mean in group anac032-1     mean in group Col-0
              -2.320548               -1.909808

$logLRP

      Two Sample t-test

data:  x by dataset_anac0321_10mM$Genotype
t = -1.5616, df = 31, p-value = 0.1285
alternative hypothesis: true difference in means between group anac032-1
and group Col-0 is not equal to 0
95 percent confidence interval:
 -0.9052967  0.1201560
sample estimates:
mean in group anac032-1     mean in group Col-0
              -2.455773               -2.063203

[1] "anac032-cc-s-1"
$logPR

      Two Sample t-test

data:  x by dataset_anac032ccs1_1mM$Genotype
t = 0.38095, df = 39, p-value = 0.7053
alternative hypothesis: true difference in means between group anac032-
cc-s-1 and group Col-0 is not equal to 0
95 percent confidence interval:
 -0.04253148  0.06226983
sample estimates:
mean in group anac032-cc-s-1          mean in group Col-0
                    1.324432                     1.314563
```

```
$logLR

	Two Sample t-test

data:  x by dataset_anac032ccs1_1mM$Genotype
t = -3.6585, df = 39, p-value = 0.0007487
alternative hypothesis: true difference in means between group anac032-
cc-s-1 and group Col-0 is not equal to 0
95 percent confidence interval:
 -0.7583294 -0.2183516
sample estimates:
mean in group anac032-cc-s-1          mean in group Col-0
                   0.7480778                    1.2364183

$logLRL

	Two Sample t-test

data:  x by dataset_anac032ccs1_1mM$Genotype
t = -3.4477, df = 39, p-value = 0.00137
alternative hypothesis: true difference in means between group anac032-
cc-s-1 and group Col-0 is not equal to 0
95 percent confidence interval:
 -1.0275769 -0.2676717
sample estimates:
mean in group anac032-cc-s-1          mean in group Col-0
                  -0.3584359                    0.2891883

$logALRL

	Two Sample t-test

data:  x by dataset_anac032ccs1_1mM$Genotype
t = -1.1288, df = 39, p-value = 0.2659
alternative hypothesis: true difference in means between group anac032-
cc-s-1 and group Col-0 is not equal to 0
95 percent confidence interval:
 -0.4446970  0.1261294
sample estimates:
mean in group anac032-cc-s-1          mean in group Col-0
                   -1.106514                    -0.947230

$logTRL

	Two Sample t-test

data:  x by dataset_anac032ccs1_1mM$Genotype
t = -2.6122, df = 39, p-value = 0.01271
alternative hypothesis: true difference in means between group anac032-
cc-s-1 and group Col-0 is not equal to 0
95 percent confidence interval:
 -0.23210999 -0.02951977
sample estimates:
```

```
mean in group anac032-cc-s-1          mean in group Col-0
                    1.516343                       1.647158

$logLRD

      Two Sample t-test

data:  x by dataset_anac032ccs1_1mM$Genotype
t = -3.7004, df = 39, p-value = 0.0006631
alternative hypothesis: true difference in means between group anac032-
cc-s-1 and group Col-0 is not equal to 0
95 percent confidence interval:
 -1.0168918 -0.2980951
sample estimates:
mean in group anac032-cc-s-1          mean in group Col-0
                   -1.682868                      -1.025374

$logLRP

      Two Sample t-test

data:  x by dataset_anac032ccs1_1mM$Genotype
t = -3.6135, df = 39, p-value = 0.0008529
alternative hypothesis: true difference in means between group anac032-
cc-s-1 and group Col-0 is not equal to 0
95 percent confidence interval:
 -0.8061010 -0.2275178
sample estimates:
mean in group anac032-cc-s-1          mean in group Col-0
                   -1.874779                      -1.357969

$logPR

      Two Sample t-test

data:  x by dataset_anac032ccs1_10mM$Genotype
t = -2.6932, df = 36, p-value = 0.01068
alternative hypothesis: true difference in means between group anac032-
cc-s-1 and group Col-0 is not equal to 0
95 percent confidence interval:
 -0.14891225 -0.02097682
sample estimates:
mean in group anac032-cc-s-1          mean in group Col-0
                    1.346393                       1.431337

$logLR

      Two Sample t-test

data:  x by dataset_anac032ccs1_10mM$Genotype
t = -2.0124, df = 36, p-value = 0.0517
alternative hypothesis: true difference in means between group anac032-
cc-s-1 and group Col-0 is not equal to 0
95 percent confidence interval:
```

```
 -0.741937595  0.002876125
sample estimates:
mean in group anac032-cc-s-1          mean in group Col-0
                   0.6325825                    1.0021133

$logLRL

	Two Sample t-test

data:  x by dataset_anac032ccs1_10mM$Genotype
t = -1.8511, df = 36, p-value = 0.07237
alternative hypothesis: true difference in means between group anac032-cc-s-1 and group Col-0 is not equal to 0
95 percent confidence interval:
 -1.1077716  0.0505435
sample estimates:
mean in group anac032-cc-s-1          mean in group Col-0
                  -0.9795225                   -0.4509084

$logALRL

	Two Sample t-test

data:  x by dataset_anac032ccs1_10mM$Genotype
t = -0.84036, df = 36, p-value = 0.4063
alternative hypothesis: true difference in means between group anac032-cc-s-1 and group Col-0 is not equal to 0
95 percent confidence interval:
 -0.5430069  0.2248403
sample estimates:
mean in group anac032-cc-s-1          mean in group Col-0
                   -1.612105                    -1.453022

$logTRL

	Two Sample t-test

data:  x by dataset_anac032ccs1_10mM$Genotype
t = -2.7993, df = 36, p-value = 0.00818
alternative hypothesis: true difference in means between group anac032-cc-s-1 and group Col-0 is not equal to 0
95 percent confidence interval:
 -0.24548121 -0.03921632
sample estimates:
mean in group anac032-cc-s-1          mean in group Col-0
                    1.467237                     1.609585

$logLRD

	Two Sample t-test

data:  x by dataset_anac032ccs1_10mM$Genotype
t = -1.6189, df = 36, p-value = 0.1142
```

```
alternative hypothesis: true difference in means between group anac032-
cc-s-1 and group Col-0 is not equal to 0
95 percent confidence interval:
 -0.9994937  0.1121547
sample estimates:
mean in group anac032-cc-s-1          mean in group Col-0
                   -2.325915                    -1.882246

$logLRP

	Two Sample t-test

data:  x by dataset_anac032ccs1_10mM$Genotype
t = -1.5834, df = 36, p-value = 0.1221
alternative hypothesis: true difference in means between group anac032-
cc-s-1 and group Col-0 is not equal to 0
95 percent confidence interval:
 -0.8810091  0.1084786
sample estimates:
mean in group anac032-cc-s-1          mean in group Col-0
                   -2.446759                    -2.060494

[1] "arf18-2"
$logPR

	Two Sample t-test

data:  x by dataset_arf182_1mM$Genotype
t = -2.5094, df = 35, p-value = 0.01687
alternative hypothesis: true difference in means between group arf18-2
and group Col-0 is not equal to 0
95 percent confidence interval:
 -0.21236721 -0.02242129
sample estimates:
mean in group arf18-2   mean in group Col-0
             1.236667              1.354062

$logLR

	Two Sample t-test

data:  x by dataset_arf182_1mM$Genotype
t = -1.6946, df = 35, p-value = 0.09903
alternative hypothesis: true difference in means between group arf18-2
and group Col-0 is not equal to 0
95 percent confidence interval:
 -0.63566534  0.05726017
sample estimates:
mean in group arf18-2   mean in group Col-0
            0.9271543             1.2163568

$logLRL

	Two Sample t-test
```

```
data:  x by dataset_arf182_1mM$Genotype
t = -1.3791, df = 35, p-value = 0.1766
alternative hypothesis: true difference in means between group arf18-2
and group Col-0 is not equal to 0
95 percent confidence interval:
 -0.7159524  0.1367134
sample estimates:
mean in group arf18-2   mean in group Col-0
            0.3164538             0.6060733

$logALRL

	Two Sample t-test

data:  x by dataset_arf182_1mM$Genotype
t = -0.0030519, df = 35, p-value = 0.9976
alternative hypothesis: true difference in means between group arf18-2
and group Col-0 is not equal to 0
95 percent confidence interval:
 -0.2777559  0.2769221
sample estimates:
mean in group arf18-2   mean in group Col-0
           -0.6107005            -0.6102835

$logTRL

	Two Sample t-test

data:  x by dataset_arf182_1mM$Genotype
t = -1.8592, df = 35, p-value = 0.07143
alternative hypothesis: true difference in means between group arf18-2
and group Col-0 is not equal to 0
95 percent confidence interval:
 -0.28466941  0.01251203
sample estimates:
mean in group arf18-2   mean in group Col-0
             1.621235              1.757313

$logLRD

	Two Sample t-test

data:  x by dataset_arf182_1mM$Genotype
t = -0.86333, df = 35, p-value = 0.3938
alternative hypothesis: true difference in means between group arf18-2
and group Col-0 is not equal to 0
95 percent confidence interval:
 -0.5772088  0.2327582
sample estimates:
mean in group arf18-2   mean in group Col-0
           -0.9202135            -0.7479883

$logLRP
```

```
	Two Sample t-test

data:  x by dataset_arf182_1mM$Genotype
t = -1.0173, df = 35, p-value = 0.316
alternative hypothesis: true difference in means between group arf18-2
and group Col-0 is not equal to 0
95 percent confidence interval:
 -0.4599348  0.1528532
sample estimates:
mean in group arf18-2   mean in group Col-0 
            -1.304781             -1.151240 

$logPR

	Two Sample t-test

data:  x by dataset_arf182_10mM$Genotype
t = -1.9774, df = 36, p-value = 0.05569
alternative hypothesis: true difference in means between group arf18-2
and group Col-0 is not equal to 0
95 percent confidence interval:
 -0.170998826  0.002162052
sample estimates:
mean in group arf18-2   mean in group Col-0 
             1.252731              1.337149 

$logLR

	Two Sample t-test

data:  x by dataset_arf182_10mM$Genotype
t = -1.0199, df = 36, p-value = 0.3146
alternative hypothesis: true difference in means between group arf18-2
and group Col-0 is not equal to 0
95 percent confidence interval:
 -0.4998818  0.1653498
sample estimates:
mean in group arf18-2   mean in group Col-0 
            0.5809599             0.7482259 

$logLRL

	Two Sample t-test

data:  x by dataset_arf182_10mM$Genotype
t = -0.57636, df = 36, p-value = 0.568
alternative hypothesis: true difference in means between group arf18-2
and group Col-0 is not equal to 0
95 percent confidence interval:
 -0.5850152  0.3260900
sample estimates:
mean in group arf18-2   mean in group Col-0 
           -0.5204753            -0.3910127 
```

```
$logALRL

      Two Sample t-test

data:  x by dataset_arf182_10mM$Genotype
t = 0.28284, df = 36, p-value = 0.7789
alternative hypothesis: true difference in means between group arf18-2 and group Col-0 is not equal to 0
95 percent confidence interval:
 -0.2332669  0.3088736
sample estimates:
mean in group arf18-2   mean in group Col-0
            -1.101435             -1.139239

$logTRL

      Two Sample t-test

data:  x by dataset_arf182_10mM$Genotype
t = -1.7298, df = 36, p-value = 0.09223
alternative hypothesis: true difference in means between group arf18-2 and group Col-0 is not equal to 0
95 percent confidence interval:
 -0.22696116  0.01801652
sample estimates:
mean in group arf18-2   mean in group Col-0
             1.424491              1.528964

$logLRD

      Two Sample t-test

data:  x by dataset_arf182_10mM$Genotype
t = -0.22118, df = 36, p-value = 0.8262
alternative hypothesis: true difference in means between group arf18-2 and group Col-0 is not equal to 0
95 percent confidence interval:
 -0.4580752  0.3679867
sample estimates:
mean in group arf18-2   mean in group Col-0
            -1.773206             -1.728162

$logLRP

      Two Sample t-test

data:  x by dataset_arf182_10mM$Genotype
t = -0.14147, df = 36, p-value = 0.8883
alternative hypothesis: true difference in means between group arf18-2 and group Col-0 is not equal to 0
95 percent confidence interval:
 -0.3832423  0.3332617
sample estimates:
mean in group arf18-2   mean in group Col-0
```

```
           -1.944967             -1.919976

[1] "arf18-3"
$logPR

	Two Sample t-test

data:  x by dataset_arf183_1mM$Genotype
t = -2.5794, df = 34, p-value = 0.0144
alternative hypothesis: true difference in means between group arf18-3 and group Col-0 is not equal to 0
95 percent confidence interval:
 -0.14395360 -0.01707881
sample estimates:
mean in group arf18-3   mean in group Col-0 
             1.309138              1.389654 

$logLR

	Two Sample t-test

data:  x by dataset_arf183_1mM$Genotype
t = -3.916, df = 34, p-value = 0.0004117
alternative hypothesis: true difference in means between group arf18-3 and group Col-0 is not equal to 0
95 percent confidence interval:
 -0.848039 -0.268569
sample estimates:
mean in group arf18-3   mean in group Col-0 
            0.4045617             0.9628657 

$logLRL

	Two Sample t-test

data:  x by dataset_arf183_1mM$Genotype
t = -3.8893, df = 34, p-value = 0.0004442
alternative hypothesis: true difference in means between group arf18-3 and group Col-0 is not equal to 0
95 percent confidence interval:
 -1.2522816 -0.3927341
sample estimates:
mean in group arf18-3   mean in group Col-0 
           -0.7165428             0.1059651 

$logALRL

	Two Sample t-test

data:  x by dataset_arf183_1mM$Genotype
t = -1.6862, df = 34, p-value = 0.1009
alternative hypothesis: true difference in means between group arf18-3 and group Col-0 is not equal to 0
95 percent confidence interval:
```

```
 -0.5826310  0.0542232
sample estimates:
mean in group arf18-3   mean in group Col-0
           -1.1211045            -0.8569006

$logTRL

      Two Sample t-test

data:  x by dataset_arf183_1mM$Genotype
t = -3.7202, df = 34, p-value = 0.0007163
alternative hypothesis: true difference in means between group arf18-3
and group Col-0 is not equal to 0
95 percent confidence interval:
 -0.30838354 -0.09049128
sample estimates:
mean in group arf18-3   mean in group Col-0
             1.455710              1.655147

$logLRD

      Two Sample t-test

data:  x by dataset_arf183_1mM$Genotype
t = -3.6928, df = 34, p-value = 0.0007734
alternative hypothesis: true difference in means between group arf18-3
and group Col-0 is not equal to 0
95 percent confidence interval:
 -1.150324 -0.333659
sample estimates:
mean in group arf18-3   mean in group Col-0
            -2.025680             -1.283689

$logLRP

      Two Sample t-test

data:  x by dataset_arf183_1mM$Genotype
t = -3.6971, df = 34, p-value = 0.0007643
alternative hypothesis: true difference in means between group arf18-3
and group Col-0 is not equal to 0
95 percent confidence interval:
 -0.9655634 -0.2805776
sample estimates:
mean in group arf18-3   mean in group Col-0
            -2.172253             -1.549182

$logPR

      Two Sample t-test

data:  x by dataset_arf183_10mM$Genotype
t = -2.6374, df = 27, p-value = 0.01369
```

```
alternative hypothesis: true difference in means between group arf18-3
and group Col-0 is not equal to 0
95 percent confidence interval:
 -0.25510414 -0.03185486
sample estimates:
mean in group arf18-3   mean in group Col-0
             1.296939              1.440418

$logLR

	Two Sample t-test

data:  x by dataset_arf183_10mM$Genotype
t = -5.527, df = 27, p-value = 7.419e-06
alternative hypothesis: true difference in means between group arf18-3
and group Col-0 is not equal to 0
95 percent confidence interval:
 -1.2643030 -0.5797303
sample estimates:
mean in group arf18-3   mean in group Col-0
            0.2132761             1.1352927

$logLRL

	Two Sample t-test

data:  x by dataset_arf183_10mM$Genotype
t = -4.3528, df = 27, p-value = 0.0001731
alternative hypothesis: true difference in means between group arf18-3
and group Col-0 is not equal to 0
95 percent confidence interval:
 -1.3662865 -0.4908565
sample estimates:
mean in group arf18-3   mean in group Col-0
           -0.7635761             0.1649954

$logALRL

	Two Sample t-test

data:  x by dataset_arf183_10mM$Genotype
t = -0.035108, df = 27, p-value = 0.9723
alternative hypothesis: true difference in means between group arf18-3
and group Col-0 is not equal to 0
95 percent confidence interval:
 -0.3896403  0.3765306
sample estimates:
mean in group arf18-3   mean in group Col-0
           -0.9768521            -0.9702973

$logTRL

	Two Sample t-test
```

```
data:  x by dataset_arf183_10mM$Genotype
t = -4.4662, df = 27, p-value = 0.0001278
alternative hypothesis: true difference in means between group arf18-3
and group Col-0 is not equal to 0
95 percent confidence interval:
 -0.3953973 -0.1464592
sample estimates:
mean in group arf18-3   mean in group Col-0
             1.436161              1.707089

$logLRD

      Two Sample t-test

data:  x by dataset_arf183_10mM$Genotype
t = -3.5963, df = 27, p-value = 0.001274
alternative hypothesis: true difference in means between group arf18-3
and group Col-0 is not equal to 0
95 percent confidence interval:
 -1.2330157 -0.3371684
sample estimates:
mean in group arf18-3   mean in group Col-0
            -2.060515             -1.275423

$logLRP

      Two Sample t-test

data:  x by dataset_arf183_10mM$Genotype
t = -3.5431, df = 27, p-value = 0.001462
alternative hypothesis: true difference in means between group arf18-3
and group Col-0 is not equal to 0
95 percent confidence interval:
 -1.0384899 -0.2767966
sample estimates:
mean in group arf18-3   mean in group Col-0
            -2.199737             -1.542093

[1] "arf18-cc-1"
$logPR

      Two Sample t-test

data:  x by dataset_arf18cc1_1mM$Genotype
t = 1.9506, df = 27, p-value = 0.06155
alternative hypothesis: true difference in means between group arf18-cc-1
and group Col-0 is not equal to 0
95 percent confidence interval:
 -0.003501654  0.138480286
sample estimates:
mean in group arf18-cc-1      mean in group Col-0
                1.398978                 1.331489

$logLR
```

```
      Two Sample t-test

data:  x by dataset_arf18cc1_1mM$Genotype
t = 2.803, df = 27, p-value = 0.009259
alternative hypothesis: true difference in means between group arf18-cc-1
and group Col-0 is not equal to 0
95 percent confidence interval:
 0.1088146 0.7032982
sample estimates:
mean in group arf18-cc-1      mean in group Col-0
               0.9171171                0.5110607

$logLRL

      Two Sample t-test

data:  x by dataset_arf18cc1_1mM$Genotype
t = 0.52646, df = 27, p-value = 0.6029
alternative hypothesis: true difference in means between group arf18-cc-1
and group Col-0 is not equal to 0
95 percent confidence interval:
 -0.3046678  0.5149699
sample estimates:
mean in group arf18-cc-1      mean in group Col-0
               -0.170124                -0.275275

$logALRL

      Two Sample t-test

data:  x by dataset_arf18cc1_1mM$Genotype
t = -2.5738, df = 27, p-value = 0.01587
alternative hypothesis: true difference in means between group arf18-cc-1
and group Col-0 is not equal to 0
95 percent confidence interval:
 -0.54078812 -0.06102247
sample estimates:
mean in group arf18-cc-1      mean in group Col-0
              -1.0872411               -0.7863358

$logTRL

      Two Sample t-test

data:  x by dataset_arf18cc1_1mM$Genotype
t = 1.422, df = 27, p-value = 0.1665
alternative hypothesis: true difference in means between group arf18-cc-1
and group Col-0 is not equal to 0
95 percent confidence interval:
 -0.02966532  0.16361131
sample estimates:
mean in group arf18-cc-1      mean in group Col-0
                1.603999                 1.537026
```

```
$logLRD

	Two Sample t-test

data:  x by dataset_arf18cc1_1mM$Genotype
t = 0.18704, df = 27, p-value = 0.853
alternative hypothesis: true difference in means between group arf18-cc-1
and group Col-0 is not equal to 0
95 percent confidence interval:
 -0.3754968  0.4508203
sample estimates:
mean in group arf18-cc-1      mean in group Col-0
               -1.569102                -1.606764

$logLRP

	Two Sample t-test

data:  x by dataset_arf18cc1_1mM$Genotype
t = 0.22933, df = 27, p-value = 0.8203
alternative hypothesis: true difference in means between group arf18-cc-1
and group Col-0 is not equal to 0
95 percent confidence interval:
 -0.3034047  0.3797609
sample estimates:
mean in group arf18-cc-1      mean in group Col-0
               -1.774123                -1.812301

$logPR

	Two Sample t-test

data:  x by dataset_arf18cc1_10mM$Genotype
t = -0.9226, df = 12, p-value = 0.3744
alternative hypothesis: true difference in means between group arf18-cc-1
and group Col-0 is not equal to 0
95 percent confidence interval:
 -0.13930418  0.05642454
sample estimates:
mean in group arf18-cc-1      mean in group Col-0
                1.318437                 1.359877

$logLR

	Two Sample t-test

data:  x by dataset_arf18cc1_10mM$Genotype
t = 1.6852, df = 12, p-value = 0.1178
alternative hypothesis: true difference in means between group arf18-cc-1
and group Col-0 is not equal to 0
95 percent confidence interval:
 -0.1539401  1.2049844
sample estimates:
mean in group arf18-cc-1      mean in group Col-0
```

```
              0.5255221               0.0000000

$logLRL

	Two Sample t-test

data:  x by dataset_arf18cc1_10mM$Genotype
t = 0.52214, df = 12, p-value = 0.6111
alternative hypothesis: true difference in means between group arf18-cc-1 and group Col-0 is not equal to 0
95 percent confidence interval:
 -1.016252  1.656844
sample estimates:
mean in group arf18-cc-1      mean in group Col-0 
              -1.119129               -1.439425 

$logALRL

	Two Sample t-test

data:  x by dataset_arf18cc1_10mM$Genotype
t = -0.50311, df = 12, p-value = 0.624
alternative hypothesis: true difference in means between group arf18-cc-1 and group Col-0 is not equal to 0
95 percent confidence interval:
 -1.0939946  0.6835428
sample estimates:
mean in group arf18-cc-1      mean in group Col-0 
              -1.644651               -1.439425 

$logTRL

	Two Sample t-test

data:  x by dataset_arf18cc1_10mM$Genotype
t = 0.22598, df = 12, p-value = 0.825
alternative hypothesis: true difference in means between group arf18-cc-1 and group Col-0 is not equal to 0
95 percent confidence interval:
 -0.1494229  0.1840043
sample estimates:
mean in group arf18-cc-1      mean in group Col-0 
               1.439899                1.422608 

$logLRD

	Two Sample t-test

data:  x by dataset_arf18cc1_10mM$Genotype
t = 0.58354, df = 12, p-value = 0.5703
alternative hypothesis: true difference in means between group arf18-cc-1 and group Col-0 is not equal to 0
95 percent confidence interval:
 -0.9889163  1.7123883
```

```
sample estimates:
mean in group arf18-cc-1      mean in group Col-0
               -2.437566                -2.799302

$logLRP

	Two Sample t-test

data:  x by dataset_arf18cc1_10mM$Genotype
t = 0.53984, df = 12, p-value = 0.5992
alternative hypothesis: true difference in means between group arf18-cc-1
and group Col-0 is not equal to 0
95 percent confidence interval:
 -0.9199339  1.5259449
sample estimates:
mean in group arf18-cc-1      mean in group Col-0
               -2.559028                -2.862033

[1] "nlp7-1"
$logPR

	Two Sample t-test

data:  x by dataset_nlp71_1mM$Genotype
t = 0.68971, df = 54, p-value = 0.4933
alternative hypothesis: true difference in means between group Col-0 and
group nlp7-1 is not equal to 0
95 percent confidence interval:
 -0.03670224  0.07519748
sample estimates:
 mean in group Col-0 mean in group nlp7-1
            1.462431             1.443183

$logLR

	Two Sample t-test

data:  x by dataset_nlp71_1mM$Genotype
t = 0.77956, df = 54, p-value = 0.4391
alternative hypothesis: true difference in means between group Col-0 and
group nlp7-1 is not equal to 0
95 percent confidence interval:
 -0.1138927  0.2588129
sample estimates:
 mean in group Col-0 mean in group nlp7-1
            1.431742             1.359282

$logLRL

	Two Sample t-test

data:  x by dataset_nlp71_1mM$Genotype
t = -4.5546, df = 54, p-value = 3.032e-05
```

```
alternative hypothesis: true difference in means between group Col-0 and
group nlp7-1 is not equal to 0
95 percent confidence interval:
 -0.4072426 -0.1582991
sample estimates:
 mean in group Col-0 mean in group nlp7-1
          -0.6100701           -0.3272993

$logALRL

      Two Sample t-test

data:  x by dataset_nlp71_1mM$Genotype
t = -3.0911, df = 54, p-value = 0.003152
alternative hypothesis: true difference in means between group Col-0 and
group nlp7-1 is not equal to 0
95 percent confidence interval:
 -0.5856357 -0.1248262
sample estimates:
 mean in group Col-0 mean in group nlp7-1
           -2.041812            -1.686581

$logTRL

      Two Sample t-test

data:  x by dataset_nlp71_1mM$Genotype
t = -3.8135, df = 54, p-value = 0.0003539
alternative hypothesis: true difference in means between group Col-0 and
group nlp7-1 is not equal to 0
95 percent confidence interval:
 -0.4509091 -0.1401640
sample estimates:
 mean in group Col-0 mean in group nlp7-1
            1.811622             2.107158

$logLRD

      Two Sample t-test

data:  x by dataset_nlp71_1mM$Genotype
t = 0.64354, df = 54, p-value = 0.5226
alternative hypothesis: true difference in means between group Col-0 and
group nlp7-1 is not equal to 0
95 percent confidence interval:
 -0.1125659  0.2189909
sample estimates:
 mean in group Col-0 mean in group nlp7-1
         -0.03068906          -0.08390156

$logLRP

      Two Sample t-test
```

```
data:  x by dataset_nlp71_1mM$Genotype
t = 0.13624, df = 54, p-value = 0.8921
alternative hypothesis: true difference in means between group Col-0 and
group nlp7-1 is not equal to 0
95 percent confidence interval:
 -0.1750950  0.2006264
sample estimates:
 mean in group Col-0 mean in group nlp7-1
           -2.421692            -2.434457

$logPR

	Two Sample t-test

data:  x by dataset_nlp71_10mM$Genotype
t = -2.2297, df = 57, p-value = 0.02972
alternative hypothesis: true difference in means between group Col-0 and
group nlp7-1 is not equal to 0
95 percent confidence interval:
 -0.31753795 -0.01704951
sample estimates:
 mean in group Col-0 mean in group nlp7-1
            1.101995             1.269289

$logLR

	Two Sample t-test

data:  x by dataset_nlp71_10mM$Genotype
t = -0.2521, df = 57, p-value = 0.8019
alternative hypothesis: true difference in means between group Col-0 and
group nlp7-1 is not equal to 0
95 percent confidence interval:
 -0.2577887  0.2001386
sample estimates:
 mean in group Col-0 mean in group nlp7-1
            1.218915             1.247740

$logLRL

	Two Sample t-test

data:  x by dataset_nlp71_10mM$Genotype
t = -4.1233, df = 57, p-value = 0.0001226
alternative hypothesis: true difference in means between group Col-0 and
group nlp7-1 is not equal to 0
95 percent confidence interval:
 -0.6679732 -0.2312622
sample estimates:
 mean in group Col-0 mean in group nlp7-1
          -0.9663224           -0.5167047

$logALRL
```

```
      Two Sample t-test

data:  x by dataset_nlp71_10mM$Genotype
t = -2.5043, df = 57, p-value = 0.01515
alternative hypothesis: true difference in means between group Col-0 and
group nlp7-1 is not equal to 0
95 percent confidence interval:
 -0.75725774 -0.08432756
sample estimates:
 mean in group Col-0 mean in group nlp7-1
           -2.185237            -1.764445

$logTRL

      Two Sample t-test

data:  x by dataset_nlp71_10mM$Genotype
t = -0.87627, df = 57, p-value = 0.3846
alternative hypothesis: true difference in means between group Col-0 and
group nlp7-1 is not equal to 0
95 percent confidence interval:
 -0.17205802  0.06731077
sample estimates:
 mean in group Col-0 mean in group nlp7-1
            1.915368             1.967742

$logLRD

      Two Sample t-test

data:  x by dataset_nlp71_10mM$Genotype
t = 0.93213, df = 57, p-value = 0.3552
alternative hypothesis: true difference in means between group Col-0 and
group nlp7-1 is not equal to 0
95 percent confidence interval:
 -0.1589981  0.4359354
sample estimates:
 mean in group Col-0 mean in group nlp7-1
          0.11691964          -0.02154901

$logLRP

      Two Sample t-test

data:  x by dataset_nlp71_10mM$Genotype
t = -3.3399, df = 57, p-value = 0.001482
alternative hypothesis: true difference in means between group Col-0 and
group nlp7-1 is not equal to 0
95 percent confidence interval:
 -0.6354130 -0.1590752
sample estimates:
 mean in group Col-0 mean in group nlp7-1
           -2.881690            -2.484446
```

```
###################################################
#                                                 #
#Two-way ANOVA for Genotype:Treatment Interaction#
#                                                 #
###################################################
[1] "anac032-1"
logPR ~ Genotype * Treatment + Image
<environment: 0x0000029574b8e9c0>
Analysis of Variance Table

Response: logPR
                    Df Sum Sq  Mean Sq F value  Pr(>F)
Genotype             1 0.0388 0.038809  2.8675 0.09111 .
Treatment            1 0.0308 0.030751  2.2721 0.13246
Image               73 7.1824 0.098390  7.2699 < 2e-16 ***
Genotype:Treatment   1 0.0399 0.039897  2.9479 0.08671 .
Residuals          428 5.7925 0.013534
---
Signif. codes:  0 ,Äò***,Äô 0.001 ,Äò**,Äô 0.01 ,Äò*,Äô 0.05 ,Äò.,Äô 0.1
,Äò ,Äô 1
logLR ~ Genotype * Treatment + Image
<environment: 0x0000029574b8e9c0>
Analysis of Variance Table

Response: logLR
                    Df Sum Sq Mean Sq F value    Pr(>F)
Genotype             1  0.012  0.0119  0.0572    0.8110
Treatment            1  3.661  3.6608 17.5455 3.408e-05 ***
Image               73 71.656  0.9816  4.7045 < 2.2e-16 ***
Genotype:Treatment   1  0.170  0.1699  0.8144    0.3673
Residuals          428 89.301  0.2086
---
Signif. codes:  0 ,Äò***,Äô 0.001 ,Äò**,Äô 0.01 ,Äò*,Äô 0.05 ,Äò.,Äô 0.1
,Äò ,Äô 1
logLRL ~ Genotype * Treatment + Image
<environment: 0x0000029574b8e9c0>
Analysis of Variance Table

Response: logLRL
                    Df  Sum Sq Mean Sq  F value    Pr(>F)
Genotype             1   0.819   0.819   2.2659   0.13299
Treatment            1  42.956  42.956 118.7953 < 2.2e-16 ***
Image               73  92.439   1.266   3.5019  4.31e-16 ***
Genotype:Treatment   1   1.216   1.216   3.3634   0.06735 .
Residuals          428 154.764   0.362
---
Signif. codes:  0 ,Äò***,Äô 0.001 ,Äò**,Äô 0.01 ,Äò*,Äô 0.05 ,Äò.,Äô 0.1
,Äò ,Äô 1
logALRL ~ Genotype * Treatment + Image
<environment: 0x0000029574b8e9c0>
Analysis of Variance Table

Response: logALRL
                    Df  Sum Sq Mean Sq F value  Pr(>F)
```

```
Genotype             1   1.029  1.0291  3.9198 0.04836 *  
Treatment            1  21.537 21.5367 82.0305 < 2e-16 ***
Image               73 110.124  1.5086  5.7459 < 2e-16 ***
Genotype:Treatment   1   0.477  0.4769  1.8166 0.17843    
Residuals          428 112.369  0.2625                    
---
Signif. codes:  0 ‚Äò***‚Äô 0.001 ‚Äò**‚Äô 0.01 ‚Äò*‚Äô 0.05 ‚Äò.‚Äô 0.1
‚Äò ‚Äô 1
logTRL ~ Genotype * Treatment + Image
<environment: 0x0000029574b8e9c0>
Analysis of Variance Table

Response: logTRL
                    Df  Sum Sq Mean Sq F value    Pr(>F)    
Genotype             1  0.2690 0.26899  9.5246 0.0021591 ** 
Treatment            1  0.4148 0.41482 14.6882 0.0001459 ***
Image               73 12.8346 0.17582  6.2254 < 2.2e-16 ***
Genotype:Treatment   1  0.0733 0.07327  2.5944 0.1079808    
Residuals          428 12.0874 0.02824                      
---
Signif. codes:  0 ‚Äò***‚Äô 0.001 ‚Äò**‚Äô 0.01 ‚Äò*‚Äô 0.05 ‚Äò.‚Äô 0.1
‚Äò ‚Äô 1
logLRD ~ Genotype * Treatment + Image
<environment: 0x0000029574b8e9c0>
Analysis of Variance Table

Response: logLRD
                    Df  Sum Sq Mean Sq F value    Pr(>F)    
Genotype             1   4.420  4.4205 13.0162 0.0003453 ***
Treatment            1  30.433 30.4331 89.6114 < 2.2e-16 ***
Image               73 184.262  2.5241  7.4324 < 2.2e-16 ***
Genotype:Treatment   1   0.816  0.8155  2.4014 0.1219643    
Residuals          428 145.354  0.3396                      
---
Signif. codes:  0 ‚Äò***‚Äô 0.001 ‚Äò**‚Äô 0.01 ‚Äò*‚Äô 0.05 ‚Äò.‚Äô 0.1
‚Äò ‚Äô 1
logLRP ~ Genotype * Treatment + Image
<environment: 0x0000029574b8e9c0>
Analysis of Variance Table

Response: logLRP
                    Df  Sum Sq Mean Sq  F value  Pr(>F)    
Genotype             1   0.149   0.149   0.6068 0.43641    
Treatment            1  34.928  34.928 141.8710 < 2e-16 ***
Image               73  86.914   1.191   4.8360 < 2e-16 ***
Genotype:Treatment   1   0.692   0.692   2.8126 0.09426 .  
Residuals          428 105.373   0.246                     
-
---
Signif. codes:  0 ‚Äò***‚Äô 0.001 ‚Äò**‚Äô 0.01 ‚Äò*‚Äô 0.05 ‚Äò.‚Äô 0.1
‚Äò ‚Äô 1
[1] "arf18-cc-1"
logPR ~ Genotype * Treatment + Image
<environment: 0x00000295738c0158>
Analysis of Variance Table

Response: logPR
                    Df Sum Sq  Mean Sq F value  Pr(>F)
```

```
Genotype             1 0.0142 0.014198  1.0996 0.29495
Treatment            1 0.0379 0.037909  2.9359 0.08736 .
Image               74 7.2179 0.097539  7.5540 < 2e-16 ***
Genotype:Treatment   1 0.0150 0.014971  1.1594 0.28220
Residuals          421 5.4360 0.012912
---
Signif. codes:  0 ,Äò***,Äô 0.001 ,Äò**,Äô 0.01 ,Äò*,Äô 0.05 ,Äò.,Äô 0.1
,Äò ,Äô 1
logLR ~ Genotype * Treatment + Image
<environment: 0x00000295738c0158>
Analysis of Variance Table

Response: logLR
                    Df Sum Sq Mean Sq F value    Pr(>F)
Genotype             1  0.246  0.2463  1.1921    0.2755
Treatment            1  3.589  3.5891 17.3719 3.732e-05 ***
Image               74 71.404  0.9649  4.6703 < 2.2e-16 ***
Genotype:Treatment   1  0.106  0.1063  0.5143    0.4737
Residuals          421 86.981  0.2066
---
Signif. codes:  0 ,Äò***,Äô 0.001 ,Äò**,Äô 0.01 ,Äò*,Äô 0.05 ,Äò.,Äô 0.1
,Äò ,Äô 1
logLRL ~ Genotype * Treatment + Image
<environment: 0x00000295738c0158>
Analysis of Variance Table

Response: logLRL
                    Df  Sum Sq Mean Sq  F value    Pr(>F)
Genotype             1   4.314   4.314  12.1396 0.0005454 ***
Treatment            1  38.124  38.124 107.2689 < 2.2e-16 ***
Image               74  90.743   1.226   3.4503 8.753e-16 ***
Genotype:Treatment   1   0.435   0.435   1.2226 0.2694803
Residuals          421 149.626   0.355
---
Signif. codes:  0 ,Äò***,Äô 0.001 ,Äò**,Äô 0.01 ,Äò*,Äô 0.05 ,Äò.,Äô 0.1
,Äò ,Äô 1
logALRL ~ Genotype * Treatment + Image
<environment: 0x00000295738c0158>
Analysis of Variance Table

Response: logALRL
                    Df  Sum Sq Mean Sq F value    Pr(>F)
Genotype             1   2.499  2.4991  9.5707  0.002109 **
Treatment            1  18.318 18.3180 70.1518 8.254e-16 ***
Image               74 108.061  1.4603  5.5924 < 2.2e-16 ***
Genotype:Treatment   1   0.111  0.1110  0.4252  0.514692
Residuals          421 109.932  0.2611
---
Signif. codes:  0 ,Äò***,Äô 0.001 ,Äò**,Äô 0.01 ,Äò*,Äô 0.05 ,Äò.,Äô 0.1
,Äò ,Äô 1
logTRL ~ Genotype * Treatment + Image
<environment: 0x00000295738c0158>
Analysis of Variance Table

Response: logTRL
                    Df  Sum Sq Mean Sq F value    Pr(>F)
Genotype             1  0.2533 0.25328  9.2080 0.0025589 **
Treatment            1  0.3355 0.33551 12.1976 0.0005292 ***
```

```
Image                74 12.8492 0.17364  6.3126 < 2.2e-16 ***
Genotype:Treatment    1  0.0002 0.00018  0.0067 0.9350032
Residuals           421 11.5803 0.02751
---
Signif. codes:  0 ‚Äò***‚Äô 0.001 ‚Äò**‚Äô 0.01 ‚Äò*‚Äô 0.05 ‚Äò.‚Äô 0.1
‚Äò ‚Äô 1
logLRD ~ Genotype * Treatment + Image
<environment: 0x00000295738c0158>
Analysis of Variance Table

Response: logLRD
                     Df  Sum Sq Mean Sq F value    Pr(>F)
Genotype              1  12.007 12.0066 35.7672 4.752e-09 ***
Treatment             1  26.131 26.1310 77.8429 < 2.2e-16 ***
Image                74 180.758  2.4427  7.2766 < 2.2e-16 ***
Genotype:Treatment    1   0.611  0.6108  1.8195    0.1781
Residuals           421 141.325  0.3357
---
Signif. codes:  0 ‚Äò***‚Äô 0.001 ‚Äò**‚Äô 0.01 ‚Äò*‚Äô 0.05 ‚Äò.‚Äô 0.1
‚Äò ‚Äô 1
logLRP ~ Genotype * Treatment + Image
<environment: 0x00000295738c0158>
Analysis of Variance Table

Response: logLRP
                     Df  Sum Sq Mean Sq  F value    Pr(>F)
Genotype              1   2.477  2.4770  10.1759  0.001529 **
Treatment             1  31.306 31.3065 128.6103 < 2.2e-16 ***
Image                74  85.442  1.1546   4.7433 < 2.2e-16 ***
Genotype:Treatment    1   0.453  0.4525   1.8591  0.173456
Residuals           421 102.480  0.2434

---
Signif. codes:  0 ‚Äò***‚Äô 0.001 ‚Äò**‚Äô 0.01 ‚Äò*‚Äô 0.05 ‚Äò.‚Äô 0.1
‚Äò ‚Äô 1
[1] "nlp7-1"
logPR ~ Genotype * Treatment + Image
<environment: 0x00000295720d08c8>
Analysis of Variance Table

Response: logPR
                     Df Sum Sq  Mean Sq F value    Pr(>F)
Genotype              1 0.0078 0.007775  0.5453  0.460623
Treatment             1 0.1341 0.134104  9.4052  0.002293 **
Image                73 7.5808 0.103846  7.2831 < 2.2e-16 ***
Genotype:Treatment    1 0.1943 0.194302 13.6271  0.000250 ***
Residuals           454 6.4733 0.014258
---
Signif. codes:  0 ‚Äò***‚Äô 0.001 ‚Äò**‚Äô 0.01 ‚Äò*‚Äô 0.05 ‚Äò.‚Äô 0.1
‚Äò ‚Äô 1
logLR ~ Genotype * Treatment + Image
<environment: 0x00000295720d08c8>
Analysis of Variance Table

Response: logLR
                     Df Sum Sq Mean Sq F value    Pr(>F)
Genotype              1 10.469 10.4688 52.6463 1.740e-12 ***
Treatment             1  3.081  3.0810 15.4940 9.575e-05 ***
```

```
Image               73 69.890  0.9574  4.8147 < 2.2e-16 ***
Genotype:Treatment   1  0.101  0.1008  0.5068    0.4769
Residuals          454 90.278  0.1989
---
Signif. codes:  0 ‚Äò***‚Äô 0.001 ‚Äò**‚Äô 0.01 ‚Äò*‚Äô 0.05 ‚Äò.‚Äô 0.1
‚Äò ‚Äô 1
logLRL ~ Genotype * Treatment + Image
<environment: 0x00000295720d08c8>
Analysis of Variance Table

Response: logLRL
                    Df  Sum Sq Mean Sq  F value    Pr(>F)
Genotype             1   4.049   4.049  12.7361 0.0003969 ***
Treatment            1  32.295  32.295 101.5747 < 2.2e-16 ***
Image               73  87.957   1.205   3.7896 < 2.2e-16 ***
Genotype:Treatment   1   0.189   0.189   0.5939 0.4413283
Residuals          454 144.346   0.318
---
Signif. codes:  0 ‚Äò***‚Äô 0.001 ‚Äò**‚Äô 0.01 ‚Äò*‚Äô 0.05 ‚Äò.‚Äô 0.1
‚Äò ‚Äô 1
logALRL ~ Genotype * Treatment + Image
<environment: 0x00000295720d08c8>
Analysis of Variance Table

Response: logALRL
                    Df  Sum Sq Mean Sq  F value    Pr(>F)
Genotype             1  27.540 27.5399 105.8896 < 2.2e-16 ***
Treatment            1  15.426 15.4260  59.3121 8.509e-14 ***
Image               73 105.879  1.4504   5.5767 < 2.2e-16 ***
Genotype:Treatment   1   0.014  0.0137   0.0527    0.8185
Residuals          454 118.077  0.2601
---
Signif. codes:  0 ‚Äò***‚Äô 0.001 ‚Äò**‚Äô 0.01 ‚Äò*‚Äô 0.05 ‚Äò.‚Äô 0.1
‚Äò ‚Äô 1
logTRL ~ Genotype * Treatment + Image
<environment: 0x00000295720d08c8>
Analysis of Variance Table

Response: logTRL
                    Df  Sum Sq Mean Sq  F value    Pr(>F)
Genotype             1  8.3718  8.3718 259.2125 < 2.2e-16 ***
Treatment            1  0.4183  0.4183  12.9523 0.0003547 ***
Image               73 11.0748  0.1517   4.6974 < 2.2e-16 ***
Genotype:Treatment   1  0.4751  0.4751  14.7117 0.0001430 ***
Residuals          454 14.6628  0.0323
---
Signif. codes:  0 ‚Äò***‚Äô 0.001 ‚Äò**‚Äô 0.01 ‚Äò*‚Äô 0.05 ‚Äò.‚Äô 0.1
‚Äò ‚Äô 1
logLRD ~ Genotype * Treatment + Image
<environment: 0x00000295720d08c8>
Analysis of Variance Table

Response: logLRD
                    Df  Sum Sq Mean Sq  F value    Pr(>F)
Genotype             1  73.652  73.652 242.2278 < 2.2e-16 ***
Treatment            1  19.152  19.152  62.9864 1.643e-14 ***
Image               73 179.294   2.456   8.0776 < 2.2e-16 ***
Genotype:Treatment   1   0.015   0.015   0.0500    0.8231
```

```
Residuals          454 138.043   0.304
---
Signif. codes:  0 ‚Äò***‚Äô 0.001 ‚Äò**‚Äô 0.01 ‚Äò*‚Äô 0.05 ‚Äò.‚Äô 0.1
‚Äò ‚Äô 1
logLRP ~ Genotype * Treatment + Image
<environment: 0x00000295720d08c8>
Analysis of Variance Table

Response: logLRP
                    Df  Sum Sq Mean Sq  F value  Pr(>F)
Genotype             1  24.066 24.0659 106.8679 < 2e-16 ***
Treatment            1  25.362 25.3622 112.6245 < 2e-16 ***
Image               73  82.584  1.1313   5.0236 < 2e-16 ***
Genotype:Treatment   1   1.263  1.2630   5.6086 0.01829 *
Residuals          454 102.237  0.2252
---
Signif. codes:  0 ‚Äò***‚Äô 0.001 ‚Äò**‚Äô 0.01 ‚Äò*‚Äô 0.05 ‚Äò.‚Äô 0.1
‚Äò ‚Äô 1
[1] "anac032-1/arf18-2"
logPR ~ Genotype * Treatment + Image
<environment: 0x0000029574f21aa8>
Analysis of Variance Table

Response: logPR
                    Df Sum Sq  Mean Sq F value    Pr(>F)
Genotype             1 0.1567 0.156685 10.3209  0.001409 **
Treatment            1 0.0037 0.003694  0.2433  0.622066
Image               74 7.4037 0.100050  6.5903 < 2.2e-16 ***
Genotype:Treatment   1 0.0099 0.009891  0.6515  0.419986
Residuals          454 6.8923 0.015181
---
Signif. codes:  0 ‚Äò***‚Äô 0.001 ‚Äò**‚Äô 0.01 ‚Äò*‚Äô 0.05 ‚Äò.‚Äô 0.1
‚Äò ‚Äô 1
logLR ~ Genotype * Treatment + Image
<environment: 0x0000029574f21aa8>
Analysis of Variance Table

Response: logLR
                    Df Sum Sq Mean Sq F value    Pr(>F)
Genotype             1  6.853  6.8531 31.8630 2.919e-08 ***
Treatment            1  3.478  3.4781 16.1711 6.776e-05 ***
Image               74 70.577  0.9537  4.4344 < 2.2e-16 ***
Genotype:Treatment   1  0.219  0.2191  1.0186    0.3134
Residuals          454 97.646  0.2151
---
Signif. codes:  0 ‚Äò***‚Äô 0.001 ‚Äò**‚Äô 0.01 ‚Äò*‚Äô 0.05 ‚Äò.‚Äô 0.1
‚Äò ‚Äô 1
logLRL ~ Genotype * Treatment + Image
<environment: 0x0000029574f21aa8>
Analysis of Variance Table

Response: logLRL
                    Df  Sum Sq Mean Sq  F value    Pr(>F)
Genotype             1   1.075   1.075   2.8750   0.09065 .
Treatment            1  40.376  40.376 107.9416 < 2.2e-16 ***
Image               74  84.382   1.140   3.0485 4.082e-13 ***
Genotype:Treatment   1   0.127   0.127   0.3390   0.56071
Residuals          454 169.821   0.374
```

```
---
Signif. codes:  0 ‚Äò***‚Äô 0.001 ‚Äò**‚Äô 0.01 ‚Äò*‚Äô 0.05 ‚Äò.‚Äô 0.1
‚Äò ‚Äô 1
logALRL ~ Genotype * Treatment + Image
<environment: 0x0000029574f21aa8>
Analysis of Variance Table

Response: logALRL
                    Df  Sum Sq Mean Sq F value    Pr(>F)    
Genotype             1   2.499  2.4990  9.0190  0.002819 ** 
Treatment            1  20.153 20.1534 72.7343 2.227e-16 ***
Image               74 107.295  1.4499  5.2328 < 2.2e-16 ***
Genotype:Treatment   1   0.679  0.6792  2.4513  0.118123    
Residuals          454 125.796  0.2771                      
---
Signif. codes:  0 ‚Äò***‚Äô 0.001 ‚Äò**‚Äô 0.01 ‚Äò*‚Äô 0.05 ‚Äò.‚Äô 0.1
‚Äò ‚Äô 1
logTRL ~ Genotype * Treatment + Image
<environment: 0x0000029574f21aa8>
Analysis of Variance Table

Response: logTRL
                    Df  Sum Sq Mean Sq F value    Pr(>F)    
Genotype             1  0.7958 0.79580 27.1752 2.828e-07 ***
Treatment            1  0.2956 0.29560 10.0943  0.001589 ** 
Image               74 12.6999 0.17162  5.8606 < 2.2e-16 ***
Genotype:Treatment   1  0.0008 0.00077  0.0261  0.871630    
Residuals          454 13.2949 0.02928                      
---
Signif. codes:  0 ‚Äò***‚Äô 0.001 ‚Äò**‚Äô 0.01 ‚Äò*‚Äô 0.05 ‚Äò.‚Äô 0.1
‚Äò ‚Äô 1
logLRD ~ Genotype * Treatment + Image
<environment: 0x0000029574f21aa8>
Analysis of Variance Table

Response: logLRD
                    Df  Sum Sq Mean Sq F value    Pr(>F)    
Genotype             1   6.115  6.1155 17.2887  3.84e-05 ***
Treatment            1  29.730 29.7304 84.0496 < 2.2e-16 ***
Image               74 175.757  2.3751  6.7145 < 2.2e-16 ***
Genotype:Treatment   1   0.208  0.2075  0.5867    0.4441    
Residuals          454 160.591  0.3537                      
---
Signif. codes:  0 ‚Äò***‚Äô 0.001 ‚Äò**‚Äô 0.01 ‚Äò*‚Äô 0.05 ‚Äò.‚Äô 0.1
‚Äò ‚Äô 1
logLRP ~ Genotype * Treatment + Image
<environment: 0x0000029574f21aa8>
Analysis of Variance Table

Response: logLRP
                    Df  Sum Sq Mean Sq  F value Pr(>F)    
Genotype             1   0.021   0.021   0.0821 0.7746    
Treatment            1  33.762  33.762 131.9704 <2e-16 ***
Image               74  81.157   1.097   4.2869 <2e-16 ***
Genotype:Treatment   1   0.108   0.108   0.4216 0.5165    
Residuals          454 116.148   0.256                    
---
```

```
#########################
#                       #
#One-way ANOVA for Col-0#
#                       #
#########################
logPR ~ Treatment
<environment: 0x000001f047039840>
Analysis of Variance Table

Response: logPR
           Df Sum Sq  Mean Sq F value Pr(>F)
Treatment   1  0.023 0.022956  0.8627 0.3535
Residuals 470 12.506 0.026609
logLR ~ Treatment
<environment: 0x000001f047039840>
Analysis of Variance Table

Response: logLR
           Df  Sum Sq Mean Sq F value   Pr(>F)
Treatment   1   2.926 2.92600  8.9326 0.002948 **
Residuals 470 153.956 0.32757
---
Signif. codes:  0 ‚Äò***‚Äô 0.001 ‚Äò**‚Äô 0.01 ‚Äò*‚Äô 0.05 ‚Äò.‚Äô 0.1
‚Äò ‚Äô 1
logLRL ~ Treatment
<environment: 0x000001f047039840>
Analysis of Variance Table

Response: logLRL
           Df  Sum Sq Mean Sq F value    Pr(>F)
Treatment   1  33.318  33.318  69.275 9.393e-16 ***
Residuals 470 226.049   0.481
---
Signif. codes:  0 ‚Äò***‚Äô 0.001 ‚Äò**‚Äô 0.01 ‚Äò*‚Äô 0.05 ‚Äò.‚Äô 0.1
‚Äò ‚Äô 1
logALRL ~ Treatment
<environment: 0x000001f047039840>
Analysis of Variance Table

Response: logALRL
           Df  Sum Sq Mean Sq F value    Pr(>F)
Treatment   1  16.497 16.4969  36.641 2.907e-09 ***
Residuals 470 211.612  0.4502
---
Signif. codes:  0 ‚Äò***‚Äô 0.001 ‚Äò**‚Äô 0.01 ‚Äò*‚Äô 0.05 ‚Äò.‚Äô 0.1
‚Äò ‚Äô 1
logTRL ~ Treatment
<environment: 0x000001f047039840>
Analysis of Variance Table

Response: logTRL
           Df  Sum Sq  Mean Sq F value  Pr(>F)
Treatment   1  0.2463 0.246267  4.8114 0.02876 *
Residuals 470 24.0567 0.051184
---
Signif. codes:  0 ‚Äò***‚Äô 0.001 ‚Äò**‚Äô 0.01 ‚Äò*‚Äô 0.05 ‚Äò.‚Äô 0.1
‚Äò ‚Äô 1
logLRD ~ Treatment
```

```
<environment: 0x000001f047039840>
Analysis of Variance Table

Response: logLRD
           Df  Sum Sq Mean Sq F value    Pr(>F)    
Treatment   1  22.363  22.363  34.192 9.355e-09 ***
Residuals 470 307.392   0.654                      
---
Signif. codes:  0 ‚Äò***‚Äô 0.001 ‚Äò**‚Äô 0.01 ‚Äò*‚Äô 0.05 ‚Äò.‚Äô 0.1
‚Äò ‚Äô 1
logLRP ~ Treatment
<environment: 0x000001f047039840>
Analysis of Variance Table

Response: logLRP
           Df  Sum Sq Mean Sq F value    Pr(>F)    
Treatment   1  27.836  27.836  74.036 < 2.2e-16 ***
Residuals 470 176.707   0.376                      
---
Signif. codes:  0 ‚Äò***‚Äô 0.001 ‚Äò**‚Äô 0.01 ‚Äò*‚Äô 0.05 ‚Äò.‚Äô 0.1
‚Äò ‚Äô 1

****************************************************************

Two-Way ANOVA
---
Signif. codes:  0 ‚Äò***‚Äô 0.001 ‚Äò**‚Äô 0.01 ‚Äò*‚Äô 0.05 ‚Äò.‚Äô 0.1
‚Äò ‚Äô 1
Signif. codes:  0 ‚Äò***‚Äô 0.001 ‚Äò**‚Äô 0.01 ‚Äò*‚Äô 0.05 ‚Äò.‚Äô 0.1
‚Äò ‚Äô 1
[1] "anac032-cc-s-1"
logPR ~ Genotype * Treatment + Image
<environment: 0x000002957087c9e0>
Analysis of Variance Table

Response: logPR
                    Df Sum Sq  Mean Sq F value  Pr(>F)    
Genotype             1 0.0020 0.002002  0.1559 0.69316    
Treatment            1 0.0160 0.016018  1.2470 0.26474    
Image               73 7.1620 0.098109  7.6379 < 2e-16 ***
Genotype:Treatment   1 0.0380 0.038047  2.9620 0.08596 .  
Residuals          434 5.5747 0.012845                    
---
Signif. codes:  0 ‚Äò***‚Äô 0.001 ‚Äò**‚Äô 0.01 ‚Äò*‚Äô 0.05 ‚Äò.‚Äô 0.1
‚Äò ‚Äô 1
logLR ~ Genotype * Treatment + Image
<environment: 0x000002957087c9e0>
Analysis of Variance Table

Response: logLR
                    Df Sum Sq Mean Sq F value    Pr(>F)    
Genotype             1  0.969 0.96853  4.4933 0.0345946 *  
Treatment            1  3.040 3.03960 14.1016 0.0001968 ***
Image               73 70.113 0.96045  4.4558 < 2.2e-16 ***
Genotype:Treatment   1  0.131 0.13128  0.6090 0.4355770    
Residuals          434 93.549 0.21555                      
---
```

```
Signif. codes:  0 ‚Äò***‚Äô 0.001 ‚Äò**‚Äô 0.01 ‚Äò*‚Äô 0.05 ‚Äò.‚Äô 0.1
‚Äò ‚Äô 1
logLRL ~ Genotype * Treatment + Image
<environment: 0x000002957087c9e0>
Analysis of Variance Table

Response: logLRL
                    Df  Sum Sq Mean Sq F value    Pr(>F)    
Genotype             1   9.565   9.565 25.7929 5.654e-07 ***
Treatment            1  37.008  37.008 99.7986 < 2.2e-16 ***
Image               73  85.360   1.169  3.1533 1.262e-13 ***
Genotype:Treatment   1   0.070   0.070  0.1893    0.6637    
Residuals          434 160.939   0.371                      
---
Signif. codes:  0 ‚Äò***‚Äô 0.001 ‚Äò**‚Äô 0.01 ‚Äò*‚Äô 0.05 ‚Äò.‚Äô 0.1
‚Äò ‚Äô 1
logALRL ~ Genotype * Treatment + Image
<environment: 0x000002957087c9e0>
Analysis of Variance Table

Response: logALRL
                    Df  Sum Sq Mean Sq F value    Pr(>F)    
Genotype             1   4.446  4.4460 16.7093 5.192e-05 ***
Treatment            1  18.835 18.8355 70.7895 5.807e-16 ***
Image               73 107.195  1.4684  5.5188 < 2.2e-16 ***
Genotype:Treatment   1   0.009  0.0095  0.0356    0.8504    
Residuals          434 115.478  0.2661                      
---
Signif. codes:  0 ‚Äò***‚Äô 0.001 ‚Äò**‚Äô 0.01 ‚Äò*‚Äô 0.05 ‚Äò.‚Äô 0.1
‚Äò ‚Äô 1
logTRL ~ Genotype * Treatment + Image
<environment: 0x000002957087c9e0>
Analysis of Variance Table

Response: logTRL
                    Df  Sum Sq Mean Sq F value    Pr(>F)    
Genotype             1  0.7502 0.75021 26.5609 3.886e-07 ***
Treatment            1  0.2697 0.26967  9.5476  0.002131 ** 
Image               73 12.5735 0.17224  6.0981 < 2.2e-16 ***
Genotype:Treatment   1  0.0002 0.00017  0.0062  0.937432    
Residuals          434 12.2583 0.02824                      
---
Signif. codes:  0 ‚Äò***‚Äô 0.001 ‚Äò**‚Äô 0.01 ‚Äò*‚Äô 0.05 ‚Äò.‚Äô 0.1
‚Äò ‚Äô 1
logLRD ~ Genotype * Treatment + Image
<environment: 0x000002957087c9e0>
Analysis of Variance Table

Response: logLRD
                    Df  Sum Sq Mean Sq F value    Pr(>F)    
Genotype             1  20.746 20.7462 60.1677 6.278e-14 ***
Treatment            1  26.011 26.0108 75.4360 < 2.2e-16 ***
Image               73 176.304  2.4151  7.0043 < 2.2e-16 ***
Genotype:Treatment   1   0.212  0.2116  0.6138    0.4338    
Residuals          434 149.646  0.3448                      
---
Signif. codes:  0 ‚Äò***‚Äô 0.001 ‚Äò**‚Äô 0.01 ‚Äò*‚Äô 0.05 ‚Äò.‚Äô 0.1
‚Äò ‚Äô 1
```

```
logLRP ~ Genotype * Treatment + Image
<environment: 0x000002957087c9e0>
Analysis of Variance Table

Response: logLRP
                    Df  Sum Sq Mean Sq  F value    Pr(>F)    
Genotype             1   4.957  4.9575  19.8339 1.076e-05 ***
Treatment            1  30.960 30.9596 123.8630 < 2.2e-16 ***
Image               73  82.200  1.1260   4.5050 < 2.2e-16 ***
Genotype:Treatment   1   0.077  0.0774   0.3096    0.5782    
Residuals          434 108.478  0.2500                       

*******************************************
```

Tomato Data

LogPR
0mM

Arf18-9b-1
The t-value is 0.61231. The p-value is .546092. The result is not significant at $p < .05$.

Arf18-9b-2
The t-value is -0.41715. The p-value is .679273. The result is not significant at $p < .05$.

Nlp7-1
The t-value is 2.00315. The p-value is .058901. The result is not significant at $p < .05$.

Nlp7-2
The t-value is 1.64683. The p-value is .115217. The result is not significant at $p < .05$.

1mM Arf18-9b-1
The t-value is -0.57884. The p-value is .567327. The result is not significant at $p < .05$.

Nlp7-1
The t-value is 0.06521. The p-value is .948402. The result is not significant at $p < .05$.

Nlp7-2
The t-value is -0.3964. The p-value is .69422. The result is not significant at $p < .05$.

10mM Arf18-9b-1
The t-value is -0.21882. The p-value is .828809. The result is not significant at $p < .05$.

Arf18-9b-2
The t-value is -0.20423. The p-value is .839707. The result is not significant at $p < .05$.

Nlp7-1
The t-value is 0.39643. The p-value is .694908. The result is not significant at $p < .05$.

Nlp7-2
The t-value is 0.56488. The p-value is .576497. The result is not significant at p < .05.
LogLR 0mM Arf18-9b-1
The t-value is 0.85284. The p-value is .401844. The result is not significant at p < .05.

Arf18-9b-2
The t-value is 0.1612. The p-value is .87289. The result is not significant at p < .05.

Nlp7-1
The t-value is -0.78371. The p-value is .439564. The result is not significant at p < .05.

Nlp7-2
The t-value is 0.02031. The p-value is .983922. The result is not significant at p < .05.

1mM Arf18-9b-1
The t-value is 0.01174. The p-value is .990721. The result is not significant at p < .05. Arf18-9b-2
The t-value is 0.29096. The p-value is .773084. The result is not significant at p < .05. Nlp7-1
The t-value is 0.41708. The p-value is .679493. The result is not significant at p < .05. Nlp7-2
The t-value is 0.28014. The p-value is .781175. The result is not significant at p < .05.

10mM Arf18-9b-1
The t-value is -2.52796. The p-value is .020001. The result is significant at p < .05. Arf18-9b-2
The t-value is -0.707. The p-value is .486105. The result is not significant at p < .05. Nlp7-1
The t-value is -0.7709. The p-value is .447989. The result is not significant at p < .05. Nlp7-2
The t-value is 0.54191. The p-value is .592322. The result is not significant at p < .05.
LogLRL 0mM Arf18-9b-1
The t-value is 0.96786. The p-value is .342767. The result is not significant at p < .05. Arf18-9b-2
The t-value is -1.00858. The p-value is .320517. The result is not significant at p < .05. Nlp7-1
The t-value is -0.53523. The p-value is .596715. The result is not significant at p < .05. Nlp7-2
The t-value is -1.16495. The p-value is .252649. The result is not significant at p < .05.

1mM Arf18-9b-1
The t-value is 2.93724. The p-value is .006698. The result is significant at p < .05. Arf18-9b-2
The t-value is 1.18104. The p-value is .246866. The result is not significant at p < .05 Nlp7-1
The t-value is 1.30648. The p-value is .200999. The result is not significant at p < .05. Nlp7-2
The t-value is 1.96043. The p-value is .0587. The result is not significant at p < .05.

10mM Arf18-9b-1
The t-value is -0.02898. The p-value is .977154. The result is not significant at p < .05. Arf18-9b-2
The t-value is -0.18655. The p-value is .853464. The result is not significant at p < .05. Nlp7-1
The t-value is 1.30355. The p-value is .203812. The result is not significant at p < .05 Nlp7-2
The t-value is 0.87754. The p-value is .387663. The result is not significant at p < .05.
LogALRL
0mM Arf18-9b-1
The t-value is -0.1351. The p-value is .893662. The result is not significant at p < .05. Arf18-9b-2
The t-value is -1.30956. The p-value is .19939. The result is not significant at p < .05. Nlp7-1
The t-value is 0.25596. The p-value is .799852. The result is not significant at p < .05. Nlp7-2
The t-value is -1.44371. The p-value is .158539. The result is not significant at p < .05.

1mM Arf18-9b-1
The t-value is 3.97043. The p-value is .000479. The result is significant at p < .05. Arf18-9b-2
The t-value is 1.03755. The p-value is .307771. The result is not significant at p < .05. Nlp7-1
The t-value is 1.05941. The p-value is .297599. The result is not significant at p < .05. Nlp7-2
The t-value is 2.26187. The p-value is .030634. The result is significant at p < .05.

10mM Arf18-9b-1
The t-value is 3.74574. The p-value is .001191. The result is significant at p < .05. Arf18-9b-2
The t-value is 0.56733. The p-value is .575356. The result is not significant at p < .05. Nlp7-1
The t-value is 2.80398. The p-value is .00942. The result is significant at p < .05. Nlp7-2
The t-value is 0.49664. The p-value is .623323. The result is not significant at p < .05.
LogTRL 0mM Arf18-9b-1
The t-value is 0.98852. The p-value is .332759. The result is not significant at p < .05. Arf18-9b-2
The t-value is -1.2308. The p-value is .227099. The result is not significant at p < .05. Nlp7-1
The t-value is -2.11631. The p-value is .043022. The result is significant at p < .05. Nlp7-2
The t-value is -1.54. The p-value is .133097. The result is not significant at p < .05.

1mM Arf18-9b-1
The t-value is 2.20049. The p-value is .036187. The result is significant at p < .05. Arf18-9b-2
The t-value is -1.23322. The p-value is .225952. The result is not significant at p < .05. Nlp7-1
The t-value is 0.47944. The p-value is .634788. The result is not significant at p < .05. Nlp7-2
The t-value is 0.46245. The p-value is .646621. The result is not significant at p < .05.

10mM Arf18-9b-1
The t-value is 0.2305. The p-value is .819836. The result is not significant at p < .05. Arf18-9b-2
The t-value is 0.28743. The p-value is .77598. The result is not significant at p < .05. Nlp7-1
The t-value is 1.11468. The p-value is .274812. The result is not significant at p < .05. Nlp7-2
The t-value is 0.95943. The p-value is .345273. The result is not significant at p < .05.
LogLRD 0mM Arf18-9b-1
The t-value is 0.38967. The p-value is .700219. The result is not significant at p < .05. Arf18-9b-2
The t-value is 0.17567. The p-value is .861626. The result is not significant at p < .05. Nlp7-1
The t-value is 0.69905. The p-value is .490289. The result is not significant at p < .05. Nlp7-2
The t-value is 0.21295. The p-value is .832714. The result is not significant at p < .05.

1mM Arf18-9b-1
The t-value is 1.78373. The p-value is .086145. The result is not significant at p < .05. Arf18-9b-2
The t-value is 2.93212. The p-value is .006509. The result is significant at p < .05. Nlp7-1
The t-value is 2.12874. The p-value is .041601. The result is significant at p < .05 Nlp7-2
The t-value is 2.23989. The p-value is .032409. The result is significant at p < .05.

10mM Arf18-9b-1
The t-value is -1.50694. The p-value is .146722. The result is not significant at p < .05. Arf18-9b-2
The t-value is 0.01089. The p-value is .991397. The result is not significant at p < .05. Nlp7-1
The t-value is -0.72229. The p-value is .476561. The result is not significant at p < .05. Nlp7-2
The t-value is 0.19374. The p-value is .847777. The result is not significant at p < .05.
LogLRP
0mM Arf18-9b-1
The t-value is 0.65378. The p-value is .519469. The result is not significant at p < .05. Arf18-9b-2
The t-value is -0.50565. The p-value is .616462. The result is not significant at p < .05. Nlp7-1
The t-value is 0.74586. The p-value is .461969. The result is not significant at p < .05. Nlp7-2
The t-value is -0.77361. The p-value is .444841. The result is not significant at p < .05.

1mM Arf18-9b-1
The t-value is 3.20669. The p-value is .003441. The result is significant at p < .05. Arf18-9b-2
The t-value is 3.38466. The p-value is .002003. The result is significant at p < .05. Nlp7-1
The t-value is 1.69731. The p-value is .099657. The result is not significant at p < .05. Nlp7-2

The t-value is 2.18101. The p-value is .036645. The result is significant at p < .05.

10mM Arf18-9b-1
The t-value is 0.48567. The p-value is .632229. The result is not significant at p < .05. Arf18-9b-2
The t-value is 0.36774. The p-value is .716043. The result is not significant at p < .05. Nlp7-1
The t-value is 1.06344. The p-value is .297361. The result is not significant at p < .05. Nlp7-2
The t-value is 0.89422. The p-value is .378827. The result is not significant at p < .05.

```
mod1 <- lm(LogPR~Genotype * Condition, contrasts=list(Genotype=contr.sum,
Condition=contr.sum), data=x)
Anova(mod1, type=3) Anova Table (Type III tests)

Response: LogPR
Sum Sq Df F value Pr(>F) (Intercept)    108.955 1 3357.6824 < 2e-16 ***
Genotype    0.147  4    1.1290 0.34465
Condition   0.072  2    1.1129 0.33105
Genotype:Condition  0.450  8 1.7325 0.09435 .
Residuals   5.354 165
---
Signif. codes: 0 ‚Äò***‚Äô 0.001 ‚Äò**‚Äô 0.01 ‚Äò*‚Äô 0.05 ‚Äò.‚Äô 0.1
‚Äò ‚Äô 1
mod1 <- lm(LogLR~Genotype * Condition, contrasts=list(Genotype=contr.sum,
Condition=contr.sum), data=x)
Anova(mod1, type=3) Anova Table (Type III tests)

Response: LogLR
Sum Sq Df F value Pr(>F) (Intercept)    118.708 1 2184.3471 < 2e-16 ***
Genotype    0.131  4    0.6030 0.66103
Condition   0.390  2    3.5859 0.03003 *
Genotype:Condition  0.399  8 0.9170 0.50411
Residuals   8.423 155
---
Signif. codes: 0 ‚Äò***‚Äô 0.001 ‚Äò**‚Äô 0.01 ‚Äò*‚Äô 0.05 ‚Äò.‚Äô 0.1
‚Äò ‚Äô 1
mod1 <- lm(LogLRL~Genotype * Condition,
contrasts=list(Genotype=contr.sum, Condition=contr.sum), data=x)
Anova(mod1, type=3) Anova Table (Type III tests)

Response: LogLRL
Sum Sq Df F value Pr(>F) (Intercept)    167.019 1 3717.0283 < 2e-16 ***
Genotype    0.236  4    1.3138 0.26725
Condition   0.531  2    5.9041 0.00338 **
Genotype:Condition  0.483  8 1.3437 0.22588
Residuals   6.965 155
---
Signif. codes: 0 ‚Äò***‚Äô 0.001 ‚Äò**‚Äô 0.01 ‚Äò*‚Äô 0.05 ‚Äò.‚Äô 0.1
‚Äò ‚Äô 1
mod1 <- lm(LogALRL~Genotype * Condition,
contrasts=list(Genotype=contr.sum, Condition=contr.sum), data=x)
Anova(mod1, type=3) Anova Table (Type III tests)

Response: LogALRL
Sum Sq Df F value Pr(>F) (Intercept)    5.7633  1 146.0570 < 2e-16 ***
Genotype    0.4842  4  3.0679 0.01822 *
```

```
Condition   0.1309  2  1.6582 0.19385
Genotype:Condition 0.4804  8  1.5217 0.15376
Residuals   6.1162 155
---
Signif. codes: 0 ‚Äò***‚Äô 0.001 ‚Äò**‚Äô 0.01 ‚Äò*‚Äô 0.05 ‚Äò.‚Äô 0.1
‚Äò ‚Äô 1
mod1 <- lm(LogTRL~Genotype * Condition,
contrasts=list(Genotype=contr.sum, Condition=contr.sum), data=x)
Anova(mod1, type=3) Anova Table (Type III tests)

Response: LogTRL
Sum Sq Df F value Pr(>F) (Intercept)    236.836 1 5004.3532 <2e-16 ***
Genotype    0.222  4    1.1740 0.3242
Condition   0.050  2    0.5326 0.5881
Genotype:Condition  0.545  8 1.4383 0.1841
Residuals   7.809 165
---
Signif. codes: 0 ‚Äò***‚Äô 0.001 ‚Äò**‚Äô 0.01 ‚Äò*‚Äô 0.05 ‚Äò.‚Äô 0.1
‚Äò ‚Äô 1
mod1 <- lm(LogLRD~Genotype * Condition,
contrasts=list(Genotype=contr.sum, Condition=contr.sum), data=x)
Anova(mod1, type=3) Anova Table (Type III tests)

Response: LogLRD
Sum Sq Df F value Pr(>F) (Intercept)     0.0548 1 1.0562        0.3057
Genotype    0.1060  4 0.5105 0.7281
Condition   1.0774  2 10.3751 5.906e-05 ***
Genotype:Condition 0.3169  8 0.7630     0.6358
Residuals   8.0479 155
---
Signif. codes: 0 ‚Äò***‚Äô 0.001 ‚Äò**‚Äô 0.01 ‚Äò*‚Äô 0.05 ‚Äò.‚Äô 0.1
‚Äò ‚Äô 1
mod1 <- lm(LogLRP~Genotype * Condition,
contrasts=list(Genotype=contr.sum, Condition=contr.sum), data=x)
Anova(mod1, type=3) Anova Table (Type III tests)

Response: LogLRP
Sum Sq Df F value Pr(>F) (Intercept)     6.1749 1 695.6815 < 2.2e-16 ***
Genotype    0.0594  4  1.6732 0.158947
Condition   0.0982  2  5.5326 0.004776 **
Genotype:Condition 0.0899  8  1.2655 0.265395
Residuals   1.3758 155
---
Signif. codes: 0 ‚Äò***‚Äô 0.001 ‚Äò**‚Äô 0.01 ‚Äò*‚Äô 0.05 ‚Äò.‚Äô 0.1
‚Äò ‚Äô 1
```

**Supplementary Data 15.**

Alignments of the DNA binding domains of tomato and arabidopsis transcription factors.

Alignment of ARFDNA binding domains

```
AtARF9    FSKVLTASDTSTHGGFSVLRKHATECLPPLDMTQQTPTQELVAEDVHGYQWKFKHIFRGQ
SlARF9B   FCKVLTASDTSTHGGFSILRKHANECLPPLDMTQATPAQELVAKDLHGFEWHFKHIFRGQ
AtARF18   FVKILTASDTSTHGGFSVLRKHATECLPSLDMTQATPTQELVTRDLHGFEWRFKHIFRGQ
SlARF18   FCKILTASDTSTHGGFSVLRKHANECLPQLDMTQATPTQDLVAKDLHGYEWRFKHIFRGQ
AtARF2    FCKTLTASDTSTHGGFSVLRRHADECLPPLDMSRQPPTQELVAKDLHANEWRFRHIFRGQ

AtARF9    PRRHLLTTGWSTFVTSKRLVAGDTFVFLRGENGELRVGVRRAN
SlARF9B   PRRHLLTTGWSTFVSSKRLVTGDSFVFLRSGKGEVRIGIRRLA
AtARF18   PRRHLLTTGWSTFVSSKRLVAGDAFVFLRGENGDLRVGVRRLA
SlARF18   PRRHLLTTGWSTFVTSKRLVAGDAFVFLRDDSGELRVGVRRLA
AtARF2    PRRHLLQSGWSVFVSSKRLVAGDAFIFLRGENGELRVGVRRAM
```

Alignment DREB26 DNA binding domains

```
AtDREB26 KYKGVRMRSWGSWVSEIRAPNQKTRIWLGSYSTAEAAARAYDVALLCLKGPQA--NLNFP
SlDREB26 KYKGVRMRSWGSWVSEIRAPNQKTRIWLGSYSTPEAAARAYDAALLCLKGPSASSNLNFP
```

Alignment of DNA binding domains

```
AtNLP7_DBD/1-82  ----KKKTEKKRGKTEKTISLDVLQQYFTGSLKDAAKSLGVCPTTMKRICRQHGISRWPS
AtNLP6_DBD/1-86  EAKTVKKSERKRGKTEKTISLEVLQQYFAGSLKDAAKSLGVCPTTMKRICRQHGISRWPS
SlNLP7a_DBD/1-83 ---TGKKSERKRGKAEKTISLEVLQQYFAGSLKDAAKSLGVCPTTMKRICRQHGISRWPS
SlNLP7b_DBD/1-84 --TSGKKSERKRGKAEKTISLEVLQQYFAGSLKDAAKSLGVCPTTMKRICRQHGISRWPS

AtNLP7_DBD/1-82  RKIKKVNRSITKLKRVIESVQGTDGG
AtNLP6_DBD/1-86  RKINKVNRSLTRLKHVIDSVQGADGS
SlNLP7a_DBD/1-83 RKINKVNRSLSKLKRVIESVQGADGT
SlNLP7b_DBD/1-84 RKINKVNRSLSKLKCVIESVQGAEGA
```

**Supplementary Data 16 (a)** qRT-PCR verifies that the expressions of tomato nitrogen-responsive genes SlNIR1 and SlNIR2 are induced by higher nitrogen levels in WT hairy roots. Statistical analysis was performed using one-way ANOVA with Tukey post-hoc test. *p<0.05, ** p<0.01, *** p<0.001; **** p < 0.0001. (**b**)RNAseq data of canonical N regulatory genes in wild type M82 tomato roots, compared to **(c)** empty-vector transformed hairy roots. Asterisks indicate differentially expressed among N conditions by limma (false discovery rate <0.05). *** p-value < 0.001, ** p-value < 0.01,* p-value < 0.05, n.s. = not significant. **(d)** Representative images of tomato hairy roots expressing nuclear localized-GFP driven by the AtNIR1 and AtNRP promoters in 0, 1, and 10 mM KNO3 (first and second row). Third row showing nuclear localized-GFP expression driven by the AtNRP promoter in the roots of stable tomato transformants.

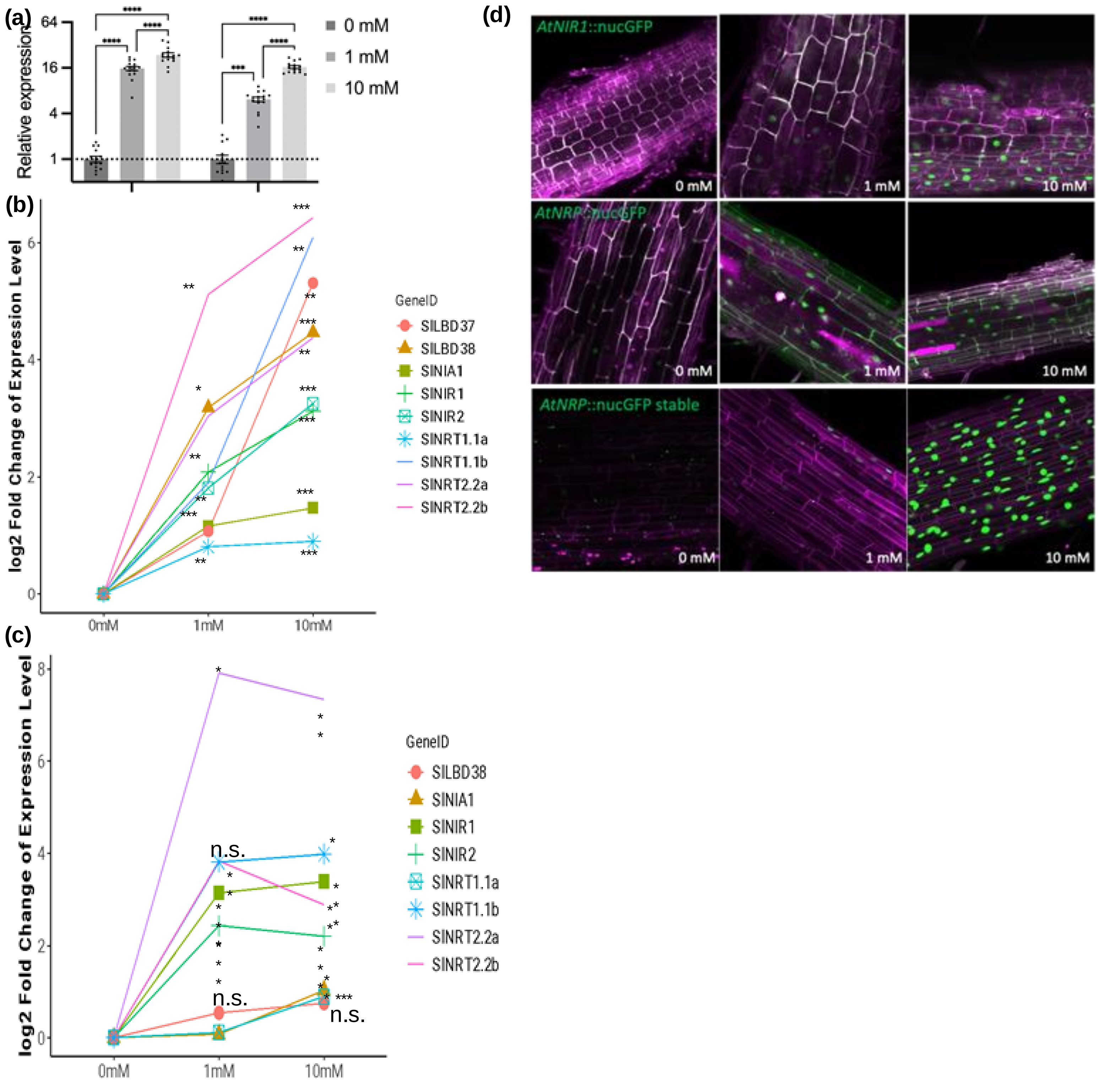

**Supplementary Data 17. Expression levels of tomato genes in modified TARGET assays**. Values represent the mean and standard error of three biological replicates of which each is the mean of two technical replicates. P-values were calculated using an unpaired two-tailed Student's t-test, *P<0.05, ** P<0.01, *** P<0.001, and **** P<0.0001. N = 3.

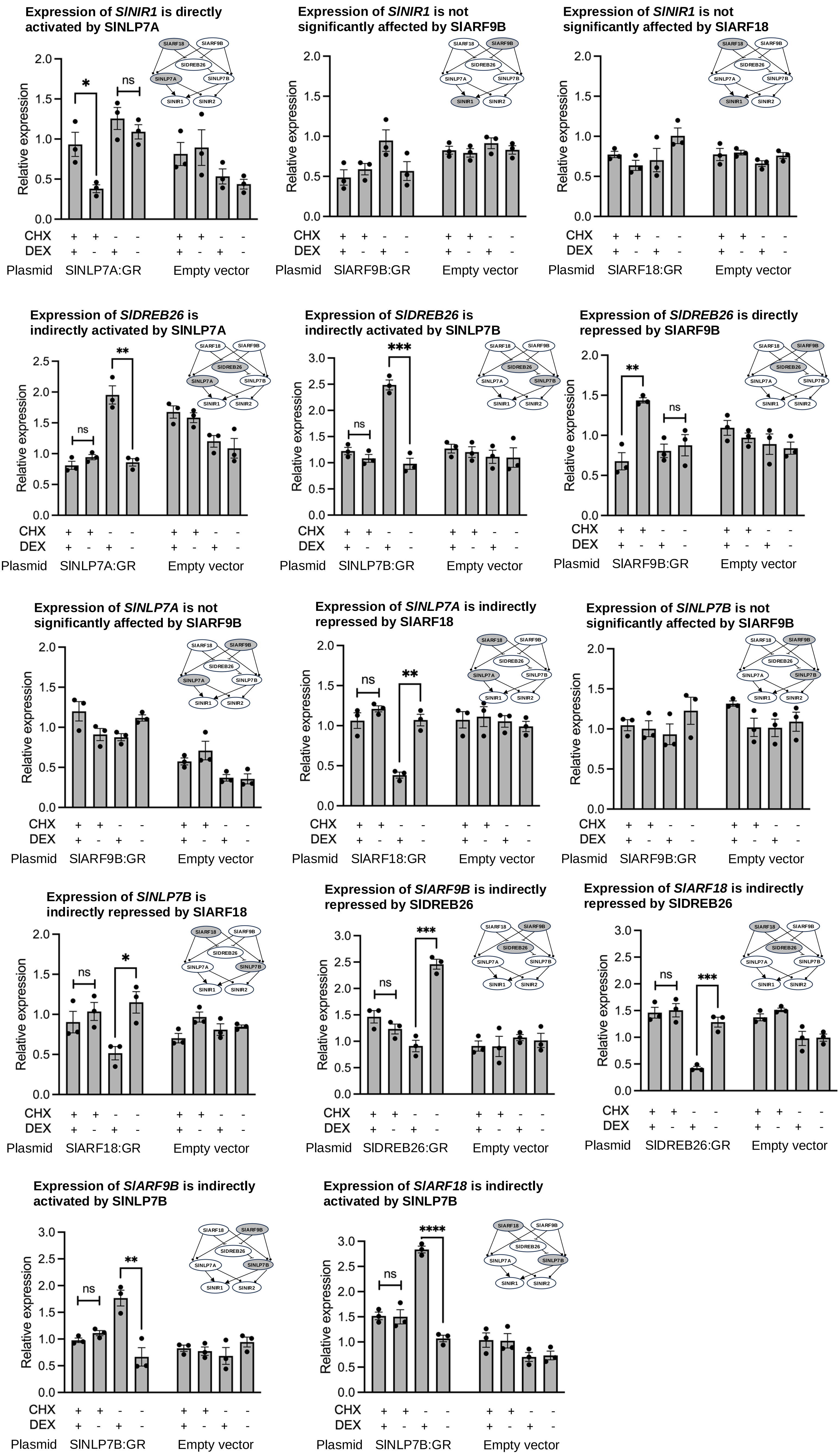

**Supplementary Data 18. Exemplification of PAROT assay. (a)** Schematic of PAROT assay in which protoplasts are transformed with two plasmids encoding (i) synthetic promoter (*AtNRP* or 4xNRE) driving the expression of emerald luciferase (ELUC), and (ii) the constitutive NOS promoter driving expression of red luciferase (RLUC). **(b)** Viability and representative images of Arabidopsis and tomato protoplasts before transfection, and after transfection & after overnight incubation in the PAROT assay. Protoplasts are stained with fluorescein diacetate (FDA) and observed under a fluorescence microscope. Viable protoplasts show fluorescence with an excitation at 470/22 nm and emission at 525/50 nm. Bright-field and fluorescence images are shown on the left and right, respectively. **(d)** The expression levels of endogenous nitrogen-responsive genes after PAROT assay was assessed by qRT-PCR. Values represent the mean and standard error of three biological replicates (independent transfections) of which each is the mean of two technical replicates (qPCR assays). **(e)** Changes in normalized luminescence of *NRP*:ELUC and *4xNRE*:ELUC in response to differing concentrations of nitrate in wild type Arabidopsis and tomato protoplasts (left) and in protoplasts of mutant lines. Values represent the mean and standard error of three biological replicates (independent transfections).

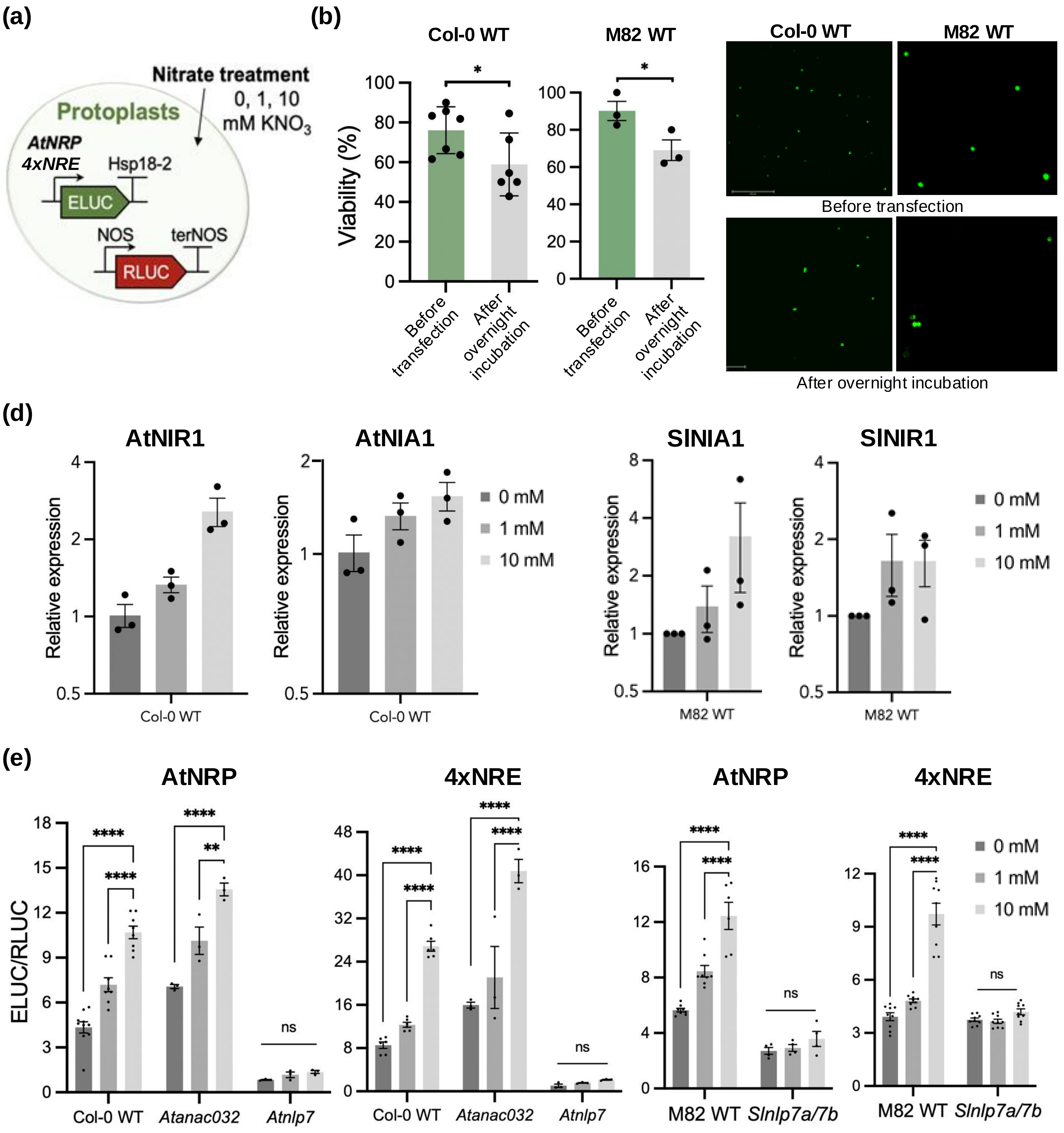

**Supplementary Figure 19. PAROT assay in Arabidopsis and tomato mutant.** Eluc/Rluc response ratio of (**a**) Arabidopsis Col-0 WT, *Atanac032*, *Atarf9b, Atarf18-2, dreb26$_c$, Atnlp6,* and *Atnlp7-1* (**b**) Arabidopsis Col-0 WT, *arf18-2/anac032 , anac032/nlp7-1, dreb26$_c$/nlp7$_c$,* and *anac032/nlp7-1* (**c**) tomato M82 WT, *Slarf9b, Slarf18, Sldreb26, Slnlp7a, and Slnlp7b* (**d**) tomato M82 WT, *Slarf9b/18, Slarf9b/18/dreb26, Slnlp7a/7b, Sldreb26/nlp7a/7b, and Slarf9b/18/dreb26/nlp7a/7b* to 0, 1, 10 mM $KNO_3$. Error bars are standard error; N=4; P-values were calculated using two-way ANOVA with Tukey's multiple comparisons test; *p<0.05, ** p<0.01, *** p<0.001; **** p < 0.0001, ns = not significant.

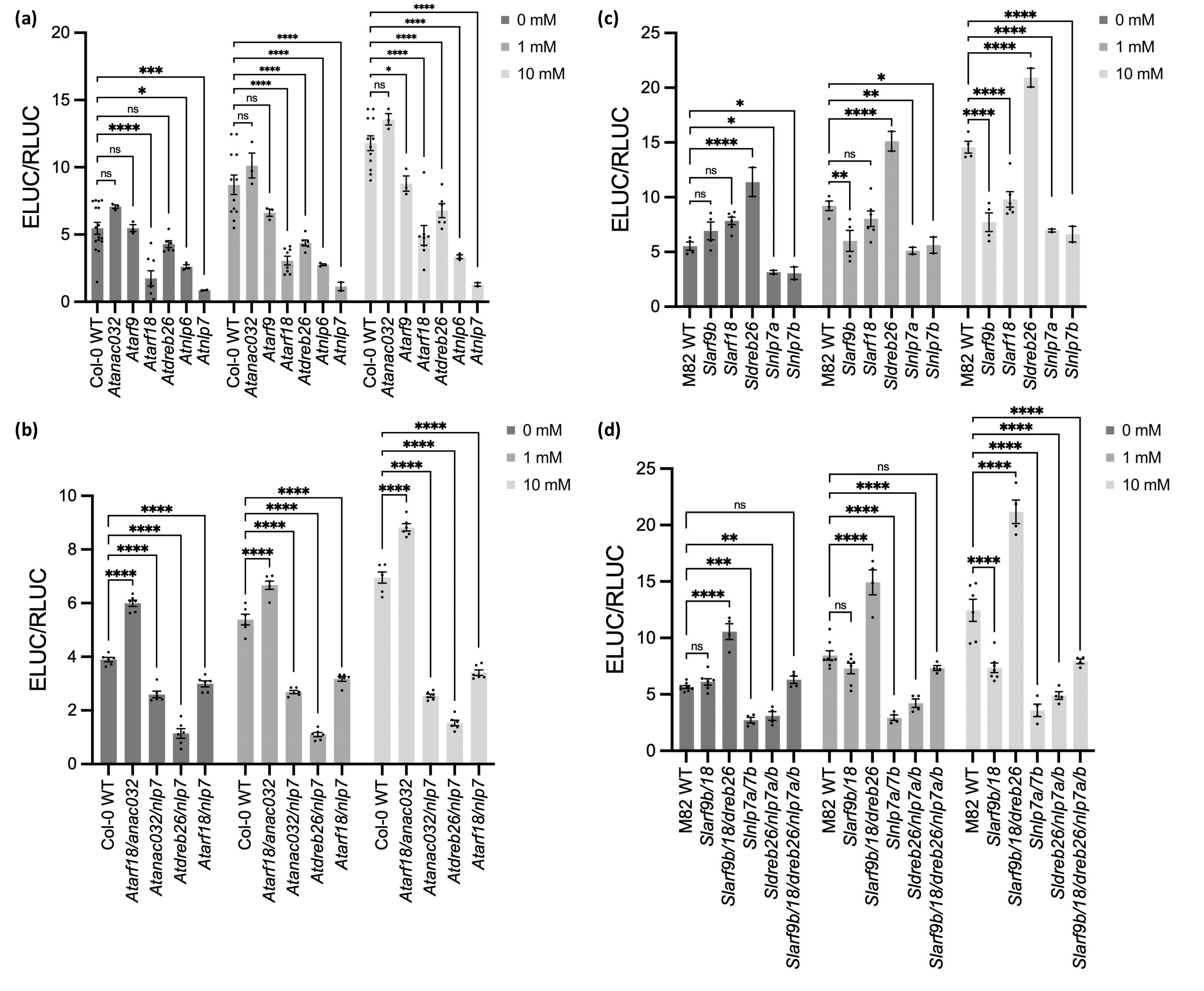

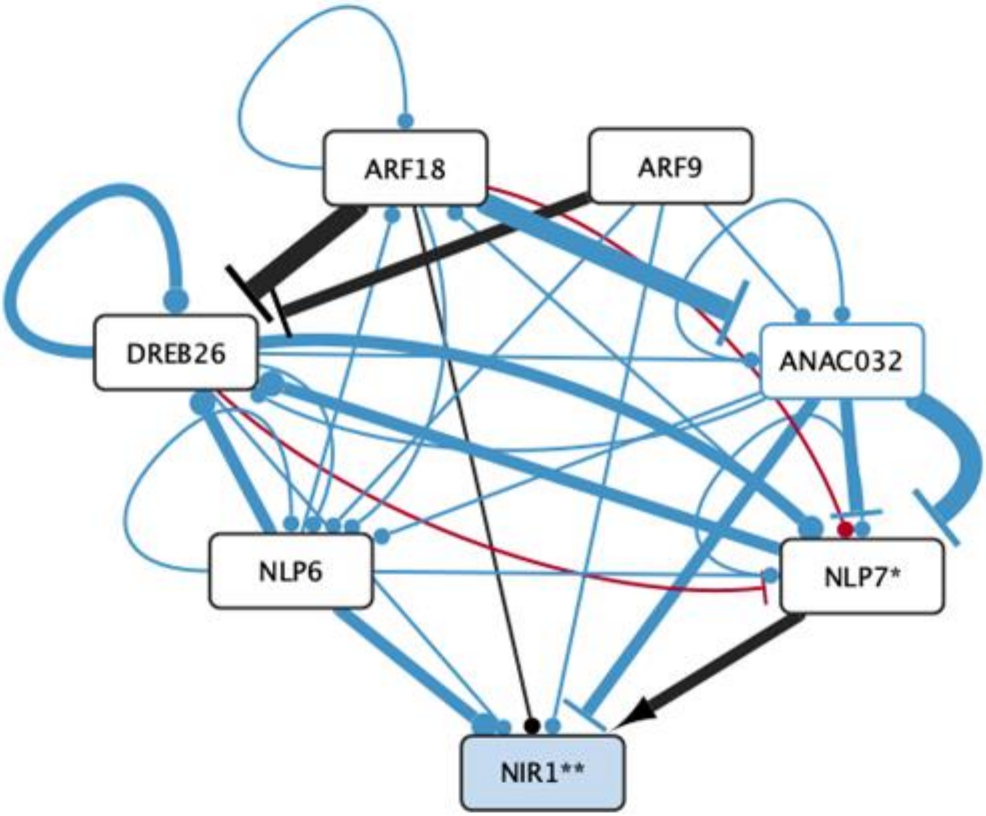

**Supplementary Data 20. A summary of interactions observed in all three assays in Arabidopsis and tomato.** Conserved interactions are indicated with black edges, Arabidopsis-specific interactions in blue, and tomato-specific interactions in red. The thickness of the edge representing the interactions indicates the number of assays in which this interaction was identified, with the thickest edge representing three (yeast one hybrid, in vitro binding assay and TARGET), and the thinnest edge representing one interaction. A perpendicular line or an arrow indicate that the interaction was demonstrated to be direct and regulatory in nature as determined by TARGET. The * in NLP7 indicates that two orthologs of NLP7 in tomato appear to be NLP7 paralogs, and the same for NIR1.

**Supplementary Data 21. Plasmids used in this study**

**Plasmids for expression of transcription factor recombinant proteins used for *in vitro* binding assays**

| Addgene# | Plasmid code | Description | Acceptor | Plasmid type | Source of plasmid |
|---|---|---|---|---|---|
| 196143 | pEPYCeGM0009 | pENTR_AtNLP6_HiBit | pDONR207 | Gateway Entry | This study |
| 196144 | pEPYCeGM0010 | pENTR_AtNLP7_HiBit | pDONR207 | Gateway Entry | This study |
| 196145 | pEPYCeGM0011 | pENTR_AtDREB26_HiBit | pDONR207 | Gateway Entry | This study |
| 196146 | pEPYCeGM0012 | pENTR_AtNAC032_HiBit | pDONR207 | Gateway Entry | This study |
| 196147 | pEPYCeGM0022 | pENTR_AtARF18_HiBit | pDONR207 | Gateway Entry | This study |
| 196150 | pEPYCdKN0009 | pH9GW_9xHis_AtNLP6_HiBit | pH9GW | Gateway Expressio | This study |
| 196151 | pEPYCdKN0010 | pH9GW_9xHis_AtNLP7_HiBit | pH9GW | Gateway Expressio | This study |
| 196152 | pEPYCdKN0011 | pH9GW_9xHis_AtDREB26_HiBit | pH9GW | Gateway Expressio | This study |
| 196153 | pEPYCdKN0012 | pH9GW_9xHis_AtNAC032_HiBit | pH9GW | Gateway Expressio | This study |
| 196154 | pEPYCdKN0022 | pH9GW_9xHis_AtARF18_HiBit | pH9GW | Gateway Expressio | This study |

**Plasmids used for TASRSET assays**

**Arabidopsis**

| Addgene# | Plasmid code | Description | Level 0 Parts | | | | Acceptor | Cloning overhang | | Source of plasmid |
|---|---|---|---|---|---|---|---|---|---|---|
| | | | PROM | CDS | CTAG | 3UTR/TERM | | 5' | 3' | |
| 197549 | pEPOZ1KN0142 | 2xCaMV35s::TMV::AtNLP6:GR::35sT | pICH51288 | pEPSW0CM0073 | pEPOZ0CM0137 | pICH41414 | pCK2 (Addgene 136696) | GCA | TAC | This study |
| 197550 | pEPOZ1KN0143 | 2xCaMV35s::TMV::AtNLP7:GR::35sT | pICH51288 | pEPSW0CM0074 | pEPOZ0CM0137 | pICH41414 | pCK2 (Addgene 136696) | GCA | TAC | This study |
| 197551 | pEPOZ1KN0144 | 2xCaMV35s::TMV::AtDREB26:GR::35sT | pICH51288 | pEPSW0CM0075 | pEPOZ0CM0137 | pICH41414 | pCK2 (Addgene 136696) | GCA | TAC | This study |
| 197552 | pEPOZ1KN0145 | 2xCaMV35s::TMV::AtANAC032:GR::35s | pICH51288 | pEPSW0CM0076 | pEPOZ0CM0137 | pICH41414 | pCK2 (Addgene 136696) | GCA | TAC | This study |
| 197554 | pEPOZ1KN0147 | 2xCaMV35s::TMV::AtARF18:GR::35sT | pICH51288 | pEPOZ0CM0138 | pEPOZ0CM0137 | pICH41414 | pCK2 (Addgene 136696) | GCA | TAC | This study |
| 197555 | pEPOZ1KN0148 | 2xCaMV35s::TMV::AtARF9:GR::35sT | pICH51288 | pEPOZ0CM0139 | pEPOZ0CM0137 | pICH41414 | pCK2 (Addgene 136696) | GCA | TAC | This study |

**Tomato**

| Addgene# | Plasmid code | Description | Acceptor | Plasmid type | Source of |
|---|---|---|---|---|---|
| | pGD0001 | pBEACON_SlNLP7A_GR | pBEACON | Gateway Expressio | This study |
| | pGD0002 | pBEACON_SlNLP7B_GR | pBEACON | Gateway Expressio | This study |
| | pGD0003 | pBEACON_SlARF9B_GR | pBEACON | Gateway Expressio | This study |
| | pGD0004 | pBEACON_SlARF18_GR | pBEACON | Gateway Expressio | This study |
| | pGD0005 | pBEACON_DREB26_GR | pBEACON | Gateway Expressio | This study |
| | pENTR-D-SlNLP7A-CDS | SlNLP7A-CDS sequence in cloning | pENTRY | Gateway Entry | This study |
| | pBEACON-5C-SlNLP7A-CDS | 35S::GR-SlNLP7A-CDS, correct | pBEACON | Gateway Expressio | This study |
| | pUC57-SlARF18-CDS | SlARF18-CDS sequence in cloning | pUC57 | Gateway Entry | This study |
| | pBEACON-5C-SlARF18-CDS | 35S::GR-SlARF18-CDS | pBEACON | Gateway Expressio | This study |
| | pENTR-D-SlNLP7B-CDS | SlNLP7B-CDS in cloning vector | pENTRY | Gateway Entry | This study |
| | pBEACON-5C-SlNLP7B-CDS | 35S::GR-SlNLP7B-CDS | pBEACON | Gateway Expressio | This study |

**Plasmids used for protoplast co-expression luciferase assays**

| Addgene# | Plasmid code | Description | Level 0 Parts | | | | | | Cloning overhang | | Source of plasmid |
|---|---|---|---|---|---|---|---|---|---|---|---|
| | | | PROM | 5UTR | CDS | CTAG | 3UTR/TERM | Acceptor | 5' | 3' | |
| 154629 | pEPYC1CB0003 | AtuNos::TMV::LucF::FLAG::nosT | pICH42211 | pICH41402 | pEPAS0CM0008 | pICSL50007 | pICH41421 | pICH47732 (Addgene 48000 | TGCC | GCAA | Cat et al., 2020 |
| 197536 | pEPSW1KN0070 | CaMV35s::TMV::LucN::FLAG::35sT | pICH51277 | - | pEPYC0CM0133 | pICSL50007 | pICH41414 | pCK1 (Addgene 136695) | ATG | GCA | This study |
| 196178 | pEPSW1KN0034 | CaMV35s::TMV::LucF::FLAG::35sT | pICH51277 | - | pEPAS0CM0008 | pICSL50007 | pICH41414 | pCK1 (Addgene 136695) | ATG | GCA | This study |
| 196177 | pEPSW1KN0035 | AtuNos::TMV::LucN::FLAG::nosT | pICH42211 | pICH41402 | pEPYC0CM0133 | pICSL50007 | pICH41421 | pCK1 (Addgene 136695) | ATG | GCA | This study |
| 196174 | pEPSW1KN0014 | AtANAC32::LucN::FLAG::nosT | pEPSW0CM0014 | - | pEPYC0CM0133 | pICSL50007 | pICH41421 | pCK1 (Addgene 136695) | ATG | GCA | This study |
| 197537 | pEPSW1KN0016 | ANR1::LucN::FLAG::nosT | pEPSW0CM0016 | - | pEPYC0CM0133 | pICSL50007 | pICH41421 | pCK1 (Addgene 136695) | ATG | GCA | This study |
| 196176 | pEPSW1KN0018 | ARF18::LucN::FLAG::nosT | pEPSW0CM0018 | - | pEPYC0CM0133 | pICSL50007 | pICH41421 | pCK1 (Addgene 136695) | ATG | GCA | This study |
| 196175 | pEPSW1KN0020 | NLP6::LucN::FLAG::nosT | pEPSW0CM0020 | - | pEPYC0CM0133 | pICSL50007 | pICH41421 | pCK1 (Addgene 136695) | ATG | GCA | This study |
| 197538 | pEPSW1KN0022 | DREB26::LucN::FLAG::nosT | pEPSW0CM0022 | - | pEPYC0CM0133 | pICSL50007 | pICH41421 | pCK1 (Addgene 136695) | ATG | GCA | This study |
| 197539 | pEPSW1KN0024 | NLP7::LucN::FLAG::nosT | pEPSW0CM0024 | - | pEPYC0CM0133 | pICSL50007 | pICH41421 | pCK1 (Addgene 136695) | ATG | GCA | This study |
| 197540 | pEPSW1KN0025 | NIR1::LucN::FLAG::nosT | pEPSW0CM0025 | - | pEPYC0CM0133 | pICSL50007 | pICH41421 | pCK1 (Addgene 136695) | ATG | GCA | This study |
| 197541 | pEPSW1KN0027 | CaMV35s::TMV::NLP6::35sT | pICH51277 | - | pEPSW0CM0027 | - | pICH41414 | pCK2 (Addgene 136696) | GCA | TAC | This study |
| 197542 | pEPSW1KN0029 | CaMV35s::TMV::NLP7::35sT | pICH51277 | - | pEPSW0CM0029 | - | pICH41414 | pCK2 (Addgene 136696) | GCA | TAC | This study |
| 197543 | pEPSW1KN0030 | CaMV35s::TMV::DREB26::35sT | pICH51277 | - | pEPSW0CM0030 | - | pICH41414 | pCK2 (Addgene 136696) | GCA | TAC | This study |
| 197544 | pEPSW1KN0031 | CaMV35s::TMV::ANAC032::35sT | pICH51277 | - | pEPSW0CM0031 | - | pICH41414 | pCK2 (Addgene 136696) | GCA | TAC | This study |
| 197545 | pEPSW1KN0032 | CaMV35s::TMV::ARF18::35sT | pICH51277 | - | pEPSW0CM0032 | - | pICH41414 | pCK2 (Addgene 136696) | GCA | TAC | This study |
| 197547 | pEPSW1KN0113 | CaMV35s::TMV::ARF9::35sT | pICH51277 | - | pEPSW0CM0112 | - | pICH41414 | pCK2 (Addgene 136696) | GCA | TAC | This study |
| 196179 | pEPOR1CB0068 | CaMV35s::TMV::YFP:NLS::35sT | pICH51277 | - | pEPOR0CM0010 | - | pICH41414 | pICH47761 (Addgene 48003 | TTAC | CAGA | This study |

**Plasmids used for nitrate reporter assay**

| Addgene# | Plasmid code | Description | Acceptor | Plasmid type | Source of |
|---|---|---|---|---|---|
| | pENTR5'-NRPpro | NRP promoter in cloning vector | pENTRY | Gateway Entry | This study |
| | pMR105-NRPpro | NRP promoter reporter | pMR105 | Gateway Expressio | This study |
| | pENTR5'-AtNIR1pro | AtNIR1 promoter in cloning vector | pENTRY | Gateway Entry | This study |
| | pMR105-AtNIR1pro | AtNIR1 promoter reporter | pMR105 | Gateway Expressio | This study |

**Plasmids used for PAROT assays**

| Addgene# | Plasmid code | Description | Acceptor | Plasmid type | Source of |
|---|---|---|---|---|---|
| 170887 | pGD0006 | pDGB2a_pNOS_RedF_NOS | pDGB2a | Expression | Gonzalez- |
| | pGD0007 | pDGBa1_NRP_ELuc_THsp18-2 | pDGBa1 | Expression | This study |

**Plasmids used to create CRISPR knockout lines**

**Arabidopsis**

| Plasmid code | Description | Cloning overhang | | Source of plasmid |
|---|---|---|---|---|
| | | 5' | 3' | |
| pEPOZ1KN0004 | AtuNOSpro::TMV::NPTII_domesticated(pEPOZ0CM0001)::AtuOCSter (backbone pCk4) | CAG (SapI) | GGT (SapI) | This study |
| pEPOZ1KN0005 | [illegible] expression cassette for SpCas9, AtYAOpro::SpCas9::RBCS-E9ter (reverse orientation) | GCA (SapI) | TAC (SapI) | This study |
| pEPOZ1KN0008 | [illegible] expression cassette for SaCas9, AtRPS5Apro::SaCas9::RBCS-E9ter (reverse orientation) | GCA (SapI) | TAC (SapI) | This study |
| pEPOZ3KN0033 | SpCas9, KanR, and 8 guides (2 guides per gene, 4 genes, ARF18, DREB26, NLP7, ANAC032) | GCA | TAC | This study |

| | | | | |
|---|---|---|---|---|
| pEPOZ3KN0103 | Level 3 contains FASTred, SaCas9, KanR, and 10 guides (2 guides per gene, 5 genes, ARF18, DREB26, NLP7, | GCA | TAC | This study |
| **Tomato** | | | | |
| **Plasmid code** | **Description** | **Cloning overhang** | | **Source of plasmid** |
| | | **5'** | **3'** | |
| pUAP4-AtHSP18.2 ter | L0 plasmid of AtHSP18.2 | GCTT | CGCT | This study |
| pCK4-35S-BAR-NOS | L1 plasmid of Basta resistence | CAG | GGT | This study |
| pCK2R-proRPS5A-SpCas9-HSP18 | L1 plasmid of Cas9 expression | GCA | TAC | This study |
| pCsA-SlNLP7A-NPTII | Binary plasmid of CRISPR-SlNLP7A, includes FASTgreen, SpCas9, 2 gRNAs, Kanamycin selection | GGAG | TACT | This study |
| pCsB-SlNLP7B-NPTII | Binary plasmid of CRISPR-SlNLP7B, includes FASTgreen, SpCas9, 2 gRNAs, Kanamycin selection | TACT | AATG | This study |
| pCsA-SlARF18-NPTII | Binary plasmid of CRISPR-SlARF18, includes FASTgreen, SpCas9, 2 gRNAs, Kanamycin selection | GGAG | TACT | This study |
| pCsB-SlARF9B-NPTII | Binary plasmid of CRISPR-SlARF9B, includes FASTgreen, SpCas9, 2 gRNAs, Kanamycin selection | TACT | AATG | This study |
| pCSA-SlANR1-NPTII | Binary plasmid of CRISPR-SlANR1, includes FASTgreen, SpCas9, 2 gRNAs, Kanamycin selection | GGAG | TACT | This study |
| PCsA-DREB26-NPTII | Binary plasmid of CRISPR-SlDREB26, includes FASTgreen, SpCas9, 2 gRNAs, Kanamycin selection | GGAG | TACT | This study |
| PCsA-ANR1-NPTII | Binary plasmid of CRISPR-SlANR1, includes FASTgreen, SpCas9, 2 gRNAs, Kanamycin selection | GGAG | TACT | This study |
| pCsA-ARF-NPTII | Binary plasmid of CRISPR-SlARF18-SlARF9B, includes FASTgreen, SpCas9, 2 gRNAs each gene, Kanamycin selection | GGAG | TACT | This study |
| pCsA-ARF-BASTA | Binary plasmid of CRISPR-SlARF18-SlARF9B, includes FASTgreen, SpCas9, 2 gRNAs each gene, Basta selection | GGAG | TACT | This study |
| pCsA-NLP-BASTA | Binary plasmid of CRISPR-SlNLP7A-SlNLP7B, includes FASTgreen, SpCas9, 2 gRNAs each gene, Basta selection | GGAG | TACT | This study |
| pCsA-NLP-NPTII | Binary plasmid of CRISPR-SlNLP7A-SlNLP7B, includes FASTgreen, SpCas9, 2 gRNAs each gene, Kanamycin selection | GGAG | TACT | This study |
| PCSA-SlANR1-SlDREB26-NPTII | Binary plasmid of CRISPR-SlDREB26-SlANR1, includes FASTgreen, SpCas9, 2 gRNAs each gene, Kanamycin selection | GGAG | TACT | This study |
| pCSA-SlARF9B-DREB26-ANR1-NPTII | Binary plasmid of CRISPR-SlARF9B-SlDREB26-SlANR1, includes FASTgreen, SpCas9, 2 gRNAs each gene, Kanamycin selection | GGAG | TACT | This study |
| pCSA-SlARF9B-DREB26-ANR1-BAR | Binary plasmid of CRISPR-SlARF9B-SlDREB26-SlANR1, includes FASTgreen, SpCas9, 2 gRNAs each gene, Basta selection | GGAG | TACT | This study |
| pCSA-SlARF18-DREB26-ANR1-NPTII | Binary plasmid of CRISPR-SlARF18-SlDREB26-SlANR1, includes FASTgreen, SpCas9, 2 gRNAs each gene, Kanamycin selection | GGAG | TACT | This study |
| pCSA-SlARF18-DREB26-ANR1-BAR | Binary plasmid of CRISPR-SlARF18-SlDREB26-SlANR1, includes FASTgreen, SpCas9, 2 gRNAs each gene, Basta selection | GGAG | TACT | This study |

| | | | | |
|---|---|---|---|---|
| PCSB-SINLP7A-7B-SIDREB26-NPTII | Binary plasmid of CRISPR-SIDREB26-SINLP7A-7B, includes FASTgreen, SpCas9, 2 gRNAs each gene, Kanamycin selection | TACT | AATG | This study |
| PCSA-SIARF18-9B-SIDREB26-NPTII | Binary plasmid of CRISPR-SIARF18-9B-SIDREB26, includes FASTgreen, SpCas9, 2 gRNAs each gene, Kanamycin selection | GGAG | TACT | This study |
| PCSA-SIARF18-9B-SINLP7A-7B-NPTII | Binary plasmid of CRISPR-SIARF18-9B-SINLP7A-7B, includes FASTgreen, SpCas9, 2 gRNAs each gene, Kanamycin selection | GGAG | TACT | This study |
| pCSA-SIARF18-SIARF9B-DREB26-ANR1- | Binary plasmid of CRISPR-SIARF18-9B-SIDREB26-SIANR1, includes FASTgreen, SpCas9, 2 gRNAs each gene, Kanamycin selection | GGAG | TACT | This study |
| pCSA-SLARF18-SIARF9B-DREB26-ANR1 | Binary plasmid of CRISPR-SIARF18-9B-SIDREB26-SIANR1, includes FASTgreen, SpCas9, 2 gRNAs each gene, Basta selection | GGAG | TACT | This study |
| pCSA-SIARF9B-DREB26-NLP7A-7B-NPTI | Binary plasmid of CRISPR-SIARF9B-SIDREB26-SINLP7A-7B, includes FASTgreen, SpCas9, 2 gRNAs each gene, Kanamycin selection | GGAG | TACT | This study |
| pCSA-SIARF9B-DREB26-NLP7A-7B-BAR | Binary plasmid of CRISPR-SIARF9B-SIDREB26-SINLP7A-7B, includes FASTgreen, SpCas9, 2 gRNAs each gene, Basta selection | GGAG | TACT | This study |
| pCSA-SIARF18-DREB26-NLP7A-7B-NPTI | Binary plasmid of CRISPR-SIARF18-SIDREB26-SINLP7A-7B, includes FASTgreen, SpCas9, 2 gRNAs each gene, Kanamycin selection | GGAG | TACT | This study |
| pCSA-SIARF18-DREB26-NLP7A-7B-BAR | Binary plasmid of CRISPR-SIARF18-SIDREB26-SINLP7A-7B, includes FASTgreen, SpCas9, 2 gRNAs each gene, Basta selection | GGAG | TACT | This study |
| pCSA-SIARF18-9B-DREB26-NLP7A-7B-N | Binary plasmid of CRISPR-SIARF18-9B-SIDREB26-SINLP7A-7B, includes FASTgreen, SpCas9, 2 gRNAs each gene, Kanamycin selection | GGAG | TACT | This study |
| pCSA-SIARF18-9B-DREB26-NLP7A-7B-B | Binary plasmid of CRISPR-SIARF18-9B-SIDREB26-SINLP7A-7B, includes FASTgreen, SpCas9, 2 gRNAs each gene, Basta selection | GGAG | TACT | This study |

**Level 0 Phytobricks used in the assemblies above**

| Addgene# | Plasmid code | Part type | Description | Compatibility with Assembly Systems | Cloning overhang (top strand) 5' | 3' | Source of plasmid |
|---|---|---|---|---|---|---|---|
| | pEPOZ0CM0001 | CDS | NPTII CDS (Kanamycin resistance | MoClo, Loop GB | AATG | GCTT | This study |
| | pEPOZ0CM0039 | PROM+5UTR | NRP promoter, 364 bp, Level 0, nitrogen responsive promoter | MoClo, Loop GB | GGAG | AATG | This study |
| 196162 | pEPSW0CM0014 | PROM+5UTR | AtANAC032 (AT1G77450) 1000 bp upstream of TSS (1 SNP to remove BsaI) | MoClo, Loop GB | GGAG | TACT | This study |
| 196164 | pEPSW0CM0018 | PROM+5UTR | AtARF18 (AT3G61830) 1000 bp upstream of TSS | MoClo, Loop GB | GGAG | TACT | This study |
| 196163 | pEPSW0CM0020 | PROM+5UTR | AtNLP6 (AT1G64530) 1000 bp upstream of TSS | MoClo, Loop GB | GGAG | TACT | This study |
| 197517 | pEPSW0CM0022 | PROM+5UTR | AtDREB26 (AT1G21910) 1000 bp upstream of TSS | MoClo, Loop GB | GGAG | TACT | This study |
| 197518 | pEPSW0CM0024 | PROM+5UTR | AtNLP7 (AT4G24020) 1000 bp upstream of TSS | MoClo, Loop GB | GGAG | TACT | This study |
| 197519 | pEPSW0CM0025 | PROM+5UTR | AtNIR1 (AT2G15620) 936 bp upstream of TSS | MoClo, Loop GB | GGAG | TACT | This study |
| 197520 | pEPSW0CM0027 | CDS | AtNLP6 | MoClo, Loop GB | AATG | GCTT | This study |
| 197521 | pEPSW0CM0029 | CDS | AtNLP7 | MoClo, Loop GB | AATG | GCTT | This study |
| 197522 | pEPSW0CM0030 | CDS | AtDREB26 | MoClo, Loop GB | AATG | GCTT | This study |
| 197523 | pEPSW0CM0031 | CDS | AtANAC032 | MoClo, Loop GB | AATG | GCTT | This study |

| | | | | | | | |
|---|---|---|---|---|---|---|---|
| 197524 | pEPSW0CM0032 | CDS | AtARF18 | MoClo, Loop GB | AATG | GCTT | This study |
| 197526 | pEPSW0CM0112 | CDS | AtARF9 | MoClo, Loop GB | AATG | GCTT | This study |
| 197528 | pEPSW0CM0073 | CDS | NLP6 (no stop codon) | MoClo, Loop GB | AATG | TTCG | This study |
| 197529 | pEPSW0CM0074 | CDS | NLP7 (no stop codon) | MoClo, Loop GB | AATG | TTCG | This study |
| 197530 | pEPSW0CM0075 | CDS | DREB26 (no stop codon) | MoClo, Loop GB | AATG | TTCG | This study |
| 197531 | pEPSW0CM0076 | CDS | ANAC032 (no stop codon) | MoClo, Loop GB | AATG | TTCG | This study |
| 197533 | pEPOZ0CM0138 | CDS | ARF18 (no stop codon) | MoClo, Loop GB | AATG | TTCG | This study |
| 197534 | pEPOZ0CM0139 | CDS | ARF9 (no stop codon) | MoClo, Loop GB | AATG | TTCG | This study |
| 197535 | pEPOZ0CM0137 | CTAG | C-terminal glucocorticoid receptor, GR | MoClo, Loop GB | TTCG | GCTT | This study |
| 50268 | pICH51277 | PROM+5UTR | CaMV35s_TMV | MoClo, Loop GB | GGAG | AATG | Engler et al 2014 |
| 50255 | pICH42211 | PROM | NOS promoter (Agrobacterium tumefac | MoClo, Loop GB | GGAG | TACT | Engler et al 2014 |
| 50271 | pICH87633 | PROM+5UTR | AtuNos_TMV | MoClo, Loop GB | GGAG | AATG | Engler et al 2014 |
| 50269 | pICH51288 | PROM+5UTR | 2xCaMV35s_TMV | MoClo, Loop GB | GGAG | AATG | Engler et al 2014 |
| 50285 | pICH41402 | 5UTR | TMVΩ (Tobacco Mosaic Virus) | MoClo, Loop GB | TACT | AATG | Engler et al 2014 |
| 154595 | pEPYC0CM0133 | CDS | LucN (NanoLuc) | MoClo, Loop GB | AATG | TTCG | Cai et al 2020 |
| 154594 | pEPAS0CM0008 | CDS | LucF (Firefly luciferase) | MoClo, Loop GB | AATG | TTCG | Cai et al 2020 |
| 50308 | pICSL50007 | CTAG | C terminal FLAG tag | MoClo, Loop GB | TTCG | GCTT | Engler et al 2014 |
| 50337 | pICH41414 | 3UTR_TERM | 35S terminator (Cauliflower Mosaic Vir | MoClo, Loop GB | GCTT | CGCT | Engler et al 2014 |
| 50339 | pICH41421 | 3UTR_TERM | NOS terminator (Agrobacterium tumefa | MoClo, Loop GB | GCTT | CGCT | Engler et al 2014 |
| 50343 | pICH41432 | 3UTR_TERM | OCS terminator (Agrobacterium tumefa | MoClo, Loop GB | GCTT | CGCT | Engler et al 2014 |

**Supplementary Data 22.** Primers used in this study

notes

Primers used for qPCR in TARSET assays

| Gene | Forward primer (5' - 3') | Reverse primer (5' - 3') |
|---|---|---|
| AtNIR1 | GGCTTATCGACGAACTTGGTG | AGTCTCTCCTCTCCCACTGTT |
| AtNLP7 | TGACCTGATAATTCATTCGAACTCT | AATTTGTCCAAAGAAACCCTAAACA |
| AtNLP6 | TGGATCTGAGAGTTTTGTTTCT | TGATGTGGAGAAGTGTGCTT |
| eEF-1α | AGATCAACGAGCCCAAGA | CCGTTCCAATACCACCAAT |
| AtANAC032 | TCTGTCGTAAATGCGCGTCG | ACAAAGCCATGTCGGGAAGG |
| AtDREB26 | ACCCTTTTGTCCCCCAAATCC | GAGACGGCTGATGAAGTAGGG |
| AtARF18 | CGACGTCGTTTTAGTGCGTA | TCGCCAACTTGAAAAATTGACAT |
| AtARF9 | CTGTGTTTTGTTGCGCATGG | AACCCTCTCCCCATAAACCC |
| SlDREB26 | AATGAGAAGCTGGGGATCATGG | TGCAGCATCATAGGCTCTAGC |
| SlNLP7A | TCGTGGAACTTTGGCATTGC | TCACTGCCTCAAGTGCTTTG |
| SlNLP7B | GATTTTTGCTGCTGCTGCTG | AGCACCGGAAAGTCCATTTC |
| SlARF9B | ACCATTTGTGGCATCAGTCG | AAATTGCGCTGAGGGAATCC |
| SlARF18 | ATGGAAGGGGTTTGTGTTGG | TTGTTTCGAGGGCAAAGCTC |
| SlANR1 | ATGTCCGCTGATGACACTGG | ACAAACGCTGCACCATTGTC |
| SlEXP | GCTAAGAACGCTGGACCTAATG | TGGGTGTGCCTTTCTGAATG |

Primers used for qPCR in expression assays

| Gene | Forward primer (5' - 3') | Reverse primer (5' - 3') |
|---|---|---|
| SlEXP | GCTAAGAACGCTGGACCTAATG | TGGGTGTGCCTTTCTGAATG |
| SlNIR1 | TGATGTGGTTGATCGATGAACTGGG | TAGCTCTCCTGAGCCGTACTC |
| SlNIR2 | GCTGATGATGTAGTCCCTGTTTGC | GGCATTCTCTTCACAACTTCTGC |
| AtNIA1 | AGGCTACGCTTATTCTGGAGG | TGTTCGGTTTCTCCTGGTGG |

Primers used for nitrate-responsive reporter cloning

| Gene | Forward primer (5' - 3') | Reverse primer (5' - 3') |
|---|---|---|
| AtNIR1 | CCT TTT TTC CAA ACA CGC ACT AAA CC | GAT GAT GGC GGA AGA AGG AGT TG |

Primers used for amplification of coding regions for cloning into pENTR™ /D-TOPO™

| Gene | Forward primer (5' - 3') | Reverse primer (5' - 3') |
|---|---|---|
| SlDREB26 | CACCATGGTGAAGACAGAGCAAA | TTAGCAAAAACTCCATAAAGGTA |
| SlNLP7B | CACCATGTCGGAACCGGGAGGA | TCATTTACCTGAACTCTCACAGGA |
| SlARF9B | CACCATGGAGAATCAAGGGTCTTT | TCATGTTGAAGCAGTTGTCT |

Primers used for genotyping Arabidopsis T-DNA insertional mutant lines

| Gene | Reaction | Forward primer (5' - 3') | Reverse primer (5' - 3') |
|---|---|---|---|
| AtNLP7 | Wild type reaction | GTTTTTCTTTAGACCGCCACC | AAGAATCAACCGAACAACACG |
| AtNLP7 | T-DNA reaction | ATTTTGCCGATTTCGGAAC | AAGAATCAACCGAACAACACG |
| AtARF18 | Wild type reaction | TCGGAAGTTCTTCGTCAAGTG | TCACCCCTGAAAAACACAAAG |
| AtARF18 | T-DNA reaction | ATATTGACCATCATACTCATTGC | TCACCCCTGAAAAACACAAAG |
| AtANAC032 | Wild type reaction | ACCAACAATTGTGGAAGCAAG | CTCTCCATTTGGAGGTTTTCC |
| AtANAC032 | T-DNA reaction | ATTTTGCCGATTTCGGAAC | CTCTCCATTTGGAGGTTTTCC |

Primers used for genotyping Arabidopsis CRISPR edited lines

| Gene | Forward primer (5' - 3') | Reverse primer (5' - 3') |
|---|---|---|
| AtDREB26 | TCACTCTACAAAGCCTGTACG | TCCGAATTGTACCGAGTTTGC |
| AtANAC032 | TGCTTCCTGAAAATAACAACAC | AAACAACAATAAAGGGGAAAATAGG |
| AtARF18 | CAACACGGAATCTGAAAGGAAAGG | TAAACCTCATCTGTTTCATGCTCC |
| AtNLP7 | TGATGAGCCTAGTCCCACCAGA | GCCCCAGACATGAGCATCCACG |

Primers used for genotyping of CRISPR lines in tomato hairy roots and stable lines

| | Gene | Forward primer (5' - 3') | Reverse primer (5' - 3') |
|---|---|---|---|
| For Illumina sequencing | SlNLP7A | TCCTCTGTCACGGAAGCG TCC GAG CCG GAA GAA GAA ATG | TTTAGCCTCCCCACCGAC GTA GAG AGA GCA TTT CCG TTA GGC |
| | SlNLP7B | TCCTCTGTCACGGAAGCG GAA CAA AAA TGT CGG AAC CGG G | TTTAGCCTCCCCACCGAC GGC TTA CAT ACA AGC TAC AAA TCT AGG |
| | SlDREB26 | TCCTCTGTCACGGAAGCG CAA TGT CAA TAT CAT CAA TAG CAA CAA GC | TTTAGCCTCCCCACCGAC GGA GAG AGA GTT GTA CAA GTG TGA TG |
| | SlARF18 | TCCTCTGTCACGGAAGCG GTA TGG GAA CTG AGG ATT TGT ATA G | TTTAGCCTCCCCACCGAC GCA CAC AAA CTA AAA GCA ATT ACT ATA G |
| | SlARF9B | TCCTCTGTCACGGAAGCG GAA TTT GTT GTG GAA AAA GTT TCA GGT G | TTTAGCCTCCCCACCGAC CTT AAC AAA GGC GAT CCC GTA TC |
| | SlANR1 | TCCTCTGTCACGGAAGCG CTT GCT TCT TGG TTA GGG TTT CTC C | TTTAGCCTCCCCACCGAC GCA TTT TCT CCT TTG ACT AGG TGA CC |
| For Sanger sequencing | SlNLP7A | CAA GTC TCA ATG GAA TCA CAC TCC TC | GTA TAA GCC CGA AGC TTA CAA GTG AC |
| | SlNLP7B | GCA AGT GGC AAC TTA ATA AGA TTC C | GCT TTC CAT TGT TTC TGT TTC TGC |
| | SlDREB26 | ATG GTG AAG ACA GAG CAA AAA AGT CTA TC | GCA AAA ACT CCA TAA AGG TAT ATC TGC TG |
| | SlARF18 | AGT TCT GTA ACA ACG ACT GCT GAG TTC | AAC AAA GTG AAA GAA CTA CCA GTT GCG |
| | SlARF9B | GTC AAA CCA GCA GCA TCA GCA AC | GAA TGA GAG GTG GAA CTA ACA ATG GTG G |
| | SlANR1 | ATG TCC GCT GAT GAC ACT GGA TC | CTT CAG CAT TAC AGT CTG TCC GTA G |
